# There is more than chitin synthase in insect resistance to benzoylureas: Molecular markers associated with teflubenzuron resistance in *Spodoptera frugiperda*

**DOI:** 10.1101/2020.11.11.376582

**Authors:** Antonio Rogério Bezerra do Nascimento, Vitor Antonio Corrêa Pavinato, Juliana Gonzales Rodrigues, Karina Lucas Silva-Brandão, Fernando Luis Consoli, Andrew Michel, Celso Omoto

## Abstract

Chitin synthesis inhibitors are successfully used in pest control and are an excellent option for use in integrated pest management programs due to their low non-target effects. Reports on field-evolved resistance of lepidopteran pests to chitin synthesis inhibitors and the selection of laboratory resistant strains to these products require a detailed investigation on the resistance mechanisms and on the identification of molecular markers to support the implementation of efficient monitoring and resistance management programs. Teflubenzuron is a chitin synthesis inhibitor highly effective in controlling lepidopteran pests, including nowadays the world widely distributed fall armyworm (FAW), *Spodoptera frugiperda* (J.E. Smith, 1797) (Lepidoptera: Noctuidae). We report the selection of a laboratory strain of *S. frugiperda* resistant to teflubenzuron, and its use for the characterization of the inheritance of resistance, evaluation of cross-resistance to other chitin-synthesis inhibitors and the identification of a set of single nucleotide polymorphisms (SNPs) for use as candidate molecular markers for monitoring the evolution of resistance of *S. frugiperda* to teflubenzuron. The resistance of the selected strain of *S. frugiperda* to teflubenzuron was characterized as polygenic, autosomal, and incompletely recessive. The resistance ratio observed was nearly 1,365-fold. Teflubenzuron-resistant strain showed some cross-resistance to lufenuron and novaluron but not to chlorfluazuron. We also detected a set of 72 SNPs that could support monitoring of the resistance frequency to teflubenzuron in field populations. Our data contribute to the understanding of the resistance mechanisms and the inheritance of polygenic resistance of *S. frugiperda* to benzoylureas. We also contribute with candidate markers as tools for monitoring the emergence and spread of teflubenzuron resistance in *S. frugiperda*.

## Introduction

Insect resistance evolution to insecticides and *Bacillus thuringiensis (Bt)*-genetically modified-crops is of great concern to biologists, farmers, industry, and government agencies. The strong selection pressure impinged both by numerous insecticides sprays and the wide adoption of *Bt*-crops increased resistance frequency in many agroecosystems, especially in the successive crop systems used in the central Cerrado savanna region in Brazil (Bolzan et al., 2019; Carvalho, Omoto, Field, Williamson, & Bass, 2013; Farias, Horikoshi, Santos, & Omoto, 2014; Lira et al., 2020; Okuma et al., 2018; Nascimento, Farias, Bernardi, Horikoshi, & Omoto, 2016).

The fall armyworm (FAW) *Spodoptera frugiperda* (J.E. Smith, 1797) (Lepidoptera: Noctuidae) is a polyphagous specie native to American tropical regions which gained worldwide distribution after invading Africa a couple of years (Goergen, Kumar, Sankung, Togola, & Samò, 2016). FAW is a serious pest of several economically important crops, such as maize (Cruz, 1995), cotton (Santos, 2007) and soybean (Bueno, Bueno, Moscardi, Parra, & Hoffman-Campo, 2011). Currently, *Bt* crops and insecticides are the main tactics in use for FAW management in the world (Assefa & Ayalew, 2019).

Insecticides from the benzoylurea group, which were introduced in the early 1970s, act as chitin synthesis inhibitors (van Daalen, Meltzer, Mulder, & Wellinga, 1972). These insecticides have been successfully applied to control several pest species in the field. They have high acaricidal and insecticidal activity, exhibiting larvicidal activity against lepidopterans, coleopterans, hemipterans, and dipterans (Yu, 2014). Moreover, the low non-target effects of benzoylureas allow their use in association with other control strategies within well-designed integrated pest management programs (Beeman, 1982; Oberlander & Silhacek, 1998; Post & Vincent, 1973).

Initial studies on the evolution of resistance to chitin synthesis inhibitors under laboratory conditions failed in selecting resistant populations even after 20 generations of directed-selection pressure (Perng, Yao, Hung, & Sun, 1988). However, their broad use increased the frequency of resistance, leading to evolution of field-evolved resistance of *Plutella xylostella* in China and *Spodoptera littura* in Paskitan (Ahmad, Sayyed, Saleem, & Ahmad, 2008; Lin, Hung, & Sun, 1989), and the selection of resistant strains of *Spodoptera frugiperda* under laboratory conditions (Nascimento et al., 2016).

The evolution of resistance in field-conditions demonstrates that the selection pressure and the resistance frequency are high enough to allow the selection of homozygous-resistant phenotypes, resulting in the complete failure of the resistant management strategies taken in place. The implementation of suitable resistant management plans is required in order to maintain benzoylureas available as a technology for pest control in areas where the resistance frequency is still manageable. The development of a reliable and successful resistance management strategy requires the understanding of the resistance mechanism and its heritability in ways that molecular markers could be consistently used in resistance monitoring programs.

Benzoylureas act as chitin synthesis inhibitors by interfering with the synthesis or deposition of chitin in the exoskeleton and other chitinous structures of insects (Merzendorfer & Zimoch, 2003; Merzendorfer, 2006). The exactly mode of action of benzoylureas has been debated as they were thought to indirectly affect chitin biosynthesis upon binding to sulfonylurea receptors, resulting in vesicle trafficking alterations; however, the role of the ABC transporter sulfonylurea receptor in chitin synthesis was arguable (Abo-Elghar, Fujiyoshi, & Matsumura, 2004; Meyer, Flötenmeyer, & Moussian, 2013). The use of bulk segregants mapping analysis to investigate the resistance mechanism of *Tetranychus urticae* to the mite chitin synthesis inhibitor etoxazole, led to the characterization of resistance of field populations as monogenic and recessive (Van Leeuwen & et al., 2012). They also identified a single nonsynonymous mutation (I1017F) in chitin synthase 1 as the resistance mechanism, demonstrating the direct effect of this acaricide on chitin synthase (Van Leeuwen et al., 2012). Their proposition that benzoylurea with insecticide activity could also target chitin synthase due the similarities with etoxazole, led to the identification of a mutation at the same position for the isoleucine residue in the chitin synthase I in different insect species resistant to chitin synthesis inhibitors (Douris et al., 2016; Fotakis et al., 2020).

Currently, benzoylureas as chlorfluazuron, diflubenzuron, lufenuron, flufenoxuron, novaluron, triflumuron and teflubenzuron are used to control insects in soybean, cotton, and maize crops in Brazil (MAPA 2020). The high selection pressure caused by this group of insecticides has decreased the susceptibility of *S. frugiperda* to benzoylureas (Schmidt, 2002). For example, FAW populations in central Brazil evolved resistance to lufenuron, carrying a high resistance ratio, and an autosomal and polygenic inheritance of resistance (Nascimento et al., 2016).

Teflubenzuron, 1-(3,5-Dichloro-2,4-difluorophenyl)-3-(2,6-difluorobenzoyl) urea, has been used to control lepidopterans, coleopterans, and dipterans larvae (Yu, 2014), since the ovicidal and larvicidal activities of this product were first demonstrated (Ascher & Nemny, 1984). Despite the efficacy of this insecticide in controlling insect pests, there are reports of insect resistance to teflubenzuron as early as in the 1980’s, such as for *Plutella xylostella* (Lepidoptera: Plutellidae) (Iqbal & Wright, 1997; Lin & Sun, 1989; Perng et al., 1988) and *Spodoptera littoralis* (Lepidoptera: Noctuidae) (Ya & Klein, 1990). In Brazil, the increased number of control failures with pyrethroids, organophosphates, and benzoylureas (mainly lufenuron) early in 2000s (Goergen et al., 2016) stimulated the use of teflubenzuron to control *S. frugiperda* in cotton, maize and soybean crops.

In this study, we characterized the genetic basis of resistance of *S. frugiperda* to teflubenzuron. By selecting a resistant strain, we 1) characterized the inheritance of resistance, 2) evaluated the cross-resistance to other chitin-synthesis inhibitors, and 3) used a genome scanning approach to identify genomic regions and single nucleotide polymorphisms (SNPs) associated to teflubenzuron resistance for further use as molecular markers to monitor resistance evolution in field conditions.

## Material and Methods

### Insects

The susceptible *S. frugiperda* strain (Sf-ss) has been maintained on an artificial diet based on bean, wheat germ and casein (Kasten Junior, Precetti, & Parra, 1978) in the Arthropod Resistance Laboratory (University of São Paulo, campus ESALQ, Brazil) without insecticide selection for 25 years. The resistant strain (Tef-rr) was selected from field-collected larvae from maize fields from the state of Mato Grosso, Brazil (13°25′35.9″S; 58°38′17.84″W), during the 2014–2015 crop season.

### Selection of teflubenzuron-resistant *S. frugiperda* strain

Selection for teflubenzuron resistance was carried out using the F_2_ screen method proposed by (Andow & Alstad, 1998). This method increases the likelihood of obtaining a resistant genotype in the F_2_ population and creates large linkage disequilibrium block in the genome that facilitates the genotype to phenotype. About 1,000 field-collected larvae were reared under controlled laboratory conditions (25 ± 2 °C; 60 ± 10% RH; 14 h photophase) up to pupation. From the field-collected larvae, we isolated 33 individual couples (families). The progeny obtained for each family was reared under the same controlled conditions described before until adult emergence. Adults originating from these families, which correspond to a F_1_ generation, were allowed to sib-mate. All eggs laid from each family were collected. Then, 120 larvae from each sib-mated line (i.e. from the F_2_ generation) were used in insecticide bioassays. We used a common diet-overlay bioassay to select for teflubenzuron resistance. The artificial diet (Kasten Junior et al., 1978) was poured into 24-well acrylic plates (Costar^®^, Corning^®^), and 30 µL/well of the diagnostic concentration (10 µg/mL) of teflubenzuron (Nomolt® 150, teflubenzuron 150 g/L, BASF S.A., São Paulo, Brazil) in a water (v:v) solution containing 0.1% Triton X-100 was applied on the diet surface. The control diet was treated only with distilled water and 0.1% Triton X-100. Plates were kept under a laminar flow hood for drying the diet surface, and each well was inoculated with one *S. frugiperda* third instar larva. We allowed larvae to feed for five days under controlled conditions (25 ± 2 °C; 60 ± 10% RH; photophase of 14 h). After five days, we collected and transferred the surviving *S. frugiperda* larvae from both experimental and control diets to plastic cups (100 mL) containing 50 mL of artificial diet for their rearing until pupation. All surviving individuals from the insecticide treatment were combined and used for successive selection rounds with teflubenzuron until the establishment of the teflubenzuron-resistant strain (Tef-rr).

### Characterization of *S. frugiperda* resistance to teflubenzuron

The susceptible (Sf-ss) and resistant (Tef-rr) strains of *S. frugiperda* to teflubenzuron were subjected to concentration-response assays with five to 12 logarithmically spaced concentrations between 0.1 and 3,200 μg.mL^-1^ of teflubenzuron. Larval bioassays were conducted using the diet overlay assays earlier described. The lethal concentration 50 (LC_50_) was estimated with Probit analysis (Finney, 1949; Lewis & Finney, 1971; Lin & Sun 1989) using the POLO software (Robertson et al., 2007). The resistance ratio of Tef-rr was calculated by dividing the LC_50_ of the Tef-rr by that of the Sf-ss strain.

### Estimation of dominance levels

Newly emerged adults from the susceptible (Sf-ss) and resistant (Tef-rr) strains were reciprocally crossed: Tef-rr males × Sf-ss females (RC-1) and Tef-rr females × Sf-ss males (RC-2). Adults (10 couples/cage) were kept in PVC cages (20 cm high × 15 cm diameter) lined with paper to serve as substrate for egg laying. Adults were fed with a 10% honey solution that was replaced every other day. The progenies obtained from each reciprocal crosses (F_1_) were reared on artificial diet until the third instar. Afterwards, third-instars from the reciprocal crosses were exposed to teflubenzuron using the diet-overlay assay explained before.

We estimated the dominance level of resistance from (Bourguet, Genissel, & Raymond, 2000), 

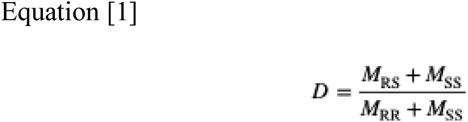

where *M*_*SS*_,*M*_*RR*_ and *M*_*RS*_ are the mortalities of the Sf-ss, Tef-rr and heterozygous strains, respectively, exposed to different concentrations of teflubenzuron. *D* values close to zero (*D* = 0) represent completely recessive inheritance, and values close to 1 (*D* = 1) represent completely dominant inheritance.

We also estimated dominance level by applying equation [2] (Stone, 1968), where D is the degree of dominance and X_F_, X_R_, X_S_ are the LC_50_ values, respectively, for the heterozygote (offspring from reciprocal cross RC1 or RC2), Tef-rr and Sf-ss. 

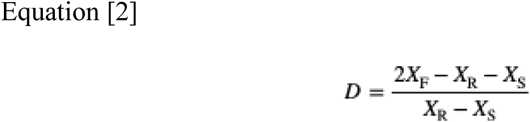

### Genetic inheritance associated with teflubenzuron resistance in *S. frugiperda*

We used the method proposed by Roush and Daly (1990) and Tsukamoto (1983) to test the hypothesis that a single gene is responsible for teflubenzuron resistance of *S. frugiperda*. We backcrossed the offspring resulting from the twenty-mating pair Tef-rr♂x Sf-ss♀ (heterozygous) with individuals from the resistant strain Tef-rr. We performed diet-overlay bioassays, using eight concentrations of teflubenzuron as earlier described.

The possibility of monogenic inheritance was calculated by using the chi-square test [Equation 3] (Sokal & Rohlf, 2012), where *Ni* is the mortality observed in backcrossed larvae at each concentration *i*; *ni* is the number of individuals tested; *q* is the expected survival, and *p* is the expected mortality calculated from the Mendelian model [Equation 4] (Georghiou, 2012), where *a* is the mortality obtained for the parental strain Tef-rr, and *b* is the mortality of the heterozygote derived from the reciprocal crosses (Tef-rr♂x Sf-ss♀). The hypothesis of monogenic inheritance is rejected when the calculated chi-square is equal or higher than the tabulated chi-square value, with 1 degree of freedom. 

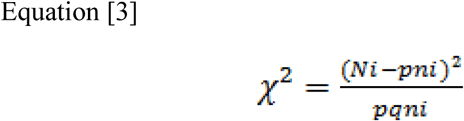

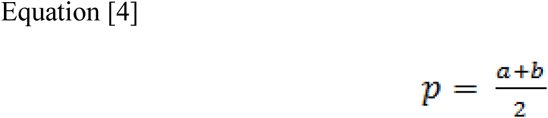

### Cross-resistance to other benzoylurea insecticides

Cross-resistance assays of the Tef-rr strain with three other benzoylureas were carried out using the diet-overlay bioassay earlier described utilizing commercial formulations of lufenuron (Match®, 50 g/L, Syngenta, Basel, Switzerland), novaluron (Rimon Supra®, 100 g/L, Syngenta) and chlorfluazuron (Atabron, 50 g/L, ISK Biosciences). For each insecticide we performed concentration-response bioassays for Tef-rr and Sf-ss as already described. Larval mortality was assessed five days after treatment, and larval mortality was characterized by larval unresponsiveness to stimulation with a fine brush or the occurrence of body malformations. LC_50_ values were estimated using the POLO software (Robertson et al., 2007).

#### Genetic Crossing and Isolation of the Pools

The pool sequencing was designed to highlight potential markers associated with resistance. The offspring from the e reciprocal crosses described above (RC-1 and RC-2), we established backcrosses with the resistant strain (Tef-rr), e.g. BC1 = (RC-1♂x Tef-rr♀), BC2 = (RC-1♀ x Tef-rr♂), BC3 = (RC-2♂x Tef-rr♀), BC4 = (RC-2♀ x Tef-rr♂). Each backcross was split in two groups of insects: 1) individuals randomly collected (BC - random) and 2) individuals that survived the selection pressure by exposure to the diagnostic concentration of 10 µg teflubenzuron/mL (BC - selected) (Figure 1).

**Figure 1.**
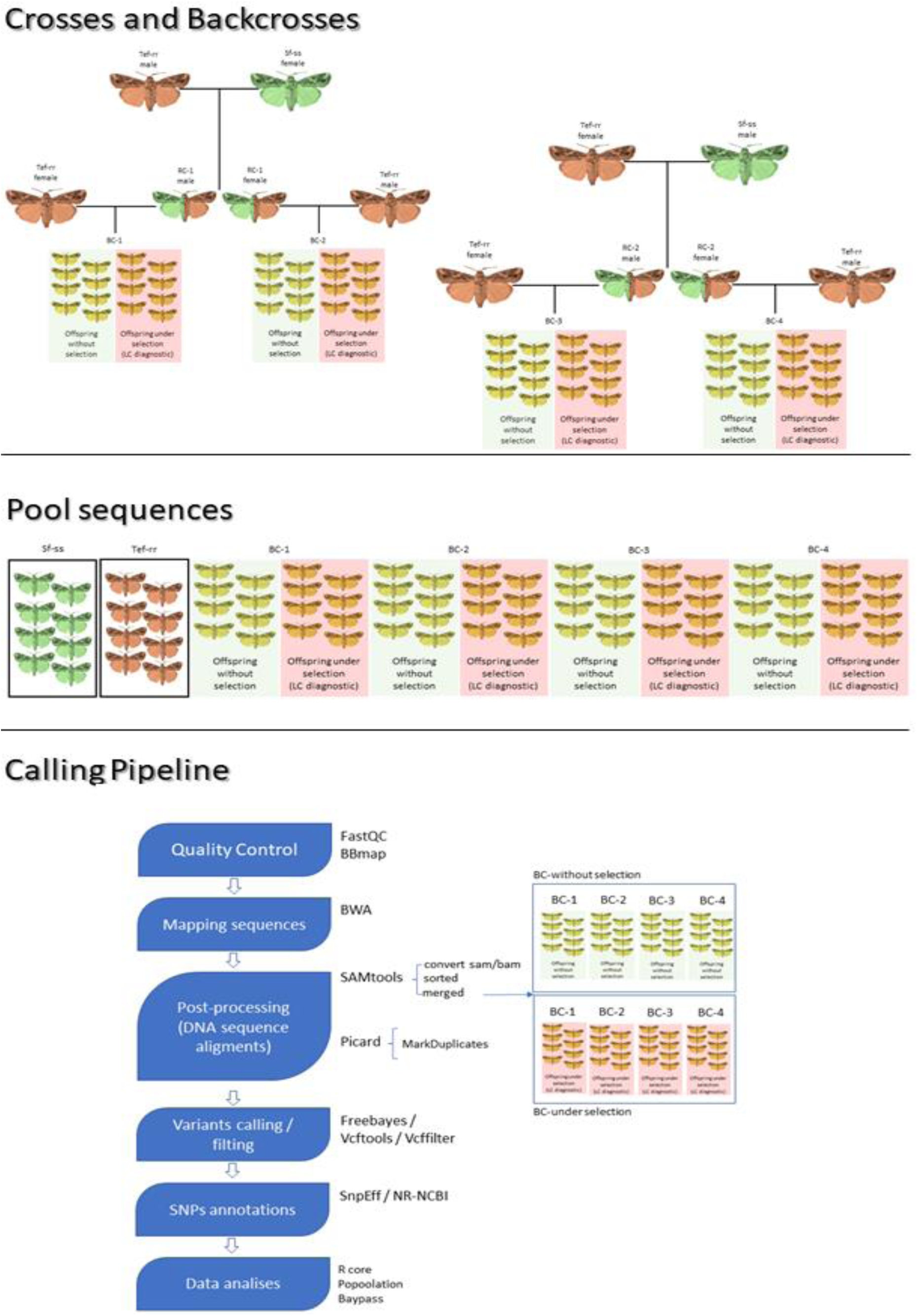
Experimental design, pool sequences and snp-calling pileline.

#### DNA Extraction and Sequencing

DNA was extracted from nine larvae from each parental line, Tef-rr (resistant) and Sf-ss (susceptible), and from both groups of each backcross BC1, BC2, BC3 and BC4. Larval genomic DNA was obtained with the modified CTAB method (Doyle, Doyle, & Hortorium, 1990). Briefly, 50 mg of tissue from each individual larva was macerated in 650 μL of extraction buffer containing 2% cetyltrimethyl ammonium bromide (CTAB), 1.4 M NaCl, 100 mM tris(hydroxymethyl)aminomethane (Tris-HCl) at pH 8.0, 20 mM ethylenediaminetetraacetic acid (EDTA) at pH 8.0, 1% polyvinylpyrrolidone, 0.2% β-mercaptoethanol, and 20 µL of proteinase K (0.1 μg·mL^−1^). Samples were incubated at 55 °C for 1 h, added with 650 μL of chloroform: isoamyl alcohol (24:1), and mixed until emulsion. Samples were centrifuged (14,000 *g* × 5 min × 4 °C), and then the supernatant was collected and transferred to new tubes, where 200 μL of the same extraction buffer (minus the β-mercaptoethanol and proteinase K), was added followed by the addition of the same volume of chloroform: isoamyl alcohol (24:1). The emulsion was thoroughly vortexed, centrifuged (14,000 *<u>g</u>* × 5 min × 4 °C), and the supernatant collected; we repeated this process 3 times. Samples were combined with 650 μL of cold isopropanol and incubated at –20 °C overnight before centrifugation (14,000 *g* × 5 min × 4 °C). The pelleted DNA was washed twice with 1 mL of 70% ethanol. The pellet was dried at room temperature, resuspended in 40 µL TE and Rnase A (10 μg.m L^−1^), and stored at –20 °C until further analyses. Genomic DNA was evaluated quantitatively with a Qubit fluorometer (Thermo Fisher Scientific, USA) and checked for degradation by agarose gel electrophoresis. Finally, 7 ng of DNA from each one of nine larvae sampled were combined into a single tube for each treatment.

Briefly, we sheared total pooled DNA into ∼300–400 bp fragments in an ultrasonicator and used it to build sequencing libraries with an NEBNext Ultra DNA library prep kit (New England Biolabs), according to the manufacturer’s instructions. The whole genome (WGS) for each pool was sequenced in a Miseq platform (Illumina, Inc., San Diego, CA, USA) at the Molecular and Cellular Imaging Center at the Ohio State University.

#### Sequencing Data Processing

The quality of raw paired-end reads was assessed using FastQC (Andrews et al., 2015), and reads were filtered using BBmap (http://jgi.doe.gov/data-and-tools/bbtools) by excluding nucleotides with a Phred quality score < 30 from subsequent analyses. Afterwards, the filtered reads were mapped against the *S. frugiperda* pseudo-genome assembly available at BIPAA - Bioinformatics Platform for Agroecosystem Arthropods (https://bipaa.genouest.org/sp/spodoptera_frugiperda_pub/), using the BWA-MEM (Li & Durbin, 2010). Alignment files were converted to SAM/BAM files using SAMtools (Li, 2011). Alignment in BAM format from the BC1-random, BC2-random, BC3-random and BC4-random were combined on a single BAM file (BC-random), whereas BC1-selected, BC2-selected, BC3-selected and BC4-selected were combined on a single BAM file (BC-selected). Read alignments with PCR duplicates were removed using the *MarkDuplicates* from Picard software (https://broadinstitute.github.io/picard/), and SNP calling was performed using freebayes (Garrison & Marth, 2012). SNPs called were subject to quality filters (quality score > 20 and depth > 10) using the programs Vcftools (Danecek et al., 2011) and Vcffilter (Müller & Jimenez, 2017).

#### Analyses

The vcfR package was used to visualize and manipulate the vcf format in the. The global F_ST_ was calculated for all SNPs using the R package PoolFstat. Tajima’s π and D were calculated for each pooled DNA sample in a 5 kb sliding window with a 5 kb step size for each comparison groups using Popoolation v.1.2.2 (Kofler et al., 2011).

Candidate SNPs associated with the resistance of *S. frugiperda* to teflubenzuron were identified using two approaches: 1) a population genomics-based approach, which uses the genetic differentiation between the pools to identify genomic regions potentially targeted by selection (Pool-GWAS), and 2) a BSA-seq approach, which identify genomic regions enriched with SNPs that have allele frequencies (either the reference or the alternative allele) highly differentiate between the segregating pools (here the backcrosses-derived individuals that were randomly selected to form the BC-random) and those that survived to the insecticidal application (here the treated BC pool BC-selected).

For the population genomics-based approach, SNP count data was analyzed using two different implementations of the bayesian hierarchical models available in the Baypass version 2.2 (Gautier, 2015). First, we applied the core model (Coop, Witonsky, Rienzo, & Pritchard, 2010; Nicholson et al., 2002) to identify loci with significant allele frequency differences. This method is equivalent to the methods that searches for loci with higher intra-locus F_ST_. However, in this model, a variance-covariance matrix of population allele frequencies (Ω matrix) that works as a kinship matrix is used to control for population structure. Controlling for population structure reduces the likelihood of spurious association between the marker and the phenotype. This method is covariate-free and was expanded to include the calibration of the XtX statistics as proposed by (Günther & Coop, 2013). Second, we employed the STD model representing an extension of core model, which allows the evaluation of the association of SNP allele frequency with an covariates (Gautier, 2015). For the covariate model we conducted two independent analysis: 1) using the the LC_50_ as covariate, and 2) using the mortality obtained with the diagnostic concentration for each bulk (susceptible, resistant and the BC-random and BC-selected) as a covariate. Because we performed the analysis with the two backcrosses-derived and the parental pools, both genome scan – the FST-like method and the covariate association method were insensitive to the identification of false positive associations between the markers and the phenotype. These methods are prone to the identification of the strongest signal that might highlight higher differences mostly associated with demography and drift (because the resistant and the susceptible populations share a common ancestor many generations ago), not with selection. The identification of true positives can be done with the identification of SNPs in linkage disequilibrium with the causal gene in the pool that was subjected to selection the BC-selected, and in loci that presents allele frequencies differences between the BC before and after selection.

#### Functional annotation and identification of putative markers associated with the resistant phenotype

Annotation of loci associated with SNPs was proposed using the *gff* file available for the *S. frugiperda* genome (https://bipaa.genouest.org/sp/spodoptera_frugiperda_pub/) using SnpEff (Cingolani et al., 2012). Genes with no functional annotation in the available genome were annotated after heuristic search using the BLASTx algorithm against the non-redundant protein database available at the NCBI.

## Results

### Characterization of teflubenzuron resistance of *S. frugiperda*

Eleven out of the 33 lines of the F_2_ generation subjected to selection yielded survivors, which were considered to carry resistance traits. The high resistance traits selected in the teflubenzuron-resistant strains were observed by comparing the concentration response curves of the Tef-rr against the Sf-ss and their backcrosses (Figure 2), and the obtained values of LC_50_. The LC_50_ for the resistant Tef-rr (641.47 μg.mL^−1^; IC = 213.05 – 2,748.81 μg.mL^−1^) was nearly 1,365-fold the LC_50_ for the susceptible Sf-ss strain (0.47 μg.mL^−1^; IC = 0.35 – 0.63 μg.mL^−1^). Both Sf-ss (*P* = 0.02) and Tef-rr (*P* = 0.03) showed no evidence of distortion at χ^2^ > 0.01, indicating a good fit to the probit inheritance model of resistance (Table 1).

**Table 1.** Concentration – mortality to teflubenzuron of susceptible (Sf-ss) and resistant (Tef-rr) *S. frugiperda* strains and progenies of reciprocal crosses between Sf-ss and Tef-rr strains

| Strains | n | Slope±SE | LC<sub>50</sub> (95% CI)*<br>(µg AI mL<sup>-1</sup>) | χ<sup>2</sup> | df** | RR*** |
| --- | --- | --- | --- | --- | --- | --- |
| Sf-ss | 715 | 3.07 ± 0.24 | 0.47(0.35 – 0.63)<sup>a</sup> | 11.630 | 4 | - |
| Tef-rr | 840 | 0.64 ± 0.09 | 641.47 (213.05 – 2,748.81)<sup>c</sup> | 10.590 | 4 | 1,364.83 |
| Tef-rr♂ vs Sf-ss♀ | 936 | 2.05 ± 0.14 | 4.88 (3.58 – 6.31)<sup>b</sup> | 8.054 | 5 | 10.38 |
| Tef-rr♀ vs Sf-ss♂ | 983 | 2.29 ± 0.17 | 3.94 (3.13 – 4.78)<sup>b</sup> | 7.604 | 6 | 8.38 |

1. \*LC<sub>50</sub> values followed by the same letter in the columns do not differ significantly for the confidence intervals (95%). The significance of the confidence intervals was determined by the likelihood ratio test, followed by multiple comparisons. 2. \*\*df = degrees of freedom. 3. \*\*\*Resistance ratio (RR) = LC<sub>50</sub> of resistant strain/LC<sub>50</sub> of susceptible strain.

**Figure 2.**
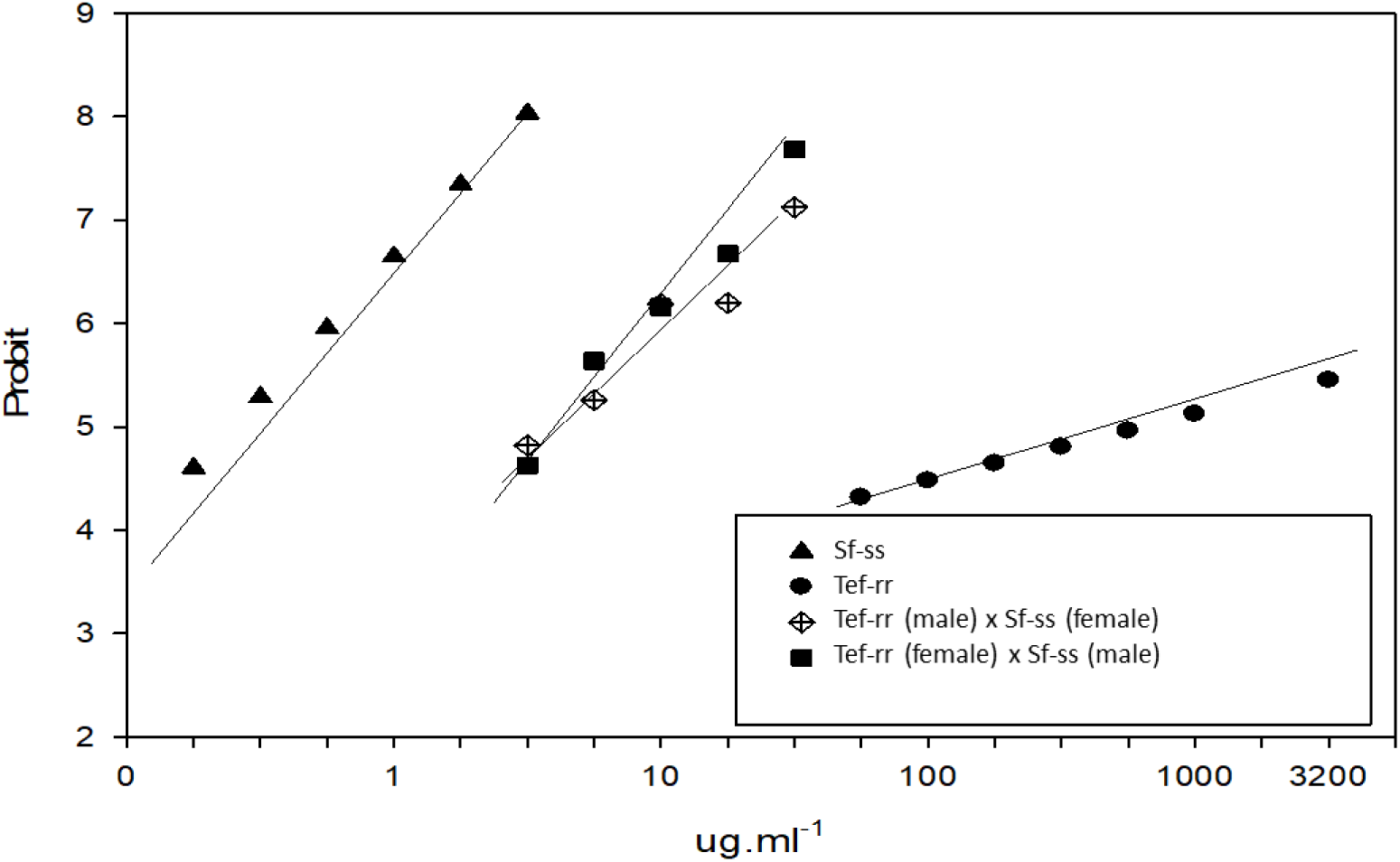
Log concentration–probit of susceptible (Sf-ss) and resistant (Tef-rr) *S. frugiperda* strains and progenies of reciprocal crosses between susceptible and resistant strains

Bioassays with progenies of the two reciprocal crosses showed no significant differences, since there was an overlap of 95% CI of the LC_50_ values (Table 1). Therefore, the hypothesis of parallelism was not rejected (*P* = 0.247, df = 1). The overlap of the confidence intervals indicated that inheritance of teflubenzuron resistance of *S. frugiperda* is autosomal, and not related to maternal effects or sex linked.

The dominance values for the reciprocal crosses of the offspring estimated following Stone (1968) were 0.32 (Tef-rr♂× Sf-ss♀) and 0.29 (Tef-rr♀ × Sf-ss♂). The dominance level estimated using the Bourguet-Genissel-Raymond method showed decreased dominance with increased teflubenzuron concentrations (Figure 3). In both cases, teflubenzuron resistance of *S. frugiperda* was shown to have an incompletely recessive inheritance.

**Figure 3.**
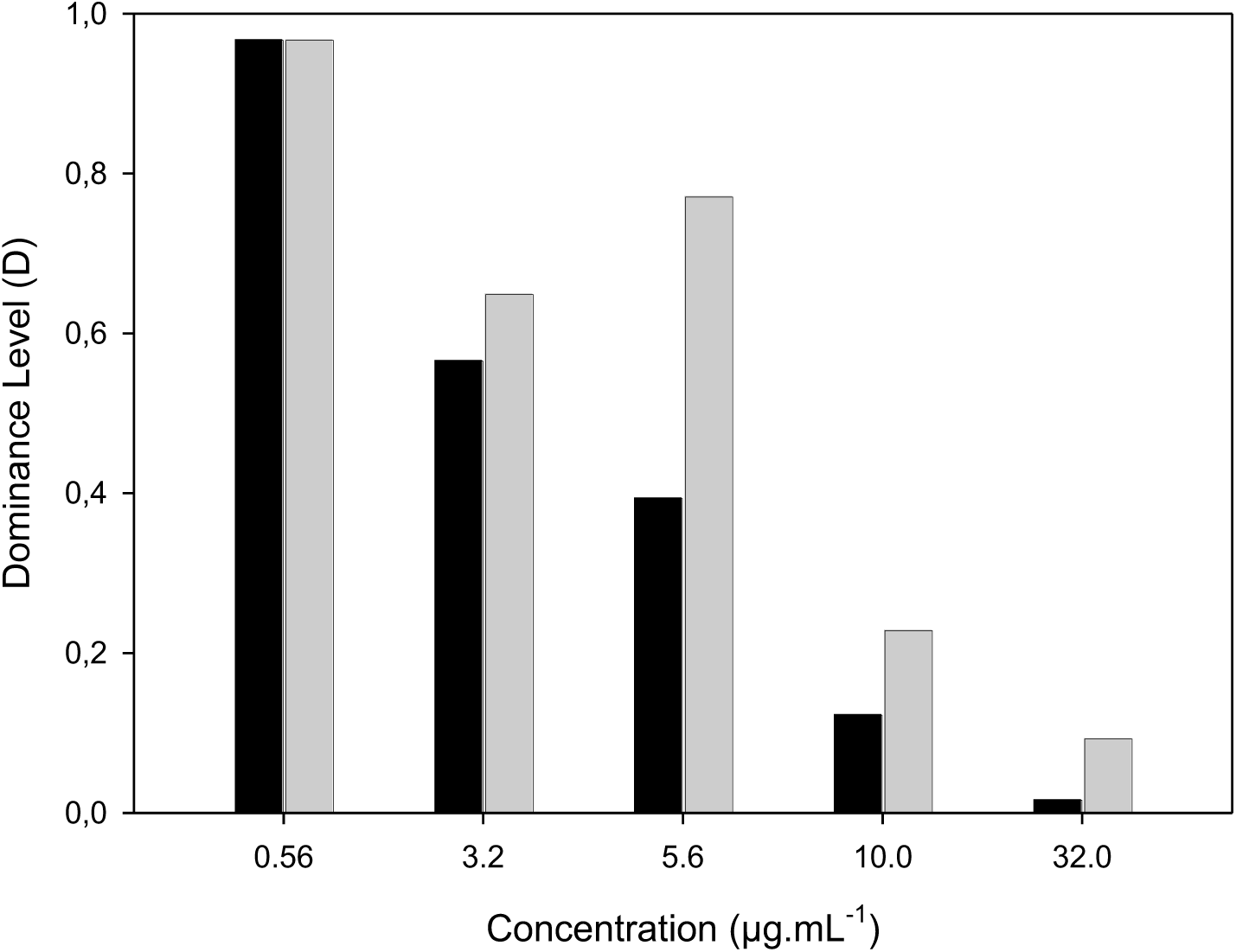
Level of dominance of *S. frugiperda* resistance as a function of teflubeunzuron concentration.

The direct hypothesis test for monogenic inheritance of the teflubenzuron resistance of *S. frugiperda* based on larval mortality of the F_1_ × Tef-rr backcross was significant (P < 0.01) for concentrations between 1 and 10 μg.mL^−1^.This result allows the rejection of the hypothesis that teflubenzuron resistance in the selected strain is monogenic (Table 2).

**Table 2.** Chi-square analysis of the mortality data from backcross between the progeny of reciprocal cross (Tef-rr♂x Sf-ss♀) and Tef-rr *S. frugiperda* strain (F1 progeny) exposed to different concentrations of teflubenzuron

| Concentration<br>µg AI mL<sup>-1</sup> | Expected<br>mortality | Observed<br>mortality | $\chi^2$ (df = 1) | P |
| --- | --- | --- | --- | --- |
| 1 | 1.042 | 5.042 | 18.47 | < 0.00001* |
| 3.2 | 21.677 | 9.574 | 8.10 | 0.0044* |
| 10 | 44.167 | 22.222 | 14.06 | 0.0001* |
| 32 | 49.167 | 48.958 | 0.001 | 0.9674 |
| 100 | 64.600 | 68.750 | 0.72 | 0.3951 |
| 320 | 70.795 | 78.125 | 2.49 | 0.1142 |
| 1000 | 80.530 | 82.291 | 0.19 | 0.6629 |

### Cross-resistance

Both susceptible and teflubenzuron-resistant strains were tested against chlorfluazuron, lufenuron and novaluron (Table 3). Teflubenzuron-resistant strain showed some cross-resistance to lufenuron (121.75-fold) and novaluron (75.8-fold), no cross-resistance of the Tef-rr strain to chlorfluazuron (4-fold) was detected (Table 3).

**Table 3.**
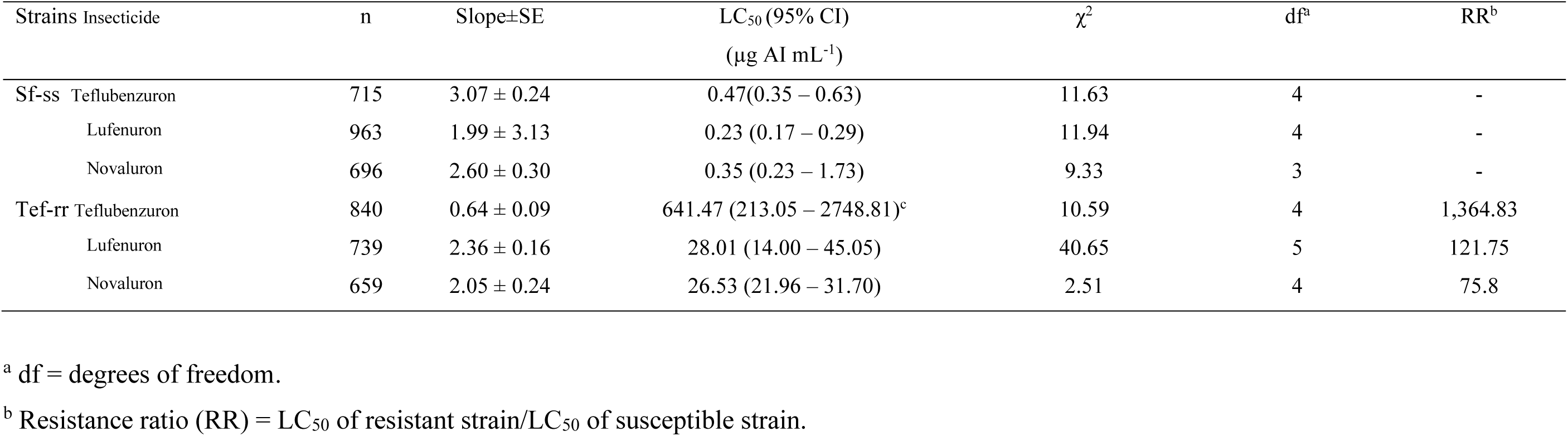
Cross-resistance of *S. frugiperda* resistant strain (Tef-rr) to benzoylphenyl ureas insecticides

### Genome-scan of Tef-rr and Sf-ss strains of *S. frugiperda*

The sequencing of the pooled WGS libraries generated 60.5 million high-quality paired-end reads after adapter removal and quality trimming. The read mean length was 229 bp and the maximum length was 300 bp, resulting in paired-end fragments with an average of 429 bp in length, with an estimated coverage of 108-fold (Table S1). After mapping and filtering the reads against the *S. frugiperda* reference genome, 890,209 SNPs were called.

The global genetic differentiation among the resistant, the susceptible and the two backcrosses samples were moderate and significant (F_ST_ = 0.10169 with 95% CI of 0.10138 - 0.10205). The pairwise measurements between BC-random and BC-selected showed that both backcross pools were virtually identical (F_ST_ = -0.01454 95% CI of - 0.0146 - -0.01442). However, the pairwise F_ST_ between Sf-ss and Tef-rr was high (0.37581 with 95% CI of 0.37489 - 0.37684). Genetic differentiation was also high between Sf-ss and BC-random (F_ST_ = 0.20461 with 95% CI of 0.20394 - 0.20532), and Sf-ss and BC-selected (F_ST_ = 0.21186 with 95% CI of 0.21116 - 0.21253) (Figure S2).

The variation within the sliding window estimates for π resulted in nucleotide diversities of 0.0192, 0.0285, 0.0336, and 0.0337, respectively for the Sf-ss and Tef-rr strains and their back crosses. We identified genomic regions with reduced diversity when comparing the Sf-ss and Tef-rr strains. Large genomic regions with reduced diversity were observed when comparing the resistant and the susceptible strains (Tajima’s π < 0.002), including 600 scaffolds from *S. frugiperda* genome. Regions of low nucleotide diversity can be observed in Table S3.

The total number of SNPs called was narrowed to 9,161 after the application of *XtX* statistics (*XtX p-value* < 0.001) with evidence of divergent selection. We also identified 4,120 SNPs with eBF > 3 db, supporting a moderate evidence of association. Four SNPs exceeded the threshold for strong evidence of association with LC_50_ values according to the Jeffreys’ rule, whereas 38 SNPs supported moderate evidence of association with mortality using the diagnostic concentration as variable. Finally, 537 SNPs overlapped with both the neutrality model and the association with environmental variables model (Figure S4), distributed in 232 scaffolds.

Scaffold annotation indicated that most of the variants under selection were located in intergenic (47%) and regulatory regions (downstream regions - 18%; upstream regions - 18%), with 2% presenting missense effect (Figure S5). GO terms distribution demonstrated an impressive number of variants on processes related to primary metabolism, metabolism of organic compounds, and components of membrane (Figure S4). We identified 19 SNPs with non-synonymous effects (Table 4), distributed in 19 scaffolds. Most of the variants are in regulatory regions (Table 5).

**Table 4.** Missense mutation under selection in *S. frugiperda* resistant to teflubenzuron.

| scaffold | Position | Nucleotide<br>modification | Protein<br>modification | Gene | Description |
| --- | --- | --- | --- | --- | --- |
| pseudoscaff_1708 | 178922 | A2228G | Asn743Ser | GSSPFG00030930001 | GATA zinc finger domain-containing protein 14-like |
| pseudoscaff_3427 | 17439 | T1358C | Val453Ala | GSSPFG00031119001.<br>2 | cytochrome CYP340AB1 |
| pseudoscaff_3907 | 6057 | C616T | Pro206Ser | GSSPFG00030391001 | nose resistant to fluoxetine 6-like |
| pseudoscaff_653 | 4817 | C764T | Ser255Leu | GSSPFG00012677001 | cuticle CPG4855 |
| pseudoscaff_957 | 60700 | T731A | Met244Lys | GSSPFG00023674001 | probable serine threonine- kinase kinX |
| pseudoscaff_1765 | 469701 | T644G | Leu215Arg | GSSPFG00027555001 | serine protease |
| pseudoscaff_3391 | 127787 | C2146T | Pro716Ser | GSSPFG00003324001 | uncharacterized protein LOC110384046 isoform X1 |
| pseudoscaff_3689 | 183291 | C518T | Thr173Met | GSSPFG00005692001.<br>1 | uncharacterized protein LOC110370494 |
| scaffold_168 | 131785 | C1496T | Pro499Leu | GSSPFG00028493001 | UPF0061 Pfl01_0444-like |
| pseudoscaff_2533 | 256404 | G200C | Trp67Ser | GSSPFG00005566001 | unknown |
| pseudoscaff_3278 | 1521 | C971A | Thr324Asn | GSSPFG00015719001 | rho GTPase-activating 11A-like |
| pseudoscaff_1247 | 66548 | C203T | Thr68Ile | GSSPFG00033321001.<br>3 | chemosensory ionotropic receptor IR40a |
| pseudoscaff_1409 | 38337 | G160A | Ala54Thr | GSSPFG00025253001 | sid 3 |
| pseudoscaff_1573 | 23130 | T428G | Leu143Trp | GSSPFG00009731001 | peptidyl-tRNA hydrolase mitochondrial-like |
| pseudoscaff_2560 | 3374 | C2099T | Thr700Met | GSSPFG00009517001.<br>3 | ubiquitin carboxyl-terminal hydrolase 36 isoform X1 |
| superscaffold_605 | 216090 | C341T | Pro114Leu | GSSPFG00011282001 | alpha-1,3/1,6-mannosyltransferase ALG2 |
| pseudoscaff_3041 | 7075 | C1067T | Ala356Val | GSSPFG00001156001 | serpin B8-like |
| pseudoscaff_2143 | 20100 | G833A | Arg278Lys | GSSPFG00012527001 | eukaryotic initiation factor 4A-III |
| pseudoscaff_3248 | 90010 | T754C | Cys252Arg | GSSPFG00017635001 | spindle assembly abnormal 6 homolog |

**Table 5.** Candidate SNPs on downstream, upstream, intronic and splice regions, SNP-index and functional description.

| Scaffold | Amido acid | Region of variant | Functional description |
| --- | --- | --- | --- |
| pseudoscaff_1297 | A22203C | downstream | protein charybde-like |
| pseudoscaff_1307 | A232656G | downstream | insulin-like precursor polypeptide 2 |
| pseudoscaff_1362 | G67863C | downstream | U4 tri-snRNP-associated 1 isoform X2 |
| pseudoscaff_1648 | C137023A | downstream | alpha-tubulin N-acetyltransferase-like isoform X2 |
| pseudoscaff_1648 | C137251T | downstream | alpha-tubulin N-acetyltransferase-like isoform X2 |
| pseudoscaff_1775 | T199512C | downstream | transcription initiation factor IIA subunit 2 |
| pseudoscaff_2258 | G33649A | downstream | methyltransferase-like 26 |
| pseudoscaff_2598 | C12345T | downstream | allatostatin-A receptor |
| pseudoscaff_2606 | G19017A | downstream | acyl-desaturase |
| pseudoscaff_2896 | T22770C | downstream | insulin-like growth factor-binding complex acid labile subunit isoform X3 |
| pseudoscaff_3278 | C1521A | downstream | F-box only 11 |
| pseudoscaff_3604 | T173487C | downstream | homeobox orthopedia-like isoform X2 |
| pseudoscaff_3799 | A16059G | downstream | E3 ubiquitin-protein ligase MARCH1-like isoform X1 |
| pseudoscaff_4151 | G20990A | downstream | STAGA complex 65 subunit gamma-like |
| pseudoscaff_4151 | C21023T | downstream | STAGA complex 65 subunit gamma-like |
| scaffold_168 | G131785A | downstream | DNA repair and recombination RAD54-like |
| superscaffold_161 | T93320C | downstream | beclin 1-associated autophagy-related key regulator |
| pseudoscaff_1108 | C68071T | intron | gamma-aminobutyric acid type B receptor subunit 1- |
| pseudoscaff_1494 | T585C | intron | paraplegin |
| pseudoscaff_1648 | A135286C | intron | alpha-tubulin N-acetyltransferase-like isoform X2 |
| pseudoscaff_1648 | A126799G | intron | alpha-tubulin N-acetyltransferase-like isoform X2 |
| pseudoscaff_1877 | C65279T | intron | EH domain-containing 3 |
| pseudoscaff_2418 | T62036G | intron | G- coupled receptor Mth2-like |
| pseudoscaff_2486 | C31733G | intron | scm-like with four MBT domains 2 |
| pseudoscaff_2598 | G6055A | intron | allatostatin-A receptor |
| pseudoscaff_26 | T40740A | intron | Guanylate cyclase |
| pseudoscaff_4046 | A194050T | intron | probable 3-hydroxyacyl- dehydrogenase isoform X2 |
| pseudoscaff_838 | T36912C | intron | UDP-glucuronic acid decarboxylase 1 isoform X1 |
| pseudoscaff_10 | G53445T | splice_region | eukaryotic translation initiation factor 3 subunit |
| pseudoscaff_1095 | G68217A | upstream | BTB/POZ domain-containing protein 6-B |
| pseudoscaff_1230 | G44162C | upstream | Coup transcription factor |
| pseudoscaff_1244 | T24415C | upstream | small GTPase Rab4b |
| pseudoscaff_1282 | A22175T | upstream | cytochrome P450 337B3v2 |
| pseudoscaff_1394 | A541334T | upstream | cytochrome P450 CYP321A9 |
| pseudoscaff_1456 | C16151G | upstream | probable phosphatase 2C |
| pseudoscaff_1648 | T166815C | upstream | tyrosine- kinase |
| pseudoscaff_165 | A42337G | upstream | mannose-1-phosphate guanyltransferase beta |
| pseudoscaff_1745 | T20426C | upstream | pancreatic triacylglycerol lipase-like |
| pseudoscaff_2012 | G288294A | upstream | aldo-keto reductase |
| pseudoscaff_2143 | G20100A | upstream | DDB1- and CUL4-associated factor 10 isoform X2 |
| pseudoscaff_215 | T100348A | upstream | disks large 1 tumor suppressor isoform X6 |
| pseudoscaff_2153 | A41682T | upstream | chromatin modification-related eaf-1-like isoform X1 |
| pseudoscaff_2411 | A307493G | upstream | ADP-ribosylation factor 6-interacting 4 |
| pseudoscaff_3028 | G39454A | upstream | exocyst complex component 8 |
| pseudoscaff_3790 | A99724T | upstream | UDP-glucose 4-epimerase-like |
| pseudoscaff_3799 | A16059G | upstream | eukaryotic translation initiation factor 6 |
| pseudoscaff_4013 | A27061G | upstream | ras-related Rab-39B |
| pseudoscaff_4148 | C20818T | upstream | triosephosphate isomerase |
| pseudoscaff_597 | T129616A | upstream | eukaryotic translation initiation factor 2A |
| pseudoscaff_597 | T58764A | upstream | MRN complex-interacting |
| pseudoscaff_838 | T36912C | upstream | multidrug resistance-associated 4-like |
| scaffold_15106 | T1653C | upstream | steroidogenic acute regulatory-like isoform X1 |
| scaffold_4023 | A1781G | upstream | E3 ubiquitin-ligase RNF13 isoform X3 |

\* SNP index is the ratio of the number of reads with mutant allele at a SNPS locus to the total number of reads covering this SNP locus.

## Discussion

In the present study, we selected a strain highly resistant (≈ 1,365-fold) to the teflubenzuron, from a field-collected population of *S. frugiperda* in the state of Mato Grosso, Brazil. Teflubezuron resistance was found to be polygenic, incompletely recessive with an autosomal mode of inheritance. *Spodoptera frugiperda* resistant to teflubenzuron showed evidence of cross resistance to lufenuron and novaluron, but not to chlorfluazuron.

The pattern of genetic inheritance of *S. frugiperda* resistance to teflubenzuron is similar and common to lepidopteran species resistant to several insecticides and *Bt* toxins, e.g., Dipel resistance and Cry1Ab resistance in *Ostrinia nubilalis* (Crambidae) (Huang, Buschman, Higgins, & McGaughey, 1999), and in *S. frugiperda* resistant to lufenuron (Nascimento et al., 2016), spinosad (Okuma et al., 2018), and spinetoram (Lira et al., 2020).

The inheritance mechanism of teflubenzuron resistance was influenced by insecticide concentration. At lower concentrations resistance inheritance assumes incompletely dominant features, but at higher concentrations it becomes incompletely recessive. The higher concentration is close to the recommended concentration currently used in field applications for *S. frugiperda* control. In resistance management, the level of dominance is a variable feature, resulting not only from the genetic background, but also from the interaction between phenotypes and environmental conditions (Bourguet et al., 2000). The level of dominance is one of the most important features for successful IRM (Lenormand & Raymond, 1998), since the frequency of resistant insects could be related to the level of dominance. But if resistance inheritance is recessive, the evolution of resistance is delayed because the resistant phenotype is present only in homozygotes, and the alleles that confer resistance are rare (Ffrench-Constant, 2013). However, the use of concentrations lower than the level recommended for field application helps to maintain heterozygous individuals in the system and increases the frequency of the resistant alleles within the population. Thus, the continued exposure to the selection pressure (teflubenzuron applications) favors the rapid increase of the degree of individual resistance, leading to a concomitant increase in the likelihood of heterozygous mating, and an ultimate production of resistant homozygotes.

The significant deviation between the observed and expected mortalities observed for the offspring of the resistant – susceptible backcrosses in three concentrations of teflubenzuron indicates resistance is based on more than one gene. Multiple genes with additive and quantitative effects can also lead to the generation of resistance features, and in these cases, it has been difficult to identify a specific gene or genetic marker associated with such evolutionary process. The polygenic nature of resistance of *S. frugiperda* to teflubenzuron agrees with the resistance characterized for another strain of this insect to the benzoylurea lufenuron (Nascimento et al., 2016).

The cross-resistance of teflubenzuron-resistant insects to lufenuron and novaluron may be related to strong selection of insects with overexpression of the detoxification genes, such as cytochrome P450 (CYP), glutathione S-transferases (GSTs), UDP-glucosyltransferases (UGTs), and esterases (CCEs) (Nascimento, Fresia, Cônsoli, & Omoto, 2015). These genes are largely associated with detoxification of xenobiotic compounds in several lepidopteran species. Therefore, selection of these genes within these superfamilies may be responsible for the evolution of resistance to different insecticide compounds within the same IRAC group.

The non-cross resistance detected to chlorfluazuron as compared to the high cross-resistance levels observed for lufenuron and novaluron agree with data available on cross-resistance among different benzoylurea compounds. Cross-resistance to benzoylureas with other chemical compounds is rarely reported, and few are the reports of cross-resistance among benzoylurea compounds (Perng et al., 1988). Chlorfluazuron cross-resistance to other benzoylureas, if detected, is always very low (cross resistance to teflubenzuron = 9.9-fold) (Fahmy, Sinchaisri, & Miyata, 1991). The lack of cross-resistance of other benzoylureas to chlorfluazuron is likely linked to the higher toxicity and delayed excretion of chlorfluazuron (Gazit, Ishaaya, & Perry, 1989; Guyer & Neumann, 1988), but mostly due to the structural nature of this compound. Insect resistance to benzoylureas has been reported to occur due to the overexpression of cytochrome P450 monooxygenases (P450) (Bogwitz et al., 2005) or site mutations in the chitin synthase gene (Douris et al., 2016; Fotakis et al., 2020; Suzuki, Shiotsuki, Jouraku, Miura, & Minakuchi, 2017). In one particular case, resistance of a natural population of *Drosophila melanogaster* to the benzoylurea lufenuron due to the overexpression of a P450 (*Cyp6g1*) has evolved as a result of cross-resistance to chemical compounds *D. melanogaster* commonly encounters in nature (Daborn et al., 2002; Wilson & Cain, 1997). But chlorfluazuron resistance of a highly resistant (resistance ratio > 2,000-fold) strain of *P. xylostella* was not affected when using the synergist piperonyl butoxide (PBO), a P450 inhibitor (QingJun, GuoRen, JianZhou, Xing, & Xiwu, 1998). Although PBO had been shown earlier to reduce the resistance ratio of a chlorfluazuron selected strain of *P. xylostella* (resistance ratio = 50-fold) in 4.3-fold, it was suggested the microsomal enzymes acting on this pesticide might be different from other benzoylurea compounds (Ismail & Wright, 1992). In fact, glutathione-S-transferases were later suggested to play a role in *P. xylostella* resistance to chlorfluazuron (Sonoda & Tsumuki, 2005).

Several candidate SNPs identified showed signals of strong positive selection, supporting the polygenic nature of the resistance of *S. frugiperda* to teflubenzuron. Although resistance of insects to benzoylureas has been associated with site mutations in *chitin synthase* (Douris et al., 2016; Fotakis et al., 2020; Suzuki et al., 2017), the resistant population of *S. frugiperda* analyzed did not display any missense variants in this gene. Missense variants were identified in two other genes. One is up-regulated in a chlorantraniliprole resistant strain of *S. exigua*, the cytochrome P450 monooxygenase *Cyp340AB1* (Wang et al., 2018). The other is the cuticular protein *CPG4855*, a gene that participates in the formation of the larval and pupal endocuticle in *S. exigua* (Liu et al., 2017). Although P450 enzymes have been implicated in the metabolization of several pesticides including benzoylureas, as earlier discussed, we do not believe the resistance mechanism of *S. frugiperda* to teflubenzuron would be associated with a point mutation in the *Cyp340AB1* gene, a P450 belonging to the CYP4 clan of P450. The *Cyp* genes mostly commonly involved in the metabolization of xenobiotics, including insecticides, belongs to the family *Cyp6* of the CYP3 Clan of P450 enzymes (Feyereisen, 2012), as the *Cyp6g1* gene earlier reported in lufenuron-resistant *D. melanogaster* (Daborn et al., 2002; Wilson & Cain, 1997). On the other hand, the site mutation in the cuticular protein CPG4855 could have a close association with the resistance mechanism observed in *S. frugiperda* to teflubenzuron, once RNAi experiments targeting cuticular proteins demonstrated associated these genes with insects resistant to insecticides (Fang et al., 2015; Y. Huang et al., 2018).

The resistance of *S. frugiperda* to teflubenzuron was found to be polygenic, and as such much more likely to involve mechanisms of regulation of gene expression, as reported to other benzoylurea-resistant insects. In fact, several polymorphic SNPs were detected upstream (within 5Kb of the start codon) and downstream (within 5Kb of the stop codon) of gene regions, including intronic regions of genome. These SNP variations might be responsible for modifications leading to regulation of gene expression and protein function. In humans, GWAS analysis demonstrated that more than 90% disease-associated SNPs were located up and downstream (promoter and enhancer regions) gene regions or even in non-coding regions of the genome (Hindorff 2009; Hrdlickova, de Almeida, and Borek, & Withoff, 2014; Ricaño-Ponce & Wijmenga, 2013).

Annotation allowed the identification of several upstream, downstream, and introns variants in genes associated with several biological processes, besides biological regulation and regulation of cellular processes. These annotation results point to that multiple gene interaction and regulation play decisive roles in insecticide resistance.

Thus, the high number of SNPs in genomic regions involved in mechanisms that interfere with the regulation of gene expression of a number of genes involved in processes of metabolization and excretion of xenobiotics are thought to serve as candidate molecular markers for monitoring the polygenic teflubenzuron-resistant phenotypes in the field. These candidates involve the multidrug resistance-associated protein 4 or ATP-binding cassette subfamily C member 4 (ABCC4), a transmembrane protein involved in the efflux of organic compounds from cells (Hardy, Bill, Jawhari, & Rothnie, 2019), including xenobiotics in insects (Labbé, Caveney, & Donly, 2011). ABCCs can be functionally diverse, but they are capable to translocate a range of organic xenobiotics including insecticides, and have been involved in insecticide resistance mechanisms in insects and drug resistance in humans (Dermauw & Van Leeuwen, 2013). Moreover, ABCC transporters act in synergy with glutathione-S-transferases (GSTs) and UDP-glucosyltransferases (UGTs) which are enzymes acting in phase II of detoxification. In humans, the synergy of ABCCs, GSTs and UGTs confer resistance to drugs and carcinogens (Dermauw & Van Leeuwen, 2013).

G-protein coupled receptors (GPCRs) play a central role in cell signaling as receptor of neuromodulators, neurotransmitors, hormones and neuropeptides. GWAS analysis for the identification of SNPs in the teflubenzuron-resistant strain of *S. frugiperda* identified variants in genomic regions that can interfere with gene expression of a GPCR belonging to the *methuselah* (*mth*) subfamily of the *secretin* family. *Methuselah* is a secretin receptor reported to be insect-specific. *Mth* play a role in several biological processes in insects, such as stress response, regulation of fluid and ion secretion, and longevity among others (Araújo, Reis, Rocha, & Aguiar, 2013). An exploration of GPCRs in *Tribolium castaneum* indicated *mth* is also important for larval molt and metamorphosis (Bai, Zhu, Shah, & Palli, 2011). Moreover, *mth* gene mutation led *D. melanogaster* to become tolerant to dichlorvos (Pandey et al., 2015), but insect response to insecticide exposure was also altered with changes in the levels of *mth* expression (Cao et al., 2019; Li et al., 2015; Lucas et al., 2019; Ma, Zhang, You, Zeng, & Gao, 2020). Similarly to the early discussed synergism of ABCCs and phase II conjugation enzymes in insect response to insecticide exposure, *mth* is co-expressed with the phase I detoxification P450 enzymes, contributing to insect resistance to insecticides (Cao et al., 2019; Li et al., 2015).

The selection of a strain of *S. frugiperda* highly resistant to teflubenzuron, the identification of cross-resistance to lufenuron and novaluron, and the use of genome wide association analysis led us to identify several candidate molecular markers for monitoring resistance evolution to benzoylureas. Several SNPs identified in association with the teflubenzuron-resistant strain demonstrates that the polygenic mechanism of resistance selected in the resistant strain of *S. frugiperda* is based on a dense and intricate network of co-expressing genes, of which many are important regulatory genes. The variation in the levels of cross-resistance observed for the different benzoylureas assayed against the teflubenzuron-resistant strain of *S. frugiperda* provides us with additional tools to investigate and understand the particular differences in the target sites of the many structural compounds sharing chitin synthesis inhibition as a mode of action. The well-defined mutation in the *chitin synthase* gene related to target site mutation resistance to benzoylureas in arthropods (Van Leeuwen et al., 2012) and the several molecular marker candidates we added as sources of metabolic resistance to benzoylureas, can be strategically used for the definition of monitoring protocols for future use in the implementation of successful insecticide resistance management strategies.

## Acknowledgements

We thank PROMIP (SISBIO License #40380-5) for helping to collect insect samples. Research carried out using the computational resources of the Center for Mathematical Sciences Applied to Industry (CeMEAI) funded by FAPESP (grant 2013/07375-0)

## Funding

We thank the São Paulo Research Foundation (FAPESP) for the fellowship to ARBN (Grants #2016/09159–0 and #2014/26212–7) and the Young Investigator project to KLSB (#2012/16266-7). We also thank the Brazilian National Council for Scientific and Technological Development (CNPq) (#403851/2013-0) and the Brazilian Insecticide Resistance Action Committee (IRAC-BR) for providing partial financial support for this study. The funding bodies had no role in the study design, data collection, analysis and interpretation, or drafting of the manuscript.

## Contributions

ARBN, FLC, AM and CO conceived the study. ARBN and JGR collected the data. CO and AM provided reagents and sequencing financing. ARBN, VACP and KLSB performed the analysis. ARBN and FLC wrote the main manuscript. All authors contributed to writing and editing the manuscript.

## Notes

### Competing Interest Statement

The authors have declared no competing interest.

## References

Abo-Elghar, G. E., Fujiyoshi, P., & Matsumura, Fd. (2004). Significance of the sulfonylurea receptor (SUR) as the target of diflubenzuron in chitin synthesis inhibition in Drosophila melanogaster and Blattella germanica. Insect Biochemistry and Molecular Biology, 34(8), 743–52. https://doi.org/10.1016/j.ibmb.2004.03.009.

Ahmad, M., Sayyed, A. H., Saleem, M. A., & Ahmad, M. (2008). Evidence for field evolved resistance to newer insecticides in Spodoptera litura (Lepidoptera: Noctuidae) from Pakistan. Crop Protection, 27(10): 1367–72. https://doi.org/10.1016/j.cropro.2008.05.003.

Andow, D. A, & Alstad, D. N. (1998). F2 screen for rare resistance alleles. Journal of Economic Entomology, 91(3), 572–578. http://doi.org/10.1093/jee/91.3.572.

Andrews, Simon, F Krueger, A Seconds-Pichon, F Biggins, and S Wingett. 2015. FastQC. a quality control tool for high throughput sequence data. Babraham Bioinformatics. Babraham Institute. 2015.

Araújo, A R, Reis, M., Rocha, H. & Aguiar, B. (2013). The Drosophila melanogaster methuselah gene: a novel gene with ancient functions. PLoS One, 8(5): e63747. http://doi.org/10.1371/journal.pone.0063747.

Ascher, K. R. S., & Nemny, N. E. (1984). The effect of CME 134 on Spodoptera littoralis eggs and larvae. Phytoparasitica, 12(1), 13–27.

Assefa, F., & Ayalew, D. (2019). Status and control measures of fall armyworm (Spodoptera frugiperda) infestations in maize fields in Ethiopia: a review. Cogent Food & Agriculture, 5, 1641902. doi:10.1080/23311932.2019.1641902.

Bai, H., Zhu, F., Shah, K., & Palli, S. R. (2011). Large-scale RNAi screen of G protein-coupled receptors involved in larval growth, molting and metamorphosis in the red flour beetle. BMC Genomics, 12(388). http://doi.org/10.1186/1471-2164-12-388.

Beeman, R. W. (1982). Recent advances in mode of action of insecticides. Annual Review of Entomology, 27 : 253–281. https://doi.org/10.1146/annurev.en.27.010182.001345.

Bogwitz, M. R., Chung, H., Magoc, L., Rigby, S., Wong, W., O’Keefe, M., … & Daborn P. J. (2005). Cyp12a4 confers lufenuron resistance in a natural population of Drosophila melanogaster. Proceedings of the National Academy of Sciences of the United States of America, 102(36): 12807–12812. https://doi.org/10.1073/pnas.0503709102.

Bolzan, A., Padovez, F. E. O., Nascimento, A. R. B., Kaiser, I. S., Lira, E. C., Amaral, F. S. A., & Omoto, C. (2019). Selection and characterization of the inheritance of resistance of Spodoptera frugiperda (Lepidoptera: Noctuidae) to chlorantraniliprole and cross-resistance to other diamide insecticides. Pest Management Science, 75(10): 2682–2689. https://doi.org/10.1002/ps.5376.

Bourguet, D., Genissel, A., & Raymond, M. (2000). Insecticide resistance and dominance levels. Journal of Economic Entomology, 93(6): 1588–1595. https://doi.org/10.1603/0022-0493-93.6.1588.

Cao, C., Sun, L., Du, H., Moural, T. W., Bai, H., & Liu, P. (2019). Physiological functions of a methuselah-like G protein coupled receptor in Lymantria dispar Linnaeus. Pesticide Biochemistry and Physiology, 160, 1–10.https://doi.org/10.1016/j.pestbp.2019.07.002.

Carvalho, R. A., Omoto, C., Field, L. M., Williamson, M. S., & Bass, C. (2013). Investigating the molecular mechanisms of organophosphate and pyrethroid resistance in the fall armyworm Spodoptera frugiperda. PLoS ONE, 8(4), e62268. https://doi.org/10.1371/journal.pone.0062268.

Cingolani, P., Platts, A., Wang, L., Coon, M., Nguyen, T., Wang, L., … & Ruden, D. M. (2012). A program for annotating and predicting the effects of single nucleotide polymorphisms, SnpEff. Fly, 6(2): 80–92. http://doi.org/10.4161/fly.19695.

Coop, G., Witonsky, D., Rienzo, A., & Pritchard, J. K. (2010). Using environmental correlations to identify loci underlying local adaptation. Genetics, 185(4), 1411–1423. http://doi.org/10.1534/genetics.110.114819.

Cruz, I. (1995). Manejo integrado de pragas de milho com ênfase para o controle biológico.. In: CICLO DE PALESTRAS SOBRE O CONTROLE BIOLÓGICO DE PRAGAS, 4., 1995,

Campinas, SP. Anais. Campinas: SEB/Instituto Biológico, 1995. p.48–92.

Daalen, J. J. van, Meltzer, J., Mulder, R., & Wellinga, K. (1972). A selective insecticide with a novel mode of action. Naturwissenschaften, 59(7), 312–313. https://doi.org/10.1007/BF00593360.

Daborn, P. J., Yen, J. L., Bogwitz, M. R., Le Goff, G., Feil, E., Jeffers, S., … ffrench-Constant, R. H.(2002). A single p450 allele associated with insecticide resistance in Drosophila. Science, 297(5590), 2253–2256.http://doi.org/10.1126/science.1074170.

Danecek, P., Auton, A., Abecasis, G., Albers, C. A., Banks, E., DePristo, M. A., & 1000 Genomes Project Analysis Group. (2011). The variant call format and VCFtools. Bioinformatics, 27(15), 2156–2158. http://doi.org/10.1093/bioinformatics/btr330.

Dermauw, W., & Van Leeuwen, T. (2013). The ABC gene family in arthropods: comparative genomics and role in insecticide transport and resistance. Insect Biochemistry and Molecular Biology, 45, 89–110. http://doi.org/10.1016/j.ibmb.2013.11.001.Douris,

V., Steinbach, D., Panteleri, R., Livadaras, I., Pickett, J. A., Van Leeuwen, T., Nauen, R., & Vontas, J. (2016). Resistance mutation conserved between insects and mites unravels the benzoylurea insecticide mode of action on chitin biosynthesis. Proceedings of the National Academy of Sciences of the United States of America, 113(51), 14692–14697. https://doi.org/10.1073/pnas.1618258113.

Doyle, J. J., Doyle, J. L., & Hortorium, L. H. B. (1990). Isolation of plant DNA from fresh tissue. Focus, 12(1), 13–15.

Fahmy, A. R., Sinchaisri, N., & Miyata, T. (1991). Development of chlorfluazuron resistance and pattern of cross-resistance in the diamondback moth, Plutella xylostella. Journal of Pesticide Science, 16(4): 665–672. https://doi.org/10.1584/jpestics.16.665.

Fang, F., Wang, W., Zhang, D., Lv, Y., Zhou, D., Ma, L., … Zhu, C. (2015). The cuticle proteins: a putative role for deltamethrin resistance in Culex pipiens pallens. Parasitology Research, 114(12), 4421–4429. https://doi.org/10.1007/s00436-015-4683-9.

Farias, J. R., Horikoshi, R. J., Santos, A. C., & Omoto, C. (2014). Geographical and temporal variability in susceptibility to Cry1F toxin from Bacillus thuringiensis in Spodoptera frugiperda (Lepidoptera: Noctuidae) populations in Brazil. Journal of Economic Entomology, 107(6), 2182–2189. https://doi.org/10.1603/EC14190.

Feyereisen, R. (2012). Insect CYP Genes and P450 enzymes. Insect Molecular Biology and Biochemistry, 236–316. https://doi.org/10.1016/B978-0-12-384747-8.10008-X.

Ffrench-Constant, R. H. (2013). The molecular genetics of insecticide resistance. Genetics, 194(4), 807–815. http://doi.org/10.1534/genetics.112.141895.

Finney, D. J. (1949). The adjustment for a natural response rate in probit analysis. Annals of Applied Biology, 36(2), 187–195. https://doi.org/10.1111/j.1744-7348.1949.tb06408.x.

Fotakis, E. A., Mastrantonio, V. Grigoraki, L., Porretta, D., Puggioli, A., Chaskopoulou, A., … Vontas, J. (2020). Identification and detection of a novel point mutation in the Chitin Synthase gene of Culex pipiens associated with diflubenzuron resistance. PLoS Neglected Tropical Diseases, 14(5), 1–10. https://doi.org/10.1371/journal.pntd.0008284.

Bueno, R. C. O. F., Bueno, A. F., Moscardi, F., Parra, J. R. P., Hoffman-Campo, C. B. (2011). Lepidopteran larva consumption of soybean foliage: basis for developing multiple-species economic thresholds for pest management decisions. Pest Management Science, 67(2), 170–174. https://doi.org/10.1002/ps.2047.

Garrison, E., & Marth, G. (2012). Haplotype-based variant detection from short-read sequencing. ArXiv Preprint. Available from: https://arXiv.org/abs/1207.3907.

Gautier, M. (2015). Genome-wide scan for adaptive divergence and association with population-specific covariates. Genetics, 201(4), 1555–1579. https://doi.org/10.1534/genetics.115.181453.

Gazit, Y., Ishaaya, I., & Perry, A. S. (1989). Detoxification and synergism of diflubenzuron and chlorfluazuron in the red flour beetle Tribolium castaneum. Pesticide Biochemistry and Physiology, 34(2), 103–110. doi: https://doi.org/10.1016/0048-3575(89)90147-8.

Georghiou, G. P. (2012). Pest Resistance to Pesticides. Springer Science & Business Media.

Goergen, G., Kumar, P. L., Sankung, S. B., Togola, A, & Samò, M. (2016). First report of outbreaks of the fall armyworm Spodoptera frugiperda (JE Smith) (Lepidoptera, Noctuidae), a new alien invasive pest in West and Central Africa. PLoS ONE, 11(10), e0165632. https://doi.org/10.1371/journal.pone.0165632.

Günther, T. & Coop, G. (2013). Robust identification of local adaptation from allele frequencies. Genetics, 195(1), 205–220. https://doi.org/10.1534/genetics.113.152462.

Guyer, W., & Neumann R. (1988). Activity and fate of chlorfluazuron and diflubenzuron in the larvae of Spodoptera littoralis and Heliothis virescens. Pesticide Biochemistry and Physiology, 30(2): 166–177. https://doi.org/10.1016/0048-3575(88)90050-8.

Hardy, D., Bill, R. M., Jawhari, A, & Rothnie, A. J. (2019). Functional expression of multidrug resistance protein 4 MRP4/ABCC4. SLAS Discovery, 24(10), 1000–1008. https://doi.org/10.1177/2472555219867070.

Hindorff, L. A., Sethupathy, P. Junkins, H. A., Ramos, E. M., Mehta, J. P., Collins, F. S., & Manolio, T. A. (2009). Potential etiologic and functional implications of genome-wide association loci for human diseases and traits. Proceedings of the National Academy of Sciences of the United States of America, 106(23), 9362–9367. https://doi.org/10.1073/pnas.0903103106

Hrdlickova, B., Almeida, R. C., Borek, Z., & Withoff, S. (2014). Genetic variation in the non-coding genome: Involvement of micro-RNAs and long non-coding RNAs in disease. Biochimica et Biophysica Acta, 1842, 1910–1922. https://doi.org/10.1016/j.bbadis.2014.03.011.

Huang, F., Buschman, L. L., Higgins, R. A., & McGaughey, W. H. (1999). Inheritance of resistance to Bacillus thuringiensis toxin (Dipel ES) in the european corn borer. Science, 284(5416), 965–967. http://doi.org/10.1126/science.284.5416.965.

Huang, Y., Guo, Q., Sun, X., Zhang, C., Xu, N., Xu, Y., … Shen, B. (2018). Culex pipiens pallens cuticular protein cplcg5 participates in pyrethroid resistance by forming a rigid matrix. Parasites & Vectors, 11(6), 1–10. https://doi.org/10.1186/s13071-017-2567-9.

Iqbal, M., & Wright, D. J. (1997). Evaluation of resistance, cross-resistance and synergism of abamectin and teflubenzuron in a multi-resistant field population of Plutella xylostella (Lepidoptera: Plutellidae). Bulletin of Entomological Research, 87(5), 481–486. https://doi.org/10.1017/S0007485300041341.

Ismail, F., & Wright, D. J. (1992). Synergism of teflubenzuron and chlorfluazuron in an acylurea-resistant field strain of Plutella xylostella L. (Lepidoptera: Yponomeutidae). Pesticide Science, 34(3): 221–226. http://doi.org/10.1002/ps.2780340306.

Kasten Junior, P., Precetti, A. A. C. M., & Parra, J. R. P. (1978). Dados biológicos comparativos de. Spodoptera frugiperda (J.E. Smith, 1797) em duas dietas artificiais e substrato natural. Brazilian Journal of Agriculture, 53(1): 68–78.

Kofler, R., Orozco-terWengel, P., Maio, N., Pandey, R. V., Nolte, V., Futschik, A., & Schlötterer, C. 2011. PoPoolation: a toolbox for population genetic analysis of next generation sequencing data from pooled individuals. PLoS ONE, 6(1), e15925.http://doi.org/10.1371/journal.pone.0015925.

Labbé, R, Caveney, S., & Donly, C. (2011). Genetic analysis of the xenobiotic resistance-associated ABC gene subfamilies of the Lepidoptera. Insect Molecular Biology, 20(2), 243–256. https://doi.org/10.1111/j.1365-2583.2010.01064.x.

Leeuwen, T. Van, Demaeght, P., Osborne, E. J., Dermauw, W., Gohlke, S., Nauen, R., … Clark, R. M. (2012). Population bulk segregant mapping uncovers resistance mutations and the mode of action of a chitin synthesis inhibitor in arthropods.” Proceedings of the National Academy of Sciences of the United States of America, 109(12), 4407–4412. https://doi.org/10.1073/pnas.1200068109.

Lenormand, T., & Raymond, M. (1998). Resistance management: the stable zone strategy. Proceedings of the Royal Society B: Biological Sciences, 265(1409), 1985–1990. https://doi.org/10.1098/rspb.1998.0529.

Lewis, J. A., & Finney, D. (1971). Probit Analysis (3rd ed.). Cambridge, UK: Cambridge University Press.

Li, H. (2011). A statistical framework for SNP calling, mutation discovery, association mapping and population genetical parameter estimation from sequencing data. Bioinformatics, 27(21): 2987–2993. https://doi.org/10.1093/bioinformatics/btr509.

Li, H., & Durbin, R. (2010). Fast and accurate long-read alignment with Burrows-Wheeler transform. Bioinformatics, 26(5): 589–595. http://doi.org/10.1093/bioinformatics/btp698.

Li, T, Cao, C., Yang, T., Zhang, L., He, L., Xi, Z., & Bian, G. (2015). A G-protein-coupled receptor regulation pathway in cytochrome P450-mediated permethrin-resistance in mosquitoes, Culex quinquefasciatus. Scientific Report, 5:17772. http://doi.org/10.1038/srep17772.

Lin, J. G., Hung, C. F., & Sun, C. N. (1989). Teflubenzuron resistance and microsomal monooxygenases in larvae of the diamondback moth. Pesticide Biochemistry and Physiology, 35(1): 20–25. https://doi.org/10.1016/0048-3575(89)90098-9.

Lira, E. C., Bolzan, A., Nascimento, A. R. B., Amaral, F. S. A., Kanno, R. H., Kaiser, I. S., & Omoto, C. 2020. Resistance of Spodoptera frugiperda (Lepidoptera: Noctuidae) to spinetoram: inheritance and cross-resistance to spinosad. Pest Management Science, 76(8): 2674–2680. https://doi.org/10.1002/ps.5812.

Liu, S., Hafeez, M., Zhang, X., Dawar, F. U., Guo, J., … Wang, M. (2017). Isolation and functional identification of three cuticle protein genes during metamorphosis of the beet armyworm, Spodoptera exigua. Scientific Reports, 7: 16061. https://doi.org/10.1038/s41598-017-16435-w.

Lucas, E. R., Rockett, K. A., Lynd, A., Essandoh, J., Grisales, N., Kemei, B., … Donnely, M. J. (2019). A high throughput multi-locus insecticide resistance marker panel for tracking resistance emergence and spread in Anopheles gambiae. Scientific Reports, 9: 13335. https://doi.org/10.1038/s41598-019-49892-6.

Ma, Z., Zhang, Y., You, C., Zeng, X., & Gao, X. (2020). The role of G protein-coupled receptor-related genes in cytochrome P450-mediated resistance of the house fly, Musca domestica (Diptera: Muscidae), to imidacloprid. Insect Molecular Biology, 29(1), 92–103. https://doi.org/10.1111/imb.12615.

MAPA. (2020). Sistema de Agrotóxicos Fitossanitários. Agrofit. 2020. http://agrofit.agricultura.gov.br/agrofit_cons/principal_agrofit_cons.

Merzendorfer, H. (2006). Insect chitin synthases: a review. Journal of Comparative Physiology B: Biochemical, Systemic, and Environmental Physiology, 176, 1–15. https://doi.org/10.1007/s00360-005-0005-3.

Merzendorfer, H., & Zimoch, L. (2003). Chitin metabolism in insects: structure, function and regulation of chitin synthases and chitinases. The Journal of Experimental Biology, 206, 4393–4412. https://doi.org/10.1242/jeb.00709.

Meyer, F., Flötenmeyer, M. & Moussian, B. (2013). The sulfonylurea receptor Sur is dispensable for chitin synthesis in Drosophila melanogaster embryos. Pest Management Science, 69(10): 1136–1140. https://doi.org/10.1002/ps.3476.

Müller, H., & Jimenez, R. (2017). VCF.Filter: interactive prioritization of disease-linked genetic variants from sequencing data. Nucleic Acids Research, 45, W567–W572. https://doi.org/10.1093/nar/gkx425.

Nascimento, A. R. B., Farias, J. R., Bernardi, D., Horikoshi, R. J., & Omoto, C. (2016). Genetic basis of Spodoptera frugiperda (Lepidoptera: Noctuidae) resistance to the chitin synthesis inhibitor lufenuron. Pest Management Science, 72(4): 810–815. https://doi.org/10.1002/ps.4057.

Nascimento, A. R. B., Fresia, P., Cônsoli, F. L., & Omoto, C. (2015). Comparative transcriptome analysis of lufenuron-resistant and susceptible strains of Spodoptera frugiperda (Lepidoptera: Noctuidae). BMC Genomics, 16(1). https://doi.org/10.1186/s12864-015-2183-z.

Nicholson, G., Smith, A. V., Jonsson, F., Gustafsson, O., Stefansson, K., & Donnelly, P. (2002). Assessing population differentiation and isolation from single-nucleotide polymorphism data. Journal of The Royal Statistical Society Series B (Statistical Methodology), 64(4), 695–715. https://doi.org/10.1111/1467-9868.00357.

Oberlander, H., & Silhacek, D. L. (1998). Mode of action of insect growth regulators in lepidopteran tissue culture. Pesticide Science, 54, 300–322. https://doi.org/10.1002/(SICI)1096-9063(1998110)54:3%3C300::AID-PS830%3E3.0.CO;2-8.

Okuma, D. M., Bernardi, D., Horikoshi, R. J., Bernardi, O., Silva, A. P., & Omoto, C. (2018). Inheritance and fitness costs of Spodoptera frugiperda (lepidoptera: noctuidae) resistance to spinosad in Brazil. Pest Management Science, 74(6): 1441–1448. https://doi.org/10.1002/ps.4829.

Pandey, A., Khatoon, R., Saini, S., Vimal, D., Patel, D. K., Narayan, G., & Chowdhuri, D. K. (2015). Efficacy of Methuselah gene mutation toward tolerance of dichlorvos exposure in Drosophila melanogaster. Free Radical Biology and Medicine, 84, 54–65. https://doi.org/10.1016/j.freeradbiomed.2015.02.025.

Perng, F. S., Yao, M. C., Hung, C. F., & Sun, C. N. (1988). Teflubenzuron resistance in diamondback moth (Lepidoptera: Plutellidae). Journal of Economic Entomology, 81(5), 1277–1282. http://dx.doi.org/10.1093/jee/81.5.1277.

Post, L. C., & Vincent, W. R. (1973). A new insecticide inhibits chitin synthesis. Naturwissenschaften 60, 431–432. https://doi.org/10.1007/BF00623561.

QingJun, W., GuoRen, Z., JianZhou, Z., Xing, Z, & Xiwu, G. (1998). Resistance selection of Plutella xylostella (L.) by chlorfluazuron and patterns of cross-resistance. Acta Entomologica Sinica, 41, 34–41.

Ricaño-Ponce, I. & Wijmenga, C. (2013). Mapping of immune-mediated disease genes. Annual Reviews of Genomics and Human Genetics, 14, 325–353. https://doi.org/10.1146/annurev-genom-091212-153450.

Robertson, J. L., Russell, R. M., Preisler, H. K., & Savin, N. E. (2007). Bioassays with Arthropods (2nd ed.). Boca Raton, FL: CRC Press.

Roush, R. T., & Daly, J. C. (1990). The role of population genetics in resistance research and management. In R. T. Roush & B. E. Tabashnik (Eds.), Pesticide Resistance in Arthropods (pp. 97–152). Boston, MA: Springer.

Santos, W. J. (2007). Manejo das pragas do algodão com destaque para o cerrado brasileiro. In E. C. Freire (Ed.), Algodão No Cerrado Do Brasil (pp. 403–478).

Brasília, BR: ABRAPASchmidt, F. B. (2002). Linha básica de suscetibilidade de Spodoptera frugiperda (Lepidoptera: Noctuidae) a lufenuron na cultura do milho (PhD thesis). University of São Paulo, Brazil.

Sokal, R. R., & Rohlf, F. J. (2012). Biometry: the principles and practice of statistics in biological research (4th ed.). New York, USA:W. H. Freeman.

Sonoda, S., & Tsumuki, H. (2005). Studies on glutathione s-transferase gene involved in chlorfluazuron resistance of the diamondback moth, Plutella xylostella L.(Lepidoptera: Yponomeutidae). Pesticide Biochemistry and Physiology, 82(1), 94–101. https://doi.org/10.1016/j.pestbp.2005.01.003.

Stone, B. F. (1968). A formula for determining degree of dominance in cases of monofactorial inheritance of resistance to chemicals. Bulletin of the World Health Organization, 38(2): 325–326.

Suzuki, Y., Shiotsuki, T. Jouraku, A., Miura, K., & Minakuchi, C. 2017. Benzoylurea resistance in western flower thrips Frankliniella occidentalis (Thysanoptera: Thripidae): the presence of a point mutation in chitin synthase 1. Journal of Pesticide Science, 42(3), 93–96. https://doi.org/10.1584/jpestics.D17-023.

Tsukamoto, M. (1983). Methods of genetic analysis of insecticide resistance. In G. P. Georghiou & T. Saito, Pest resistance to pesticide (pp. 71–98). New York, NY: Plenum.

Wang, X., Chen, Y., Gong, C., Yao, X., Jiang, C., Yang, Q. (2018). Molecular identification of four novel cytochrome p450 genes related to the development of resistance of Spodoptera exigua (Lepidoptera: Noctuidae) to chlorantraniliprole Pest Management Science, 74(8), 1938–1952. https://doi.org/10.1002/ps.4898.

Wilson, T. G., & Cain, J. W. (1997). Resistance to the insecticides lufenuron and propoxur in natural populations of Drosophila melanogaster (Diptera: Drosophilidae). Journal of Economic Entomology, 90(5): 1131–36. https://doi.org/10.1093/jee/90.5.1131.

Ya, I. I., & Klein, M. (1990). Response of susceptible laboratory and resistant field strains of Spodoptera littoralis (Lepidoptera: Noctuidae) to teflubenzuron. Journal of Economic Entomology, 83(1), 59–62. https://doi.org/10.1093/jee/83.1.59.

Yu, S. J. (2014). The Toxicology and Biochemistry of Insecticides. (2nd ed.). Boca Raton, FL: CRC Press.

