## Supplementary material for "There is more than chitin synthase in insect resistance to benzoylureas: Molecular markers associated with teflubenzuron resistance in *Spodoptera frugiperda*": Table S1

**Table S.1.** Summary of Miseq sequence using pooled *S.frugiperda* strains resistant and susceptible to teflubenzuron

| | |
|---|---|
| Raw total sequences | 220,088,414 |
| Reads mapped | 159,842,968 |
| Average length | 229 bp |
| Maximum length | 300 bp |
| Insert size average | 429,7 |

| | **Nº of Reads** | **Mapping Ratio** |
|---|---|---|
| Tef-rr | 21,729,006 | 65.87 |
| Sf-ss | 21,758,520 | 81.81 |
| BC-random | 85,699,458 | 72.62 |
| BC-selected | 90,901,430 | 72.05 |
