## Supplementary material for "There is more than chitin synthase in insect resistance to benzoylureas: Molecular markers associated with teflubenzuron resistance in *Spodoptera frugiperda*": Table S3

Table S3 - Sliding-indow analyses of nucleotide diversity (Tajima's π)

| Scaffold | Window | Sf-ss | Tef-rr | BC-random | BC-selected |
|---|---|---|---|---|---|
| pseudoscaff_261 | 12500 | 0.001850946 | 0.028905 | 0.026178463 | 0.024963915 |
| pseudoscaff_261 | 17500 | 0.001161453 | 0.003924 | 0.034124115 | 0.03096947 |
| pseudoscaff_261 | 27500 | 0.001021823 | 0.024558 | 0.025610933 | 0.024897644 |
| pseudoscaff_261 | 82500 | 0.000291445 | 0.013632 | 0.017895837 | 0.019967977 |
| pseudoscaff_261 | 92500 | 0.000120602 | 0.02657 | 0.020383592 | 0.02411522 |
| pseudoscaff_261 | 97500 | 0 | 0.007951 | 0.021515505 | 0.027161495 |
| pseudoscaff_261 | 102500 | 0.001741561 | 0.01862 | 0.028688717 | 0.027773691 |
| pseudoscaff_261 | 107500 | 0.000083417 | 0.02511 | 0.019860435 | 0.02139541 |
| pseudoscaff_261 | 112500 | 0.000946754 | 0.021892 | 0.02139615 | 0.019496919 |
| pseudoscaff_261 | 122500 | 0 | 0.008005 | 0.009722198 | 0.011429688 |
| pseudoscaff_261 | 132500 | 0.000228245 | 0.019515 | 0.015454657 | 0.013125341 |
| pseudoscaff_261 | 137500 | 0.000619605 | 0.008334 | 0.010450807 | 0.008416742 |
| pseudoscaff_261 | 142500 | 0.000652784 | 0.002775 | 0.003291907 | 0.003329372 |
| pseudoscaff_261 | 152500 | 0.000886673 | 0.029196 | 0.011668128 | 0.010321929 |
| pseudoscaff_261 | 157500 | 0 | 0.003527 | 0.005574128 | 0.005225694 |
| pseudoscaff_261 | 167500 | 0.001271646 | 0.009667 | 0.010012068 | 0.006534525 |
| pseudoscaff_261 | 172500 | 0.001009306 | 0.001518 | 0.012939722 | 0.01511417 |
| pseudoscaff_261 | 177500 | 0.00159054 | 0.003721 | 0.006584694 | 0.006464144 |
| pseudoscaff_261 | 182500 | 0.000267396 | 0.00484 | 0.004616512 | 0.004556662 |
| pseudoscaff_261 | 187500 | 0.000346169 | 0.00156 | 0.004865013 | 0.00408352 |
| pseudoscaff_261 | 192500 | 0 | 0.005214 | 0.00885591 | 0.008770057 |
| pseudoscaff_261 | 202500 | 0.000431456 | 0.005309 | 0.005424204 | 0.005274645 |
| pseudoscaff_261 | 207500 | 0.000877026 | 0.005839 | 0.004811499 | 0.004992086 |
| pseudoscaff_261 | 212500 | 0.000163128 | 0.004144 | 0.004231082 | 0.004307533 |
| pseudoscaff_261 | 217500 | 0.001976285 | 0.003444 | 0.006781643 | 0.006941877 |
| pseudoscaff_261 | 222500 | 0.000298042 | 0.004785 | 0.007868585 | 0.008094978 |
| pseudoscaff_261 | 232500 | 0.000097317 | 0.024457 | 0.007785622 | 0.005894883 |
| pseudoscaff_261 | 237500 | 0.000217537 | 0.022954 | 0.012210864 | 0.010281368 |
| pseudoscaff_261 | 262500 | 0.00006715 | 0.002405 | 0.004068532 | 0.003725351 |
| pseudoscaff_261 | 297500 | 0.001239245 | 0.01477 | 0.00577938 | 0.006103144 |
| pseudoscaff_261 | 302500 | 0.001146498 | 0.001051 | 0.006893342 | 0.008203996 |
| pseudoscaff_261 | 307500 | 0.001686698 | 0.005464 | 0.008937472 | 0.008829977 |
| pseudoscaff_261 | 312500 | 0 | 0.006231 | 0.015280983 | 0.012117543 |
| pseudoscaff_261 | 322500 | 0.001701671 | 0.007782 | 0.008579357 | 0.006979378 |
| pseudoscaff_261 | 327500 | 0 | 0.001431 | 0.004873812 | 0.00475507 |
| pseudoscaff_261 | 332500 | 0.000219986 | 0.003587 | 0.008192954 | 0.008141891 |
| pseudoscaff_261 | 337500 | 0.000212547 | 0.005745 | 0.004611093 | 0.006481296 |
| pseudoscaff_261 | 342500 | 0.000274155 | 0.007182 | 0.012384408 | 0.010742302 |
| pseudoscaff_261 | 352500 | 0.000194542 | 0.00446 | 0.009582769 | 0.007288275 |
| pseudoscaff_261 | 357500 | 0.000172089 | 0.009222 | 0.007998405 | 0.008041443 |
| pseudoscaff_2242 | 2500 | 0.001507146 | 0.019265 | 0.029141628 | 0.02799427 |
| pseudoscaff_2242 | 7500 | 0.000475145 | 0.021839 | 0.029037565 | 0.032781722 |
| pseudoscaff_3071 | 7500 | 0.000589359 | 0.007592 | 0.004631239 | 0.00469371 |
| pseudoscaff_3071 | 12500 | 0.000521893 | 0.003937 | 0.001845514 | 0.001383965 |
| pseudoscaff_3071 | 17500 | 0.000744736 | 0.004326 | 0.00754644 | 0.00789025 |
| pseudoscaff_3608 | 7500 | 0.001135825 | 0.008063 | 0.006019172 | 0.005506454 |
| pseudoscaff_3608 | 22500 | 0.001740075 | 0.005143 | 0.007355219 | 0.00502385 |
| pseudoscaff_3023 | 2500 | 0.000807968 | 0 | 0.002281265 | 0.003332035 |

| | | | | | |
|---|---|---|---|---|---|
| pseudoscaff_3023 | 7500 | 0.000418683 | 0 | 0.002797939 | 0.002829809 |
| pseudoscaff_3023 | 12500 | 0.001703787 | 0 | 0.001969738 | 0.002141708 |
| pseudoscaff_3023 | 17500 | 0.001570731 | 0 | 0.00263224 | 0.002880853 |
| pseudoscaff_3023 | 22500 | 0.000575661 | 0.000466 | 0.002894415 | 0.00278105 |
| pseudoscaff_3023 | 42500 | 0.000170876 | 0 | 0.001848451 | 0.001917411 |
| pseudoscaff_3023 | 57500 | 0.000428839 | 0 | 0.000674803 | 0.000276751 |
| pseudoscaff_3023 | 62500 | 0.000966499 | 0.001399 | 0.003557895 | 0.004452885 |
| pseudoscaff_137 | 2500 | 0.001061735 | 0.028464 | 0.034227072 | 0.035117633 |
| pseudoscaff_2285 | 147500 | 0.00191271 | 0.017134 | 0.015804722 | 0.01623621 |
| pseudoscaff_3718 | 7500 | 0 | 0.018519 | 0.018402744 | 0.018205146 |
| pseudoscaff_3718 | 12500 | 0.000411278 | 0.001488 | 0.00499865 | 0.004946139 |
| pseudoscaff_3718 | 17500 | 0 | 0.006193 | 0.007579648 | 0.007642752 |
| pseudoscaff_3718 | 22500 | 0.000060007 | 0.006837 | 0.012202545 | 0.012184684 |
| pseudoscaff_3718 | 32500 | 0 | 0.010354 | 0.007808002 | 0.01282489 |
| pseudoscaff_3718 | 37500 | 0 | 0.010574 | 0.013420133 | 0.013287124 |
| pseudoscaff_3718 | 42500 | 0 | 0.002597 | 0.01579806 | 0.015463526 |
| pseudoscaff_3718 | 47500 | 0.000524898 | 0.013785 | 0.024020143 | 0.0225652 |
| pseudoscaff_3718 | 52500 | 0.000132252 | 0.016967 | 0.01645493 | 0.015759499 |
| pseudoscaff_3718 | 62500 | 0 | 0.006432 | 0.008868561 | 0.006432061 |
| pseudoscaff_3718 | 72500 | 0 | 0.011782 | 0.014734123 | 0.014779859 |
| pseudoscaff_3718 | 87500 | 0 | 0.003347 | 0.010025123 | 0.011212132 |
| pseudoscaff_3718 | 92500 | 0 | 0.008465 | 0.01570152 | 0.014117616 |
| pseudoscaff_1511 | 32500 | 0.000495269 | 0.015459 | 0.007286657 | 0.009521276 |
| pseudoscaff_1511 | 42500 | 0.001419755 | 0.011588 | 0.008925953 | 0.011718227 |
| pseudoscaff_1511 | 47500 | 0 | 0.009699 | 0.007523224 | 0.006272074 |
| pseudoscaff_1511 | 57500 | 0 | 0.001221 | 0.003582806 | 0.003155271 |
| pseudoscaff_1511 | 62500 | 0 | 0.00234 | 0.003638964 | 0.003389118 |
| pseudoscaff_1511 | 72500 | 0 | 0.021376 | 0.008310119 | 0.007595464 |
| pseudoscaff_1511 | 82500 | 0 | 0.010975 | 0.01035051 | 0.009236815 |
| pseudoscaff_1511 | 87500 | 0.000217373 | 0.011522 | 0.00507329 | 0.006704579 |
| pseudoscaff_1511 | 92500 | 0.001256238 | 0.012798 | 0.009900962 | 0.014179434 |
| pseudoscaff_3753 | 2500 | 0.001040197 | 0.001524 | 0.006623733 | 0.009639073 |
| pseudoscaff_3753 | 7500 | 0.000164383 | 0.000328 | 0.009371507 | 0.01019925 |
| pseudoscaff_3095 | 32500 | 0.000290123 | 0.023172 | 0.033220807 | 0.036411869 |
| pseudoscaff_2560 | 2500 | 0.000057942 | 0.004783 | 0.004972567 | 0.005279494 |
| pseudoscaff_2560 | 7500 | 0.000075717 | 0.001588 | 0.008986909 | 0.009453431 |
| pseudoscaff_2560 | 12500 | 0 | 0.010405 | 0.007057806 | 0.00834454 |
| pseudoscaff_2560 | 17500 | 0 | 0.01019 | 0.003208671 | 0.003609847 |
| pseudoscaff_2560 | 22500 | 0.000108083 | 0.002016 | 0.001497221 | 0.001434207 |
| pseudoscaff_2560 | 27500 | 0 | 0.00261 | 0.002623856 | 0.003534805 |
| pseudoscaff_2560 | 32500 | 0 | 0.006656 | 0.007709648 | 0.009235084 |
| pseudoscaff_2560 | 37500 | 0 | 0.024486 | 0.017471546 | 0.020822027 |
| pseudoscaff_754 | 17500 | 0 | 0.003077 | 0.003987226 | 0.003981335 |
| pseudoscaff_754 | 22500 | 0 | 0.009481 | 0.017242775 | 0.016899146 |
| pseudoscaff_754 | 37500 | 0 | 0.031206 | 0.040759167 | 0.038053369 |
| pseudoscaff_2033 | 17500 | 0.000095873 | 0.027935 | 0.025665236 | 0.029295657 |
| pseudoscaff_3069 | 2500 | 0 | 0.002642 | 0.002100619 | 0.003505926 |
| pseudoscaff_3069 | 7500 | 0 | 0.002505 | 0.006466455 | 0.006258772 |
| pseudoscaff_3069 | 12500 | 0.000731857 | 0.005476 | 0.002941042 | 0.00449481 |
| pseudoscaff_3069 | 17500 | 0 | 0.014397 | 0.004421595 | 0.005586153 |

| | | | | | |
|---|---|---|---|---|---|
| pseudoscaff_3069 | 27500 | 0 | 0.002894 | 0.0016046 | 0.001887868 |
| pseudoscaff_3069 | 32500 | 0.000169307 | 0.001648 | 0.002793373 | 0.003084351 |
| pseudoscaff_3069 | 37500 | 0.0012158 | 0.003136 | 0.001712138 | 0.002788026 |
| pseudoscaff_3069 | 42500 | 0 | 0.00208 | 0.001837669 | 0.001659955 |
| pseudoscaff_3069 | 47500 | 0 | 0.004252 | 0.002344599 | 0.002779203 |
| pseudoscaff_3069 | 52500 | 0.000190771 | 0.004285 | 0.002868888 | 0.002849322 |
| pseudoscaff_1226 | 7500 | 0.00162047 | 0.009983 | 0.016287979 | 0.016357593 |
| pseudoscaff_2186 | 22500 | 0.001372299 | 0.045085 | 0.039582134 | 0.041868015 |
| pseudoscaff_2186 | 32500 | 0.001496906 | 0.030838 | 0.032402819 | 0.03406323 |
| pseudoscaff_141 | 42500 | 0.00006462 | 0.017313 | 0.016283836 | 0.018334916 |
| pseudoscaff_141 | 47500 | 0.001769422 | 0.029917 | 0.031084461 | 0.030010124 |
| pseudoscaff_141 | 52500 | 0.000058518 | 0.032429 | 0.026424606 | 0.029319972 |
| pseudoscaff_141 | 62500 | 0.000755126 | 0.036214 | 0.023472292 | 0.022970256 |
| pseudoscaff_141 | 72500 | 0 | 0.029806 | 0.034144781 | 0.030705233 |
| pseudoscaff_141 | 77500 | 0.001618492 | 0.036154 | 0.037804342 | 0.038303427 |
| pseudoscaff_141 | 82500 | 0.000706071 | 0.041067 | 0.042158605 | 0.047742865 |
| pseudoscaff_287 | 2500 | 0.000213294 | 0.012702 | 0.013147851 | 0.012694389 |
| pseudoscaff_287 | 17500 | 0.000202452 | 0.017115 | 0.020700154 | 0.0184711 |
| pseudoscaff_287 | 32500 | 0 | 0.017424 | 0.019400273 | 0.01645001 |
| pseudoscaff_287 | 37500 | 0.000840181 | 0.019713 | 0.022113508 | 0.020285651 |
| pseudoscaff_287 | 42500 | 0.000072382 | 0.025839 | 0.019524223 | 0.01856908 |
| pseudoscaff_287 | 47500 | 0.000094091 | 0.026063 | 0.019544815 | 0.019593649 |
| pseudoscaff_287 | 57500 | 0 | 0.026133 | 0.014250735 | 0.012857253 |
| pseudoscaff_287 | 62500 | 0.000073562 | 0.014153 | 0.023485081 | 0.021925986 |
| pseudoscaff_287 | 67500 | 0 | 0.01164 | 0.00929519 | 0.007753825 |
| pseudoscaff_287 | 77500 | 0.00020687 | 0.003475 | 0.00210669 | 0.00225732 |
| pseudoscaff_2338 | 27500 | 0.001637758 | 0.005505 | 0.003200508 | 0.003616528 |
| pseudoscaff_3995 | 2500 | 0.000117055 | 0.028496 | 0.019048794 | 0.018500659 |
| pseudoscaff_3995 | 7500 | 0.000580681 | 0.019142 | 0.014389497 | 0.016464405 |
| pseudoscaff_528 | 12500 | 0.001919716 | 0.006366 | 0.0071959 | 0.014115902 |
| pseudoscaff_891 | 2500 | 0.000848006 | 0.010538 | 0.011637561 | 0.011914873 |
| pseudoscaff_891 | 7500 | 0.001466052 | 0.013364 | 0.010094402 | 0.009969307 |
| pseudoscaff_891 | 12500 | 0.001468769 | 0.020228 | 0.013317604 | 0.013391577 |
| pseudoscaff_476 | 7500 | 0.000819973 | 0.012634 | 0.019289622 | 0.019236753 |
| pseudoscaff_476 | 12500 | 0.000140296 | 0.026299 | 0.024642968 | 0.020481466 |
| pseudoscaff_3032 | 7500 | 0.000367529 | 0.02255 | 0.033601691 | 0.034187067 |
| pseudoscaff_3032 | 12500 | 0.000095146 | 0.01771 | 0.020881373 | 0.021055333 |
| pseudoscaff_3032 | 17500 | 0.000173239 | 0.025562 | 0.032299356 | 0.031618719 |
| pseudoscaff_782 | 2500 | 0.000943185 | 0.025777 | 0.02875756 | 0.03236717 |
| pseudoscaff_3620 | 32500 | 0.001403581 | 0.0247 | 0.027958142 | 0.027500274 |
| pseudoscaff_3620 | 37500 | 0 | 0.015483 | 0.014171405 | 0.014183742 |
| pseudoscaff_3620 | 52500 | 0.001263176 | 0.024609 | 0.036189345 | 0.035714189 |
| pseudoscaff_2772 | 462500 | 0.001946466 | 0.003089 | 0.00399391 | 0.00425532 |
| pseudoscaff_2772 | 467500 | 0.001752542 | 0.006419 | 0.002973936 | 0.002728852 |
| pseudoscaff_325 | 17500 | 0.00045876 | 0.033025 | 0.038472462 | 0.031349613 |
| pseudoscaff_3422 | 7500 | 0.001466575 | 0.03492 | 0.031619675 | 0.034561607 |
| pseudoscaff_2630 | 7500 | 0.00049398 | 0.046686 | 0.030824602 | 0.034306336 |
| pseudoscaff_2630 | 12500 | 0.000754707 | 0.026168 | 0.025861429 | 0.026390376 |
| pseudoscaff_2630 | 17500 | 0.000953669 | 0.033973 | 0.031912314 | 0.033149803 |
| pseudoscaff_2630 | 22500 | 0.000128585 | 0.017684 | 0.02000143 | 0.018730468 |

| | | | | | |
|---|---|---|---|---|---|
| pseudoscaff_2630 | 27500 | 0.000049942 | 0.029512 | 0.026732042 | 0.026026938 |
| pseudoscaff_2630 | 32500 | 0 | 0.020016 | 0.020801892 | 0.021161884 |
| pseudoscaff_2630 | 42500 | 0.000142464 | 0.024617 | 0.027229144 | 0.027253907 |
| pseudoscaff_2630 | 47500 | 0 | 0.016044 | 0.019049201 | 0.019736086 |
| pseudoscaff_2630 | 57500 | 0 | 0.035319 | 0.029332207 | 0.031203163 |
| pseudoscaff_2630 | 62500 | 0.000827186 | 0.017431 | 0.015185103 | 0.015403641 |
| pseudoscaff_2630 | 67500 | 0.001495284 | 0.029097 | 0.027502016 | 0.033207977 |
| pseudoscaff_2630 | 72500 | 0.000055234 | 0.026262 | 0.028571181 | 0.027564101 |
| pseudoscaff_2630 | 77500 | 0 | 0.01447 | 0.013072271 | 0.013047895 |
| pseudoscaff_2630 | 82500 | 0.000313667 | 0.033514 | 0.034897298 | 0.031824782 |
| pseudoscaff_2630 | 92500 | 0 | 0.036089 | 0.031845951 | 0.033601191 |
| pseudoscaff_363 | 2500 | 0.000368387 | 0.00661 | 0.00817164 | 0.004974607 |
| pseudoscaff_363 | 32500 | 0.000097848 | 0.006965 | 0.008542247 | 0.009239584 |
| pseudoscaff_1490 | 7500 | 0.001497726 | 0.00096 | 0.002260392 | 0.00198824 |
| pseudoscaff_1066 | 12500 | 0.001940181 | 0.002605 | 0.002967942 | 0.002975006 |
| pseudoscaff_1066 | 17500 | 0.001617702 | 0.005602 | 0.003606169 | 0.002954926 |
| pseudoscaff_1066 | 22500 | 0.001775155 | 0.003617 | 0.002733118 | 0.003436717 |
| pseudoscaff_867 | 7500 | 0.001180027 | 0.014848 | 0.007657594 | 0.008696711 |
| pseudoscaff_867 | 17500 | 0.001170153 | 0.00729 | 0.008856295 | 0.010042435 |
| pseudoscaff_867 | 22500 | 0.000750636 | 0.015337 | 0.014337162 | 0.016658419 |
| pseudoscaff_867 | 27500 | 0.000396939 | 0.005646 | 0.006417992 | 0.007699929 |
| pseudoscaff_867 | 32500 | 0.000263158 | 0.017344 | 0.015689631 | 0.016694148 |
| pseudoscaff_867 | 42500 | 0 | 0.006513 | 0.01230858 | 0.012174368 |
| pseudoscaff_867 | 47500 | 0 | 0.013249 | 0.012438594 | 0.013115753 |
| pseudoscaff_867 | 67500 | 0.000677838 | 0.013105 | 0.022453402 | 0.020353728 |
| pseudoscaff_867 | 77500 | 0.000323921 | 0.001692 | 0.015657918 | 0.013185251 |
| pseudoscaff_2122 | 12500 | 0.000226815 | 0.019043 | 0.017458131 | 0.018862605 |
| pseudoscaff_2122 | 17500 | 0.000497879 | 0.019061 | 0.019553474 | 0.017308402 |
| pseudoscaff_2122 | 22500 | 0.000110742 | 0.036226 | 0.022531255 | 0.025177417 |
| pseudoscaff_2122 | 52500 | 0.000832118 | 0.000576 | 0.002978343 | 0.002243094 |
| pseudoscaff_1795 | 72500 | 0.000357604 | 0.003681 | 0.008101199 | 0.00751672 |
| pseudoscaff_1795 | 77500 | 0.000804213 | 0.005101 | 0.01211094 | 0.01250305 |
| pseudoscaff_1795 | 92500 | 0.000393629 | 0.012931 | 0.010515697 | 0.009089651 |
| pseudoscaff_3722 | 2500 | 0.000257309 | 0.003294 | 0.00420487 | 0.004221268 |
| pseudoscaff_3526 | 107500 | 0.001891216 | 0.006084 | 0.007060572 | 0.007382559 |
| pseudoscaff_1092 | 12500 | 0.001285656 | 0.011006 | 0.017691891 | 0.014238197 |
| pseudoscaff_1092 | 17500 | 0 | 0.007536 | 0.023852321 | 0.023400975 |
| pseudoscaff_3744 | 2500 | 0.00089307 | 0.011332 | 0.014363893 | 0.013769625 |
| pseudoscaff_3744 | 17500 | 0.001501168 | 0.008352 | 0.007552346 | 0.00631561 |
| pseudoscaff_3744 | 32500 | 0.000656308 | 0.018401 | 0.014189376 | 0.013146154 |
| pseudoscaff_3744 | 52500 | 0 | 0.010046 | 0.008432832 | 0.007707813 |
| pseudoscaff_3744 | 57500 | 0.000154619 | 0.010565 | 0.013828163 | 0.013188803 |
| pseudoscaff_3744 | 62500 | 0.000084124 | 0.005784 | 0.009066987 | 0.007617309 |
| pseudoscaff_3744 | 67500 | 0.001824921 | 0.015067 | 0.013360904 | 0.012859071 |
| pseudoscaff_1449 | 17500 | 0.001306825 | 0.027427 | 0.027451585 | 0.02546524 |
| pseudoscaff_1449 | 22500 | 0 | 0.03842 | 0.032719424 | 0.036236435 |
| pseudoscaff_1449 | 42500 | 0 | 0.032606 | 0.036613691 | 0.033960437 |
| pseudoscaff_1449 | 52500 | 0.001243885 | 0.017772 | 0.017991792 | 0.020798519 |
| pseudoscaff_1449 | 57500 | 0.000256556 | 0.029205 | 0.027810431 | 0.029304042 |
| pseudoscaff_1449 | 62500 | 0.000348132 | 0.01283 | 0.019258323 | 0.01799414 |

| | | | | | |
|---|---|---|---|---|---|
| pseudoscaff_1449 | 67500 | 0 | 0.018357 | 0.016607545 | 0.016775663 |
| pseudoscaff_1449 | 77500 | 0 | 0.033139 | 0.031862045 | 0.033745688 |
| pseudoscaff_1449 | 87500 | 0 | 0.033682 | 0.031502652 | 0.032689821 |
| pseudoscaff_1449 | 97500 | 0 | 0.020635 | 0.030774981 | 0.028344563 |
| pseudoscaff_373 | 132500 | 0.001883098 | 0.022304 | 0.021376132 | 0.021447342 |
| pseudoscaff_373 | 142500 | 0.0019444 | 0.018408 | 0.014125301 | 0.01533501 |
| pseudoscaff_373 | 147500 | 0.00138121 | 0.002893 | 0.004730912 | 0.004253418 |
| pseudoscaff_3384 | 12500 | 0.000398483 | 0.041094 | 0.033278853 | 0.033777335 |
| pseudoscaff_3384 | 17500 | 0 | 0.03248 | 0.032825213 | 0.034584015 |
| pseudoscaff_3384 | 32500 | 0.000213673 | 0.022931 | 0.031894993 | 0.024033252 |
| pseudoscaff_3384 | 37500 | 0.000816896 | 0.019034 | 0.026351039 | 0.028471508 |
| pseudoscaff_3384 | 42500 | 0.001829181 | 0.026559 | 0.021882772 | 0.023904364 |
| pseudoscaff_3384 | 52500 | 0.00151643 | 0.053939 | 0.039084881 | 0.036000411 |
| pseudoscaff_2688 | 12500 | 0.000622692 | 0.030008 | 0.038633792 | 0.038511607 |
| pseudoscaff_2688 | 27500 | 0.000164806 | 0.03945 | 0.039166295 | 0.03382627 |
| pseudoscaff_2688 | 32500 | 0.000082277 | 0.033278 | 0.037494722 | 0.034288182 |
| pseudoscaff_2688 | 37500 | 0.000847804 | 0.036135 | 0.040777256 | 0.037414116 |
| pseudoscaff_2688 | 42500 | 0.000180938 | 0.027356 | 0.028575275 | 0.028828691 |
| pseudoscaff_2688 | 47500 | 0.000163671 | 0.019249 | 0.020898675 | 0.020726202 |
| pseudoscaff_2688 | 52500 | 0.000064619 | 0.017096 | 0.018687227 | 0.019329618 |
| pseudoscaff_2688 | 57500 | 0 | 0.024827 | 0.027696621 | 0.025768578 |
| pseudoscaff_2688 | 62500 | 0.000241781 | 0.037842 | 0.037200204 | 0.035947586 |
| pseudoscaff_2688 | 72500 | 0 | 0.019476 | 0.017764195 | 0.018301195 |
| pseudoscaff_2688 | 77500 | 0 | 0.00726 | 0.014042497 | 0.014130198 |
| pseudoscaff_2688 | 87500 | 0.000708536 | 0.017775 | 0.018532678 | 0.018841095 |
| pseudoscaff_2688 | 107500 | 0 | 0.000558 | 0.001299755 | 0.001245706 |
| pseudoscaff_2222 | 7500 | 0.001935812 | 0.025239 | 0.027729341 | 0.022436322 |
| pseudoscaff_1863 | 122500 | 0.000653395 | 0.030595 | 0.037396796 | 0.037931515 |
| pseudoscaff_1863 | 137500 | 0 | 0.02575 | 0.024647172 | 0.024672868 |
| pseudoscaff_1863 | 287500 | 0.000221827 | 0.004203 | 0.016596951 | 0.018489791 |
| pseudoscaff_1048 | 7500 | 0.000469408 | 0.014957 | 0.018813278 | 0.023776997 |
| pseudoscaff_1048 | 12500 | 0.000690329 | 0.003194 | 0.015883849 | 0.017083178 |
| pseudoscaff_1048 | 17500 | 0 | 0.009375 | 0.01880455 | 0.019776023 |
| pseudoscaff_1048 | 32500 | 0.001073516 | 0.015212 | 0.014387462 | 0.016385565 |
| pseudoscaff_1048 | 37500 | 0.000120189 | 0.021044 | 0.00936864 | 0.014903918 |
| pseudoscaff_4196 | 7500 | 0.001931356 | 0.003633 | 0.003773823 | 0.004161044 |
| pseudoscaff_3317 | 7500 | 0.000045017 | 0.001394 | 0.018715507 | 0.017183531 |
| pseudoscaff_3317 | 12500 | 0.001260695 | 0.009066 | 0.022756201 | 0.025125523 |
| pseudoscaff_3317 | 17500 | 0.001697563 | 0.025941 | 0.024897243 | 0.025613093 |
| pseudoscaff_1124 | 2500 | 0.000142228 | 0.003241 | 0.004205286 | 0.007693359 |
| pseudoscaff_1124 | 7500 | 0 | 0.01986 | 0.01832081 | 0.017740055 |
| pseudoscaff_1124 | 17500 | 0.000100563 | 0.017223 | 0.018562614 | 0.021595862 |
| pseudoscaff_1124 | 22500 | 0.000143312 | 0.010008 | 0.017171371 | 0.018084598 |
| pseudoscaff_1124 | 32500 | 0 | 0.006146 | 0.002749613 | 0.005126336 |
| pseudoscaff_1124 | 37500 | 0.000132453 | 0.01027 | 0.002091704 | 0.002611033 |
| pseudoscaff_1124 | 42500 | 0 | 0.000273 | 0.002820836 | 0.004480976 |
| pseudoscaff_1124 | 47500 | 0 | 0.005044 | 0.009113984 | 0.010053121 |
| pseudoscaff_1124 | 52500 | 0 | 8.01E-05 | 0.000612727 | 0.00050558 |
| pseudoscaff_2201 | 2500 | 0.001045859 | 0.026786 | 0.031171946 | 0.029974183 |
| pseudoscaff_2201 | 12500 | 0.000771953 | 0.01673 | 0.029739193 | 0.032305346 |

| | | | | | |
|---|---|---|---|---|---|
| pseudoscaff_879 | 187500 | 0.001911872 | 0.004903 | 0.005776602 | 0.005568373 |
| pseudoscaff_879 | 192500 | 0.00061873 | 0.00477 | 0.004543754 | 0.005424088 |
| pseudoscaff_879 | 207500 | 0.001905175 | 0.007475 | 0.008595136 | 0.008401736 |
| pseudoscaff_879 | 217500 | 0.001756461 | 0.010294 | 0.009430037 | 0.009309404 |
| pseudoscaff_879 | 222500 | 0.001720195 | 0.006918 | 0.0090445 | 0.009429032 |
| pseudoscaff_879 | 227500 | 0.000995028 | 0.006218 | 0.007517412 | 0.006875641 |
| pseudoscaff_879 | 232500 | 0.000621884 | 0.013423 | 0.010596363 | 0.009745398 |
| pseudoscaff_879 | 242500 | 0.001103603 | 0.008196 | 0.008091807 | 0.008404354 |
| pseudoscaff_879 | 252500 | 0.000978772 | 0.007004 | 0.007490871 | 0.007525691 |
| pseudoscaff_879 | 262500 | 0 | 0.008378 | 0.011550027 | 0.011472279 |
| pseudoscaff_3715 | 2500 | 0 | 0.018634 | 0.013822635 | 0.013008501 |
| pseudoscaff_3715 | 7500 | 0 | 0.027102 | 0.023751278 | 0.018045087 |
| pseudoscaff_3715 | 17500 | 0 | 0.017323 | 0.010800635 | 0.010187593 |
| pseudoscaff_3715 | 27500 | 0 | 0.019311 | 0.021882251 | 0.021719648 |
| pseudoscaff_2014 | 7500 | 0.001402395 | 0.00263 | 0.003536304 | 0.003605113 |
| pseudoscaff_1189 | 12500 | 0 | 0.012539 | 0.01478568 | 0.015730916 |
| pseudoscaff_424 | 22500 | 0 | 0.010045 | 0.009500053 | 0.012503142 |
| pseudoscaff_424 | 27500 | 0.000177184 | 0.014115 | 0.017554168 | 0.019195454 |
| pseudoscaff_424 | 37500 | 0.001119441 | 0.028944 | 0.026922589 | 0.030100403 |
| pseudoscaff_424 | 42500 | 0.00013996 | 0.025308 | 0.025966943 | 0.029129846 |
| pseudoscaff_424 | 47500 | 0.001175414 | 0.0229 | 0.024208726 | 0.024270928 |
| pseudoscaff_2795 | 7500 | 0 | 0.012593 | 0.008097334 | 0.008025657 |
| pseudoscaff_2795 | 12500 | 0.000216856 | 0.010576 | 0.009726542 | 0.010511709 |
| pseudoscaff_2795 | 27500 | 0 | 0.02557 | 0.007265682 | 0.010044695 |
| pseudoscaff_2795 | 52500 | 0.000837443 | 0.00878 | 0.021682437 | 0.018272197 |
| pseudoscaff_1367 | 2500 | 0 | 0.017176 | 0.016185781 | 0.016657173 |
| pseudoscaff_2455 | 22500 | 0.001385519 | 0 | 0.006334337 | 0.005989897 |
| pseudoscaff_2455 | 27500 | 0.001277952 | 0.008477 | 0.015282707 | 0.018448875 |
| pseudoscaff_2169 | 7500 | 0 | 0.018783 | 0.02247968 | 0.02501733 |
| pseudoscaff_2169 | 12500 | 0.001712359 | 0.024549 | 0.022519291 | 0.024842217 |
| pseudoscaff_2169 | 17500 | 0.001935236 | 0.021468 | 0.025810164 | 0.030542041 |
| pseudoscaff_2169 | 27500 | 0 | 0.022351 | 0.018098434 | 0.016210371 |
| pseudoscaff_2694 | 2500 | 0.001165812 | 0.012588 | 0.013326006 | 0.014446503 |
| pseudoscaff_2694 | 7500 | 0.000080662 | 0.012365 | 0.018525601 | 0.021851093 |
| pseudoscaff_2694 | 12500 | 0.001683186 | 0.028942 | 0.023977803 | 0.024930926 |
| pseudoscaff_2694 | 27500 | 0 | 0.000808 | 0.011237485 | 0.010707316 |
| pseudoscaff_2694 | 37500 | 0.000529815 | 0.012758 | 0.01242759 | 0.013295535 |
| pseudoscaff_2694 | 42500 | 0 | 0.017552 | 0.009455562 | 0.010285009 |
| pseudoscaff_2694 | 57500 | 0.000626693 | 0.013983 | 0.00859807 | 0.009879941 |
| pseudoscaff_2694 | 62500 | 0 | 0.006389 | 0.010919686 | 0.009974649 |
| pseudoscaff_2694 | 77500 | 0.000656749 | 0.006566 | 0.005452079 | 0.006329584 |
| pseudoscaff_2694 | 82500 | 0.000150934 | 0.015936 | 0.007715044 | 0.010161024 |
| pseudoscaff_2694 | 102500 | 0.001187585 | 0.011886 | 0.024449556 | 0.027097547 |
| pseudoscaff_2694 | 107500 | 0.00121884 | 0.013144 | 0.013181372 | 0.01409924 |
| pseudoscaff_2694 | 112500 | 0.001733047 | 0.005991 | 0.019573359 | 0.019881008 |
| pseudoscaff_2193 | 2500 | 0.001374263 | 0.00503 | 0.005024289 | 0.006791069 |
| pseudoscaff_2193 | 7500 | 0.000567983 | 0.009435 | 0.006908986 | 0.008529824 |
| pseudoscaff_2193 | 12500 | 0.001053501 | 0.002063 | 0.0053781 | 0.008730162 |
| pseudoscaff_2193 | 22500 | 0 | 0.000854 | 0.000691717 | 0.00160165 |
| pseudoscaff_2193 | 27500 | 0.000214634 | 0.004971 | 0.001712841 | 0.00196955 |

| | | | | | |
|---|---|---|---|---|---|
| pseudoscaff_2193 | 32500 | 0.00005645 | 0.01081 | 0.001301202 | 0.002595038 |
| pseudoscaff_2193 | 37500 | 0 | 0.008963 | 0.009274934 | 0.00923078 |
| pseudoscaff_2193 | 47500 | 0.000606223 | 0.022814 | 0.014103498 | 0.017492075 |
| pseudoscaff_398 | 32500 | 0.000450912 | 0.039542 | 0.035468318 | 0.038295399 |
| pseudoscaff_703 | 7500 | 0.000171535 | 0.028567 | 0.033691901 | 0.03144176 |
| pseudoscaff_703 | 12500 | 0.000386311 | 0.041039 | 0.050208875 | 0.051618387 |
| pseudoscaff_703 | 37500 | 0 | 0.01965 | 0.020105095 | 0.02097782 |
| pseudoscaff_703 | 47500 | 0.001743932 | 0.031658 | 0.038138315 | 0.039532371 |
| pseudoscaff_2378 | 322500 | 0.000215866 | 0.029689 | 0.021774393 | 0.019968195 |
| pseudoscaff_2378 | 327500 | 0 | 0.022525 | 0.023670292 | 0.024142506 |
| pseudoscaff_2378 | 332500 | 0.000982142 | 0.026833 | 0.029975204 | 0.03149033 |
| pseudoscaff_2253 | 12500 | 0.001783492 | 0.032617 | 0.035601057 | 0.034598999 |
| pseudoscaff_2253 | 17500 | 0 | 0.023331 | 0.023014497 | 0.025172978 |
| pseudoscaff_2253 | 22500 | 0.000167307 | 0.035455 | 0.031432623 | 0.033680754 |
| pseudoscaff_2253 | 32500 | 0 | 0.019496 | 0.01934998 | 0.019047954 |
| pseudoscaff_1493 | 2500 | 0.0003259 | 0.006794 | 0.011044413 | 0.010675218 |
| pseudoscaff_1493 | 7500 | 0.000246786 | 0.0184 | 0.011804032 | 0.012616583 |
| pseudoscaff_1493 | 12500 | 0.001188773 | 0.029063 | 0.015555915 | 0.015517317 |
| pseudoscaff_1493 | 17500 | 0.001158135 | 0.01259 | 0.012207316 | 0.013796368 |
| pseudoscaff_1493 | 22500 | 0.000141435 | 0.013709 | 0.023667859 | 0.02412153 |
| pseudoscaff_1493 | 27500 | 0 | 0.0115 | 0.020300709 | 0.016438594 |
| pseudoscaff_1493 | 32500 | 0.000578358 | 0.009495 | 0.021654196 | 0.019667041 |
| pseudoscaff_1493 | 37500 | 0 | 0.006007 | 0.018438192 | 0.016452092 |
| pseudoscaff_2582 | 7500 | 0.00072617 | 0.008033 | 0.022419558 | 0.020315462 |
| pseudoscaff_3481 | 12500 | 0 | 0.021233 | 0.021167109 | 0.022138275 |
| pseudoscaff_3481 | 17500 | 0 | 0.02779 | 0.026632844 | 0.028507065 |
| pseudoscaff_82 | 22500 | 0 | 0.011064 | 0.011235465 | 0.010863465 |
| pseudoscaff_82 | 32500 | 0 | 0.023144 | 0.022579429 | 0.024585136 |
| pseudoscaff_82 | 37500 | 0.000265045 | 0.011934 | 0.020571259 | 0.020609823 |
| pseudoscaff_82 | 172500 | 0.00135088 | 0.001469 | 0.002492266 | 0.002475094 |
| pseudoscaff_82 | 207500 | 0 | 0.014283 | 0.015883253 | 0.0148678 |
| pseudoscaff_3210 | 2500 | 0.001943347 | 0.019136 | 0.020809748 | 0.021212623 |
| pseudoscaff_42 | 2500 | 0 | 0 | 0.000083423 | 0.000056608 |
| pseudoscaff_119 | 37500 | 0.001131538 | 0.042875 | 0.041640207 | 0.039475728 |
| pseudoscaff_119 | 52500 | 0.00117301 | 0.025318 | 0.012077561 | 0.013994282 |
| pseudoscaff_3827 | 207500 | 0.001355998 | 0.036785 | 0.039082305 | 0.038227988 |
| pseudoscaff_1309 | 157500 | 0.001306475 | 0.00472 | 0.005129911 | 0.005664225 |
| pseudoscaff_3453 | 7500 | 0.000411598 | 0.00098 | 0.007260811 | 0.007385792 |
| pseudoscaff_3972 | 7500 | 0.000992566 | 0.018669 | 0.025110024 | 0.028598155 |
| pseudoscaff_3413 | 27500 | 0 | 0.021196 | 0.02441959 | 0.026215976 |
| pseudoscaff_2724 | 2500 | 0.000170302 | 0.019798 | 0.02104033 | 0.022469821 |
| pseudoscaff_3636 | 2500 | 0.000115431 | 0.000522 | 0.000883324 | 0.001234044 |
| pseudoscaff_2419 | 17500 | 0 | 0.004064 | 0.005085318 | 0.004826297 |
| pseudoscaff_2419 | 27500 | 0.000123111 | 0.002811 | 0.001908832 | 0.002342101 |
| pseudoscaff_2419 | 32500 | 0 | 0.003634 | 0.003707766 | 0.003223178 |
| pseudoscaff_2419 | 42500 | 0.000223134 | 0.014186 | 0.006119964 | 0.007094448 |
| pseudoscaff_1727 | 2500 | 0.001799532 | 0.022873 | 0.030677095 | 0.032444254 |
| pseudoscaff_1727 | 7500 | 0.001107345 | 0.03733 | 0.030156413 | 0.031331417 |
| pseudoscaff_1727 | 57500 | 0.001011787 | 0.006295 | 0.007115124 | 0.007310622 |
| pseudoscaff_1727 | 67500 | 0.001083214 | 0.006243 | 0.008560637 | 0.008987678 |

| | | | | | |
|---|---|---|---|---|---|
| pseudoscaff_1727 | 72500 | 0.00193312 | 0.008673 | 0.007542363 | 0.00817026 |
| pseudoscaff_1727 | 77500 | 0.001789019 | 0.007914 | 0.007893219 | 0.007810436 |
| pseudoscaff_1727 | 82500 | 0.001033089 | 0.007935 | 0.007426909 | 0.008056289 |
| pseudoscaff_1727 | 92500 | 0.000844315 | 0.015739 | 0.01147848 | 0.013503017 |
| pseudoscaff_1727 | 97500 | 0.000297121 | 0.007388 | 0.006046318 | 0.006062079 |
| pseudoscaff_1727 | 102500 | 0.001978266 | 0.005579 | 0.009701945 | 0.009081759 |
| pseudoscaff_1727 | 107500 | 0.00030215 | 0.003663 | 0.012315993 | 0.011672027 |
| pseudoscaff_1723 | 42500 | 0.001111886 | 0.007914 | 0.005848508 | 0.005226062 |
| pseudoscaff_1119 | 12500 | 0.00165122 | 0.007688 | 0.006126123 | 0.006538465 |
| pseudoscaff_1888 | 27500 | 0.000341692 | 0.036096 | 0.036317161 | 0.035086412 |
| pseudoscaff_1888 | 32500 | 0 | 0.02389 | 0.026140758 | 0.026366062 |
| pseudoscaff_1888 | 37500 | 0 | 0.028466 | 0.027706271 | 0.027262877 |
| pseudoscaff_1888 | 42500 | 0.001165068 | 0.036019 | 0.040728031 | 0.038753812 |
| pseudoscaff_1888 | 52500 | 0 | 0.026999 | 0.036625884 | 0.035290002 |
| pseudoscaff_1888 | 57500 | 0.000138266 | 0.022129 | 0.027764876 | 0.025319251 |
| pseudoscaff_1888 | 62500 | 0 | 0.021791 | 0.026639833 | 0.029839969 |
| pseudoscaff_1888 | 67500 | 0.0016031 | 0.015527 | 0.023814201 | 0.02427468 |
| pseudoscaff_3624 | 2500 | 0.000390931 | 0.023641 | 0.026649465 | 0.028199066 |
| pseudoscaff_460 | 22500 | 0.000580452 | 0.004858 | 0.004162494 | 0.004628272 |
| pseudoscaff_460 | 47500 | 0.001809041 | 0.005478 | 0.010212466 | 0.00710789 |
| pseudoscaff_460 | 57500 | 0.00195782 | 0.007093 | 0.006608904 | 0.006034535 |
| pseudoscaff_3936 | 2500 | 0 | 0.003758 | 0.005837313 | 0.006310957 |
| pseudoscaff_3936 | 7500 | 0.00074333 | 0.00342 | 0.010698135 | 0.010686116 |
| pseudoscaff_3078 | 12500 | 0.000178815 | 0.012838 | 0.011725631 | 0.012487342 |
| pseudoscaff_276 | 52500 | 0.001095792 | 0.012472 | 0.011209548 | 0.01022456 |
| pseudoscaff_276 | 67500 | 0.001833812 | 0.022757 | 0.020335436 | 0.021034928 |
| pseudoscaff_276 | 117500 | 0.00030563 | 0.028428 | 0.027115025 | 0.028135721 |
| pseudoscaff_276 | 167500 | 0.001594287 | 0.025287 | 0.024875241 | 0.026696279 |
| pseudoscaff_276 | 232500 | 0.000157677 | 0.000165 | 0.003235506 | 0.0034142 |
| pseudoscaff_276 | 457500 | 0.000616588 | 0.037815 | 0.037222534 | 0.038130467 |
| pseudoscaff_276 | 622500 | 0.000282374 | 0.027434 | 0.026846394 | 0.026888718 |
| pseudoscaff_2788 | 2500 | 0 | 0.004688 | 0.017855791 | 0.014064608 |
| pseudoscaff_2788 | 7500 | 0 | 0.006638 | 0.020882216 | 0.018531834 |
| pseudoscaff_2788 | 22500 | 0.001991006 | 0.006678 | 0.006880634 | 0.007012876 |
| pseudoscaff_2764 | 7500 | 0 | 0.004964 | 0.006442571 | 0.005883274 |
| pseudoscaff_2764 | 12500 | 0.000075596 | 0.007379 | 0.007580685 | 0.007916164 |
| pseudoscaff_2764 | 17500 | 0 | 0.009876 | 0.010224233 | 0.010348696 |
| pseudoscaff_2764 | 27500 | 0.000308099 | 0.019025 | 0.024681684 | 0.027983459 |
| pseudoscaff_2764 | 42500 | 0 | 0.034081 | 0.03725518 | 0.033617406 |
| pseudoscaff_2764 | 52500 | 0.00145969 | 0.034141 | 0.033750493 | 0.035267818 |
| pseudoscaff_2764 | 67500 | 0.000841272 | 0.018332 | 0.024525783 | 0.024911328 |
| pseudoscaff_2764 | 72500 | 0.001002733 | 0.024786 | 0.025750821 | 0.02695612 |
| pseudoscaff_2381 | 12500 | 0.000851636 | 0.017392 | 0.020738086 | 0.019718955 |
| pseudoscaff_2381 | 22500 | 0.00187259 | 0.022279 | 0.018314415 | 0.019215925 |
| pseudoscaff_2381 | 37500 | 0.001654944 | 0.027908 | 0.030286087 | 0.032006885 |
| pseudoscaff_2381 | 47500 | 0.000678558 | 0.023947 | 0.027616472 | 0.027372638 |
| pseudoscaff_2381 | 52500 | 0.001382696 | 0.02842 | 0.029304084 | 0.028742647 |
| pseudoscaff_2381 | 67500 | 0.000413937 | 0.037905 | 0.042143201 | 0.04365089 |
| pseudoscaff_2965 | 7500 | 0.000126584 | 0.0375 | 0.042392362 | 0.043480212 |
| pseudoscaff_656 | 7500 | 0.000147485 | 0.036544 | 0.042614461 | 0.038923221 |

| | | | | | |
|---|---|---|---|---|---|
| pseudoscaff_656 | 12500 | 0.000317253 | 0.033638 | 0.032067721 | 0.032956399 |
| pseudoscaff_656 | 17500 | 0.000126625 | 0.021701 | 0.027449683 | 0.025376951 |
| pseudoscaff_656 | 22500 | 0.001034756 | 0.022078 | 0.018409 | 0.018418265 |
| pseudoscaff_656 | 27500 | 0 | 0.007713 | 0.014903426 | 0.012974134 |
| pseudoscaff_656 | 62500 | 0.000365817 | 0.010052 | 0.016848482 | 0.015879285 |
| pseudoscaff_656 | 67500 | 0 | 0.004867 | 0.01170666 | 0.0105871 |
| pseudoscaff_656 | 72500 | 0 | 0.003503 | 0.014653572 | 0.014201555 |
| pseudoscaff_656 | 77500 | 0.000337702 | 0.010292 | 0.012283357 | 0.011459132 |
| pseudoscaff_656 | 82500 | 0 | 0.018153 | 0.022726069 | 0.024700661 |
| pseudoscaff_656 | 87500 | 0 | 0.012451 | 0.014235971 | 0.012347394 |
| pseudoscaff_656 | 92500 | 0 | 0.006114 | 0.015680687 | 0.016499324 |
| pseudoscaff_656 | 107500 | 0.000644346 | 0.016676 | 0.013892802 | 0.012164661 |
| pseudoscaff_656 | 112500 | 0 | 0.013166 | 0.004726126 | 0.006895814 |
| pseudoscaff_656 | 117500 | 0.00009179 | 0.002851 | 0.014988523 | 0.014286153 |
| pseudoscaff_656 | 122500 | 0 | 0.010244 | 0.009368985 | 0.009648699 |
| pseudoscaff_656 | 127500 | 0.000323471 | 0.008456 | 0.0059822 | 0.006334978 |
| pseudoscaff_656 | 132500 | 0 | 0.023783 | 0.013753419 | 0.01344721 |
| pseudoscaff_656 | 142500 | 0.001313363 | 0.033174 | 0.025581234 | 0.024717943 |
| pseudoscaff_656 | 152500 | 0.00010339 | 0.024773 | 0.027375226 | 0.028134541 |
| pseudoscaff_656 | 157500 | 0.000061854 | 0.02637 | 0.027545556 | 0.028379602 |
| pseudoscaff_656 | 162500 | 0.000050469 | 0.02643 | 0.036001267 | 0.03560838 |
| pseudoscaff_656 | 167500 | 0.000616794 | 0.030111 | 0.036828397 | 0.032581011 |
| pseudoscaff_656 | 172500 | 0.000101386 | 0.005842 | 0.020844594 | 0.018701323 |
| pseudoscaff_656 | 177500 | 0.000288991 | 0.010526 | 0.008596046 | 0.008649627 |
| pseudoscaff_656 | 182500 | 0.000020564 | 0.024133 | 0.015220711 | 0.018775529 |
| pseudoscaff_656 | 187500 | 0.00177181 | 0.040942 | 0.037555289 | 0.037777042 |
| pseudoscaff_656 | 192500 | 0.000355926 | 0.03158 | 0.03380227 | 0.034779835 |
| pseudoscaff_3380 | 12500 | 0.00070162 | 0.003062 | 0.010579149 | 0.009328238 |
| pseudoscaff_3380 | 32500 | 0.000604992 | 0.007444 | 0.017637082 | 0.01596617 |
| pseudoscaff_3380 | 42500 | 0.001285051 | 0.00936 | 0.009830841 | 0.010123726 |
| pseudoscaff_2994 | 232500 | 0.001960885 | 0.004689 | 0.00626138 | 0.005572439 |
| pseudoscaff_838 | 442500 | 0.00111034 | 0.002674 | 0.003334495 | 0.003566166 |
| pseudoscaff_838 | 662500 | 0.000624774 | 0.008837 | 0.011291674 | 0.011510349 |
| pseudoscaff_838 | 667500 | 0.001151149 | 0.00542 | 0.005708586 | 0.005327737 |
| pseudoscaff_2446 | 52500 | 0.001956587 | 0.01579 | 0.019131978 | 0.019832362 |
| pseudoscaff_2446 | 207500 | 0.000674863 | 0.016719 | 0.023002561 | 0.022851807 |
| pseudoscaff_402 | 17500 | 0.000511039 | 0.031115 | 0.041599279 | 0.043037062 |
| pseudoscaff_852 | 7500 | 0 | 0.026873 | 0.031462051 | 0.032554606 |
| pseudoscaff_852 | 12500 | 0.000937018 | 0.012565 | 0.014381254 | 0.01515245 |
| pseudoscaff_852 | 17500 | 0 | 0.005118 | 0.007392235 | 0.005724389 |
| pseudoscaff_852 | 27500 | 0 | 0.001423 | 0.009484169 | 0.01255751 |
| pseudoscaff_3994 | 7500 | 0.000072193 | 0.00953 | 0.007820456 | 0.009059799 |
| pseudoscaff_217 | 47500 | 0.001578155 | 0.033673 | 0.047002096 | 0.0458587 |
| pseudoscaff_217 | 62500 | 0.000835121 | 0.04217 | 0.047116018 | 0.043054529 |
| pseudoscaff_217 | 67500 | 0.000161928 | 0.023082 | 0.02784009 | 0.027413976 |
| pseudoscaff_217 | 72500 | 0 | 0.032446 | 0.033970338 | 0.032103495 |
| pseudoscaff_217 | 87500 | 0.000350175 | 0.032725 | 0.039232109 | 0.036863168 |
| pseudoscaff_3468 | 57500 | 0.001548811 | 0.029605 | 0.032163154 | 0.031522806 |
| pseudoscaff_3396 | 17500 | 0.001657304 | 0.002471 | 0.006093059 | 0.006174315 |
| pseudoscaff_824 | 2500 | 0.000463412 | 0.004026 | 0.003329079 | 0.003090948 |

| | | | | | |
|---|---|---|---|---|---|
| pseudoscaff_824 | 7500 | 0.000581897 | 0.004428 | 0.003223308 | 0.0031385 |
| pseudoscaff_4055 | 32500 | 0.001134224 | 0.023835 | 0.02658504 | 0.03042447 |
| pseudoscaff_4055 | 37500 | 0.001158096 | 0.025225 | 0.029289401 | 0.030250465 |
| pseudoscaff_4055 | 112500 | 0.001038322 | 0.027326 | 0.03180718 | 0.032836404 |
| pseudoscaff_4055 | 167500 | 0.001088651 | 0.031565 | 0.026958585 | 0.033341104 |
| pseudoscaff_4055 | 192500 | 0.000281386 | 0.032334 | 0.038032856 | 0.038462632 |
| pseudoscaff_4055 | 207500 | 0.000561357 | 0.041801 | 0.034525322 | 0.034960354 |
| pseudoscaff_4055 | 257500 | 0.001242612 | 0.024647 | 0.023421696 | 0.025017425 |
| pseudoscaff_3387 | 7500 | 0.001308354 | 0.032952 | 0.03021794 | 0.030276877 |
| pseudoscaff_3387 | 12500 | 0.001258612 | 0.014117 | 0.027795508 | 0.029542247 |
| pseudoscaff_3387 | 52500 | 0.000218941 | 0.021642 | 0.026899654 | 0.030087127 |
| pseudoscaff_3387 | 297500 | 0.000713201 | 0.023127 | 0.031417147 | 0.03259034 |
| pseudoscaff_3140 | 2500 | 0 | 0.000239 | 0.005370086 | 0.007734684 |
| pseudoscaff_3140 | 7500 | 0.001174334 | 0.006419 | 0.01178919 | 0.009079729 |
| pseudoscaff_3140 | 12500 | 0.001565234 | 0.004327 | 0.009610391 | 0.009994581 |
| pseudoscaff_3140 | 22500 | 0.000589486 | 0.001416 | 0.012460424 | 0.018729345 |
| pseudoscaff_3140 | 27500 | 0.000918068 | 0.000811 | 0.008043916 | 0.007996074 |
| pseudoscaff_3140 | 32500 | 0.000228603 | 0.001859 | 0.002599776 | 0.002546123 |
| pseudoscaff_2800 | 302500 | 0.000324139 | 0.007433 | 0.009607399 | 0.008856114 |
| pseudoscaff_281 | 32500 | 0.000460308 | 0.046631 | 0.039055965 | 0.039980142 |
| pseudoscaff_281 | 47500 | 0.000283736 | 0.037453 | 0.033070114 | 0.033949026 |
| pseudoscaff_2870 | 102500 | 0.00171375 | 0.005738 | 0.010007066 | 0.012852899 |
| pseudoscaff_543 | 17500 | 0.000076807 | 0.006175 | 0.005734976 | 0.00590334 |
| pseudoscaff_543 | 47500 | 0.000910874 | 0.018859 | 0.021648056 | 0.020946219 |
| pseudoscaff_543 | 52500 | 0.000072922 | 0.010339 | 0.014347814 | 0.014405901 |
| pseudoscaff_543 | 57500 | 0.00150008 | 0.017537 | 0.02008746 | 0.019607651 |
| pseudoscaff_543 | 62500 | 0.000059505 | 0.013178 | 0.01682771 | 0.017453404 |
| pseudoscaff_543 | 72500 | 0.001672952 | 0.010208 | 0.009437679 | 0.010322559 |
| pseudoscaff_543 | 77500 | 0.000427292 | 0.018894 | 0.017248484 | 0.01673447 |
| pseudoscaff_543 | 82500 | 0.000163081 | 0.012014 | 0.014210856 | 0.014117809 |
| pseudoscaff_543 | 87500 | 0.000566807 | 0.015404 | 0.013942963 | 0.014401925 |
| pseudoscaff_543 | 97500 | 0.000209154 | 0.025469 | 0.028811786 | 0.031579658 |
| pseudoscaff_543 | 107500 | 0.001015247 | 0.018013 | 0.018569182 | 0.017610205 |
| pseudoscaff_543 | 112500 | 0 | 0.031105 | 0.031246819 | 0.0318293 |
| pseudoscaff_543 | 122500 | 0.000814919 | 0.022884 | 0.024956888 | 0.025266497 |
| pseudoscaff_543 | 177500 | 0.001378199 | 0.026082 | 0.033455047 | 0.032843436 |
| pseudoscaff_2433 | 2500 | 0.000188763 | 0.016533 | 0.00900284 | 0.008578373 |
| pseudoscaff_2932 | 12500 | 0.001481304 | 0.004922 | 0.007922945 | 0.008824213 |
| pseudoscaff_256 | 517500 | 0.001537633 | 0.014418 | 0.013636459 | 0.013828344 |
| pseudoscaff_256 | 527500 | 0.000575362 | 0.012401 | 0.010583613 | 0.00843493 |
| pseudoscaff_256 | 532500 | 0.001128542 | 0.007543 | 0.007790611 | 0.008472388 |
| pseudoscaff_3014 | 2500 | 0.000110344 | 0.008252 | 0.010515287 | 0.010123672 |
| pseudoscaff_3014 | 7500 | 0.001253267 | 0.020931 | 0.030031864 | 0.029307171 |
| pseudoscaff_3143 | 7500 | 0 | 0 | 0.005553811 | 0.004323271 |
| pseudoscaff_3143 | 12500 | 0.001349084 | 0.005935 | 0.010394393 | 0.010502261 |
| pseudoscaff_3143 | 17500 | 0.000609102 | 0 | 0.00536682 | 0.003857426 |
| pseudoscaff_3143 | 22500 | 0.000610729 | 0.000549 | 0.00568381 | 0.00525945 |
| pseudoscaff_3524 | 212500 | 0.000269251 | 0.004493 | 0.021156041 | 0.014554004 |
| pseudoscaff_3295 | 2500 | 0.001826312 | 0.014361 | 0.013495691 | 0.01360532 |
| pseudoscaff_3271 | 2500 | 0 | 0.020846 | 0.024386379 | 0.026087069 |

| | | | | | |
|---|---|---|---|---|---|
| pseudoscaff_3271 | 7500 | 0.000147668 | 0.017773 | 0.026352113 | 0.026352822 |
| pseudoscaff_3271 | 12500 | 0.000255941 | 0.01672 | 0.012679238 | 0.0148573 |
| pseudoscaff_412 | 2500 | 0 | 0.001632 | 0.002232112 | 0.002062969 |
| pseudoscaff_125 | 2500 | 0.001877984 | 0.024728 | 0.033275563 | 0.035304665 |
| pseudoscaff_125 | 7500 | 0.000618696 | 0.01648 | 0.017928799 | 0.017712184 |
| pseudoscaff_125 | 12500 | 0 | 0.023338 | 0.018689628 | 0.017766643 |
| pseudoscaff_125 | 17500 | 0 | 0.012997 | 0.017906675 | 0.018946363 |
| pseudoscaff_125 | 22500 | 0.001321084 | 0.015274 | 0.016438408 | 0.015825979 |
| pseudoscaff_125 | 27500 | 0 | 0.003748 | 0.006216131 | 0.006453025 |
| pseudoscaff_125 | 32500 | 0.001840455 | 0.017132 | 0.022272743 | 0.022122647 |
| pseudoscaff_125 | 37500 | 0.00009851 | 0.018203 | 0.016599437 | 0.01798823 |
| pseudoscaff_125 | 42500 | 0.000304334 | 0.012183 | 0.011627739 | 0.011713304 |
| pseudoscaff_125 | 47500 | 0.000200332 | 0.008961 | 0.013887784 | 0.014061392 |
| pseudoscaff_125 | 57500 | 0.000337058 | 0.028123 | 0.021110417 | 0.019532769 |
| pseudoscaff_125 | 67500 | 0.000555232 | 0.022572 | 0.033920501 | 0.030897492 |
| pseudoscaff_125 | 77500 | 0.0000938 | 0.017683 | 0.018254949 | 0.020715871 |
| pseudoscaff_125 | 82500 | 0.001072271 | 0.014531 | 0.018847446 | 0.022133782 |
| pseudoscaff_125 | 87500 | 0.000183944 | 0.008654 | 0.010940203 | 0.009608334 |
| pseudoscaff_125 | 92500 | 0.00015151 | 0.011432 | 0.015153699 | 0.014967197 |
| pseudoscaff_125 | 97500 | 0 | 0.008294 | 0.016515138 | 0.018606693 |
| pseudoscaff_125 | 102500 | 0 | 0.001835 | 0.003254475 | 0.00268804 |
| pseudoscaff_125 | 107500 | 0.001080576 | 0.029916 | 0.033744422 | 0.035340248 |
| pseudoscaff_125 | 112500 | 0.000833247 | 0.02084 | 0.028321378 | 0.026486106 |
| pseudoscaff_3530 | 17500 | 0.00150575 | 0.005793 | 0.004837027 | 0.005889807 |
| pseudoscaff_3664 | 17500 | 0.001149193 | 0.0156 | 0.010738771 | 0.012708206 |
| pseudoscaff_3664 | 32500 | 0.000706143 | 0.004911 | 0.007752121 | 0.010016892 |
| pseudoscaff_3664 | 47500 | 0 | 0.012913 | 0.003227667 | 0.00652266 |
| pseudoscaff_3664 | 52500 | 0 | 0.007872 | 0.008103912 | 0.007980423 |
| pseudoscaff_1797 | 2500 | 0 | 0.001142 | 0.011932653 | 0.009504728 |
| pseudoscaff_1797 | 7500 | 0 | 0.004345 | 0.006513377 | 0.006298407 |
| pseudoscaff_1797 | 17500 | 0.000319585 | 0.001401 | 0.005041073 | 0.005178306 |
| pseudoscaff_1797 | 22500 | 0.000864588 | 0.005704 | 0.008427232 | 0.008619694 |
| pseudoscaff_1797 | 27500 | 0.000636002 | 0.004154 | 0.005314428 | 0.004005249 |
| pseudoscaff_1797 | 32500 | 0 | 0.004425 | 0.003938356 | 0.00452489 |
| pseudoscaff_1797 | 37500 | 0.000422083 | 0.010958 | 0.008820385 | 0.008590467 |
| pseudoscaff_1797 | 42500 | 0 | 0.006424 | 0.006169276 | 0.005859945 |
| pseudoscaff_1797 | 47500 | 0.000766141 | 0.005994 | 0.01050413 | 0.009845993 |
| pseudoscaff_1797 | 62500 | 0.000182101 | 0.004258 | 0.011815569 | 0.010325046 |
| pseudoscaff_1797 | 72500 | 0.001318443 | 0.019496 | 0.012262875 | 0.012318177 |
| pseudoscaff_1797 | 77500 | 0 | 0.014555 | 0.017176409 | 0.019680325 |
| pseudoscaff_1797 | 82500 | 0.001442228 | 0.010912 | 0.011459711 | 0.010854101 |
| pseudoscaff_1797 | 87500 | 0.000203455 | 0.007323 | 0.013190748 | 0.008725954 |
| pseudoscaff_1797 | 102500 | 0.001250254 | 0.011532 | 0.016600685 | 0.017249201 |
| pseudoscaff_1797 | 107500 | 0 | 0.012287 | 0.01337167 | 0.01129055 |
| pseudoscaff_3152 | 77500 | 0.001874623 | 0.000891 | 0.001986675 | 0.001922138 |
| pseudoscaff_957 | 7500 | 0.000477259 | 0.017376 | 0.013142658 | 0.01359902 |
| pseudoscaff_957 | 62500 | 0 | 0.021194 | 0.036276292 | 0.035986835 |
| pseudoscaff_1648 | 7500 | 0.000128563 | 0.018697 | 0.001354204 | 0.00159992 |
| pseudoscaff_1648 | 12500 | 0.00027445 | 0.001565 | 0.001388996 | 0.001420112 |
| pseudoscaff_1648 | 17500 | 0 | 0.000941 | 0.00357206 | 0.002986319 |

| | | | | | |
|---|---|---|---|---|---|
| pseudoscaff_1648 | 22500 | 0 | 0.001914 | 0.009945917 | 0.008070665 |
| pseudoscaff_1648 | 27500 | 0.001276715 | 0.012662 | 0.008459674 | 0.009105559 |
| pseudoscaff_1648 | 42500 | 0 | 0.016881 | 0.012898449 | 0.015868435 |
| pseudoscaff_1648 | 47500 | 0 | 0.003311 | 0.001609954 | 0.00216007 |
| pseudoscaff_1648 | 52500 | 0 | 0.006726 | 0.002375877 | 0.002492698 |
| pseudoscaff_1648 | 57500 | 0 | 0.002891 | 0.001690669 | 0.001530144 |
| pseudoscaff_1648 | 67500 | 0.000138925 | 0.001814 | 0.023557194 | 0.016468943 |
| pseudoscaff_1648 | 92500 | 0.000282014 | 0.02408 | 0.024669692 | 0.020954828 |
| pseudoscaff_1648 | 97500 | 0.000486359 | 0.03314 | 0.015888127 | 0.015636083 |
| pseudoscaff_1648 | 117500 | 0.00124649 | 0.015571 | 0.005356627 | 0.006795342 |
| pseudoscaff_1648 | 127500 | 0.000210163 | 0.008277 | 0.004552111 | 0.006166927 |
| pseudoscaff_1648 | 132500 | 0.00145016 | 0.015153 | 0.007308824 | 0.007711848 |
| pseudoscaff_1648 | 137500 | 0.000145748 | 0.012294 | 0.005324344 | 0.008129698 |
| pseudoscaff_1648 | 142500 | 0.000129717 | 0.012673 | 0.007093153 | 0.0061566 |
| pseudoscaff_1648 | 162500 | 0.001445538 | 0.014819 | 0.003941327 | 0.008453299 |
| pseudoscaff_1648 | 167500 | 0 | 0.015707 | 0.004334389 | 0.00555817 |
| pseudoscaff_1648 | 177500 | 0.000428032 | 0.024376 | 0.00932869 | 0.014199333 |
| pseudoscaff_1648 | 182500 | 0.000108557 | 0.025599 | 0.007533165 | 0.010704802 |
| pseudoscaff_1648 | 187500 | 0.00076656 | 0.008346 | 0.006385836 | 0.006997915 |
| pseudoscaff_1648 | 197500 | 0.00037874 | 0.018512 | 0.00284023 | 0.00281856 |
| pseudoscaff_1648 | 202500 | 0.000493727 | 0.010211 | 0.00255865 | 0.003097389 |
| pseudoscaff_1648 | 222500 | 0 | 0.012169 | 0.003878293 | 0.003226127 |
| pseudoscaff_2835 | 2500 | 0.001522776 | 0.008005 | 0.005168964 | 0.006540273 |
| pseudoscaff_2835 | 7500 | 0.00072841 | 0.018003 | 0.016286693 | 0.016379527 |
| pseudoscaff_2835 | 12500 | 0 | 0.024007 | 0.011518265 | 0.012987318 |
| pseudoscaff_2835 | 17500 | 0.001873996 | 0.000496 | 0.000644774 | 0.000950202 |
| pseudoscaff_2835 | 27500 | 0.000519733 | 0 | 0.000557504 | 0.000561344 |
| pseudoscaff_2835 | 32500 | 0 | 0 | 0.000096684 | 0.000100528 |
| pseudoscaff_2195 | 2500 | 0.001151318 | 0.028165 | 0.031784355 | 0.034958918 |
| pseudoscaff_2195 | 12500 | 0 | 0.022357 | 0.024466036 | 0.023661295 |
| pseudoscaff_4058 | 2500 | 0.001733795 | 0.002083 | 0.010976602 | 0.009615822 |
| pseudoscaff_748 | 7500 | 0.000124894 | 0.020873 | 0.026705577 | 0.027963304 |
| pseudoscaff_748 | 32500 | 0.000330838 | 0.023195 | 0.033755049 | 0.033545653 |
| pseudoscaff_748 | 52500 | 0.001797861 | 0.012441 | 0.024234401 | 0.022519764 |
| pseudoscaff_3256 | 37500 | 0 | 0.010349 | 0.020915618 | 0.020660943 |
| pseudoscaff_3256 | 72500 | 0.000707327 | 0.01494 | 0.020780383 | 0.021570159 |
| pseudoscaff_3256 | 77500 | 0 | 0.017092 | 0.019305243 | 0.021090206 |
| pseudoscaff_3256 | 157500 | 0 | 0.02031 | 0.034502016 | 0.036887831 |
| pseudoscaff_2602 | 2500 | 0.000311563 | 0.006526 | 0.002112845 | 0.005838013 |
| pseudoscaff_2602 | 7500 | 0.001012705 | 0.005743 | 0.001519054 | 0.003234555 |
| pseudoscaff_3437 | 7500 | 0.001070383 | 0.035303 | 0.033643242 | 0.035974999 |
| pseudoscaff_3417 | 7500 | 0.001498556 | 0.008704 | 0.016904894 | 0.013435487 |
| pseudoscaff_1952 | 2500 | 0.000288669 | 0.002896 | 0.003468856 | 0.00327248 |
| pseudoscaff_1952 | 7500 | 0.000354109 | 0.007943 | 0.006442459 | 0.005875967 |
| pseudoscaff_1952 | 12500 | 0 | 0.002289 | 0.0061449 | 0.006228009 |
| pseudoscaff_1952 | 22500 | 0.001561505 | 0.008868 | 0.008070141 | 0.0075458 |
| pseudoscaff_2712 | 17500 | 0 | 0.008076 | 0.010348766 | 0.012157099 |
| pseudoscaff_2712 | 22500 | 0 | 0.006064 | 0.008083183 | 0.009638064 |
| pseudoscaff_2712 | 27500 | 0 | 0.002947 | 0.009420146 | 0.01151997 |
| pseudoscaff_2712 | 32500 | 0 | 0.003813 | 0.009525879 | 0.00826142 |

| | | | | | |
|---|---|---|---|---|---|
| pseudoscaff_2712 | 47500 | 0 | 0.00689 | 0.009458122 | 0.009252867 |
| pseudoscaff_2712 | 92500 | 0 | 0.016046 | 0.013643707 | 0.01386855 |
| pseudoscaff_2712 | 112500 | 0 | 0.001063 | 0.002113706 | 0.002082051 |
| pseudoscaff_3700 | 2500 | 0.000612278 | 0.010482 | 0.009881525 | 0.008636291 |
| pseudoscaff_3700 | 7500 | 0.000371738 | 0.019711 | 0.011717666 | 0.010267246 |
| pseudoscaff_3700 | 12500 | 0 | 0.004893 | 0.006419137 | 0.007302485 |
| pseudoscaff_3700 | 17500 | 0.000443934 | 0.002059 | 0.00637694 | 0.006655596 |
| pseudoscaff_3700 | 22500 | 0.000088045 | 0.005191 | 0.003617394 | 0.003283323 |
| pseudoscaff_3700 | 32500 | 0 | 0.021426 | 0.00570994 | 0.006404237 |
| pseudoscaff_3700 | 42500 | 0.000313392 | 0.009249 | 0.01244196 | 0.0124329 |
| pseudoscaff_3700 | 47500 | 0 | 0.010402 | 0.010995263 | 0.010512122 |
| pseudoscaff_3700 | 52500 | 0 | 0.005938 | 0.00914741 | 0.009212513 |
| pseudoscaff_3700 | 67500 | 0.000281521 | 0.006645 | 0.016216889 | 0.018329049 |
| pseudoscaff_3700 | 72500 | 0.000476833 | 0.017034 | 0.015563948 | 0.014820126 |
| pseudoscaff_3700 | 82500 | 0.00025766 | 0.013397 | 0.005939772 | 0.00843031 |
| pseudoscaff_3700 | 87500 | 0.000592017 | 0.013561 | 0.016279452 | 0.015860211 |
| pseudoscaff_3700 | 92500 | 0.000108212 | 0.003004 | 0.017161701 | 0.016297606 |
| pseudoscaff_3700 | 97500 | 0 | 0.006897 | 0.007128111 | 0.006733742 |
| pseudoscaff_3700 | 102500 | 0.00074346 | 0.00593 | 0.009933769 | 0.009550462 |
| pseudoscaff_3700 | 107500 | 0.000136218 | 0.012549 | 0.013178402 | 0.014022135 |
| pseudoscaff_3700 | 112500 | 0 | 0.006023 | 0.01153605 | 0.011987822 |
| pseudoscaff_3700 | 117500 | 0 | 0.00304 | 0.013718158 | 0.012137103 |
| pseudoscaff_3700 | 127500 | 0.000380805 | 0.020186 | 0.016157673 | 0.016401285 |
| pseudoscaff_3700 | 132500 | 0.000317584 | 0.010264 | 0.029764217 | 0.02691035 |
| pseudoscaff_3700 | 137500 | 0.000424895 | 0.021442 | 0.020652412 | 0.020430231 |
| pseudoscaff_3700 | 142500 | 0.000474302 | 0.015446 | 0.018082026 | 0.017515388 |
| pseudoscaff_3700 | 147500 | 0.000339911 | 0.000993 | 0.001996365 | 0.002057819 |
| pseudoscaff_3700 | 152500 | 0 | 0.005805 | 0.017169398 | 0.014523307 |
| pseudoscaff_3700 | 157500 | 0 | 0.006703 | 0.029120401 | 0.027293597 |
| pseudoscaff_3700 | 162500 | 0.001039319 | 0.001865 | 0.00347324 | 0.00406736 |
| pseudoscaff_3700 | 167500 | 0.00082402 | 0.006838 | 0.005787522 | 0.007736865 |
| pseudoscaff_833 | 112500 | 0.00138888 | 0.001179 | 0.002393914 | 0.002504843 |
| pseudoscaff_2057 | 7500 | 0 | 0.036555 | 0.043310232 | 0.044067612 |
| pseudoscaff_2057 | 12500 | 0.000186929 | 0.036776 | 0.03529612 | 0.035193534 |
| pseudoscaff_2057 | 17500 | 0.001015334 | 0.033535 | 0.033275267 | 0.031231595 |
| pseudoscaff_619 | 12500 | 0.001887319 | 0.04119 | 0.041478572 | 0.041890579 |
| pseudoscaff_3028 | 37500 | 0.000945932 | 0.01501 | 0.022755252 | 0.021043953 |
| pseudoscaff_1259 | 17500 | 0.000485725 | 0.000345 | 0.005260267 | 0.005000319 |
| pseudoscaff_1522 | 2500 | 0.000105672 | 0.013387 | 0.011986928 | 0.01289508 |
| pseudoscaff_1522 | 7500 | 0.000316838 | 0.021661 | 0.026769372 | 0.027432888 |
| pseudoscaff_1522 | 22500 | 0.001359576 | 0.017798 | 0.032471634 | 0.032376025 |
| pseudoscaff_1522 | 27500 | 0.001546174 | 0.032364 | 0.029107555 | 0.028970162 |
| pseudoscaff_1522 | 37500 | 0.000684872 | 0.033439 | 0.038609151 | 0.039873783 |
| pseudoscaff_1522 | 57500 | 0 | 0.036083 | 0.033142656 | 0.033664975 |
| pseudoscaff_1522 | 67500 | 0.000265292 | 0.022128 | 0.021793127 | 0.0228203 |
| pseudoscaff_1522 | 77500 | 0.000123151 | 0.021133 | 0.024191158 | 0.025381947 |
| pseudoscaff_1522 | 82500 | 0.001794594 | 0.026824 | 0.029858077 | 0.030819758 |
| pseudoscaff_850 | 7500 | 0.000151982 | 0.023327 | 0.026932627 | 0.026011706 |
| pseudoscaff_850 | 12500 | 0.001844261 | 0.026355 | 0.023450832 | 0.022868137 |
| pseudoscaff_850 | 22500 | 0.001907189 | 0.029979 | 0.028653032 | 0.030935089 |

| | | | | | |
|---|---|---|---|---|---|
| pseudoscaff_850 | 27500 | 0 | 0.028798 | 0.0361309 | 0.03538242 |
| pseudoscaff_850 | 42500 | 0.000907014 | 0.031918 | 0.032518683 | 0.031342718 |
| pseudoscaff_850 | 47500 | 0.001803618 | 0.027077 | 0.026628236 | 0.02832379 |
| pseudoscaff_850 | 52500 | 0.001123919 | 0.03729 | 0.033197332 | 0.032472461 |
| pseudoscaff_850 | 67500 | 0 | 0.017168 | 0.035202425 | 0.034783581 |
| pseudoscaff_850 | 72500 | 0.001008932 | 0.034337 | 0.030219355 | 0.031379485 |
| pseudoscaff_850 | 77500 | 0.000550741 | 0.011938 | 0.016538084 | 0.015050785 |
| pseudoscaff_850 | 97500 | 0.000149333 | 0.031357 | 0.032507636 | 0.030155239 |
| pseudoscaff_850 | 107500 | 0.000111353 | 0.026114 | 0.025997054 | 0.025935311 |
| pseudoscaff_850 | 117500 | 0.000356461 | 0.025063 | 0.023675031 | 0.023062628 |
| pseudoscaff_850 | 122500 | 0.000173128 | 0.024915 | 0.025214057 | 0.020609917 |
| pseudoscaff_850 | 127500 | 0 | 0.021933 | 0.024502348 | 0.023710759 |
| pseudoscaff_1744 | 227500 | 0.000311547 | 0.014069 | 0.017653049 | 0.017666812 |
| pseudoscaff_1744 | 242500 | 0 | 0.022551 | 0.039293433 | 0.039545228 |
| pseudoscaff_1744 | 312500 | 0.001795524 | 0.031951 | 0.04058159 | 0.043801111 |
| pseudoscaff_1744 | 337500 | 0.000340394 | 0.008478 | 0.032495145 | 0.031392286 |
| pseudoscaff_1744 | 342500 | 0.000426943 | 0.023826 | 0.032162148 | 0.031362159 |
| pseudoscaff_1744 | 417500 | 0 | 0.024926 | 0.034667882 | 0.032136756 |
| pseudoscaff_1744 | 427500 | 0 | 0.001357 | 0.001210663 | 0.001196646 |
| pseudoscaff_1744 | 432500 | 0 | 0.006331 | 0.006421243 | 0.006684784 |
| pseudoscaff_1744 | 437500 | 0 | 0.009413 | 0.011126394 | 0.011774943 |
| pseudoscaff_1744 | 442500 | 0 | 0.013168 | 0.014146089 | 0.014750122 |
| pseudoscaff_1744 | 447500 | 0 | 0.02738 | 0.026144866 | 0.025782686 |
| pseudoscaff_1744 | 482500 | 0.001203428 | 0.008393 | 0.024307159 | 0.024043387 |
| pseudoscaff_1744 | 492500 | 0.000642013 | 0.034559 | 0.034054147 | 0.033195777 |
| pseudoscaff_1744 | 497500 | 0.000135864 | 0.036539 | 0.037071566 | 0.035330526 |
| pseudoscaff_1744 | 502500 | 0.000630826 | 0.017349 | 0.026870949 | 0.028998022 |
| pseudoscaff_1744 | 512500 | 0.001059758 | 0.033571 | 0.042427107 | 0.043414953 |
| pseudoscaff_1744 | 562500 | 0.000200247 | 0.024897 | 0.034374926 | 0.037408089 |
| pseudoscaff_1744 | 567500 | 0 | 0.021112 | 0.024719883 | 0.024210756 |
| pseudoscaff_1744 | 572500 | 0.000354018 | 0.021013 | 0.025373392 | 0.025495871 |
| pseudoscaff_1744 | 602500 | 0 | 0.022488 | 0.027176159 | 0.028765158 |
| pseudoscaff_1744 | 622500 | 0.000329291 | 0.039704 | 0.030883222 | 0.036848609 |
| pseudoscaff_3331 | 347500 | 0.000434295 | 0.02521 | 0.023913363 | 0.022628282 |
| pseudoscaff_3214 | 7500 | 0.000219739 | 0.014329 | 0.019782033 | 0.01849887 |
| pseudoscaff_3474 | 12500 | 0.001161112 | 0.004391 | 0.003206105 | 0.003287377 |
| pseudoscaff_3853 | 442500 | 0.001077398 | 0.005624 | 0.007431976 | 0.007130039 |
| pseudoscaff_3853 | 462500 | 0.001423794 | 0.013402 | 0.008840678 | 0.010239379 |
| pseudoscaff_2401 | 7500 | 0.000566144 | 0.011848 | 0.019245658 | 0.017645965 |
| pseudoscaff_2401 | 17500 | 0.000872272 | 0.029418 | 0.018845016 | 0.016991507 |
| pseudoscaff_1711 | 12500 | 0 | 0.01206 | 0.015222171 | 0.015505613 |
| pseudoscaff_1711 | 17500 | 0 | 0.014982 | 0.01837885 | 0.018367996 |
| pseudoscaff_1711 | 22500 | 0.000638561 | 0.01181 | 0.015433065 | 0.015736449 |
| pseudoscaff_1711 | 27500 | 0.001379358 | 0.026184 | 0.026604505 | 0.027817195 |
| pseudoscaff_1711 | 37500 | 0.000281017 | 0.025293 | 0.028839339 | 0.02721943 |
| pseudoscaff_1711 | 47500 | 0.000966386 | 0.024325 | 0.029937061 | 0.028059669 |
| pseudoscaff_1711 | 57500 | 0.00026748 | 0.01882 | 0.02681085 | 0.026023052 |
| pseudoscaff_1711 | 67500 | 0.001289859 | 0.030531 | 0.03603664 | 0.035971506 |
| pseudoscaff_1711 | 107500 | 0 | 0.010594 | 0.01337505 | 0.013650937 |
| pseudoscaff_1711 | 117500 | 0 | 0.002227 | 0.010923689 | 0.010449278 |

| | | | | | |
|---|---|---|---|---|---|
| pseudoscaff_1711 | 132500 | 0.000524599 | 0.022464 | 0.028315334 | 0.028510109 |
| pseudoscaff_1711 | 137500 | 0.000101652 | 0.026304 | 0.020315314 | 0.021885233 |
| pseudoscaff_1711 | 162500 | 0.000747799 | 0.021146 | 0.030682257 | 0.029829605 |
| pseudoscaff_1711 | 167500 | 0.000811971 | 0.038538 | 0.033225785 | 0.034173083 |
| pseudoscaff_1711 | 187500 | 0.000400247 | 0.021862 | 0.023163268 | 0.025484259 |
| pseudoscaff_1711 | 212500 | 0.001047912 | 0.0222 | 0.023962987 | 0.022328362 |
| pseudoscaff_1711 | 222500 | 0.000820631 | 0.014128 | 0.021064735 | 0.021027621 |
| pseudoscaff_1711 | 227500 | 0 | 0.01284 | 0.018020667 | 0.01789701 |
| pseudoscaff_1711 | 232500 | 0 | 0.021806 | 0.019635526 | 0.019305934 |
| pseudoscaff_1711 | 242500 | 0.000913613 | 0.015491 | 0.018483965 | 0.017783214 |
| pseudoscaff_1711 | 247500 | 0.00020726 | 0.016807 | 0.017047308 | 0.017682641 |
| pseudoscaff_1518 | 42500 | 0.000766288 | 0.016361 | 0.022909439 | 0.024063174 |
| pseudoscaff_3253 | 157500 | 0.001881944 | 0.036084 | 0.031947017 | 0.032125529 |
| pseudoscaff_225 | 92500 | 0.000359286 | 0.016532 | 0.028434655 | 0.028213453 |
| pseudoscaff_62 | 7500 | 0.000408 | 0.009307 | 0.011721896 | 0.010787292 |
| pseudoscaff_62 | 12500 | 0.00005235 | 0.009339 | 0.00861266 | 0.009613537 |
| pseudoscaff_62 | 42500 | 0.001664682 | 0.016622 | 0.015385416 | 0.018754095 |
| pseudoscaff_3679 | 2500 | 0.000095065 | 0.013098 | 0.012657104 | 0.012055016 |
| pseudoscaff_3679 | 7500 | 0.000056063 | 0.01182 | 0.013861168 | 0.013411123 |
| pseudoscaff_3679 | 12500 | 0.000725407 | 0.015741 | 0.013271727 | 0.012698371 |
| pseudoscaff_3838 | 37500 | 0.001086841 | 0.001308 | 0.002122493 | 0.001916102 |
| pseudoscaff_3838 | 47500 | 0.000465606 | 0.003383 | 0.005216594 | 0.004362996 |
| pseudoscaff_885 | 2500 | 0.000929612 | 0.042571 | 0.042423656 | 0.041636513 |
| pseudoscaff_885 | 42500 | 0.001441265 | 0.011953 | 0.010687031 | 0.010640822 |
| pseudoscaff_885 | 57500 | 0.001701283 | 0.012415 | 0.014035319 | 0.014336936 |
| pseudoscaff_885 | 62500 | 0.001786385 | 0.002379 | 0.006003862 | 0.00448146 |
| pseudoscaff_1964 | 2500 | 0.000190395 | 0.000511 | 0.000428546 | 0.000710307 |
| pseudoscaff_1162 | 7500 | 0.000397998 | 0.033214 | 0.016781048 | 0.016562786 |
| pseudoscaff_1162 | 12500 | 0.00104553 | 0.039546 | 0.017613148 | 0.018154254 |
| pseudoscaff_1162 | 17500 | 0.000074462 | 0.037314 | 0.019576348 | 0.019297175 |
| pseudoscaff_1162 | 22500 | 0 | 0.029567 | 0.016880609 | 0.016809722 |
| pseudoscaff_1162 | 27500 | 0.000105649 | 0.031735 | 0.015519497 | 0.015219462 |
| pseudoscaff_1162 | 32500 | 0.000562927 | 0.034639 | 0.017958786 | 0.019444443 |
| pseudoscaff_1162 | 37500 | 0 | 0.023636 | 0.013363922 | 0.012709406 |
| pseudoscaff_1162 | 42500 | 0.000133626 | 0.033211 | 0.032210436 | 0.032782729 |
| pseudoscaff_1162 | 47500 | 0 | 0.040803 | 0.030367452 | 0.031469363 |
| pseudoscaff_1162 | 52500 | 0.000048586 | 0.046401 | 0.037360984 | 0.037970833 |
| pseudoscaff_1162 | 57500 | 0.000155383 | 0.034317 | 0.029036007 | 0.027975028 |
| pseudoscaff_1162 | 62500 | 0.000405224 | 0.028152 | 0.023766146 | 0.024704298 |
| pseudoscaff_2123 | 12500 | 0.000406304 | 0.005549 | 0.009627145 | 0.011462109 |
| pseudoscaff_2123 | 22500 | 0 | 0.028848 | 0.013902598 | 0.018621452 |
| pseudoscaff_2123 | 37500 | 0.000189975 | 0.003384 | 0.005606349 | 0.006505648 |
| pseudoscaff_2123 | 42500 | 0 | 0.012788 | 0.005942151 | 0.008611574 |
| pseudoscaff_2123 | 47500 | 0.000368739 | 0.004842 | 0.00854482 | 0.008367772 |
| pseudoscaff_2123 | 52500 | 0 | 0.007365 | 0.009117019 | 0.009376052 |
| pseudoscaff_2123 | 72500 | 0.000456153 | 0.008974 | 0.011565618 | 0.011935089 |
| pseudoscaff_2123 | 82500 | 0.000226637 | 0.003075 | 0.006874951 | 0.006614468 |
| pseudoscaff_2123 | 92500 | 0.000408976 | 0.004356 | 0.005385431 | 0.005595647 |
| pseudoscaff_2123 | 97500 | 0.001052173 | 0.004087 | 0.00251366 | 0.003282764 |
| pseudoscaff_2123 | 102500 | 0.000131793 | 0.002005 | 0.00264702 | 0.003084696 |

| | | | | | |
|---|---|---|---|---|---|
| pseudoscaff_2123 | 112500 | 0 | 0.002437 | 0.001688441 | 0.00180724 |
| pseudoscaff_2123 | 117500 | 0.000702382 | 0.000322 | 0.000846266 | 0.001044606 |
| pseudoscaff_2123 | 122500 | 0.000325853 | 0.005702 | 0.009263793 | 0.009448305 |
| pseudoscaff_2123 | 207500 | 0.000689143 | 0.025876 | 0.025861375 | 0.027847121 |
| pseudoscaff_2123 | 212500 | 0.000229544 | 0.010523 | 0.01127764 | 0.010952915 |
| pseudoscaff_2911 | 47500 | 0.000106617 | 0.022194 | 0.03258126 | 0.03362763 |
| pseudoscaff_2911 | 52500 | 0 | 0.034912 | 0.036380884 | 0.039239164 |
| pseudoscaff_2911 | 57500 | 0.000118486 | 0.018339 | 0.02398809 | 0.026175269 |
| pseudoscaff_2911 | 72500 | 0.001200187 | 0.038082 | 0.045664285 | 0.046473783 |
| pseudoscaff_2911 | 87500 | 0.000312725 | 0.039157 | 0.033525661 | 0.03627579 |
| pseudoscaff_2911 | 92500 | 0.001717814 | 0.026201 | 0.031457679 | 0.029916616 |
| pseudoscaff_2911 | 97500 | 0.000954635 | 0.024916 | 0.023577943 | 0.024971375 |
| pseudoscaff_2911 | 102500 | 0 | 0.025838 | 0.024293909 | 0.025513679 |
| pseudoscaff_2911 | 112500 | 0.000344049 | 0.02942 | 0.037776634 | 0.039108617 |
| pseudoscaff_2911 | 117500 | 0 | 0.031225 | 0.030496005 | 0.028505984 |
| pseudoscaff_2911 | 122500 | 0.000883662 | 0.024408 | 0.0274901 | 0.025702872 |
| pseudoscaff_2911 | 127500 | 0.000062864 | 0.014298 | 0.021000872 | 0.01991511 |
| pseudoscaff_2911 | 132500 | 0 | 0.017224 | 0.023531418 | 0.025758201 |
| pseudoscaff_2911 | 137500 | 0.000137419 | 0.024135 | 0.021083059 | 0.022759691 |
| pseudoscaff_2911 | 142500 | 0.00035621 | 0.032779 | 0.026539798 | 0.030563442 |
| pseudoscaff_2911 | 147500 | 0.000493277 | 0.022007 | 0.023820245 | 0.024259801 |
| pseudoscaff_2911 | 152500 | 0.001374597 | 0.016324 | 0.027009741 | 0.02700138 |
| pseudoscaff_2911 | 157500 | 0.001124173 | 0.027956 | 0.030936569 | 0.026607268 |
| pseudoscaff_2911 | 162500 | 0.000745138 | 0.021482 | 0.020096924 | 0.021414658 |
| pseudoscaff_2911 | 167500 | 0.001367465 | 0.020465 | 0.021957547 | 0.024020042 |
| pseudoscaff_2911 | 172500 | 0 | 0.012261 | 0.014887522 | 0.015655208 |
| pseudoscaff_2911 | 177500 | 0.000554681 | 0.016902 | 0.01817103 | 0.018846771 |
| pseudoscaff_2911 | 182500 | 0 | 0.016629 | 0.018812148 | 0.02152564 |
| pseudoscaff_2911 | 187500 | 0.000851066 | 0.019865 | 0.019324101 | 0.02049593 |
| pseudoscaff_2911 | 192500 | 0.001334185 | 0.019348 | 0.024188171 | 0.025669044 |
| pseudoscaff_2911 | 197500 | 0 | 0.010196 | 0.008498635 | 0.009490872 |
| pseudoscaff_2911 | 202500 | 0.000185487 | 0.021186 | 0.023015069 | 0.023545213 |
| pseudoscaff_3714 | 17500 | 0.001381195 | 0.016687 | 0.008314203 | 0.008336186 |
| pseudoscaff_3714 | 42500 | 0.000473611 | 0.027928 | 0.021387908 | 0.024020625 |
| pseudoscaff_3999 | 17500 | 0.000254369 | 0.030824 | 0.022126212 | 0.02410602 |
| pseudoscaff_801 | 2500 | 0.000220237 | 0.006433 | 0.003600397 | 0.004836196 |
| pseudoscaff_801 | 12500 | 0.000552975 | 0.024322 | 0.022516186 | 0.020673379 |
| pseudoscaff_801 | 17500 | 0.000420678 | 0.021498 | 0.019853287 | 0.020488634 |
| pseudoscaff_144 | 22500 | 0 | 0.022848 | 0.013739482 | 0.013225659 |
| pseudoscaff_144 | 27500 | 0.001444165 | 0.012479 | 0.021318373 | 0.020958017 |
| pseudoscaff_3633 | 57500 | 0.000782555 | 0.01955 | 0.010601725 | 0.009808813 |
| pseudoscaff_3483 | 57500 | 0.001974073 | 0.014871 | 0.016385578 | 0.017709082 |
| pseudoscaff_3041 | 17500 | 0.001667868 | 0.004812 | 0.003612209 | 0.003767706 |
| pseudoscaff_1817 | 2500 | 0 | 0.008738 | 0.008559425 | 0.006549589 |
| pseudoscaff_1817 | 7500 | 0.001715186 | 0.010264 | 0.010693305 | 0.008706581 |
| pseudoscaff_1817 | 12500 | 0 | 0.00164 | 0.002590991 | 0.002485692 |
| pseudoscaff_1817 | 17500 | 0.000056002 | 0.002417 | 0.003812475 | 0.00348989 |
| pseudoscaff_1817 | 22500 | 0 | 0.004842 | 0.005063751 | 0.005315298 |
| pseudoscaff_1167 | 7500 | 0 | 0.015036 | 0.016337871 | 0.017506474 |
| pseudoscaff_1167 | 12500 | 0 | 0.011709 | 0.019837212 | 0.019783724 |

| | | | | | |
|---|---|---|---|---|---|
| pseudoscaff_1167 | 17500 | 0.000687461 | 0.020604 | 0.02693701 | 0.027554569 |
| pseudoscaff_1167 | 22500 | 0 | 0.026191 | 0.02278969 | 0.02461701 |
| pseudoscaff_1167 | 27500 | 0.00007185 | 0.00976 | 0.019706357 | 0.017834331 |
| pseudoscaff_1167 | 32500 | 0.000423522 | 0.012155 | 0.021842204 | 0.020961935 |
| pseudoscaff_1167 | 37500 | 0 | 0.026102 | 0.034885435 | 0.031929096 |
| pseudoscaff_1351 | 82500 | 0.001142327 | 0.015674 | 0.032479774 | 0.034232693 |
| pseudoscaff_1351 | 117500 | 0.001511483 | 0.026173 | 0.023981297 | 0.022998117 |
| pseudoscaff_1351 | 122500 | 0.001560337 | 0.042714 | 0.034639066 | 0.036666615 |
| pseudoscaff_1351 | 137500 | 0.000907705 | 0.005228 | 0.019941649 | 0.016494849 |
| pseudoscaff_1351 | 167500 | 0.000529566 | 0.030268 | 0.028353376 | 0.029251648 |
| pseudoscaff_2036 | 47500 | 0 | 0.013909 | 0.020290527 | 0.016255928 |
| pseudoscaff_2036 | 72500 | 0.000099351 | 0.018093 | 0.013241604 | 0.013452793 |
| pseudoscaff_2036 | 87500 | 0.000377319 | 0.014064 | 0.01766386 | 0.015800496 |
| pseudoscaff_2036 | 97500 | 0 | 0.011322 | 0.024741043 | 0.022848575 |
| pseudoscaff_2036 | 102500 | 0 | 0.013012 | 0.016810059 | 0.016339271 |
| pseudoscaff_2036 | 107500 | 0.000373517 | 0.01151 | 0.012463391 | 0.012791298 |
| pseudoscaff_2036 | 112500 | 0 | 0.011489 | 0.011710977 | 0.010789543 |
| pseudoscaff_2036 | 117500 | 0 | 0.017452 | 0.02528347 | 0.025307432 |
| pseudoscaff_2036 | 122500 | 0.000213761 | 0.025656 | 0.032342563 | 0.032676471 |
| pseudoscaff_2036 | 127500 | 0.001033861 | 0.016673 | 0.020876237 | 0.019783701 |
| pseudoscaff_2036 | 132500 | 0 | 0.0057 | 0.009929875 | 0.008955556 |
| pseudoscaff_2036 | 142500 | 0 | 0.01919 | 0.024543026 | 0.02365568 |
| pseudoscaff_2036 | 157500 | 0.001006535 | 0.027765 | 0.027527772 | 0.027266701 |
| pseudoscaff_2036 | 197500 | 0.001910494 | 0.027838 | 0.038252806 | 0.035324334 |
| pseudoscaff_2036 | 202500 | 0.000787443 | 0.01573 | 0.021793622 | 0.019813184 |
| pseudoscaff_2036 | 212500 | 0.000151074 | 0.013879 | 0.019896741 | 0.02161258 |
| pseudoscaff_2036 | 227500 | 0.00121473 | 0.007561 | 0.014613639 | 0.011460005 |
| pseudoscaff_2036 | 247500 | 0.001802829 | 0.030741 | 0.034538561 | 0.031416189 |
| pseudoscaff_2036 | 252500 | 0.001077717 | 0.032339 | 0.035264124 | 0.032819078 |
| pseudoscaff_2036 | 267500 | 0.000491647 | 0.03524 | 0.045800008 | 0.045021732 |
| pseudoscaff_4200 | 47500 | 0.000256599 | 0.002912 | 0.004110819 | 0.002858512 |
| pseudoscaff_4200 | 52500 | 0 | 0.002856 | 0.004306515 | 0.003934643 |
| pseudoscaff_2210 | 2500 | 0 | 0.007272 | 0.008017866 | 0.008545562 |
| pseudoscaff_2210 | 7500 | 0.000219452 | 0.005445 | 0.008844035 | 0.008712121 |
| pseudoscaff_2210 | 12500 | 0.000338317 | 0.006974 | 0.009579769 | 0.009429199 |
| pseudoscaff_2210 | 17500 | 0.00045881 | 0.012392 | 0.008668686 | 0.010183146 |
| pseudoscaff_2210 | 22500 | 0.000697236 | 0.009698 | 0.005949429 | 0.00723839 |
| pseudoscaff_2210 | 27500 | 0.000710624 | 0.004291 | 0.00409792 | 0.003899712 |
| pseudoscaff_3613 | 2500 | 0.000617304 | 0.02991 | 0.033949611 | 0.038740666 |
| pseudoscaff_3613 | 12500 | 0 | 0.032551 | 0.025214192 | 0.027321209 |
| pseudoscaff_400 | 2500 | 0 | 0.002947 | 0.007347566 | 0.007188819 |
| pseudoscaff_400 | 7500 | 0 | 0.002202 | 0.002433259 | 0.003482305 |
| pseudoscaff_400 | 12500 | 0.000750231 | 0.015797 | 0.020091534 | 0.021352306 |
| pseudoscaff_400 | 17500 | 0.000133682 | 0.030839 | 0.023185659 | 0.024108061 |
| pseudoscaff_400 | 22500 | 0.00045262 | 0.037188 | 0.032195061 | 0.037099288 |
| pseudoscaff_1027 | 2500 | 0.000096445 | 0.008415 | 0.009387163 | 0.010772884 |
| pseudoscaff_1027 | 12500 | 0.000057213 | 0.01989 | 0.025251692 | 0.024516787 |
| pseudoscaff_1027 | 32500 | 0.000114549 | 0.021731 | 0.020760487 | 0.021011346 |
| pseudoscaff_1027 | 37500 | 0 | 0.026237 | 0.027283506 | 0.028179161 |
| pseudoscaff_1027 | 42500 | 0.001546172 | 0.022881 | 0.022052344 | 0.022355553 |

| | | | | | |
|---|---|---|---|---|---|
| pseudoscaff_1027 | 47500 | 0.000397179 | 0.028015 | 0.025768404 | 0.028728695 |
| pseudoscaff_1027 | 52500 | 0.000055969 | 0.018265 | 0.024098408 | 0.02511854 |
| pseudoscaff_1027 | 57500 | 0.001179955 | 0.0188 | 0.019694535 | 0.021801433 |
| pseudoscaff_1027 | 62500 | 0.000705723 | 0.019872 | 0.02322959 | 0.022922393 |
| pseudoscaff_1027 | 162500 | 0.001680457 | 0.018064 | 0.022735831 | 0.025417219 |
| pseudoscaff_1027 | 167500 | 0.000837667 | 0.017466 | 0.019688922 | 0.024414924 |
| pseudoscaff_1027 | 172500 | 0 | 0.020119 | 0.021609706 | 0.024249059 |
| pseudoscaff_1027 | 242500 | 0 | 0.013336 | 0.012685472 | 0.011340929 |
| pseudoscaff_3943 | 17500 | 0.001281957 | 0.007531 | 0.004267921 | 0.005531304 |
| pseudoscaff_3943 | 22500 | 0.000085312 | 0.015394 | 0.011406201 | 0.013624566 |
| pseudoscaff_3943 | 27500 | 0 | 0.010134 | 0.010837199 | 0.010771526 |
| pseudoscaff_3943 | 32500 | 0.000097323 | 0.01825 | 0.011788674 | 0.016312429 |
| pseudoscaff_3943 | 42500 | 0.000272481 | 0.013762 | 0.014970347 | 0.016418313 |
| pseudoscaff_3943 | 52500 | 0.000507075 | 0.019316 | 0.016464323 | 0.016631533 |
| pseudoscaff_3943 | 67500 | 0.001220767 | 0.013334 | 0.022495505 | 0.020466532 |
| pseudoscaff_3943 | 72500 | 0.00097086 | 0.013894 | 0.009334731 | 0.009548262 |
| pseudoscaff_1264 | 2500 | 0.001851902 | 0.0062 | 0.007759675 | 0.006428985 |
| pseudoscaff_1264 | 7500 | 0.000470711 | 0.000565 | 0.00294112 | 0.002997347 |
| pseudoscaff_1548 | 177500 | 0.000246836 | 0.014591 | 0.016413325 | 0.016105405 |
| pseudoscaff_2256 | 2500 | 0.001301766 | 0.004514 | 0.00614111 | 0.006166801 |
| pseudoscaff_2309 | 177500 | 0.00098578 | 0.014261 | 0.015548679 | 0.013614501 |
| pseudoscaff_1921 | 137500 | 0.000247224 | 0.029427 | 0.030669965 | 0.031936327 |
| pseudoscaff_1921 | 142500 | 0.000096898 | 0.027853 | 0.026034159 | 0.028583474 |
| pseudoscaff_1921 | 162500 | 0.001283228 | 0.013734 | 0.023938194 | 0.023203321 |
| pseudoscaff_1921 | 167500 | 0 | 0.004382 | 0.023740232 | 0.021890079 |
| pseudoscaff_1921 | 172500 | 0.000176975 | 0.015248 | 0.014862995 | 0.010706307 |
| pseudoscaff_1921 | 267500 | 0.000752501 | 0.005846 | 0.018785471 | 0.017055861 |
| pseudoscaff_3278 | 7500 | 0 | 0.014777 | 0.021251212 | 0.021463817 |
| pseudoscaff_3278 | 12500 | 0.000067376 | 0.018932 | 0.021064293 | 0.021497153 |
| pseudoscaff_3278 | 57500 | 0 | 0.006794 | 0.008856786 | 0.009419018 |
| pseudoscaff_3278 | 62500 | 0.000599933 | 0.031342 | 0.033555054 | 0.030595386 |
| pseudoscaff_3278 | 67500 | 0 | 0.006956 | 0.019220614 | 0.018002279 |
| pseudoscaff_3278 | 72500 | 0 | 0.022837 | 0.021030027 | 0.021099225 |
| pseudoscaff_3278 | 87500 | 0.001261603 | 0.030471 | 0.027839528 | 0.02814887 |
| pseudoscaff_3278 | 107500 | 0.000560785 | 0.024024 | 0.034401515 | 0.037020655 |
| pseudoscaff_2607 | 42500 | 0.000390443 | 0.046435 | 0.042614169 | 0.043852562 |
| pseudoscaff_2607 | 52500 | 0 | 0.027648 | 0.027393541 | 0.029782978 |
| pseudoscaff_3001 | 12500 | 0.00099653 | 0.008537 | 0.009664854 | 0.009560813 |
| pseudoscaff_4203 | 12500 | 0.000523053 | 0.01612 | 0.028452509 | 0.029522371 |
| pseudoscaff_4203 | 17500 | 0 | 0.010657 | 0.030422267 | 0.025404539 |
| pseudoscaff_4007 | 7500 | 0.000132166 | 0.010752 | 0.011321368 | 0.010241823 |
| pseudoscaff_4007 | 17500 | 0.001741825 | 0.020769 | 0.019579297 | 0.017391449 |
| pseudoscaff_4007 | 22500 | 0.000133646 | 0.005567 | 0.005999657 | 0.006315417 |
| pseudoscaff_4007 | 27500 | 0 | 0.00905 | 0.008045369 | 0.009202827 |
| pseudoscaff_4007 | 32500 | 0.00065901 | 0.006615 | 0.006340894 | 0.00631227 |
| pseudoscaff_4007 | 37500 | 0 | 0.003472 | 0.002812626 | 0.002834106 |
| pseudoscaff_4007 | 42500 | 0.000168477 | 0.002328 | 0.004100725 | 0.004478587 |
| pseudoscaff_4007 | 52500 | 0.0003583 | 0.013196 | 0.011081563 | 0.0115147 |
| pseudoscaff_4007 | 62500 | 0.001675438 | 0.018762 | 0.012666799 | 0.009258677 |
| pseudoscaff_2750 | 67500 | 0 | 0.027313 | 0.034957233 | 0.034653584 |

| | | | | | |
|---|---|---|---|---|---|
| pseudoscaff_2750 | 72500 | 0 | 0.009539 | 0.016144776 | 0.016259174 |
| pseudoscaff_2750 | 82500 | 0.000492717 | 0.019333 | 0.023733939 | 0.022314272 |
| pseudoscaff_2749 | 2500 | 0.001397657 | 0.009556 | 0.009874754 | 0.010815198 |
| pseudoscaff_2749 | 17500 | 0 | 0.013242 | 0.00284571 | 0.002504981 |
| pseudoscaff_2749 | 22500 | 0 | 0.013981 | 0.005379723 | 0.004905629 |
| pseudoscaff_2634 | 12500 | 0 | 0.014676 | 0.019560671 | 0.020335817 |
| pseudoscaff_2634 | 17500 | 0.000305926 | 0.020283 | 0.022357137 | 0.023464401 |
| pseudoscaff_2634 | 22500 | 0.000430354 | 0.008681 | 0.014567302 | 0.015848362 |
| pseudoscaff_2634 | 27500 | 0.001179096 | 0.020292 | 0.026525797 | 0.025769964 |
| pseudoscaff_2634 | 32500 | 0 | 0.013109 | 0.019922641 | 0.019309521 |
| pseudoscaff_3003 | 2500 | 0.001909187 | 0.012373 | 0.005965987 | 0.007072675 |
| pseudoscaff_3003 | 7500 | 0.000732352 | 0.009783 | 0.007028244 | 0.0069301 |
| pseudoscaff_3003 | 12500 | 0 | 0.007359 | 0.004643068 | 0.004502793 |
| pseudoscaff_2175 | 27500 | 0.000641268 | 0.015404 | 0.016929526 | 0.01795908 |
| pseudoscaff_2175 | 72500 | 0.000778553 | 0.027179 | 0.028770627 | 0.029899555 |
| pseudoscaff_2175 | 92500 | 0.001286786 | 0.029573 | 0.031379622 | 0.03158803 |
| pseudoscaff_2175 | 102500 | 0.000272203 | 0.018647 | 0.016369018 | 0.016638008 |
| pseudoscaff_2175 | 107500 | 0.0004198 | 0.031472 | 0.033690371 | 0.032838961 |
| pseudoscaff_2175 | 117500 | 0.000765197 | 0.012724 | 0.012503132 | 0.013074048 |
| pseudoscaff_2175 | 132500 | 0.000530363 | 0.019141 | 0.016940377 | 0.0172182 |
| pseudoscaff_2175 | 147500 | 0.000274244 | 0.005697 | 0.008102438 | 0.008180076 |
| pseudoscaff_2175 | 152500 | 0 | 0.006031 | 0.003389705 | 0.003477579 |
| pseudoscaff_2175 | 157500 | 0.001570234 | 0.005579 | 0.018063753 | 0.019843323 |
| pseudoscaff_2175 | 162500 | 0.000349178 | 0.001995 | 0.003972606 | 0.004534776 |
| pseudoscaff_2175 | 167500 | 0 | 0.001825 | 0.004240852 | 0.004431167 |
| pseudoscaff_2175 | 177500 | 0 | 0.001068 | 0.001805996 | 0.001829752 |
| pseudoscaff_2175 | 182500 | 0 | 0.00031 | 0.002120107 | 0.001858905 |
| pseudoscaff_2175 | 187500 | 0 | 0.00153 | 0.00372067 | 0.002309308 |
| pseudoscaff_2175 | 197500 | 0 | 0.017438 | 0.008083239 | 0.008992906 |
| pseudoscaff_2175 | 212500 | 0.000874298 | 0.021387 | 0.029150423 | 0.031422961 |
| pseudoscaff_2175 | 217500 | 0.000564559 | 0.033878 | 0.032681089 | 0.033122879 |
| pseudoscaff_2175 | 222500 | 0.00058882 | 0.015348 | 0.02240179 | 0.023084294 |
| pseudoscaff_1134 | 7500 | 0 | 0.003667 | 0.010592023 | 0.009365279 |
| pseudoscaff_1134 | 12500 | 0 | 0.002393 | 0.002399688 | 0.00797761 |
| pseudoscaff_1134 | 17500 | 0 | 0.001836 | 0.001691936 | 0.002024766 |
| pseudoscaff_1134 | 32500 | 0.001241173 | 0.010586 | 0.012119455 | 0.012572834 |
| pseudoscaff_560 | 2500 | 0.00082696 | 0.001 | 0.003519813 | 0.002642606 |
| pseudoscaff_4135 | 122500 | 0.000488091 | 0.009852 | 0.013199573 | 0.015476166 |
| pseudoscaff_953 | 12500 | 0.000976601 | 0.043922 | 0.045851913 | 0.047407413 |
| pseudoscaff_953 | 32500 | 0.000098072 | 0.026933 | 0.0288908 | 0.029200376 |
| pseudoscaff_953 | 37500 | 0.000172925 | 0.026578 | 0.030749623 | 0.029330481 |
| pseudoscaff_953 | 42500 | 0.000350606 | 0.025189 | 0.025820453 | 0.02962618 |
| pseudoscaff_953 | 57500 | 0.000439571 | 0.035461 | 0.038259189 | 0.034266545 |
| pseudoscaff_3151 | 7500 | 0.000534445 | 0.032093 | 0.030884338 | 0.030855456 |
| pseudoscaff_3151 | 12500 | 0.000975687 | 0.027779 | 0.028764506 | 0.027934192 |
| pseudoscaff_3151 | 17500 | 0.000355812 | 0.026969 | 0.035095847 | 0.036383308 |
| pseudoscaff_3151 | 22500 | 0 | 0.022698 | 0.023140497 | 0.022816619 |
| pseudoscaff_3151 | 27500 | 0.000072256 | 0.024437 | 0.024119498 | 0.024831908 |
| pseudoscaff_2215 | 2500 | 0 | 0.006638 | 0.009947001 | 0.009537147 |
| pseudoscaff_2215 | 7500 | 0.000124665 | 0.016366 | 0.012248173 | 0.012150194 |

| | | | | | |
|---|---|---|---|---|---|
| pseudoscaff_2215 | 12500 | 0.001306856 | 0.013631 | 0.014606204 | 0.014445639 |
| pseudoscaff_2215 | 17500 | 0 | 0.008032 | 0.013151784 | 0.012005302 |
| pseudoscaff_2215 | 22500 | 0 | 0.012886 | 0.021764671 | 0.021329964 |
| pseudoscaff_2215 | 32500 | 0 | 0.027193 | 0.023076157 | 0.021745962 |
| pseudoscaff_2215 | 42500 | 0.000224193 | 0.015969 | 0.020340386 | 0.019614031 |
| pseudoscaff_2215 | 47500 | 0.000178366 | 0.017086 | 0.024499641 | 0.024410874 |
| pseudoscaff_2215 | 57500 | 0 | 0.019314 | 0.02298232 | 0.023607085 |
| pseudoscaff_2215 | 67500 | 0 | 0.009097 | 0.015101667 | 0.015445248 |
| pseudoscaff_2215 | 77500 | 0.000820027 | 0.018725 | 0.01957223 | 0.019035967 |
| pseudoscaff_388 | 147500 | 0.000919609 | 0.020912 | 0.027062637 | 0.029060919 |
| pseudoscaff_2208 | 2500 | 0.000478885 | 0.010241 | 0.012656195 | 0.015582362 |
| pseudoscaff_2208 | 7500 | 0.000580921 | 0.010117 | 0.015938143 | 0.0125962 |
| pseudoscaff_2208 | 27500 | 0.000068008 | 0.029751 | 0.014228717 | 0.017729796 |
| pseudoscaff_2208 | 32500 | 0 | 0.026829 | 0.011578864 | 0.011105094 |
| pseudoscaff_2208 | 37500 | 0.000715928 | 0.011071 | 0.010848735 | 0.015958621 |
| pseudoscaff_2208 | 47500 | 0.001311855 | 0.009617 | 0.009939795 | 0.010491564 |
| pseudoscaff_2208 | 52500 | 0 | 0.011503 | 0.019946643 | 0.022028277 |
| pseudoscaff_2208 | 62500 | 0 | 0.02498 | 0.016294457 | 0.023408017 |
| pseudoscaff_2208 | 72500 | 0.00105067 | 0.014397 | 0.023762526 | 0.024874199 |
| pseudoscaff_2208 | 77500 | 0.000533175 | 0.009855 | 0.01495594 | 0.013048904 |
| pseudoscaff_2208 | 92500 | 0.000530394 | 0.008139 | 0.005484067 | 0.005289692 |
| pseudoscaff_2208 | 102500 | 0 | 0.004931 | 0.005427546 | 0.006223355 |
| pseudoscaff_2580 | 82500 | 0.000105635 | 0.001009 | 0.001163708 | 0.00166632 |
| pseudoscaff_2580 | 87500 | 0.000391523 | 0.000416 | 0.000890822 | 0.000665818 |
| pseudoscaff_2491 | 32500 | 0.000767059 | 0.014764 | 0.011762614 | 0.012337633 |
| pseudoscaff_2491 | 42500 | 0.001342556 | 0.011854 | 0.009812643 | 0.008847615 |
| pseudoscaff_3872 | 7500 | 0.001889331 | 0.017674 | 0.019377681 | 0.019217093 |
| pseudoscaff_3872 | 67500 | 0.001479005 | 0.035475 | 0.034531634 | 0.03402549 |
| pseudoscaff_3872 | 82500 | 0.001791184 | 0.022365 | 0.020429175 | 0.019236661 |
| pseudoscaff_3872 | 112500 | 0.000054393 | 0.00255 | 0.003185712 | 0.003541427 |
| pseudoscaff_3872 | 122500 | 0 | 0.000836 | 0.001250024 | 0.001198242 |
| pseudoscaff_3872 | 127500 | 0.000083367 | 0.001209 | 0.001232491 | 0.001355658 |
| pseudoscaff_3872 | 132500 | 0.000655805 | 0.000876 | 0.004463247 | 0.002953521 |
| pseudoscaff_3872 | 137500 | 0.000114908 | 0.000829 | 0.000777591 | 0.000796906 |
| pseudoscaff_3872 | 142500 | 0 | 0.002289 | 0.001018275 | 0.00106752 |
| pseudoscaff_3872 | 147500 | 0.000391503 | 0.004985 | 0.00313157 | 0.003223952 |
| pseudoscaff_3872 | 157500 | 0.001157828 | 0.003076 | 0.008076655 | 0.005997202 |
| pseudoscaff_3872 | 167500 | 0.000897838 | 0.004304 | 0.006866335 | 0.006471521 |
| pseudoscaff_3872 | 177500 | 0.001490976 | 0.004528 | 0.009567778 | 0.009315183 |
| pseudoscaff_3872 | 182500 | 0.00006782 | 0.003736 | 0.003889856 | 0.003620838 |
| pseudoscaff_3872 | 207500 | 0.000972643 | 0.019092 | 0.018781597 | 0.015981021 |
| pseudoscaff_3872 | 252500 | 0.001052584 | 0.006444 | 0.014446605 | 0.013947167 |
| pseudoscaff_3872 | 272500 | 0.000444972 | 0.014902 | 0.015548614 | 0.017322861 |
| pseudoscaff_3872 | 282500 | 0 | 0.020557 | 0.012061385 | 0.010221011 |
| pseudoscaff_3872 | 287500 | 0.000178513 | 0.013517 | 0.005904673 | 0.007837285 |
| pseudoscaff_586 | 7500 | 0.001665416 | 0.029678 | 0.02596411 | 0.0246319 |
| pseudoscaff_602 | 87500 | 0.000354033 | 0.029897 | 0.031750835 | 0.029649073 |
| pseudoscaff_602 | 92500 | 0.001609872 | 0.024307 | 0.032008457 | 0.034915201 |
| pseudoscaff_472 | 2500 | 0.000358198 | 0.024944 | 0.030743096 | 0.029033139 |
| pseudoscaff_472 | 7500 | 0 | 0.040097 | 0.033748571 | 0.028623771 |

| | | | | | |
|---|---|---|---|---|---|
| pseudoscaff_3142 | 2500 | 0.000051545 | 0.020489 | 0.014191359 | 0.018129265 |
| pseudoscaff_3154 | 42500 | 0 | 0.007972 | 0.011481517 | 0.009553482 |
| pseudoscaff_3154 | 47500 | 0 | 0.021223 | 0.023461985 | 0.0239415 |
| pseudoscaff_3154 | 82500 | 0.000131585 | 0.016472 | 0.014296283 | 0.015540002 |
| pseudoscaff_3154 | 92500 | 0 | 0.032774 | 0.037594988 | 0.038343404 |
| pseudoscaff_3154 | 107500 | 0.000056875 | 0.026805 | 0.029527968 | 0.033878508 |
| pseudoscaff_3154 | 112500 | 0 | 0.018405 | 0.020833988 | 0.020105228 |
| pseudoscaff_3154 | 117500 | 0 | 0.017362 | 0.016206016 | 0.015719312 |
| pseudoscaff_3154 | 132500 | 0.000470058 | 0.02208 | 0.022103225 | 0.021167156 |
| pseudoscaff_3154 | 162500 | 0.000150221 | 0.015646 | 0.018340237 | 0.017633413 |
| pseudoscaff_3154 | 167500 | 0 | 0.022716 | 0.026625626 | 0.026411347 |
| pseudoscaff_3154 | 172500 | 0 | 0.021663 | 0.028663976 | 0.026725677 |
| pseudoscaff_3154 | 177500 | 0.000146746 | 0.016386 | 0.019283277 | 0.021099098 |
| pseudoscaff_3154 | 182500 | 0 | 0.02038 | 0.020483232 | 0.020247514 |
| pseudoscaff_3154 | 222500 | 0 | 0.012711 | 0.016488347 | 0.015426616 |
| pseudoscaff_3154 | 227500 | 0.000183756 | 0.026647 | 0.030644532 | 0.032373599 |
| pseudoscaff_3154 | 232500 | 0.00035768 | 0.019204 | 0.024518347 | 0.023985494 |
| pseudoscaff_3154 | 237500 | 0.000438763 | 0.034335 | 0.031395231 | 0.03115471 |
| pseudoscaff_3154 | 242500 | 0 | 0.00699 | 0.009380013 | 0.009293995 |
| pseudoscaff_3154 | 247500 | 0.00172323 | 0.033559 | 0.035954516 | 0.038691716 |
| pseudoscaff_3154 | 272500 | 0 | 0.024086 | 0.030247394 | 0.030695221 |
| pseudoscaff_3154 | 427500 | 0 | 0.028926 | 0.020522792 | 0.023727322 |
| pseudoscaff_3154 | 437500 | 0.001943224 | 0.013904 | 0.01340778 | 0.012323795 |
| pseudoscaff_3154 | 447500 | 0.00086706 | 0.00141 | 0.022863247 | 0.022266671 |
| pseudoscaff_3154 | 452500 | 0.001301495 | 0.017464 | 0.02604282 | 0.032494664 |
| pseudoscaff_3154 | 497500 | 0.00027794 | 0.005221 | 0.017660611 | 0.017767651 |
| pseudoscaff_3154 | 587500 | 0.00127805 | 0.040497 | 0.044169185 | 0.045803886 |
| pseudoscaff_3154 | 627500 | 0.000758429 | 0.019271 | 0.029984105 | 0.030954646 |
| pseudoscaff_3154 | 662500 | 0.000113302 | 0.017892 | 0.023850089 | 0.025214712 |
| pseudoscaff_3154 | 667500 | 0 | 0.007037 | 0.01162462 | 0.0115365 |
| pseudoscaff_3154 | 687500 | 0 | 0.022197 | 0.029962835 | 0.029618252 |
| pseudoscaff_3154 | 692500 | 0 | 0.004371 | 0.004935403 | 0.004526285 |
| pseudoscaff_3154 | 722500 | 0.000247164 | 0.009251 | 0.009857421 | 0.010893912 |
| pseudoscaff_3154 | 727500 | 0 | 0.015504 | 0.022837036 | 0.023271777 |
| pseudoscaff_3154 | 742500 | 0.001839564 | 0.015571 | 0.021019401 | 0.019349679 |
| pseudoscaff_3154 | 747500 | 0.001605245 | 0.031778 | 0.027590687 | 0.025767288 |
| pseudoscaff_3154 | 757500 | 0.000177829 | 0.025578 | 0.026174665 | 0.025144676 |
| pseudoscaff_3154 | 762500 | 0.001118504 | 0.022391 | 0.016100301 | 0.015683423 |
| pseudoscaff_3154 | 767500 | 0 | 0.021122 | 0.019709498 | 0.02170314 |
| pseudoscaff_3154 | 772500 | 0 | 0.022429 | 0.019503103 | 0.019349634 |
| pseudoscaff_3154 | 777500 | 0.000127807 | 0.017524 | 0.019507257 | 0.019289627 |
| pseudoscaff_3154 | 782500 | 0.000672465 | 0.019259 | 0.016998868 | 0.01723081 |
| pseudoscaff_2783 | 2500 | 0.000871356 | 0.016004 | 0.011942338 | 0.010054511 |
| pseudoscaff_2783 | 22500 | 0.000323927 | 0.012504 | 0.013666673 | 0.013449869 |
| pseudoscaff_2040 | 7500 | 0 | 0.017583 | 0.023437944 | 0.024183349 |
| pseudoscaff_2040 | 32500 | 0 | 0.017128 | 0.023936959 | 0.02056865 |
| pseudoscaff_2040 | 42500 | 0 | 0.006511 | 0.018775057 | 0.016574311 |
| pseudoscaff_2040 | 47500 | 0.000130515 | 0.014954 | 0.015168121 | 0.014567035 |
| pseudoscaff_2040 | 52500 | 0.000285339 | 0.006324 | 0.00747656 | 0.008592876 |
| pseudoscaff_2040 | 57500 | 0 | 0.008523 | 0.008131165 | 0.008707707 |

| | | | | | |
|---|---|---|---|---|---|
| pseudoscaff_2040 | 62500 | 0.00144472 | 0.011621 | 0.025721124 | 0.020921687 |
| pseudoscaff_2040 | 67500 | 0.000583968 | 0.019788 | 0.013897565 | 0.012227827 |
| pseudoscaff_2040 | 72500 | 0 | 0.008853 | 0.010855117 | 0.010440196 |
| pseudoscaff_2040 | 77500 | 0 | 0.029709 | 0.014592874 | 0.017370568 |
| pseudoscaff_2040 | 87500 | 0.000409323 | 0.006003 | 0.005607469 | 0.005357209 |
| pseudoscaff_1666 | 17500 | 0.001151361 | 0.00033 | 0.002811136 | 0.00333622 |
| pseudoscaff_1666 | 22500 | 0.001624414 | 0.00229 | 0.007035182 | 0.009870728 |
| pseudoscaff_2492 | 82500 | 0.001436172 | 0.006489 | 0.004795926 | 0.00431595 |
| pseudoscaff_2492 | 87500 | 0.001074901 | 0.003149 | 0.003095309 | 0.002520945 |
| pseudoscaff_3556 | 2500 | 0 | 0.021751 | 0.025666522 | 0.024777826 |
| pseudoscaff_3556 | 7500 | 0 | 0.011844 | 0.015055555 | 0.015496329 |
| pseudoscaff_3556 | 12500 | 0 | 0.014075 | 0.013448881 | 0.01304908 |
| pseudoscaff_3556 | 22500 | 0.000155385 | 0.023491 | 0.019590832 | 0.01828846 |
| pseudoscaff_3556 | 27500 | 0.00004629 | 0.019307 | 0.015157626 | 0.015511308 |
| pseudoscaff_3556 | 32500 | 0 | 0.016118 | 0.014447818 | 0.014403665 |
| pseudoscaff_3556 | 37500 | 0 | 0.017515 | 0.020745911 | 0.02034831 |
| pseudoscaff_3556 | 42500 | 0.001586741 | 0.032067 | 0.032185969 | 0.032236571 |
| pseudoscaff_3556 | 52500 | 0.00087916 | 0.027465 | 0.038040239 | 0.038426027 |
| pseudoscaff_3556 | 62500 | 0 | 0.02499 | 0.028324966 | 0.030053865 |
| pseudoscaff_3556 | 72500 | 0.001256441 | 0.029879 | 0.034247339 | 0.035078426 |
| pseudoscaff_2509 | 152500 | 0.000153288 | 0.000189 | 0.000529161 | 0.000865143 |
| pseudoscaff_2509 | 157500 | 0 | 9.03E-05 | 0.001977286 | 0.002213864 |
| pseudoscaff_2509 | 162500 | 0 | 0.000362 | 0.000345155 | 0.000277737 |
| pseudoscaff_2509 | 177500 | 0.001405794 | 0.00179 | 0.00111349 | 0.001267595 |
| pseudoscaff_3459 | 7500 | 0.000300467 | 0.000508 | 0.001108044 | 0.001042787 |
| pseudoscaff_3459 | 17500 | 0.001758174 | 0.003583 | 0.002764155 | 0.004181672 |
| pseudoscaff_1777 | 212500 | 0.001644213 | 0.003154 | 0.004252826 | 0.002461887 |
| pseudoscaff_3648 | 27500 | 0.001252245 | 0.034349 | 0.035373934 | 0.029264765 |
| pseudoscaff_3697 | 2500 | 0.001034541 | 0.003437 | 0.004645164 | 0.004344788 |
| pseudoscaff_3697 | 12500 | 0.001354166 | 0.006444 | 0.006168383 | 0.006692506 |
| pseudoscaff_840 | 2500 | 0.001516963 | 0.039738 | 0.030445502 | 0.029919393 |
| pseudoscaff_840 | 37500 | 0.001451288 | 0.026387 | 0.025836231 | 0.025099945 |
| pseudoscaff_329 | 2500 | 0.00043474 | 0.014619 | 0.008756645 | 0.012737966 |
| pseudoscaff_329 | 7500 | 0.000483095 | 0.025116 | 0.01586278 | 0.01174365 |
| pseudoscaff_329 | 17500 | 0 | 0.003256 | 0.002137117 | 0.002480031 |
| pseudoscaff_329 | 22500 | 0 | 0.022258 | 0.007388365 | 0.01133046 |
| pseudoscaff_329 | 27500 | 0.00009982 | 0.016799 | 0.008254222 | 0.011868364 |
| pseudoscaff_329 | 47500 | 0.00106861 | 0.005122 | 0.013323625 | 0.011971417 |
| pseudoscaff_1190 | 7500 | 0 | 0.005384 | 0.004661934 | 0.004409191 |
| pseudoscaff_1190 | 12500 | 0.000264136 | 0.003649 | 0.008076905 | 0.006090692 |
| pseudoscaff_1190 | 17500 | 0 | 0.004143 | 0.006920317 | 0.006040114 |
| pseudoscaff_1190 | 22500 | 0.000629334 | 0.016767 | 0.015826691 | 0.01539464 |
| pseudoscaff_1190 | 27500 | 0 | 0.012106 | 0.009379692 | 0.007391759 |
| pseudoscaff_1190 | 32500 | 0.000252228 | 0.013237 | 0.01305704 | 0.011643 |
| pseudoscaff_1190 | 37500 | 0.000421865 | 0.020301 | 0.026146363 | 0.026232548 |
| pseudoscaff_1190 | 42500 | 0 | 0.029195 | 0.023880195 | 0.020499023 |
| pseudoscaff_1190 | 47500 | 0.00010729 | 0.016813 | 0.023293839 | 0.016715592 |
| pseudoscaff_1190 | 52500 | 0.000112626 | 0.020717 | 0.019920972 | 0.018135974 |
| pseudoscaff_1190 | 57500 | 0.000137527 | 0.033132 | 0.027314761 | 0.031135609 |
| pseudoscaff_1190 | 67500 | 0.001888339 | 0.044136 | 0.046931246 | 0.044786634 |

| | | | | | |
|---|---|---|---|---|---|
| pseudoscaff_1190 | 72500 | 0.000610322 | 0.034275 | 0.037300546 | 0.04179596 |
| pseudoscaff_1190 | 82500 | 0.001979174 | 0.033877 | 0.043815796 | 0.045121466 |
| pseudoscaff_1190 | 102500 | 0 | 0.009864 | 0.025161787 | 0.025458134 |
| pseudoscaff_1190 | 107500 | 0.000200403 | 0.021655 | 0.034761848 | 0.027051798 |
| pseudoscaff_1190 | 142500 | 0 | 0.030794 | 0.028023611 | 0.030024836 |
| pseudoscaff_4201 | 2500 | 0.001703774 | 0.010945 | 0.019133957 | 0.01880903 |
| pseudoscaff_4201 | 17500 | 0.000305724 | 0.016872 | 0.017013083 | 0.017050771 |
| pseudoscaff_4201 | 22500 | 0.000277151 | 0.014424 | 0.014740287 | 0.015191955 |
| pseudoscaff_4201 | 27500 | 0.000151452 | 0.018682 | 0.022382059 | 0.023001457 |
| pseudoscaff_701 | 12500 | 0 | 0.006051 | 0.004291556 | 0.004361328 |
| pseudoscaff_701 | 22500 | 0 | 0.002115 | 0.00251826 | 0.002511696 |
| pseudoscaff_701 | 27500 | 0.000897489 | 0.000613 | 0.003898363 | 0.004131285 |
| pseudoscaff_701 | 37500 | 0.000130766 | 0.008992 | 0.008206776 | 0.006678868 |
| pseudoscaff_701 | 52500 | 0.000197562 | 0.00321 | 0.001625371 | 0.001637679 |
| pseudoscaff_701 | 67500 | 0 | 0.001734 | 0.00501768 | 0.004736887 |
| pseudoscaff_701 | 72500 | 0 | 0.000779 | 0.004289517 | 0.004610991 |
| pseudoscaff_701 | 77500 | 0.000951131 | 0.001069 | 0.003682531 | 0.003225174 |
| pseudoscaff_701 | 82500 | 0.000995906 | 0.003658 | 0.002290202 | 0.002292337 |
| pseudoscaff_701 | 157500 | 0.001782655 | 0.009362 | 0.012712255 | 0.012169507 |
| pseudoscaff_701 | 167500 | 0.000185371 | 0.002026 | 0.005851185 | 0.005392342 |
| pseudoscaff_701 | 202500 | 0.001337882 | 0.022057 | 0.020294155 | 0.02345547 |
| pseudoscaff_701 | 222500 | 0.000175348 | 0.013473 | 0.008345772 | 0.010595083 |
| pseudoscaff_1767 | 2500 | 0.000257181 | 0.003571 | 0.001784951 | 0.002432842 |
| pseudoscaff_1767 | 7500 | 0.001454217 | 0.00186 | 0.002435842 | 0.002469816 |
| pseudoscaff_1767 | 17500 | 0 | 0.007982 | 0.003010379 | 0.00526238 |
| pseudoscaff_1767 | 22500 | 0.001166175 | 0.016235 | 0.005196061 | 0.010724784 |
| pseudoscaff_1767 | 27500 | 0.000478345 | 0.003307 | 0.007019447 | 0.007256735 |
| pseudoscaff_1767 | 37500 | 0.000143476 | 0.00377 | 0.008575957 | 0.010885232 |
| pseudoscaff_1767 | 52500 | 0.000862428 | 0.001052 | 0.002127542 | 0.002034356 |
| pseudoscaff_1767 | 57500 | 0.000437967 | 0.001841 | 0.002055619 | 0.002078009 |
| pseudoscaff_1767 | 62500 | 0 | 0.001396 | 0.006285335 | 0.004586792 |
| pseudoscaff_3973 | 22500 | 0.001475218 | 0.019313 | 0.020633241 | 0.021640716 |
| pseudoscaff_3973 | 27500 | 0 | 0.013857 | 0.01263382 | 0.013551971 |
| pseudoscaff_3973 | 32500 | 0.00010777 | 0.009105 | 0.011902782 | 0.012484536 |
| pseudoscaff_3973 | 37500 | 0.000116308 | 0.02453 | 0.027497252 | 0.024149716 |
| pseudoscaff_3973 | 47500 | 0.000193539 | 0.021629 | 0.020511532 | 0.01928447 |
| pseudoscaff_1247 | 27500 | 0.00048673 | 0.022943 | 0.029850308 | 0.036293617 |
| pseudoscaff_1247 | 32500 | 0 | 0.028313 | 0.023937831 | 0.024494398 |
| pseudoscaff_1247 | 42500 | 0.000152772 | 0.021927 | 0.02522203 | 0.02623918 |
| pseudoscaff_1247 | 47500 | 0.000509138 | 0.022026 | 0.028298842 | 0.026195611 |
| pseudoscaff_1247 | 92500 | 0.001820842 | 0.037551 | 0.036590254 | 0.037156584 |
| pseudoscaff_1247 | 102500 | 0.000937149 | 0.030453 | 0.029779704 | 0.028904125 |
| pseudoscaff_1247 | 182500 | 0.000707432 | 0.029736 | 0.036628237 | 0.031796009 |
| pseudoscaff_1247 | 202500 | 0 | 0.028132 | 0.02607383 | 0.026581981 |
| pseudoscaff_1247 | 272500 | 0.001307535 | 0.029423 | 0.031071718 | 0.033261058 |
| pseudoscaff_1247 | 297500 | 0.000296247 | 0.019013 | 0.036092563 | 0.03424181 |
| pseudoscaff_1247 | 417500 | 0.001457395 | 0.02415 | 0.034283277 | 0.036599085 |
| pseudoscaff_1247 | 587500 | 0.000089188 | 0.026048 | 0.027353539 | 0.028284239 |
| pseudoscaff_1247 | 597500 | 0 | 0.029907 | 0.038351786 | 0.037692968 |
| pseudoscaff_1247 | 602500 | 0.00151245 | 0.020299 | 0.022928707 | 0.027108657 |

| | | | | | |
|---|---|---|---|---|---|
| pseudoscaff_1247 | 607500 | 0.001177012 | 0.035342 | 0.032440187 | 0.033800366 |
| pseudoscaff_1247 | 617500 | 0.000449764 | 0.02503 | 0.025335179 | 0.025573403 |
| pseudoscaff_1247 | 622500 | 0.000065102 | 0.01361 | 0.015018083 | 0.016013449 |
| pseudoscaff_1247 | 627500 | 0.000532392 | 0.016384 | 0.015815498 | 0.017305154 |
| pseudoscaff_1247 | 632500 | 0.000236088 | 0.018759 | 0.016747328 | 0.016698746 |
| pseudoscaff_1247 | 647500 | 0.000273563 | 0.022992 | 0.024642427 | 0.025509467 |
| pseudoscaff_1247 | 652500 | 0.001515038 | 0.020669 | 0.023748557 | 0.023586219 |
| pseudoscaff_1247 | 662500 | 0 | 0.011469 | 0.013245256 | 0.012917461 |
| pseudoscaff_1247 | 667500 | 0.000366909 | 0.018715 | 0.017139466 | 0.016897028 |
| pseudoscaff_333 | 12500 | 0.000867201 | 0.007125 | 0.013023296 | 0.015279899 |
| pseudoscaff_1254 | 2500 | 0 | 0.005109 | 0.010794986 | 0.010353223 |
| pseudoscaff_569 | 7500 | 0.001112749 | 0.006271 | 0.013399601 | 0.01335988 |
| pseudoscaff_569 | 17500 | 0.000082134 | 0.03067 | 0.009133185 | 0.008786078 |
| pseudoscaff_569 | 27500 | 0 | 0.001563 | 0.003473589 | 0.004115878 |
| pseudoscaff_569 | 37500 | 0 | 0.004445 | 0.003235024 | 0.003736237 |
| pseudoscaff_569 | 67500 | 0.000525899 | 0.036774 | 0.022510914 | 0.027103537 |
| pseudoscaff_569 | 72500 | 0.001139263 | 0.008851 | 0.022103879 | 0.019417888 |
| pseudoscaff_569 | 77500 | 0.000991775 | 0.006588 | 0.011317988 | 0.010494385 |
| pseudoscaff_569 | 82500 | 0 | 0.006244 | 0.005096516 | 0.005302422 |
| pseudoscaff_569 | 87500 | 0 | 0.012769 | 0.010619091 | 0.010208025 |
| pseudoscaff_569 | 92500 | 0 | 0.011997 | 0.009442526 | 0.012190087 |
| pseudoscaff_411 | 12500 | 0.000692127 | 0.000792 | 0.000892037 | 0.000759915 |
| pseudoscaff_411 | 17500 | 0.000962497 | 0.001189 | 0.001212241 | 0.001501178 |
| pseudoscaff_1104 | 2500 | 0.000602171 | 0.032178 | 0.028368019 | 0.027991068 |
| pseudoscaff_1104 | 7500 | 0 | 0.038925 | 0.028024931 | 0.029964127 |
| pseudoscaff_1104 | 12500 | 0 | 0.017326 | 0.012889714 | 0.013794666 |
| pseudoscaff_1104 | 17500 | 0 | 0.019719 | 0.023999715 | 0.024778859 |
| pseudoscaff_1104 | 22500 | 0 | 0.014625 | 0.017244719 | 0.016571799 |
| pseudoscaff_1104 | 27500 | 0 | 0.012987 | 0.015855266 | 0.016183535 |
| pseudoscaff_875 | 2500 | 0.001536938 | 0.003862 | 0.006931222 | 0.007117767 |
| pseudoscaff_4101 | 17500 | 0.001143441 | 0.001551 | 0.001641194 | 0.002616921 |
| pseudoscaff_4101 | 22500 | 0.001634079 | 0.001 | 0.001680287 | 0.001998701 |
| pseudoscaff_4101 | 32500 | 0.001553986 | 0.005499 | 0.008082984 | 0.005593608 |
| pseudoscaff_848 | 12500 | 0.00082647 | 0.03033 | 0.03017208 | 0.028785858 |
| pseudoscaff_848 | 17500 | 0.001761009 | 0.027991 | 0.032343964 | 0.031676152 |
| pseudoscaff_848 | 22500 | 0.000438792 | 0.008153 | 0.027237603 | 0.021819468 |
| pseudoscaff_848 | 32500 | 0 | 0.019694 | 0.02214785 | 0.022167092 |
| pseudoscaff_848 | 37500 | 0.0001031 | 0.021129 | 0.025098268 | 0.025092378 |
| pseudoscaff_848 | 67500 | 0.00164773 | 0.023488 | 0.033163463 | 0.035134591 |
| pseudoscaff_848 | 82500 | 0.000163784 | 0.018885 | 0.023505783 | 0.022580829 |
| pseudoscaff_848 | 87500 | 0 | 0.018278 | 0.028517176 | 0.024851337 |
| pseudoscaff_848 | 92500 | 0.000528188 | 0.014643 | 0.019365298 | 0.016483918 |
| pseudoscaff_848 | 97500 | 0.001158873 | 0.010204 | 0.011507272 | 0.010604045 |
| pseudoscaff_848 | 102500 | 0.000554934 | 0.015342 | 0.015780949 | 0.013533313 |
| pseudoscaff_848 | 112500 | 0 | 0.023692 | 0.033730749 | 0.032281975 |
| pseudoscaff_848 | 117500 | 0.000342694 | 0.030925 | 0.039183198 | 0.036313178 |
| pseudoscaff_848 | 122500 | 0.000113069 | 0.032967 | 0.024246396 | 0.023686184 |
| pseudoscaff_848 | 132500 | 0 | 0.003365 | 0.012781063 | 0.008943161 |
| pseudoscaff_176 | 12500 | 0.001174448 | 0.004679 | 0.008900459 | 0.009042837 |
| pseudoscaff_2068 | 37500 | 0.001559077 | 0.001749 | 0.002620583 | 0.002485278 |

| | | | | | |
|---|---|---|---|---|---|
| pseudoscaff_2068 | 42500 | 0.00041556 | 0.003981 | 0.003570161 | 0.00356712 |
| pseudoscaff_15 | 2500 | 0.000572704 | 0.006098 | 0.006484628 | 0.006618637 |
| pseudoscaff_15 | 7500 | 0 | 0.002309 | 0.012831317 | 0.012241971 |
| pseudoscaff_15 | 12500 | 0 | 0.005423 | 0.005799241 | 0.0063302 |
| pseudoscaff_15 | 17500 | 0.000264238 | 0.013422 | 0.007702554 | 0.012162918 |
| pseudoscaff_15 | 22500 | 0.000482442 | 0.015026 | 0.009924358 | 0.012410479 |
| pseudoscaff_15 | 27500 | 0 | 0.020236 | 0.013629472 | 0.013665446 |
| pseudoscaff_3319 | 97500 | 0.001205159 | 0.001928 | 0.003686824 | 0.003825186 |
| pseudoscaff_4148 | 22500 | 0.000106202 | 0.012005 | 0.007405343 | 0.006406707 |
| pseudoscaff_597 | 72500 | 0.001371526 | 0.006347 | 0.006544964 | 0.00618113 |
| pseudoscaff_1489 | 107500 | 0.001918572 | 0.006437 | 0.005825419 | 0.005504807 |
| pseudoscaff_3085 | 2500 | 0.001054746 | 0.02595 | 0.0360944 | 0.038374282 |
| pseudoscaff_3085 | 17500 | 0 | 0.05548 | 0.042002231 | 0.041593625 |
| pseudoscaff_3085 | 22500 | 0.000981054 | 0.027987 | 0.033842342 | 0.032049747 |
| pseudoscaff_3085 | 77500 | 0 | 0.018278 | 0.026384349 | 0.027088578 |
| pseudoscaff_1175 | 172500 | 0.001847499 | 0.022056 | 0.02431244 | 0.024280184 |
| pseudoscaff_3270 | 12500 | 0 | 0.018749 | 0.020777223 | 0.019350996 |
| pseudoscaff_3270 | 52500 | 0.000819069 | 0.01459 | 0.019496981 | 0.019933217 |
| pseudoscaff_3270 | 62500 | 0.000532643 | 0.0189 | 0.019601382 | 0.019499375 |
| pseudoscaff_3270 | 67500 | 0 | 0.014375 | 0.019850719 | 0.020405808 |
| pseudoscaff_3270 | 77500 | 0.000383885 | 0.016688 | 0.018196243 | 0.021249823 |
| pseudoscaff_3270 | 87500 | 0 | 0.021174 | 0.026052723 | 0.028528064 |
| pseudoscaff_3270 | 97500 | 0.001954013 | 0.031528 | 0.029177897 | 0.028354693 |
| pseudoscaff_3270 | 107500 | 0.000587908 | 0.011972 | 0.012420492 | 0.011583565 |
| pseudoscaff_3270 | 122500 | 0 | 0.02296 | 0.028905083 | 0.030081695 |
| pseudoscaff_3270 | 132500 | 0 | 0.026165 | 0.025920282 | 0.024699223 |
| pseudoscaff_3270 | 162500 | 0 | 0.015903 | 0.019675837 | 0.019043518 |
| pseudoscaff_3270 | 167500 | 0.000185557 | 0.016712 | 0.015663461 | 0.016248144 |
| pseudoscaff_3270 | 172500 | 0.000986619 | 0.008803 | 0.015413886 | 0.017598751 |
| pseudoscaff_3270 | 187500 | 0.000290177 | 0.026565 | 0.029561636 | 0.027819281 |
| pseudoscaff_3270 | 197500 | 0.001889679 | 0.019674 | 0.020138523 | 0.020191781 |
| pseudoscaff_106 | 2500 | 0.000883363 | 0.000491 | 0.003550718 | 0.003291513 |
| pseudoscaff_3359 | 2500 | 0.000141433 | 0.012399 | 0.009057368 | 0.007693911 |
| pseudoscaff_3359 | 12500 | 0 | 0.011159 | 0.014848851 | 0.015598377 |
| pseudoscaff_3359 | 17500 | 0 | 0.008009 | 0.012025485 | 0.013003604 |
| pseudoscaff_3359 | 22500 | 0 | 0.017781 | 0.022239873 | 0.02185467 |
| pseudoscaff_3359 | 27500 | 0 | 0.01706 | 0.018954166 | 0.020760482 |
| pseudoscaff_3359 | 32500 | 0.000546545 | 0.017696 | 0.023406477 | 0.02223639 |
| pseudoscaff_3359 | 37500 | 0.0004148 | 0.019484 | 0.018979289 | 0.019604421 |
| pseudoscaff_3359 | 42500 | 0.000253627 | 0.02676 | 0.036761914 | 0.037896522 |
| pseudoscaff_3359 | 57500 | 0.000187033 | 0.037424 | 0.034741883 | 0.034786113 |
| pseudoscaff_3359 | 62500 | 0.001371231 | 0.02641 | 0.028240472 | 0.027643768 |
| pseudoscaff_3359 | 67500 | 0.000221252 | 0.011541 | 0.032050171 | 0.031058245 |
| pseudoscaff_3359 | 72500 | 0.000075791 | 0.033759 | 0.03845363 | 0.037771789 |
| pseudoscaff_3359 | 82500 | 0.000364619 | 0.019212 | 0.027221939 | 0.026206972 |
| pseudoscaff_2159 | 12500 | 0.000788962 | 0.00567 | 0.004892436 | 0.004611003 |
| pseudoscaff_1704 | 2500 | 0.001419557 | 0.007005 | 0.010940447 | 0.01178225 |
| pseudoscaff_3881 | 17500 | 0.000656536 | 0.002061 | 0.004433027 | 0.006723616 |
| pseudoscaff_3881 | 42500 | 0.001512079 | 0.007799 | 0.006330238 | 0.007562738 |
| pseudoscaff_3881 | 57500 | 0.000647987 | 0.002247 | 0.00573069 | 0.005450852 |

| | | | | | |
|---|---|---|---|---|---|
| pseudoscaff_3881 | 67500 | 0.00039436 | 0.015679 | 0.013725548 | 0.012937139 |
| pseudoscaff_3881 | 87500 | 0.001216208 | 0.011499 | 0.021284842 | 0.018328264 |
| pseudoscaff_3881 | 97500 | 0.001684761 | 0.007441 | 0.009331789 | 0.007928633 |
| pseudoscaff_3881 | 102500 | 0.001601513 | 0.005322 | 0.008180801 | 0.006698161 |
| pseudoscaff_3881 | 107500 | 0.000256556 | 0.004752 | 0.005979667 | 0.006133515 |
| pseudoscaff_3881 | 112500 | 0.000136494 | 0.003147 | 0.005552897 | 0.00529802 |
| pseudoscaff_3881 | 117500 | 0 | 0.008519 | 0.00913418 | 0.009646683 |
| pseudoscaff_3881 | 152500 | 0.001997726 | 0.013348 | 0.027954058 | 0.027725606 |
| pseudoscaff_3389 | 42500 | 0.000224212 | 0.021937 | 0.032439073 | 0.033462412 |
| pseudoscaff_3389 | 47500 | 0.001815379 | 0.025143 | 0.036175337 | 0.03595812 |
| pseudoscaff_3389 | 57500 | 0 | 0.033883 | 0.029695188 | 0.028583823 |
| pseudoscaff_3389 | 62500 | 0 | 0.037983 | 0.037311953 | 0.037531162 |
| pseudoscaff_3389 | 72500 | 0.000295 | 0.021849 | 0.029737 | 0.029999036 |
| pseudoscaff_3389 | 82500 | 0.000220306 | 0.026834 | 0.039655236 | 0.034065743 |
| pseudoscaff_3389 | 87500 | 0.001033611 | 0.024697 | 0.033195054 | 0.031358065 |
| pseudoscaff_3389 | 97500 | 0.000980414 | 0.030191 | 0.032402792 | 0.032056984 |
| pseudoscaff_899 | 292500 | 0 | 0.010048 | 0.015576763 | 0.016302343 |
| pseudoscaff_899 | 302500 | 0.001464194 | 0.014429 | 0.013171621 | 0.014329162 |
| pseudoscaff_899 | 307500 | 0 | 0.007769 | 0.013691232 | 0.014027997 |
| pseudoscaff_899 | 312500 | 0.000803762 | 0.021 | 0.017299597 | 0.018387309 |
| pseudoscaff_899 | 317500 | 0 | 0.013513 | 0.014766994 | 0.015050031 |
| pseudoscaff_899 | 322500 | 0.000649331 | 0.017008 | 0.016870886 | 0.018378869 |
| pseudoscaff_899 | 327500 | 0 | 0.01682 | 0.017797246 | 0.017483574 |
| pseudoscaff_899 | 332500 | 0 | 0.013248 | 0.01257005 | 0.011410778 |
| pseudoscaff_899 | 337500 | 0.001942617 | 0.020777 | 0.023027595 | 0.02410956 |
| pseudoscaff_899 | 342500 | 0.000422416 | 0.026056 | 0.018593145 | 0.020173539 |
| pseudoscaff_899 | 347500 | 0 | 0.012551 | 0.01355131 | 0.014760155 |
| pseudoscaff_899 | 357500 | 0.00152354 | 0.016797 | 0.018203601 | 0.019668086 |
| pseudoscaff_899 | 362500 | 0 | 0.017509 | 0.016184602 | 0.017606193 |
| pseudoscaff_899 | 367500 | 0 | 0.015583 | 0.015410888 | 0.015843482 |
| pseudoscaff_899 | 372500 | 0 | 0.022971 | 0.030290875 | 0.031393545 |
| pseudoscaff_899 | 377500 | 0.000990444 | 0.015212 | 0.022350421 | 0.026044034 |
| pseudoscaff_899 | 382500 | 0 | 0.012116 | 0.01431828 | 0.015025759 |
| pseudoscaff_3391 | 142500 | 0.001789873 | 0.011867 | 0.012333968 | 0.012191289 |
| pseudoscaff_4127 | 2500 | 0.00102521 | 0.000284 | 0.000975584 | 0.001370213 |
| pseudoscaff_4127 | 12500 | 0.001212687 | 0.000802 | 0.001608634 | 0.001374537 |
| pseudoscaff_4127 | 22500 | 0.001559774 | 0.001408 | 0.00659953 | 0.004857635 |
| pseudoscaff_2174 | 37500 | 0.001595833 | 0.018432 | 0.017500759 | 0.015851846 |
| pseudoscaff_2174 | 62500 | 0.00055783 | 0.002866 | 0.001225697 | 0.001397514 |
| pseudoscaff_2174 | 67500 | 0.001895717 | 0.00844 | 0.002105379 | 0.002740956 |
| pseudoscaff_2174 | 72500 | 0.001143808 | 0.013551 | 0.01119983 | 0.009624225 |
| pseudoscaff_2174 | 77500 | 0.001111294 | 0.004369 | 0.002480507 | 0.002910795 |
| pseudoscaff_3133 | 77500 | 0 | 0.010659 | 0.010203451 | 0.011567389 |
| pseudoscaff_3133 | 82500 | 0 | 0.017757 | 0.018791853 | 0.019658608 |
| pseudoscaff_3133 | 87500 | 0 | 0.013833 | 0.020409836 | 0.020815991 |
| pseudoscaff_3133 | 97500 | 0 | 0.008049 | 0.012138004 | 0.011579547 |
| pseudoscaff_3133 | 102500 | 0.001341766 | 0.019072 | 0.017842494 | 0.018995769 |
| pseudoscaff_3133 | 107500 | 0.000064923 | 0.01113 | 0.016685458 | 0.018149392 |
| pseudoscaff_3133 | 112500 | 0 | 0.008024 | 0.013632779 | 0.014555895 |
| pseudoscaff_3133 | 117500 | 0.000076622 | 0.005883 | 0.009757691 | 0.010456039 |

| | | | | | |
|---|---|---|---|---|---|
| pseudoscaff_3133 | 122500 | 0.000199904 | 0.008898 | 0.0219472 | 0.022178714 |
| pseudoscaff_3133 | 127500 | 0.000150266 | 0.011722 | 0.016239818 | 0.015781423 |
| pseudoscaff_2825 | 102500 | 0.000730927 | 0.017874 | 0.019354168 | 0.018275805 |
| pseudoscaff_1280 | 72500 | 0.00016059 | 0.006084 | 0.005072424 | 0.005013989 |
| pseudoscaff_2294 | 42500 | 0.00011354 | 0.021169 | 0.023471286 | 0.023053665 |
| pseudoscaff_2294 | 47500 | 0 | 0.021899 | 0.032825975 | 0.033125198 |
| pseudoscaff_2294 | 187500 | 0 | 0.031629 | 0.038658069 | 0.034961094 |
| pseudoscaff_2294 | 192500 | 0 | 0.006396 | 0.009105023 | 0.007035712 |
| pseudoscaff_993 | 2500 | 0 | 0.001228 | 0.001684781 | 0.001739889 |
| pseudoscaff_993 | 12500 | 0.000658715 | 0.002219 | 0.007398361 | 0.006759719 |
| pseudoscaff_1304 | 147500 | 0 | 0.014055 | 0.014742119 | 0.015001761 |
| pseudoscaff_3045 | 2500 | 0.000261915 | 0.009377 | 0.01552415 | 0.015506112 |
| pseudoscaff_3045 | 7500 | 0.000159485 | 0.021441 | 0.016465543 | 0.017206703 |
| pseudoscaff_3045 | 17500 | 0 | 0.018648 | 0.008032384 | 0.009035848 |
| pseudoscaff_2937 | 37500 | 0.001255157 | 0.009334 | 0.007312717 | 0.007824133 |
| pseudoscaff_2937 | 52500 | 0.001472318 | 0.005023 | 0.008311842 | 0.007187301 |
| pseudoscaff_2937 | 82500 | 0.000550031 | 0.007592 | 0.010631021 | 0.009414004 |
| pseudoscaff_1686 | 42500 | 0.001650494 | 0.016702 | 0.012698689 | 0.012558666 |
| pseudoscaff_1842 | 12500 | 0.00029643 | 0.030827 | 0.037213301 | 0.03735059 |
| pseudoscaff_1842 | 17500 | 0.001004652 | 0.032131 | 0.040064907 | 0.039695614 |
| pseudoscaff_1842 | 32500 | 0 | 0.020542 | 0.025725714 | 0.027805234 |
| pseudoscaff_301 | 22500 | 0.000537869 | 0.026354 | 0.021821089 | 0.023934459 |
| pseudoscaff_301 | 32500 | 0.001547576 | 0.012136 | 0.018029846 | 0.02149588 |
| pseudoscaff_301 | 42500 | 0.000570722 | 0.00159 | 0.007242863 | 0.010050902 |
| pseudoscaff_301 | 62500 | 0 | 0.036826 | 0.03406356 | 0.034340436 |
| pseudoscaff_4014 | 2500 | 0.00175802 | 0.006555 | 0.004009443 | 0.006632661 |
| pseudoscaff_4014 | 12500 | 0.00116138 | 0.003648 | 0.004536964 | 0.004024256 |
| pseudoscaff_2305 | 32500 | 0.000144584 | 0.019493 | 0.020639011 | 0.019774429 |
| pseudoscaff_3011 | 17500 | 0.001309661 | 0.004545 | 0.008220195 | 0.008548048 |
| pseudoscaff_3011 | 32500 | 0.000554588 | 0.009573 | 0.009965455 | 0.008766186 |
| pseudoscaff_3011 | 52500 | 0.000287918 | 0.010658 | 0.007614229 | 0.009538054 |
| pseudoscaff_3011 | 57500 | 0.000131106 | 0.010694 | 0.011526358 | 0.011283155 |
| pseudoscaff_3011 | 87500 | 0 | 0.019318 | 0.010302231 | 0.008944771 |
| pseudoscaff_3011 | 92500 | 0 | 0.007953 | 0.009693117 | 0.009305212 |
| pseudoscaff_3011 | 97500 | 0.001025277 | 0.007159 | 0.006713573 | 0.008313493 |
| pseudoscaff_3011 | 102500 | 0.000544289 | 0.008879 | 0.013264998 | 0.01161907 |
| pseudoscaff_3011 | 107500 | 0 | 0.022711 | 0.009010855 | 0.013210882 |
| pseudoscaff_3011 | 112500 | 0.000511444 | 0.031973 | 0.014154165 | 0.014441537 |
| pseudoscaff_3011 | 117500 | 0.000829383 | 0.013815 | 0.016326016 | 0.012636025 |
| pseudoscaff_925 | 2500 | 0 | 0.005398 | 0.009273468 | 0.008006879 |
| pseudoscaff_2143 | 12500 | 0.000335133 | 0.01962 | 0.020254897 | 0.019794766 |
| pseudoscaff_2143 | 17500 | 0 | 0.011437 | 0.012985231 | 0.013466702 |
| pseudoscaff_2143 | 22500 | 0 | 0.010464 | 0.01663219 | 0.018524866 |
| pseudoscaff_3338 | 47500 | 0.001681655 | 0.003658 | 0.005061662 | 0.002811357 |
| pseudoscaff_1065 | 7500 | 0.000946379 | 0.016861 | 0.026360934 | 0.025663602 |
| pseudoscaff_1065 | 12500 | 0 | 0.024046 | 0.024448679 | 0.025459428 |
| pseudoscaff_1065 | 17500 | 0.001936948 | 0.044047 | 0.035913035 | 0.03663021 |
| pseudoscaff_4020 | 7500 | 0.001352222 | 0.000961 | 0.002870155 | 0.002219555 |
| pseudoscaff_4020 | 37500 | 0.001897462 | 0.004961 | 0.005121657 | 0.005254887 |
| pseudoscaff_4020 | 42500 | 0.001399026 | 0.001121 | 0.002556498 | 0.002714451 |

| | | | | | |
|---|---|---|---|---|---|
| pseudoscaff_4020 | 52500 | 0.001608164 | 0.00567 | 0.006263905 | 0.006142561 |
| pseudoscaff_4020 | 82500 | 0.001680322 | 0.002273 | 0.001740487 | 0.002017723 |
| pseudoscaff_4020 | 92500 | 0.000144788 | 0 | 0.000526945 | 0.000404773 |
| pseudoscaff_4020 | 102500 | 0.00042665 | 0.001201 | 0.001776854 | 0.001678101 |
| pseudoscaff_2918 | 102500 | 0.001593648 | 0.011402 | 0.008481126 | 0.008201413 |
| pseudoscaff_2918 | 107500 | 0.001750499 | 0.01071 | 0.014794445 | 0.014663249 |
| pseudoscaff_2918 | 162500 | 0.001949069 | 0.006953 | 0.007396855 | 0.006807202 |
| pseudoscaff_2438 | 7500 | 0.000969141 | 0.03028 | 0.017571032 | 0.016475093 |
| pseudoscaff_2438 | 12500 | 0.000959313 | 0.00771 | 0.011650083 | 0.012011391 |
| pseudoscaff_2438 | 17500 | 0 | 0.002884 | 0.00270388 | 0.003232203 |
| pseudoscaff_2438 | 22500 | 0.000157962 | 0.004344 | 0.004521135 | 0.004313871 |
| pseudoscaff_2438 | 37500 | 0.000180052 | 0.009252 | 0.008402929 | 0.009208267 |
| pseudoscaff_95 | 2500 | 0.000267655 | 0.030411 | 0.035134623 | 0.038214974 |
| pseudoscaff_3067 | 312500 | 0 | 0.002612 | 0.004644318 | 0.004185945 |
| pseudoscaff_3067 | 317500 | 0 | 0.006553 | 0.005171203 | 0.005860676 |
| pseudoscaff_2551 | 7500 | 0 | 0.021495 | 0.002774101 | 0.004469568 |
| pseudoscaff_2551 | 17500 | 0.000286238 | 0.002634 | 0.000843985 | 0.000161076 |
| pseudoscaff_2551 | 22500 | 0.000601759 | 0.003719 | 0.000399386 | 0.000399635 |
| pseudoscaff_2551 | 27500 | 0.000310409 | 0.005128 | 0.000743381 | 0.000631932 |
| pseudoscaff_2551 | 37500 | 0.000897408 | 0.002037 | 0.000928987 | 0.001373999 |
| pseudoscaff_2551 | 42500 | 0 | 0.000276 | 0.0007205 | 0.000842723 |
| pseudoscaff_2551 | 47500 | 0 | 0.013871 | 0.000140985 | 0.000140347 |
| pseudoscaff_2551 | 62500 | 0.001971707 | 0.001876 | 0.0001047 | 0.000326689 |
| pseudoscaff_2737 | 27500 | 0 | 0.015486 | 0.017495999 | 0.015995101 |
| pseudoscaff_2737 | 32500 | 0 | 0.019345 | 0.021276566 | 0.021518168 |
| pseudoscaff_2737 | 37500 | 0 | 0.007257 | 0.010112959 | 0.010587636 |
| pseudoscaff_2737 | 42500 | 0.001545946 | 0.037557 | 0.036719432 | 0.034313675 |
| pseudoscaff_2737 | 57500 | 0.00078712 | 0.035713 | 0.047849082 | 0.047574605 |
| pseudoscaff_3494 | 2500 | 0.001429393 | 0.011679 | 0.019955343 | 0.019637205 |
| pseudoscaff_3494 | 7500 | 0 | 0.006795 | 0.011551427 | 0.010039096 |
| pseudoscaff_3494 | 12500 | 0.001338753 | 0.005386 | 0.007923109 | 0.007464021 |
| pseudoscaff_3494 | 22500 | 0.000921093 | 0.023801 | 0.012443718 | 0.012203559 |
| pseudoscaff_3494 | 27500 | 0.000370128 | 0.00987 | 0.007416367 | 0.004973326 |
| pseudoscaff_3494 | 102500 | 0.001443403 | 0.034845 | 0.05043559 | 0.043388345 |
| pseudoscaff_3494 | 212500 | 0 | 0.030261 | 0.033027493 | 0.032104058 |
| pseudoscaff_3494 | 217500 | 0 | 0.023686 | 0.031617863 | 0.033741934 |
| pseudoscaff_3494 | 247500 | 0.000711597 | 0.037969 | 0.045512302 | 0.047314335 |
| pseudoscaff_3494 | 267500 | 0.000275635 | 0.014726 | 0.024470316 | 0.024684178 |
| pseudoscaff_3494 | 287500 | 0.00026332 | 0.026788 | 0.032039455 | 0.030554339 |
| pseudoscaff_1673 | 7500 | 0.000863724 | 0.039727 | 0.040600904 | 0.03957493 |
| pseudoscaff_1673 | 12500 | 0 | 0.036726 | 0.030371074 | 0.029578776 |
| pseudoscaff_1673 | 32500 | 0.0002112 | 0.04239 | 0.035670372 | 0.039682322 |
| pseudoscaff_1673 | 42500 | 0 | 0.050231 | 0.043907732 | 0.041837795 |
| pseudoscaff_2821 | 2500 | 0 | 0.000604 | 0.001037916 | 0.000809581 |
| pseudoscaff_2821 | 22500 | 0.001753007 | 0.00861 | 0.005760572 | 0.005330225 |
| pseudoscaff_2821 | 27500 | 0.000125924 | 0.008407 | 0.008024584 | 0.00797552 |
| pseudoscaff_2821 | 32500 | 0.000187969 | 0.002792 | 0.003630025 | 0.004018593 |
| pseudoscaff_2821 | 42500 | 0.001290622 | 0.007472 | 0.010738924 | 0.009633656 |
| pseudoscaff_2821 | 47500 | 0.001140133 | 0.001977 | 0.00728568 | 0.006664899 |
| pseudoscaff_2821 | 52500 | 0 | 0.005026 | 0.003176834 | 0.003113321 |

| | | | | | |
|---|---|---|---|---|---|
| pseudoscaff_2821 | 67500 | 0.001152175 | 0.00846 | 0.008164965 | 0.011392427 |
| pseudoscaff_2821 | 72500 | 0.001260917 | 0.00668 | 0.010335974 | 0.015356429 |
| pseudoscaff_977 | 7500 | 0.000939244 | 0.007959 | 0.006816228 | 0.006488426 |
| pseudoscaff_977 | 12500 | 0.000447147 | 0.003707 | 0.004542697 | 0.006627424 |
| pseudoscaff_977 | 22500 | 0 | 0.003979 | 0.003761884 | 0.003710229 |
| pseudoscaff_977 | 37500 | 0.001477098 | 0.014613 | 0.008392602 | 0.00863522 |
| pseudoscaff_977 | 42500 | 0.000204373 | 0.003533 | 0.005346857 | 0.00581205 |
| pseudoscaff_1405 | 2500 | 0.000597183 | 0.002841 | 0.000560208 | 0.00044669 |
| pseudoscaff_1405 | 12500 | 0.001216597 | 0.003141 | 0.001565415 | 0.001650611 |
| pseudoscaff_1405 | 32500 | 0.001668041 | 0.003995 | 0.002154836 | 0.002956085 |
| pseudoscaff_2490 | 72500 | 0.000620675 | 0.003967 | 0.003436069 | 0.003378239 |
| pseudoscaff_2490 | 82500 | 0.001782898 | 0.001677 | 0.00369631 | 0.0043982 |
| pseudoscaff_2490 | 87500 | 0.001820572 | 0.00462 | 0.004040693 | 0.004243192 |
| pseudoscaff_2490 | 132500 | 0.001363185 | 0.014091 | 0.013372616 | 0.010447068 |
| pseudoscaff_2490 | 142500 | 0.001986085 | 0.03164 | 0.015750733 | 0.017653674 |
| pseudoscaff_3987 | 97500 | 0.000430996 | 0.036911 | 0.038290469 | 0.040004364 |
| pseudoscaff_3179 | 7500 | 0.000703274 | 0.017013 | 0.021188911 | 0.021527603 |
| pseudoscaff_3179 | 12500 | 0.000246783 | 0.006921 | 0.007174564 | 0.007335068 |
| pseudoscaff_3179 | 22500 | 0.001201548 | 0.006444 | 0.005699772 | 0.005918535 |
| pseudoscaff_865 | 7500 | 0.000284758 | 0.0194 | 0.022973253 | 0.02179618 |
| pseudoscaff_865 | 17500 | 0.001047557 | 0.020702 | 0.021248731 | 0.023104792 |
| pseudoscaff_865 | 22500 | 0.00018397 | 0.010524 | 0.01748414 | 0.018544112 |
| pseudoscaff_865 | 27500 | 0.000159109 | 0.017465 | 0.023343061 | 0.020749685 |
| pseudoscaff_865 | 47500 | 0.000090735 | 0.026614 | 0.023236176 | 0.02364176 |
| pseudoscaff_865 | 52500 | 0 | 0.011884 | 0.023860133 | 0.024243142 |
| pseudoscaff_865 | 57500 | 0.001020344 | 0.009389 | 0.009155995 | 0.008379794 |
| pseudoscaff_865 | 62500 | 0 | 0.007654 | 0.021617954 | 0.014470044 |
| pseudoscaff_865 | 67500 | 0.000161814 | 0.010549 | 0.014031916 | 0.012959917 |
| pseudoscaff_865 | 72500 | 0.000364946 | 0.009306 | 0.011194448 | 0.011620394 |
| pseudoscaff_865 | 77500 | 0.000782473 | 0.015081 | 0.012775598 | 0.013689785 |
| pseudoscaff_865 | 82500 | 0.000669134 | 0.01195 | 0.010421578 | 0.010895071 |
| pseudoscaff_865 | 87500 | 0.000111099 | 0.002862 | 0.002490822 | 0.002913795 |
| pseudoscaff_865 | 92500 | 0.000082727 | 0.006195 | 0.009303294 | 0.009651568 |
| pseudoscaff_865 | 97500 | 0.000392211 | 0.017118 | 0.019267899 | 0.0179416 |
| pseudoscaff_865 | 107500 | 0.001215686 | 0.006683 | 0.01292708 | 0.012322781 |
| pseudoscaff_793 | 7500 | 0.00047704 | 0.003714 | 0.013972855 | 0.015029305 |
| pseudoscaff_793 | 12500 | 0.001076851 | 0.014461 | 0.019019094 | 0.017367069 |
| pseudoscaff_3548 | 7500 | 0 | 0.018172 | 0.026734841 | 0.028200462 |
| pseudoscaff_2703 | 22500 | 0.000396349 | 0.036755 | 0.045340688 | 0.046409427 |
| pseudoscaff_2703 | 127500 | 0.001455626 | 0.023193 | 0.03068641 | 0.02977144 |
| pseudoscaff_4171 | 197500 | 0.001700976 | 0.020364 | 0.021693267 | 0.022459134 |
| pseudoscaff_3164 | 107500 | 0.001405829 | 0.001951 | 0.001865418 | 0.002005391 |
| pseudoscaff_3164 | 112500 | 0.00133841 | 0.011532 | 0.007276916 | 0.007209679 |
| pseudoscaff_3164 | 132500 | 0.001307599 | 0.018882 | 0.00979348 | 0.012282802 |
| pseudoscaff_757 | 2500 | 0.001479715 | 0.030462 | 0.033448723 | 0.03575806 |
| pseudoscaff_757 | 12500 | 0.000528695 | 0.017846 | 0.033445501 | 0.035265087 |
| pseudoscaff_757 | 22500 | 0.001328083 | 0.021183 | 0.029903227 | 0.02669324 |
| pseudoscaff_617 | 7500 | 0 | 0.018208 | 0.035863405 | 0.034525775 |
| pseudoscaff_3716 | 2500 | 0.001821736 | 0.022004 | 0.013482391 | 0.014442329 |
| pseudoscaff_3716 | 12500 | 0.000957456 | 0.014639 | 0.012335341 | 0.012893069 |

| | | | | | |
|---|---|---|---|---|---|
| pseudoscaff_3716 | 17500 | 0.000316932 | 0.003773 | 0.004635989 | 0.004602717 |
| pseudoscaff_3716 | 42500 | 0.000643786 | 0.017391 | 0.008467416 | 0.008286078 |
| pseudoscaff_3716 | 52500 | 0.000125249 | 0.012627 | 0.009388864 | 0.005881405 |
| pseudoscaff_3716 | 57500 | 0 | 0.014185 | 0.006685834 | 0.009534668 |
| pseudoscaff_731 | 37500 | 0.001335366 | 0.006109 | 0.0080871 | 0.007910127 |
| pseudoscaff_2745 | 87500 | 0 | 0.034148 | 0.041139259 | 0.041851883 |
| pseudoscaff_2745 | 97500 | 0.000217997 | 0.015441 | 0.032127973 | 0.033532925 |
| pseudoscaff_2745 | 107500 | 0.000836641 | 0.025845 | 0.0268682 | 0.027620986 |
| pseudoscaff_2745 | 112500 | 0.000547524 | 0.018284 | 0.02712015 | 0.026400938 |
| pseudoscaff_2745 | 117500 | 0.000947022 | 0.026337 | 0.030181989 | 0.032886145 |
| pseudoscaff_2745 | 122500 | 0 | 0.029086 | 0.022662813 | 0.025798597 |
| pseudoscaff_2745 | 127500 | 0 | 0.027444 | 0.031225305 | 0.030657947 |
| pseudoscaff_2745 | 132500 | 0.000089888 | 0.030141 | 0.03171825 | 0.031363175 |
| pseudoscaff_2745 | 137500 | 0.000089615 | 0.021932 | 0.018137318 | 0.018734774 |
| pseudoscaff_2745 | 142500 | 0.001501801 | 0.035796 | 0.034562788 | 0.032201363 |
| pseudoscaff_3990 | 2500 | 0.000264399 | 0.021088 | 0.021452615 | 0.021292335 |
| pseudoscaff_3990 | 7500 | 0.000723123 | 0.017696 | 0.021493223 | 0.021690934 |
| pseudoscaff_3990 | 12500 | 0.001033014 | 0.024708 | 0.027289321 | 0.027886553 |
| pseudoscaff_3990 | 22500 | 0.001623459 | 0.016884 | 0.016857221 | 0.017032217 |
| pseudoscaff_3990 | 32500 | 0.001883662 | 0.026951 | 0.026415531 | 0.026380209 |
| pseudoscaff_3990 | 37500 | 0.000517908 | 0.017486 | 0.017487767 | 0.017251429 |
| pseudoscaff_3990 | 42500 | 0.001013153 | 0.007491 | 0.010857278 | 0.011347833 |
| pseudoscaff_1273 | 12500 | 0.000073813 | 0.007694 | 0.011808368 | 0.011828302 |
| pseudoscaff_1273 | 27500 | 0.000630136 | 0.005699 | 0.013988742 | 0.012768919 |
| pseudoscaff_1273 | 32500 | 0.000137357 | 0.005146 | 0.010914655 | 0.012055335 |
| pseudoscaff_1273 | 42500 | 0.001933924 | 0.032377 | 0.026356108 | 0.023123324 |
| pseudoscaff_1273 | 52500 | 0.000560357 | 0.004232 | 0.005220687 | 0.004734614 |
| pseudoscaff_1273 | 57500 | 0.000975913 | 0.016788 | 0.010582833 | 0.009815842 |
| pseudoscaff_1273 | 67500 | 0.00081042 | 0.001863 | 0.00701054 | 0.00486834 |
| pseudoscaff_1273 | 77500 | 0 | 0.0369 | 0.012659496 | 0.014886591 |
| pseudoscaff_1273 | 82500 | 0.001267967 | 0.007323 | 0.012953145 | 0.012325539 |
| pseudoscaff_1273 | 87500 | 0.000203357 | 0.010382 | 0.018710548 | 0.015786259 |
| pseudoscaff_1273 | 92500 | 0 | 0.008136 | 0.00746871 | 0.009195246 |
| pseudoscaff_1273 | 102500 | 0.000507871 | 0.008502 | 0.014434848 | 0.0172494 |
| pseudoscaff_1273 | 117500 | 0.000381357 | 0.00502 | 0.011277202 | 0.010584186 |
| pseudoscaff_1273 | 122500 | 0.000117718 | 0.004684 | 0.003904119 | 0.005265787 |
| pseudoscaff_1273 | 127500 | 0.000671104 | 0.012048 | 0.006204034 | 0.004534699 |
| pseudoscaff_1273 | 132500 | 0.001895092 | 0.011088 | 0.013816264 | 0.010289577 |
| pseudoscaff_1273 | 142500 | 0.001436503 | 0.012966 | 0.008604551 | 0.008502234 |
| pseudoscaff_1845 | 2500 | 0.001160744 | 0.000376 | 0.001019276 | 0.001361949 |
| pseudoscaff_1845 | 12500 | 0.001273043 | 0.001492 | 0.002398794 | 0.002302232 |
| pseudoscaff_1845 | 17500 | 0.000202165 | 0.000785 | 0.001511521 | 0.001435534 |
| pseudoscaff_1845 | 22500 | 0.000160891 | 0.00075 | 0.002130062 | 0.002027494 |
| pseudoscaff_2160 | 2500 | 0.001099303 | 0.007906 | 0.008073125 | 0.006965946 |
| pseudoscaff_2160 | 17500 | 0.000301073 | 0.007078 | 0.005465204 | 0.005700937 |
| pseudoscaff_2893 | 7500 | 0.001037357 | 0.013027 | 0.013312301 | 0.012387549 |
| pseudoscaff_2893 | 12500 | 0.000739128 | 0.017166 | 0.018135585 | 0.019551116 |
| pseudoscaff_2893 | 32500 | 0.001345088 | 0.006756 | 0.010047495 | 0.008819635 |
| pseudoscaff_1800 | 47500 | 0.001293134 | 0.022299 | 0.04355444 | 0.042697283 |
| pseudoscaff_749 | 42500 | 0.001863326 | 0.001906 | 0.002833357 | 0.002875779 |

| | | | | | |
|---|---|---|---|---|---|
| pseudoscaff_2515 | 2500 | 0 | 0.02095 | 0.013262276 | 0.011840164 |
| pseudoscaff_2515 | 7500 | 0.000335171 | 0.040439 | 0.016960211 | 0.01234967 |
| pseudoscaff_2515 | 72500 | 0.001692416 | 0.003945 | 0.007154987 | 0.007201375 |
| pseudoscaff_2515 | 77500 | 0.000913446 | 0.00406 | 0.003034415 | 0.00342808 |
| pseudoscaff_2515 | 87500 | 0.00037702 | 0.018155 | 0.016957719 | 0.017286635 |
| pseudoscaff_2515 | 97500 | 0 | 0.001923 | 0.001493863 | 0.001494131 |
| pseudoscaff_2515 | 102500 | 0.000070923 | 0.003022 | 0.00242234 | 0.002560559 |
| pseudoscaff_2515 | 107500 | 0 | 0.00258 | 0.002756038 | 0.002392288 |
| pseudoscaff_2515 | 112500 | 0.001027308 | 0.002827 | 0.009159954 | 0.006896353 |
| pseudoscaff_2515 | 117500 | 0.001093606 | 0.008808 | 0.009983285 | 0.010650512 |
| pseudoscaff_743 | 2500 | 0.000596743 | 0.020194 | 0.015439137 | 0.015817431 |
| pseudoscaff_743 | 7500 | 0.000873182 | 0.012217 | 0.014351521 | 0.011198045 |
| pseudoscaff_245 | 32500 | 0 | 0.02723 | 0.025808071 | 0.028211257 |
| pseudoscaff_2364 | 27500 | 0.001692428 | 0.02092 | 0.024186442 | 0.022307634 |
| pseudoscaff_2364 | 37500 | 0.000087478 | 0.02044 | 0.020727316 | 0.020304852 |
| pseudoscaff_2364 | 42500 | 0.00065056 | 0.02978 | 0.031589795 | 0.032826069 |
| pseudoscaff_2364 | 57500 | 0.000467668 | 0.024128 | 0.021603118 | 0.021231496 |
| pseudoscaff_2364 | 62500 | 0.001238117 | 0.036584 | 0.034273604 | 0.035076419 |
| pseudoscaff_2364 | 67500 | 0.001797297 | 0.030499 | 0.031961888 | 0.032571867 |
| pseudoscaff_2364 | 72500 | 0.001085461 | 0.031812 | 0.03228718 | 0.033372134 |
| pseudoscaff_2364 | 82500 | 0.000105212 | 0.027657 | 0.02760004 | 0.025815592 |
| pseudoscaff_2364 | 97500 | 0 | 0.020996 | 0.023787123 | 0.025516984 |
| pseudoscaff_2364 | 117500 | 0.000355876 | 0.035254 | 0.033953283 | 0.0355966 |
| pseudoscaff_2364 | 122500 | 0.000205193 | 0.016916 | 0.026105805 | 0.025510242 |
| pseudoscaff_2364 | 127500 | 0.000504789 | 0.024278 | 0.031455788 | 0.032522113 |
| pseudoscaff_2364 | 142500 | 0 | 0.018106 | 0.025170063 | 0.024545633 |
| pseudoscaff_2364 | 147500 | 0 | 0.020849 | 0.025518587 | 0.026305142 |
| pseudoscaff_2364 | 152500 | 0.000384892 | 0.020367 | 0.025679575 | 0.024905116 |
| pseudoscaff_2364 | 157500 | 0.000415401 | 0.011665 | 0.009507364 | 0.009602701 |
| pseudoscaff_2364 | 162500 | 0.000582644 | 0.012243 | 0.010511651 | 0.009937373 |
| pseudoscaff_2364 | 167500 | 0.001391752 | 0.018334 | 0.019672525 | 0.017442406 |
| pseudoscaff_2364 | 187500 | 0 | 0.023185 | 0.032105418 | 0.03436232 |
| pseudoscaff_2364 | 192500 | 0.000081373 | 0.019564 | 0.021683008 | 0.021629041 |
| pseudoscaff_2364 | 197500 | 0.000266992 | 0.031693 | 0.031062497 | 0.03480711 |
| pseudoscaff_2364 | 202500 | 0.000905991 | 0.022887 | 0.026415487 | 0.024252701 |
| pseudoscaff_2364 | 207500 | 0.000639863 | 0.022347 | 0.020292222 | 0.02088903 |
| pseudoscaff_2364 | 212500 | 0.000301555 | 0.031086 | 0.02861965 | 0.029507736 |
| pseudoscaff_2364 | 227500 | 0.001930334 | 0.014913 | 0.01988294 | 0.020650176 |
| pseudoscaff_2364 | 247500 | 0 | 0.024225 | 0.030239489 | 0.029561415 |
| pseudoscaff_2364 | 257500 | 0.000304665 | 0.014514 | 0.022463612 | 0.023418722 |
| pseudoscaff_2364 | 262500 | 0 | 0.010975 | 0.027985703 | 0.026727662 |
| pseudoscaff_2364 | 267500 | 0.000759362 | 0.028162 | 0.028024923 | 0.026650743 |
| pseudoscaff_83 | 117500 | 0.00065865 | 0.02581 | 0.030204681 | 0.029716542 |
| pseudoscaff_83 | 127500 | 0.001601149 | 0.027537 | 0.034134652 | 0.03708536 |
| pseudoscaff_83 | 137500 | 0.00076912 | 0.025939 | 0.038472282 | 0.038955919 |
| pseudoscaff_83 | 152500 | 0.000946636 | 0.017073 | 0.029576704 | 0.024713791 |
| pseudoscaff_2815 | 12500 | 0.000341506 | 0.029196 | 0.028887698 | 0.027475496 |
| pseudoscaff_2782 | 17500 | 0.001825876 | 0.011697 | 0.010323965 | 0.00997108 |
| pseudoscaff_3998 | 32500 | 0.000620693 | 0.00163 | 0.009093404 | 0.008336531 |
| pseudoscaff_3998 | 37500 | 0.001686132 | 0.004678 | 0.006186365 | 0.006628004 |

| | | | | | |
|---|---|---|---|---|---|
| pseudoscaff_3998 | 47500 | 0 | 0.001632 | 0.001688435 | 0.0017899 |
| pseudoscaff_3998 | 52500 | 0.0018448 | 0.003205 | 0.003071729 | 0.002267669 |
| pseudoscaff_1519 | 2500 | 0.000606602 | 0.004279 | 0.005018742 | 0.005006101 |
| pseudoscaff_3462 | 67500 | 0.001121544 | 0.007022 | 0.00667203 | 0.006597681 |
| pseudoscaff_698 | 2500 | 0.001992546 | 0.020271 | 0.022161557 | 0.020042114 |
| pseudoscaff_698 | 12500 | 0.001399294 | 0.021929 | 0.029349639 | 0.031297298 |
| pseudoscaff_2676 | 22500 | 0.001496444 | 0.030901 | 0.034003967 | 0.038280361 |
| pseudoscaff_1753 | 42500 | 0 | 0.021107 | 0.029000052 | 0.027661966 |
| pseudoscaff_1753 | 47500 | 0.000743473 | 0.015528 | 0.031161747 | 0.029466646 |
| pseudoscaff_2808 | 7500 | 0 | 0.013213 | 0.010333051 | 0.009877251 |
| pseudoscaff_3444 | 2500 | 0.000929386 | 0.004562 | 0.009064341 | 0.009004228 |
| pseudoscaff_3444 | 7500 | 0 | 0.002676 | 0.006045255 | 0.005220873 |
| pseudoscaff_3444 | 12500 | 0 | 0.005356 | 0.003061896 | 0.004357832 |
| pseudoscaff_544 | 2500 | 0.000619001 | 0.014606 | 0.035081128 | 0.030187103 |
| pseudoscaff_544 | 12500 | 0.000295495 | 0.019743 | 0.02801735 | 0.024392157 |
| pseudoscaff_544 | 17500 | 0 | 0.013276 | 0.015880726 | 0.012723226 |
| pseudoscaff_544 | 27500 | 0.00190158 | 0.013173 | 0.027766032 | 0.029094867 |
| pseudoscaff_544 | 37500 | 0.001042685 | 0.022813 | 0.02290611 | 0.018373228 |
| pseudoscaff_544 | 52500 | 0 | 0.016484 | 0.014477071 | 0.015880959 |
| pseudoscaff_544 | 57500 | 0.000761075 | 0.022908 | 0.014058447 | 0.013406587 |
| pseudoscaff_544 | 62500 | 0 | 0.02124 | 0.009047459 | 0.009250729 |
| pseudoscaff_544 | 67500 | 0.000585623 | 0.009132 | 0.015958196 | 0.018638137 |
| pseudoscaff_544 | 77500 | 0.00015934 | 0.014263 | 0.030697386 | 0.029232368 |
| pseudoscaff_544 | 82500 | 0.000693113 | 0.010064 | 0.023058311 | 0.021413388 |
| pseudoscaff_544 | 87500 | 0.000879243 | 0.005474 | 0.005845954 | 0.005538863 |
| pseudoscaff_544 | 92500 | 0.001385013 | 0.012712 | 0.016195235 | 0.015043952 |
| pseudoscaff_744 | 7500 | 0.000539374 | 0.015233 | 0.012744453 | 0.012030425 |
| pseudoscaff_2972 | 2500 | 0 | 0.016913 | 0.017936235 | 0.017015393 |
| pseudoscaff_1081 | 22500 | 0.001674482 | 0.01996 | 0.032086598 | 0.031061102 |
| pseudoscaff_2621 | 2500 | 0.000271707 | 0.002849 | 0.003997925 | 0.003207524 |
| pseudoscaff_2621 | 7500 | 0 | 0.004681 | 0.010346711 | 0.009831119 |
| pseudoscaff_2621 | 17500 | 0 | 0.003741 | 0.018017617 | 0.017923868 |
| pseudoscaff_2621 | 22500 | 0.000533557 | 0.001918 | 0.003057024 | 0.002660888 |
| pseudoscaff_2621 | 27500 | 0.000833473 | 0.0043 | 0.002366326 | 0.002812247 |
| pseudoscaff_2621 | 32500 | 0 | 0.001489 | 0.002208548 | 0.001681969 |
| pseudoscaff_2621 | 42500 | 0 | 0.027382 | 0.007937545 | 0.00715882 |
| pseudoscaff_2621 | 52500 | 0 | 0.009118 | 0.001712432 | 0.001773327 |
| pseudoscaff_2621 | 57500 | 0.000150788 | 0.025959 | 0.019844455 | 0.014144371 |
| pseudoscaff_4 | 32500 | 0 | 0.02031 | 0.03820503 | 0.037689486 |
| pseudoscaff_4 | 37500 | 0 | 0.016607 | 0.021792792 | 0.022052488 |
| pseudoscaff_4 | 47500 | 0.001668614 | 0.024742 | 0.043827777 | 0.042700122 |
| pseudoscaff_2985 | 17500 | 0.001137095 | 0.036593 | 0.043321182 | 0.044189826 |
| pseudoscaff_2953 | 7500 | 0.00012368 | 0.008134 | 0.015013334 | 0.020460485 |
| pseudoscaff_2953 | 22500 | 0 | 0.024775 | 0.010476898 | 0.016021649 |
| pseudoscaff_1488 | 2500 | 0.001045515 | 0.000656 | 0.000855114 | 0.001026741 |
| pseudoscaff_1488 | 7500 | 0.001328327 | 0.002771 | 0.002576665 | 0.002485382 |
| pseudoscaff_3662 | 147500 | 0.000172688 | 0.021755 | 0.042119113 | 0.034537513 |
| pseudoscaff_3662 | 172500 | 0 | 0.0324 | 0.031955405 | 0.03402967 |
| pseudoscaff_3662 | 182500 | 0.000805927 | 0.024028 | 0.038447749 | 0.041704434 |
| pseudoscaff_3662 | 197500 | 0.000263206 | 0.030408 | 0.03695811 | 0.041602923 |

| | | | | | |
|---|---|---|---|---|---|
| pseudoscaff_3662 | 202500 | 0.000302384 | 0.02682 | 0.037345893 | 0.039905771 |
| pseudoscaff_3662 | 252500 | 0.000321208 | 0.017691 | 0.022292703 | 0.020305085 |
| pseudoscaff_3662 | 257500 | 0 | 0.015701 | 0.01833245 | 0.018268736 |
| pseudoscaff_3662 | 267500 | 0.001019082 | 0.011143 | 0.016399622 | 0.017219543 |
| pseudoscaff_3662 | 292500 | 0.001678042 | 0.013051 | 0.020051929 | 0.019885111 |
| pseudoscaff_3662 | 297500 | 0.00073372 | 0.01863 | 0.025391763 | 0.027682628 |
| pseudoscaff_3662 | 302500 | 0.001623274 | 0.021069 | 0.030078205 | 0.037057394 |
| pseudoscaff_3662 | 307500 | 0.000824365 | 0.041429 | 0.034429359 | 0.036695549 |
| pseudoscaff_4009 | 2500 | 0.000369481 | 0.00229 | 0.001499367 | 0.001607495 |
| pseudoscaff_4009 | 7500 | 0.001963049 | 0.001616 | 0.002022005 | 0.001785776 |
| pseudoscaff_4009 | 22500 | 0.000770925 | 0.001628 | 0.001976389 | 0.001305891 |
| pseudoscaff_4009 | 32500 | 0.001213343 | 0.000712 | 0.00175818 | 0.001812075 |
| pseudoscaff_4009 | 37500 | 0.001064521 | 0.002722 | 0.002207903 | 0.001922666 |
| pseudoscaff_80 | 77500 | 0.000855524 | 0.038338 | 0.037649449 | 0.03612848 |
| pseudoscaff_959 | 2500 | 0.000503606 | 0.002954 | 0.003213601 | 0.003479943 |
| pseudoscaff_959 | 12500 | 0.001213266 | 0.003386 | 0.004602973 | 0.003836171 |
| pseudoscaff_959 | 22500 | 0 | 0.001803 | 0.001816782 | 0.001438067 |
| pseudoscaff_959 | 27500 | 0 | 0.008769 | 0.01563401 | 0.015287521 |
| pseudoscaff_959 | 47500 | 0.000633636 | 0.005291 | 0.005472715 | 0.005237737 |
| pseudoscaff_1645 | 27500 | 0 | 0.035889 | 0.01141303 | 0.016970181 |
| pseudoscaff_2019 | 2500 | 0.000488744 | 0.000884 | 0.000854778 | 0.001100218 |
| pseudoscaff_1652 | 2500 | 0 | 0.02089 | 0.022390717 | 0.02291394 |
| pseudoscaff_1652 | 7500 | 0 | 0.016187 | 0.018171911 | 0.018161163 |
| pseudoscaff_1652 | 12500 | 0.001342 | 0.028745 | 0.032538809 | 0.034405072 |
| pseudoscaff_1652 | 22500 | 0.000298735 | 0.028886 | 0.028154608 | 0.028695694 |
| pseudoscaff_1652 | 27500 | 0 | 0.011932 | 0.019056747 | 0.018716266 |
| pseudoscaff_1652 | 32500 | 0.000066911 | 0.01685 | 0.02055196 | 0.021613182 |
| pseudoscaff_1652 | 67500 | 0 | 0.012362 | 0.013571335 | 0.013611655 |
| pseudoscaff_1439 | 7500 | 0.000419543 | 0.019587 | 0.020911997 | 0.022771057 |
| pseudoscaff_1439 | 17500 | 0.000728834 | 0.011598 | 0.026280613 | 0.028160712 |
| pseudoscaff_1439 | 32500 | 0 | 0.021928 | 0.027450731 | 0.0297025 |
| pseudoscaff_1439 | 37500 | 0 | 0.021028 | 0.026340533 | 0.026795838 |
| pseudoscaff_1439 | 42500 | 0.000623359 | 0.02095 | 0.03129654 | 0.029674177 |
| pseudoscaff_3531 | 2500 | 0.001492092 | 0.026427 | 0.032788875 | 0.036878152 |
| pseudoscaff_3531 | 17500 | 0 | 0.051791 | 0.04305907 | 0.038597685 |
| pseudoscaff_3531 | 22500 | 0.001566075 | 0.045487 | 0.04717817 | 0.042661351 |
| pseudoscaff_3531 | 27500 | 0.001584253 | 0.038408 | 0.03239557 | 0.03693907 |
| pseudoscaff_3531 | 32500 | 0 | 0.025481 | 0.029603126 | 0.029188704 |
| pseudoscaff_3531 | 37500 | 0.000545617 | 0.0338 | 0.038536171 | 0.037161144 |
| pseudoscaff_3531 | 47500 | 0.000159793 | 0.02752 | 0.032845842 | 0.037718463 |
| pseudoscaff_3531 | 52500 | 0.000217229 | 0.021245 | 0.030761615 | 0.029843667 |
| pseudoscaff_3531 | 62500 | 0 | 0.025706 | 0.022875398 | 0.023820621 |
| pseudoscaff_3531 | 72500 | 0 | 0.031394 | 0.029781265 | 0.027331241 |
| pseudoscaff_806 | 82500 | 0.00195519 | 0.044252 | 0.04419536 | 0.045086969 |
| pseudoscaff_806 | 97500 | 0.001790242 | 0.046791 | 0.041035256 | 0.044449027 |
| pseudoscaff_3209 | 2500 | 0.000310967 | 0.000511 | 0.001100581 | 0.001839567 |
| pseudoscaff_3209 | 7500 | 0 | 0 | 0.000352117 | 0.000761406 |
| pseudoscaff_3209 | 12500 | 0 | 0.00013 | 0.000671656 | 0.000905038 |
| pseudoscaff_3209 | 17500 | 0.000146791 | 0.0003 | 0.000597819 | 0.000942242 |
| pseudoscaff_3209 | 22500 | 0.000296535 | 0.002843 | 0.004774069 | 0.006411143 |

| | | | | | |
|---|---|---|---|---|---|
| pseudoscaff_3209 | 27500 | 0.000776408 | 0.002445 | 0.001846592 | 0.003356288 |
| pseudoscaff_3209 | 47500 | 0.001853783 | 0.003713 | 0.004505604 | 0.00379765 |
| pseudoscaff_3209 | 57500 | 0.000110228 | 0.00498 | 0.004398306 | 0.004369921 |
| pseudoscaff_3209 | 67500 | 0 | 0.000546 | 0.002441554 | 0.001898929 |
| pseudoscaff_3209 | 72500 | 0 | 0.000749 | 0.001640511 | 0.001661453 |
| pseudoscaff_3209 | 77500 | 0.001529106 | 0.006557 | 0.010262291 | 0.008627534 |
| pseudoscaff_3209 | 107500 | 0.0009423 | 0.016041 | 0.015448455 | 0.01500887 |
| pseudoscaff_1333 | 137500 | 0.001253161 | 0.014012 | 0.012254531 | 0.009391047 |
| pseudoscaff_1333 | 167500 | 0.001064006 | 0.015562 | 0.020705944 | 0.021492863 |
| pseudoscaff_1333 | 187500 | 0.000204605 | 0.002187 | 0.004269124 | 0.003768164 |
| pseudoscaff_1333 | 192500 | 0.000202265 | 0.000601 | 0.002755962 | 0.002829156 |
| pseudoscaff_1333 | 197500 | 0.001004641 | 0.002928 | 0.00349708 | 0.002690552 |
| pseudoscaff_1333 | 202500 | 0.001963657 | 0.003213 | 0.00247739 | 0.002814044 |
| pseudoscaff_1333 | 212500 | 0 | 0.003092 | 0.002616793 | 0.002624763 |
| pseudoscaff_1333 | 217500 | 0 | 0.004209 | 0.003841921 | 0.003578264 |
| pseudoscaff_1333 | 222500 | 0.001520994 | 0.005209 | 0.002720037 | 0.002631347 |
| pseudoscaff_1333 | 227500 | 0.000153751 | 0.002394 | 0.001956775 | 0.002033138 |
| pseudoscaff_1333 | 232500 | 0.001017281 | 0.001747 | 0.00231213 | 0.001521798 |
| pseudoscaff_1333 | 242500 | 0.000766656 | 0.000827 | 0.002457006 | 0.003033516 |
| pseudoscaff_1333 | 247500 | 0.000964402 | 0.005555 | 0.00163545 | 0.003737769 |
| pseudoscaff_1333 | 252500 | 0.000646297 | 0.002702 | 0.002116316 | 0.002132862 |
| pseudoscaff_1333 | 257500 | 0 | 0.001232 | 0.000836789 | 0.001167288 |
| pseudoscaff_425 | 42500 | 0.001589708 | 0.002655 | 0.00154798 | 0.001640781 |
| pseudoscaff_425 | 47500 | 0.001217089 | 0.00368 | 0.001214088 | 0.000987128 |
| pseudoscaff_425 | 62500 | 0.000992833 | 0.003177 | 0.00125035 | 0.001320512 |
| pseudoscaff_425 | 67500 | 0.000756616 | 0.001794 | 0.000651521 | 0.001089822 |
| pseudoscaff_3165 | 27500 | 0.000106641 | 0.003337 | 0.004292427 | 0.004423196 |
| pseudoscaff_3165 | 32500 | 0.000621441 | 0.00258 | 0.002358349 | 0.002943967 |
| pseudoscaff_3165 | 37500 | 0 | 0.003201 | 0.002043164 | 0.002270792 |
| pseudoscaff_3165 | 47500 | 0.000148268 | 0.003567 | 0.004700148 | 0.004793029 |
| pseudoscaff_3165 | 57500 | 0.000698179 | 0.009031 | 0.007663894 | 0.007232627 |
| pseudoscaff_3165 | 62500 | 0.000707223 | 0.01071 | 0.0097655 | 0.008393132 |
| pseudoscaff_3165 | 67500 | 0.001809404 | 0.006921 | 0.006523034 | 0.005881844 |
| pseudoscaff_3165 | 72500 | 0.000627401 | 0.009078 | 0.008388847 | 0.007947038 |
| pseudoscaff_3165 | 87500 | 0.001787624 | 0.008205 | 0.011780591 | 0.011324181 |
| pseudoscaff_3165 | 92500 | 0 | 0.006698 | 0.00738899 | 0.007392734 |
| pseudoscaff_3165 | 97500 | 0.000149846 | 0.005953 | 0.006064992 | 0.006696268 |
| pseudoscaff_3165 | 172500 | 0.001374451 | 0.005054 | 0.007967249 | 0.00868109 |
| pseudoscaff_3165 | 197500 | 0.00070412 | 0.006682 | 0.005166999 | 0.00532125 |
| pseudoscaff_3165 | 202500 | 0.00089753 | 0.003921 | 0.00686415 | 0.006861631 |
| pseudoscaff_3165 | 217500 | 0 | 0.002413 | 0.007722142 | 0.006980568 |
| pseudoscaff_3165 | 242500 | 0.001473788 | 0.003657 | 0.00689692 | 0.006493517 |
| pseudoscaff_3469 | 2500 | 0.000454125 | 0.021925 | 0.026256567 | 0.027734824 |
| pseudoscaff_3469 | 42500 | 0.001667727 | 0.015967 | 0.018879645 | 0.016992764 |
| pseudoscaff_3469 | 57500 | 0.000466534 | 0.03408 | 0.033462007 | 0.035896145 |
| pseudoscaff_3469 | 67500 | 0.001095644 | 0.023881 | 0.025233011 | 0.025876759 |
| pseudoscaff_3469 | 72500 | 0 | 0.009475 | 0.013330184 | 0.014021667 |
| pseudoscaff_3469 | 77500 | 0.001872966 | 0.017089 | 0.017846776 | 0.017083737 |
| pseudoscaff_3469 | 82500 | 0 | 0.016591 | 0.019402226 | 0.021083485 |
| pseudoscaff_3469 | 87500 | 0 | 0.021108 | 0.022397053 | 0.022030299 |

| | | | | | |
|---|---|---|---|---|---|
| pseudoscaff_3469 | 147500 | 0.000182 | 0.041721 | 0.035705099 | 0.036244987 |
| pseudoscaff_3469 | 292500 | 0.001819477 | 0.031615 | 0.033203936 | 0.036844851 |
| pseudoscaff_3469 | 322500 | 0 | 0.032653 | 0.031861466 | 0.031220989 |
| pseudoscaff_3469 | 327500 | 0 | 0.022231 | 0.027160259 | 0.025791486 |
| pseudoscaff_3469 | 332500 | 0 | 0.022685 | 0.017299981 | 0.017364706 |
| pseudoscaff_3469 | 352500 | 0.000367789 | 0.030842 | 0.030255742 | 0.031112844 |
| pseudoscaff_3469 | 357500 | 0.000259745 | 0.012745 | 0.014412951 | 0.015227101 |
| pseudoscaff_3469 | 362500 | 0.000228466 | 0.019757 | 0.02482676 | 0.024977066 |
| pseudoscaff_3469 | 387500 | 0.000152519 | 0.03621 | 0.030531971 | 0.030576397 |
| pseudoscaff_3469 | 392500 | 0 | 0.03582 | 0.033149589 | 0.03351004 |
| pseudoscaff_3469 | 412500 | 0 | 0.018198 | 0.017185414 | 0.017666783 |
| pseudoscaff_3469 | 452500 | 0.001118471 | 0.007935 | 0.007562192 | 0.009778377 |
| pseudoscaff_3469 | 457500 | 0 | 0.015365 | 0.015596972 | 0.016497126 |
| pseudoscaff_3469 | 472500 | 0.000229236 | 0.028679 | 0.034498127 | 0.036511295 |
| pseudoscaff_3469 | 477500 | 0.000077296 | 0.009636 | 0.013639434 | 0.01376717 |
| pseudoscaff_3469 | 482500 | 0 | 0.029952 | 0.029616777 | 0.031680621 |
| pseudoscaff_3469 | 487500 | 0.001014116 | 0.038489 | 0.042119247 | 0.0424671 |
| pseudoscaff_3469 | 492500 | 0 | 0.035556 | 0.03389668 | 0.034464729 |
| pseudoscaff_3469 | 497500 | 0.000253515 | 0.03112 | 0.030271654 | 0.029459116 |
| pseudoscaff_3469 | 502500 | 0.000426611 | 0.037161 | 0.032599529 | 0.032476073 |
| pseudoscaff_3469 | 507500 | 0.000862921 | 0.043825 | 0.041619106 | 0.044747101 |
| pseudoscaff_3469 | 512500 | 0.000649717 | 0.042072 | 0.041250401 | 0.039781671 |
| pseudoscaff_3469 | 532500 | 0 | 0.024307 | 0.022711978 | 0.023358365 |
| pseudoscaff_3469 | 537500 | 0.00114152 | 0.028537 | 0.026923299 | 0.031304659 |
| pseudoscaff_3469 | 667500 | 0.000475195 | 0.020442 | 0.023744754 | 0.024234674 |
| pseudoscaff_3469 | 692500 | 0.000532068 | 0.029702 | 0.029211456 | 0.034894442 |
| pseudoscaff_3469 | 697500 | 0.000100734 | 0.013223 | 0.016298978 | 0.015668257 |
| pseudoscaff_3469 | 702500 | 0 | 0.008966 | 0.00791742 | 0.007456113 |
| pseudoscaff_3469 | 707500 | 0.000502193 | 0.011819 | 0.011545544 | 0.011994702 |
| pseudoscaff_2234 | 7500 | 0.000979421 | 0.008217 | 0.00770982 | 0.00812348 |
| pseudoscaff_2234 | 17500 | 0.000346519 | 0.000793 | 0.003588401 | 0.003476017 |
| pseudoscaff_3555 | 2500 | 0.001638748 | 0.016178 | 0.012589578 | 0.013468411 |
| pseudoscaff_3555 | 7500 | 0.000492341 | 0.0095 | 0.008548616 | 0.008441473 |
| pseudoscaff_3555 | 12500 | 0.00051892 | 0.036432 | 0.040029846 | 0.041966659 |
| pseudoscaff_3555 | 17500 | 0 | 0.02396 | 0.023458331 | 0.024322764 |
| pseudoscaff_3555 | 22500 | 0.001918336 | 0.020398 | 0.020508037 | 0.01888519 |
| pseudoscaff_3555 | 27500 | 0 | 0.037271 | 0.038752735 | 0.034720842 |
| pseudoscaff_3555 | 37500 | 0.000359079 | 0.025934 | 0.029287663 | 0.025535854 |
| pseudoscaff_3555 | 42500 | 0.001316775 | 0.032932 | 0.031918808 | 0.029067249 |
| pseudoscaff_3555 | 47500 | 0.000061481 | 0.032691 | 0.026249477 | 0.028609662 |
| pseudoscaff_3555 | 52500 | 0 | 0.035597 | 0.039598628 | 0.038656841 |
| pseudoscaff_3555 | 62500 | 0.000135521 | 0.03017 | 0.035429046 | 0.036701058 |
| pseudoscaff_3555 | 97500 | 0.000671366 | 0.025701 | 0.031101988 | 0.03085844 |
| pseudoscaff_3555 | 102500 | 0.000190867 | 0.038842 | 0.041070959 | 0.037106587 |
| pseudoscaff_3555 | 117500 | 0 | 0.05245 | 0.042344028 | 0.041103538 |
| pseudoscaff_3555 | 122500 | 0.00170836 | 0.029757 | 0.035086778 | 0.032071253 |
| pseudoscaff_3555 | 132500 | 0.000198853 | 0.04066 | 0.037757192 | 0.036348595 |
| pseudoscaff_3555 | 142500 | 0 | 0.029506 | 0.033866835 | 0.035485227 |
| pseudoscaff_3555 | 147500 | 0 | 0.030401 | 0.034417494 | 0.033667091 |
| pseudoscaff_3555 | 152500 | 0 | 0.037846 | 0.039089614 | 0.03554453 |

| | | | | | |
|---|---|---|---|---|---|
| pseudoscaff_3555 | 157500 | 0.000860106 | 0.011422 | 0.019519518 | 0.017488519 |
| pseudoscaff_3893 | 27500 | 0.000777428 | 0.024582 | 0.026282477 | 0.027074771 |
| pseudoscaff_3893 | 42500 | 0 | 0.006336 | 0.007042189 | 0.006916087 |
| pseudoscaff_3893 | 57500 | 0 | 0.023336 | 0.020561718 | 0.020653341 |
| pseudoscaff_3893 | 92500 | 0.000809756 | 0.03947 | 0.023959462 | 0.025565393 |
| pseudoscaff_3893 | 107500 | 0 | 0.024823 | 0.017374819 | 0.017483257 |
| pseudoscaff_486 | 2500 | 0.000140952 | 0.033841 | 0.029981796 | 0.029382125 |
| pseudoscaff_2917 | 182500 | 0.000429906 | 0.033811 | 0.05042364 | 0.047863516 |
| pseudoscaff_4215 | 12500 | 0.000749446 | 0.046396 | 0.048444799 | 0.047045163 |
| pseudoscaff_4215 | 37500 | 0.000520374 | 0.033208 | 0.042955538 | 0.04316042 |
| pseudoscaff_4215 | 42500 | 0.000115844 | 0.02516 | 0.03403808 | 0.037474955 |
| pseudoscaff_2638 | 7500 | 0 | 0.025057 | 0.024565861 | 0.02642359 |
| pseudoscaff_4204 | 2500 | 0.000401861 | 0.008916 | 0.007719337 | 0.007449097 |
| pseudoscaff_4204 | 12500 | 0 | 0.009499 | 0.007384005 | 0.007422285 |
| pseudoscaff_4204 | 17500 | 0 | 0.005241 | 0.005183917 | 0.005203754 |
| pseudoscaff_4204 | 27500 | 0 | 0.00488 | 0.004080886 | 0.004530527 |
| pseudoscaff_4204 | 32500 | 0 | 0.004463 | 0.003433329 | 0.003223222 |
| pseudoscaff_4204 | 42500 | 0.000470301 | 0.003259 | 0.003150734 | 0.003273936 |
| pseudoscaff_4204 | 47500 | 0 | 0.003356 | 0.003156639 | 0.002071397 |
| pseudoscaff_4204 | 52500 | 0 | 0.008597 | 0.006508056 | 0.004972427 |
| pseudoscaff_3970 | 2500 | 0.000236047 | 0.019509 | 0.023890376 | 0.021876293 |
| pseudoscaff_3970 | 12500 | 0.001015374 | 0.020146 | 0.020429972 | 0.021163918 |
| pseudoscaff_3970 | 17500 | 0.00095862 | 0.021723 | 0.025251051 | 0.025395321 |
| pseudoscaff_3970 | 27500 | 0.000697641 | 0.014189 | 0.016320023 | 0.017215212 |
| pseudoscaff_3970 | 32500 | 0 | 0.020553 | 0.015008588 | 0.016979749 |
| pseudoscaff_3970 | 42500 | 0.000503074 | 0.027851 | 0.033959893 | 0.033075899 |
| pseudoscaff_1419 | 12500 | 0 | 0.029357 | 0.032832689 | 0.033781862 |
| pseudoscaff_1419 | 17500 | 0 | 0.023412 | 0.02825933 | 0.026068495 |
| pseudoscaff_1419 | 22500 | 0.000072704 | 0.031155 | 0.040743852 | 0.044140912 |
| pseudoscaff_2648 | 97500 | 0.00028921 | 0.017183 | 0.019283027 | 0.017535725 |
| pseudoscaff_2648 | 347500 | 0 | 0.010209 | 0.014160408 | 0.013856687 |
| pseudoscaff_2648 | 352500 | 0.000085982 | 0.021631 | 0.026744601 | 0.026528821 |
| pseudoscaff_2648 | 357500 | 0.000574308 | 0.026174 | 0.034109461 | 0.036616753 |
| pseudoscaff_2648 | 362500 | 0.000836415 | 0.047439 | 0.04464444 | 0.047230343 |
| pseudoscaff_3198 | 27500 | 0.000459056 | 0.002484 | 0.004182044 | 0.004193617 |
| pseudoscaff_3198 | 52500 | 0.000805885 | 0.015797 | 0.015826211 | 0.014546316 |
| pseudoscaff_2202 | 27500 | 0.00133515 | 0.009421 | 0.009080401 | 0.00868094 |
| pseudoscaff_2202 | 327500 | 0.001942451 | 0.0084 | 0.008410487 | 0.008095288 |
| pseudoscaff_3416 | 7500 | 0.000449099 | 0.008324 | 0.015033146 | 0.012938173 |
| pseudoscaff_3416 | 12500 | 0.000297711 | 0.007821 | 0.004796262 | 0.006177988 |
| pseudoscaff_3416 | 17500 | 0.00133817 | 0.006053 | 0.007638203 | 0.007354879 |
| pseudoscaff_3416 | 22500 | 0 | 0.019881 | 0.026027002 | 0.020722498 |
| pseudoscaff_3416 | 27500 | 0.001124834 | 0.007876 | 0.01679517 | 0.017706187 |
| pseudoscaff_3416 | 52500 | 0 | 0.006456 | 0.015177553 | 0.013896269 |
| pseudoscaff_3416 | 57500 | 0.00088767 | 0.010505 | 0.007472537 | 0.007399145 |
| pseudoscaff_3416 | 72500 | 0.000422535 | 0.025303 | 0.028711291 | 0.026385042 |
| pseudoscaff_3416 | 77500 | 0 | 0.016719 | 0.018205785 | 0.018879273 |
| pseudoscaff_1586 | 7500 | 0.001193173 | 0.006661 | 0.004039371 | 0.003135697 |
| pseudoscaff_1586 | 12500 | 0.00083745 | 0.006166 | 0.00315155 | 0.003075408 |
| pseudoscaff_1240 | 2500 | 0.000863892 | 0.029183 | 0.027564836 | 0.023507182 |

| | | | | | |
|---|---|---|---|---|---|
| pseudoscaff_1240 | 27500 | 0.000258054 | 0.015169 | 0.0153117 | 0.014509238 |
| pseudoscaff_729 | 127500 | 0.000170431 | 0.016085 | 0.002110588 | 0.002314328 |
| pseudoscaff_729 | 157500 | 0 | 0.001231 | 0.00217056 | 0.00198419 |
| pseudoscaff_729 | 162500 | 0.000481613 | 0.013959 | 0.003466905 | 0.003924369 |
| pseudoscaff_729 | 182500 | 0 | 0.004495 | 0.002860886 | 0.003903879 |
| pseudoscaff_729 | 187500 | 0 | 0.003749 | 0.002870477 | 0.002079565 |
| pseudoscaff_729 | 197500 | 0.000335888 | 0.000443 | 0.002747858 | 0.00596937 |
| pseudoscaff_729 | 227500 | 0.001648622 | 0.005571 | 0.004108169 | 0.005303556 |
| pseudoscaff_729 | 247500 | 0.000856176 | 0.002685 | 0.002106972 | 0.002055037 |
| pseudoscaff_729 | 257500 | 0.000256022 | 0.003662 | 0.004621783 | 0.004392727 |
| pseudoscaff_729 | 262500 | 0.000925405 | 0.003123 | 0.003994158 | 0.004960398 |
| pseudoscaff_729 | 267500 | 0 | 0.006581 | 0.010775246 | 0.012825035 |
| pseudoscaff_729 | 272500 | 0 | 0.005182 | 0.004613119 | 0.005927337 |
| pseudoscaff_729 | 277500 | 0 | 0.003048 | 0.003060758 | 0.002981267 |
| pseudoscaff_729 | 287500 | 0.000935122 | 0.002226 | 0.004912968 | 0.005725451 |
| pseudoscaff_729 | 292500 | 0.00010369 | 0.002135 | 0.002133834 | 0.003050738 |
| pseudoscaff_729 | 297500 | 0.000209087 | 0.001393 | 0.002067618 | 0.002617806 |
| pseudoscaff_729 | 302500 | 0 | 0.000588 | 0.002675814 | 0.002684895 |
| pseudoscaff_729 | 382500 | 0.001141334 | 0.011366 | 0.026709406 | 0.028718401 |
| pseudoscaff_729 | 467500 | 0.000105743 | 0.009205 | 0.008544566 | 0.008597588 |
| pseudoscaff_729 | 472500 | 0.000135108 | 0.000799 | 0.001600994 | 0.001806982 |
| pseudoscaff_729 | 492500 | 0.001155874 | 0.007149 | 0.010569781 | 0.010497757 |
| pseudoscaff_729 | 502500 | 0.000572216 | 0.004708 | 0.003550542 | 0.003976465 |
| pseudoscaff_2153 | 2500 | 0.000338187 | 0.029612 | 0.034006862 | 0.034679317 |
| pseudoscaff_2153 | 32500 | 0.00025372 | 0.032969 | 0.034183308 | 0.038709635 |
| pseudoscaff_2153 | 37500 | 0.000340292 | 0.02081 | 0.027208847 | 0.028357798 |
| pseudoscaff_2153 | 57500 | 0.00187325 | 0.011504 | 0.015921856 | 0.01532556 |
| pseudoscaff_2153 | 62500 | 0.00059348 | 0.010971 | 0.011003436 | 0.010285938 |
| pseudoscaff_990 | 22500 | 0.001460946 | 0.022539 | 0.017973058 | 0.019499348 |
| pseudoscaff_2776 | 7500 | 0.00057024 | 0.009836 | 0.013623775 | 0.010351272 |
| pseudoscaff_2776 | 12500 | 0.000996463 | 0.010903 | 0.014152063 | 0.012554285 |
| pseudoscaff_1487 | 2500 | 0 | 0.014094 | 0.021851267 | 0.022788571 |
| pseudoscaff_1487 | 7500 | 0 | 0.009496 | 0.017280399 | 0.016242338 |
| pseudoscaff_1487 | 12500 | 0.000071524 | 0.009495 | 0.01136017 | 0.011154602 |
| pseudoscaff_1487 | 17500 | 0 | 0.009048 | 0.015373822 | 0.014643282 |
| pseudoscaff_1487 | 22500 | 0 | 0.006772 | 0.008644059 | 0.008629513 |
| pseudoscaff_1487 | 27500 | 0 | 0.022738 | 0.02700929 | 0.026204117 |
| pseudoscaff_1487 | 32500 | 0.000083072 | 0.014526 | 0.016391163 | 0.014570829 |
| pseudoscaff_1487 | 37500 | 0 | 0.018189 | 0.018213997 | 0.018008374 |
| pseudoscaff_1487 | 42500 | 0 | 0.015441 | 0.019597495 | 0.018939887 |
| pseudoscaff_1487 | 47500 | 0 | 0.011039 | 0.015433771 | 0.015410885 |
| pseudoscaff_1487 | 52500 | 0.000237085 | 0.009772 | 0.012811816 | 0.011980648 |
| pseudoscaff_1487 | 57500 | 0 | 0.008272 | 0.008423274 | 0.008608183 |
| pseudoscaff_1487 | 62500 | 0 | 0.010444 | 0.009934885 | 0.010293574 |
| pseudoscaff_1487 | 67500 | 0 | 0.006149 | 0.007512645 | 0.007309386 |
| pseudoscaff_1487 | 72500 | 0 | 0.012535 | 0.010149388 | 0.010252882 |
| pseudoscaff_1487 | 77500 | 0 | 0.014549 | 0.013183064 | 0.012852531 |
| pseudoscaff_1487 | 82500 | 0.000112793 | 0.010882 | 0.012171858 | 0.011190158 |
| pseudoscaff_1487 | 87500 | 0 | 0.015684 | 0.017057816 | 0.016958424 |
| pseudoscaff_1487 | 97500 | 0 | 0.009415 | 0.009873989 | 0.009824547 |

| | | | | | |
|---|---|---|---|---|---|
| pseudoscaff_1487 | 102500 | 0.00044478 | 0.006447 | 0.007777047 | 0.008034384 |
| pseudoscaff_1487 | 107500 | 0 | 0.007325 | 0.006152855 | 0.006162631 |
| pseudoscaff_1487 | 112500 | 0 | 0.009971 | 0.010083782 | 0.009328908 |
| pseudoscaff_1487 | 117500 | 0 | 0.015589 | 0.02356306 | 0.025307569 |
| pseudoscaff_1487 | 122500 | 0 | 0.004326 | 0.008591546 | 0.004519197 |
| pseudoscaff_1487 | 172500 | 0.000330256 | 0.029377 | 0.03495996 | 0.033849388 |
| pseudoscaff_1487 | 222500 | 0.001530195 | 0.028372 | 0.025800136 | 0.02447063 |
| pseudoscaff_1487 | 227500 | 0.000613586 | 0.040352 | 0.041892668 | 0.03848567 |
| pseudoscaff_1487 | 242500 | 0 | 0.016146 | 0.018532111 | 0.019763598 |
| pseudoscaff_1487 | 252500 | 0.001249517 | 0.023933 | 0.026952571 | 0.028956256 |
| pseudoscaff_2635 | 7500 | 0.001176466 | 0.038652 | 0.039231746 | 0.037263913 |
| pseudoscaff_2635 | 32500 | 0 | 0.023499 | 0.026322642 | 0.026640337 |
| pseudoscaff_2635 | 42500 | 0 | 0.034186 | 0.033396794 | 0.037280608 |
| pseudoscaff_2635 | 47500 | 0.000233654 | 0.036226 | 0.033107366 | 0.03000168 |
| pseudoscaff_2635 | 52500 | 0.001535369 | 0.042085 | 0.04334266 | 0.039820793 |
| pseudoscaff_2635 | 82500 | 0.00159788 | 0.043341 | 0.041821545 | 0.041526749 |
| pseudoscaff_2635 | 87500 | 0.000788294 | 0.047899 | 0.046343272 | 0.042994493 |
| pseudoscaff_2635 | 92500 | 0 | 0.043734 | 0.037547776 | 0.038064533 |
| pseudoscaff_2635 | 137500 | 0.000987211 | 0.040287 | 0.037455304 | 0.037611583 |
| pseudoscaff_2635 | 142500 | 0 | 0.032482 | 0.031503897 | 0.031762212 |
| pseudoscaff_2635 | 152500 | 0.001441341 | 0.04319 | 0.042431764 | 0.044193375 |
| pseudoscaff_2635 | 157500 | 0.001125023 | 0.031547 | 0.034109798 | 0.036683536 |
| pseudoscaff_2635 | 162500 | 0 | 0.028918 | 0.031013206 | 0.029721015 |
| pseudoscaff_2635 | 182500 | 0.001757465 | 0.03552 | 0.035468413 | 0.036213618 |
| pseudoscaff_2635 | 187500 | 0 | 0.045907 | 0.038703313 | 0.038415707 |
| pseudoscaff_2635 | 242500 | 0 | 0.019848 | 0.018059625 | 0.015898006 |
| pseudoscaff_2635 | 252500 | 0 | 0.017043 | 0.017103153 | 0.015970564 |
| pseudoscaff_2635 | 257500 | 0.000081192 | 0.040742 | 0.03882142 | 0.040735495 |
| pseudoscaff_2635 | 262500 | 0.000169226 | 0.043602 | 0.032935459 | 0.033209031 |
| pseudoscaff_2635 | 287500 | 0.000521724 | 0.041333 | 0.045855627 | 0.042600381 |
| pseudoscaff_2635 | 292500 | 0.000452453 | 0.036878 | 0.036065209 | 0.035139868 |
| pseudoscaff_2635 | 297500 | 0.00010231 | 0.02397 | 0.024073914 | 0.024350417 |
| pseudoscaff_2635 | 302500 | 0.000309233 | 0.027325 | 0.029294343 | 0.027362409 |
| pseudoscaff_2635 | 307500 | 0.000097265 | 0.031945 | 0.031751066 | 0.029530557 |
| pseudoscaff_2635 | 327500 | 0 | 0.030638 | 0.028397738 | 0.028486525 |
| pseudoscaff_2635 | 332500 | 0.001617891 | 0.035583 | 0.033159227 | 0.033464245 |
| pseudoscaff_2635 | 337500 | 0.001062463 | 0.040704 | 0.038140278 | 0.038756486 |
| pseudoscaff_2635 | 342500 | 0.000648958 | 0.038236 | 0.0404813 | 0.03998327 |
| pseudoscaff_2635 | 347500 | 0.000882036 | 0.042808 | 0.035916858 | 0.035441801 |
| pseudoscaff_2635 | 357500 | 0.001522446 | 0.021445 | 0.019225703 | 0.019261007 |
| pseudoscaff_2635 | 367500 | 0.000916487 | 0.015177 | 0.013089997 | 0.011868015 |
| pseudoscaff_2635 | 372500 | 0.000584608 | 0.01522 | 0.015864706 | 0.014899879 |
| pseudoscaff_2635 | 377500 | 0.000275466 | 0.01081 | 0.012338968 | 0.012374734 |
| pseudoscaff_2635 | 387500 | 0.000065576 | 0.01274 | 0.011827449 | 0.01134063 |
| pseudoscaff_2635 | 392500 | 0.000369757 | 0.01148 | 0.013216736 | 0.013622641 |
| pseudoscaff_2635 | 397500 | 0.000361179 | 0.015677 | 0.01219184 | 0.0109694 |
| pseudoscaff_2635 | 407500 | 0.000805671 | 0.012013 | 0.011513396 | 0.013602287 |
| pseudoscaff_2635 | 417500 | 0 | 0.007431 | 0.008605714 | 0.008443909 |
| pseudoscaff_2635 | 422500 | 0.000654034 | 0.00663 | 0.009614802 | 0.009217976 |
| pseudoscaff_1331 | 12500 | 0 | 0.000528 | 0.003373002 | 0.004131174 |

| | | | | | |
|---|---|---|---|---|---|
| pseudoscaff_2510 | 7500 | 0.000645723 | 0.003725 | 0.007008532 | 0.007658152 |
| pseudoscaff_2510 | 132500 | 0 | 0.001939 | 0.003008521 | 0.002159737 |
| pseudoscaff_2510 | 177500 | 0.000275662 | 0.043727 | 0.008163337 | 0.009846202 |
| pseudoscaff_2510 | 182500 | 0.000220275 | 0.003323 | 0.003701381 | 0.002815859 |
| pseudoscaff_429 | 12500 | 0.000929023 | 0.012606 | 0.015642435 | 0.016448567 |
| pseudoscaff_3984 | 117500 | 0.000949205 | 0.006992 | 0.00222594 | 0.003291689 |
| pseudoscaff_3984 | 122500 | 0.000750053 | 0.005483 | 0.001101521 | 0.000622204 |
| pseudoscaff_3713 | 2500 | 0 | 0.000575 | 0.00353956 | 0.008586335 |
| pseudoscaff_630 | 122500 | 0.000313545 | 0.018318 | 0.022771045 | 0.02244173 |
| pseudoscaff_630 | 132500 | 0.001050794 | 0.030537 | 0.029744192 | 0.029349319 |
| pseudoscaff_630 | 147500 | 0.000732011 | 0.034937 | 0.04081923 | 0.041184595 |
| pseudoscaff_630 | 157500 | 0.001588718 | 0.042392 | 0.036686007 | 0.037549773 |
| pseudoscaff_630 | 162500 | 0.000206067 | 0.027889 | 0.031526105 | 0.033099244 |
| pseudoscaff_630 | 167500 | 0.001240618 | 0.016911 | 0.020160167 | 0.019493025 |
| pseudoscaff_630 | 172500 | 0.001416712 | 0.037662 | 0.037040791 | 0.033667022 |
| pseudoscaff_630 | 187500 | 0.001589741 | 0.034352 | 0.037901851 | 0.037337307 |
| pseudoscaff_630 | 202500 | 0.000626502 | 0.018007 | 0.027788909 | 0.028127956 |
| pseudoscaff_630 | 212500 | 0.000265552 | 0.023833 | 0.025198309 | 0.024661872 |
| pseudoscaff_630 | 247500 | 0.000076467 | 0.009086 | 0.009817562 | 0.010329936 |
| pseudoscaff_630 | 252500 | 0.000154322 | 0.017599 | 0.017038812 | 0.017456678 |
| pseudoscaff_630 | 282500 | 0.000357941 | 0.015272 | 0.017949749 | 0.017362605 |
| pseudoscaff_630 | 292500 | 0 | 0.009357 | 0.014623771 | 0.014253055 |
| pseudoscaff_630 | 307500 | 0.001594865 | 0.033094 | 0.035058203 | 0.037813035 |
| pseudoscaff_630 | 312500 | 0.000058392 | 0.02562 | 0.027490868 | 0.028483887 |
| pseudoscaff_630 | 322500 | 0.000218082 | 0.019856 | 0.031204503 | 0.030038287 |
| pseudoscaff_630 | 332500 | 0 | 0.008466 | 0.032296784 | 0.030071325 |
| pseudoscaff_630 | 337500 | 0 | 0.017954 | 0.02496061 | 0.026230877 |
| pseudoscaff_630 | 342500 | 0.000070332 | 0.021405 | 0.029402186 | 0.026321572 |
| pseudoscaff_630 | 347500 | 0.000750487 | 0.017673 | 0.01630208 | 0.017370066 |
| pseudoscaff_630 | 352500 | 0.000391237 | 0.017038 | 0.017067091 | 0.017510538 |
| pseudoscaff_630 | 357500 | 0.000631736 | 0.011243 | 0.012152864 | 0.012407104 |
| pseudoscaff_630 | 362500 | 0.001824917 | 0.010723 | 0.016598281 | 0.016165858 |
| pseudoscaff_630 | 367500 | 0.000216333 | 0.00888 | 0.009571777 | 0.008960648 |
| pseudoscaff_630 | 377500 | 0.000283443 | 0.016627 | 0.02267123 | 0.022927932 |
| pseudoscaff_630 | 382500 | 0.001090074 | 0.019975 | 0.029394471 | 0.029915102 |
| pseudoscaff_630 | 387500 | 0.000153052 | 0.009086 | 0.009932858 | 0.011047896 |
| pseudoscaff_630 | 392500 | 0.000399355 | 0.016898 | 0.016349753 | 0.017120096 |
| pseudoscaff_630 | 397500 | 0.000307092 | 0.018235 | 0.019836635 | 0.020080504 |
| pseudoscaff_630 | 402500 | 0 | 0.015103 | 0.01464623 | 0.013539013 |
| pseudoscaff_630 | 407500 | 0 | 0.010526 | 0.012260383 | 0.011730452 |
| pseudoscaff_630 | 412500 | 0.000729927 | 0.031864 | 0.027162195 | 0.027160292 |
| pseudoscaff_630 | 422500 | 0.001785451 | 0.010278 | 0.013079268 | 0.013443672 |
| pseudoscaff_630 | 427500 | 0.001751299 | 0.014751 | 0.020065037 | 0.01864781 |
| pseudoscaff_178 | 2500 | 0.000090664 | 0.043016 | 0.037602145 | 0.037056285 |
| pseudoscaff_2538 | 1302500 | 0.000354048 | 0.011987 | 0.019617955 | 0.022879743 |
| pseudoscaff_2538 | 1307500 | 0.000156919 | 0.016867 | 0.024466321 | 0.02688941 |
| pseudoscaff_3360 | 182500 | 0.001399855 | 0.015535 | 0.008894254 | 0.010347147 |
| pseudoscaff_3360 | 467500 | 0.0011325 | 0.030334 | 0.026594002 | 0.030971429 |
| pseudoscaff_686 | 22500 | 0.001552859 | 0.002195 | 0.004586727 | 0.004839227 |
| pseudoscaff_328 | 27500 | 0.000877789 | 0.027444 | 0.03841786 | 0.044452571 |

| | | | | | |
|---|---|---|---|---|---|
| pseudoscaff_328 | 42500 | 0 | 0.033737 | 0.038183598 | 0.03843644 |
| pseudoscaff_3559 | 2500 | 0 | 0.012893 | 0.006800357 | 0.006975742 |
| pseudoscaff_3559 | 7500 | 0 | 0.002678 | 0.011408977 | 0.008304179 |
| pseudoscaff_3559 | 12500 | 0.000275522 | 0.006302 | 0.010337196 | 0.010794587 |
| pseudoscaff_3559 | 22500 | 0.001330917 | 0.008737 | 0.015811936 | 0.014663652 |
| pseudoscaff_3559 | 27500 | 0.001976923 | 0.007976 | 0.008735504 | 0.007541329 |
| pseudoscaff_3559 | 32500 | 0.001730907 | 0.004291 | 0.006987402 | 0.006729527 |
| pseudoscaff_3559 | 37500 | 0.000737312 | 0.010322 | 0.010251976 | 0.010909962 |
| pseudoscaff_3559 | 47500 | 0.000254946 | 0.009658 | 0.009406607 | 0.009342352 |
| pseudoscaff_3559 | 57500 | 0 | 0.004699 | 0.006375464 | 0.007829155 |
| pseudoscaff_3559 | 62500 | 0.000092596 | 0.021139 | 0.010390411 | 0.011218628 |
| pseudoscaff_3559 | 67500 | 0 | 0.006971 | 0.006514452 | 0.006714857 |
| pseudoscaff_3559 | 72500 | 0.000742418 | 0.009589 | 0.01165109 | 0.010808974 |
| pseudoscaff_3559 | 77500 | 0 | 0.011366 | 0.009025885 | 0.008568081 |
| pseudoscaff_3559 | 82500 | 0 | 0.004746 | 0.010562409 | 0.009333749 |
| pseudoscaff_396 | 12500 | 0.000398384 | 0.004162 | 0.010695384 | 0.010484302 |
| pseudoscaff_396 | 17500 | 0.001758488 | 0.014918 | 0.024128108 | 0.015893525 |
| pseudoscaff_396 | 22500 | 0.001364121 | 0.008386 | 0.021130275 | 0.014613747 |
| pseudoscaff_396 | 27500 | 0.00108562 | 0.006939 | 0.013575787 | 0.011958026 |
| pseudoscaff_396 | 32500 | 0.000400737 | 0.00879 | 0.012901417 | 0.012938358 |
| pseudoscaff_3750 | 2500 | 0.001841057 | 0.007301 | 0.00724453 | 0.007662005 |
| pseudoscaff_3750 | 42500 | 0.0005057 | 0.001438 | 0.002566693 | 0.002968836 |
| pseudoscaff_1781 | 2500 | 0.001740273 | 0.016118 | 0.018944652 | 0.021448711 |
| pseudoscaff_1781 | 7500 | 0.001510349 | 0.018506 | 0.015027093 | 0.017357297 |
| pseudoscaff_1781 | 22500 | 0.000579969 | 0.028128 | 0.025853972 | 0.026632521 |
| pseudoscaff_1781 | 27500 | 0.000446011 | 0.035731 | 0.03806841 | 0.039296762 |
| pseudoscaff_1781 | 32500 | 0.00043127 | 0.030549 | 0.032133566 | 0.03315018 |
| pseudoscaff_1781 | 37500 | 0.00150035 | 0.026299 | 0.032044565 | 0.034108544 |
| pseudoscaff_1781 | 42500 | 0.000874187 | 0.024875 | 0.033166048 | 0.030918317 |
| pseudoscaff_1781 | 52500 | 0.000568089 | 0.015754 | 0.013186834 | 0.0127189 |
| pseudoscaff_1781 | 57500 | 0.001525682 | 0.014623 | 0.011767691 | 0.012368311 |
| pseudoscaff_1781 | 62500 | 0.000580533 | 0.01303 | 0.013262438 | 0.013902509 |
| pseudoscaff_1781 | 67500 | 0.000946642 | 0.009889 | 0.011563241 | 0.011453552 |
| pseudoscaff_1781 | 82500 | 0.001671074 | 0.013403 | 0.014083139 | 0.015125563 |
| pseudoscaff_1781 | 87500 | 0.001265433 | 0.006419 | 0.007114553 | 0.006985197 |
| pseudoscaff_1781 | 92500 | 0.000636021 | 0.012342 | 0.013506662 | 0.01289086 |
| pseudoscaff_1781 | 97500 | 0.000536513 | 0.011621 | 0.010120292 | 0.011646025 |
| pseudoscaff_1781 | 102500 | 0 | 0.008759 | 0.008557819 | 0.008487188 |
| pseudoscaff_1781 | 107500 | 0.001358444 | 0.012285 | 0.012123847 | 0.013045558 |
| pseudoscaff_854 | 2500 | 0.000123364 | 0.000664 | 0.002619775 | 0.002473533 |
| pseudoscaff_2827 | 2500 | 0.000481738 | 0.001243 | 0.002569519 | 0.003846056 |
| pseudoscaff_4018 | 12500 | 0.00039175 | 0.028415 | 0.025798189 | 0.027738601 |
| pseudoscaff_4018 | 17500 | 0 | 0.006584 | 0.011389754 | 0.016215194 |
| pseudoscaff_4018 | 37500 | 0.000074607 | 0.002491 | 0.01257774 | 0.009739054 |
| pseudoscaff_4018 | 47500 | 0 | 0.006034 | 0.00723329 | 0.008029004 |
| pseudoscaff_4018 | 62500 | 0.000662769 | 0.012762 | 0.017040171 | 0.013081862 |
| pseudoscaff_4018 | 67500 | 0.000126045 | 0.009754 | 0.013151288 | 0.011927956 |
| pseudoscaff_4018 | 82500 | 0 | 0.003225 | 0.012183985 | 0.011444536 |
| pseudoscaff_2387 | 2500 | 0 | 0.003175 | 0.005645233 | 0.00510354 |
| pseudoscaff_2387 | 12500 | 0 | 0.002352 | 0.005054374 | 0.00604216 |

| | | | | | |
|---|---|---|---|---|---|
| pseudoscaff_2387 | 17500 | 0 | 0.002701 | 0.003828186 | 0.004238633 |
| pseudoscaff_2387 | 22500 | 0 | 0.004234 | 0.005330264 | 0.007155003 |
| pseudoscaff_2387 | 37500 | 0.000449881 | 0.008627 | 0.007084344 | 0.006471722 |
| pseudoscaff_2387 | 47500 | 0.001373781 | 0.003022 | 0.019047828 | 0.016816099 |
| pseudoscaff_2387 | 57500 | 0 | 0.00478 | 0.013198065 | 0.013706904 |
| pseudoscaff_2387 | 62500 | 0.000438176 | 0.00487 | 0.003423597 | 0.003602651 |
| pseudoscaff_2387 | 67500 | 0 | 0.021351 | 0.006299855 | 0.010260338 |
| pseudoscaff_1918 | 2500 | 0 | 0.004946 | 0.002614214 | 0.002560283 |
| pseudoscaff_1918 | 12500 | 0.000925151 | 0.009088 | 0.0071927 | 0.00688985 |
| pseudoscaff_1918 | 22500 | 0.000590511 | 0.006263 | 0.012962972 | 0.014563449 |
| pseudoscaff_1872 | 2500 | 0.000512491 | 0.036921 | 0.034250844 | 0.03417342 |
| pseudoscaff_221 | 2500 | 0.000092671 | 0.017372 | 0.013640217 | 0.016194025 |
| pseudoscaff_1659 | 2500 | 0.001029724 | 0.01086 | 0.014757886 | 0.015013447 |
| pseudoscaff_895 | 47500 | 0.001329963 | 0.002448 | 0.003831686 | 0.004004027 |
| pseudoscaff_2119 | 2500 | 0.000460157 | 0.016055 | 0.018100085 | 0.016441724 |
| pseudoscaff_2119 | 7500 | 0.000086768 | 0.020689 | 0.016170679 | 0.016256835 |
| pseudoscaff_3654 | 32500 | 0.001340685 | 0.011264 | 0.012367276 | 0.014054952 |
| pseudoscaff_1377 | 27500 | 0.000167169 | 0.011754 | 0.016298066 | 0.017312957 |
| pseudoscaff_1377 | 32500 | 0.001382795 | 0.001178 | 0.00467288 | 0.007205188 |
| pseudoscaff_1377 | 37500 | 0.000130171 | 0.008744 | 0.004777687 | 0.005015868 |
| pseudoscaff_1377 | 42500 | 0.000416147 | 0.006701 | 0.007161331 | 0.007105976 |
| pseudoscaff_1377 | 47500 | 0.000205228 | 0.003053 | 0.004397165 | 0.004100528 |
| pseudoscaff_1377 | 52500 | 0.000204219 | 0.007216 | 0.006614682 | 0.0064647 |
| pseudoscaff_1377 | 77500 | 0.001699713 | 0.005194 | 0.007683579 | 0.007379745 |
| pseudoscaff_1377 | 102500 | 0.000135221 | 0.004395 | 0.003317603 | 0.004060646 |
| pseudoscaff_1377 | 117500 | 0.000729579 | 0.001626 | 0.001621861 | 0.001930067 |
| pseudoscaff_1377 | 207500 | 0.00019656 | 0.003688 | 0.004049515 | 0.003760497 |
| pseudoscaff_1377 | 217500 | 0.000974417 | 0.001865 | 0.00294783 | 0.003792524 |
| pseudoscaff_1377 | 222500 | 0.000198956 | 0.001038 | 0.00345725 | 0.003591422 |
| pseudoscaff_624 | 47500 | 0.000163222 | 0.0007 | 0.001121156 | 0.001660669 |
| pseudoscaff_3339 | 7500 | 0 | 0.017544 | 0.016487969 | 0.016763093 |
| pseudoscaff_3339 | 12500 | 0.001685356 | 0.045058 | 0.042136696 | 0.041842357 |
| pseudoscaff_3339 | 17500 | 0 | 0.014875 | 0.018298077 | 0.017804409 |
| pseudoscaff_3339 | 22500 | 0 | 0.036615 | 0.03835174 | 0.038249045 |
| pseudoscaff_3339 | 27500 | 0.001133291 | 0.037682 | 0.041029755 | 0.042966813 |
| pseudoscaff_1409 | 47500 | 0.00046787 | 0.003583 | 0.003356872 | 0.003146939 |
| pseudoscaff_1409 | 52500 | 0.000744295 | 0.006364 | 0.007359375 | 0.007326179 |
| pseudoscaff_3592 | 37500 | 0 | 0.046904 | 0.04832815 | 0.048705237 |
| pseudoscaff_4207 | 2500 | 0.000818961 | 0.004864 | 0.013874905 | 0.013887763 |
| pseudoscaff_4207 | 22500 | 0.000145667 | 0.026093 | 0.032559077 | 0.025170089 |
| pseudoscaff_4207 | 42500 | 0 | 0.035841 | 0.031743016 | 0.029490192 |
| pseudoscaff_4207 | 52500 | 0.000789836 | 0.00741 | 0.012458193 | 0.011331241 |
| pseudoscaff_4207 | 177500 | 0.001150737 | 0.020512 | 0.026724804 | 0.027444648 |
| pseudoscaff_4207 | 182500 | 0 | 0.028993 | 0.032708186 | 0.031148177 |
| pseudoscaff_4207 | 187500 | 0.000371126 | 0.02685 | 0.026923692 | 0.028078901 |
| pseudoscaff_4207 | 212500 | 0.000280921 | 0.026787 | 0.034164693 | 0.031974486 |
| pseudoscaff_1869 | 537500 | 0.001344624 | 0.003012 | 0.002779359 | 0.003275986 |
| pseudoscaff_463 | 22500 | 0.00068944 | 0.031379 | 0.037276004 | 0.037592986 |
| pseudoscaff_463 | 82500 | 0.000792383 | 0.032986 | 0.037388462 | 0.035353203 |
| pseudoscaff_463 | 117500 | 0.000218455 | 0.014209 | 0.019552219 | 0.019914273 |

| | | | | | |
|---|---|---|---|---|---|
| pseudoscaff_463 | 122500 | 0.001564322 | 0.018607 | 0.019453481 | 0.021242833 |
| pseudoscaff_463 | 127500 | 0.000059523 | 0.014793 | 0.019150053 | 0.021014864 |
| pseudoscaff_463 | 142500 | 0.000893927 | 0.022585 | 0.030311866 | 0.034188462 |
| pseudoscaff_463 | 157500 | 0.000287874 | 0.022074 | 0.03062281 | 0.029552174 |
| pseudoscaff_463 | 167500 | 0.001972066 | 0.044577 | 0.043179485 | 0.038848013 |
| pseudoscaff_463 | 177500 | 0.001834737 | 0.035747 | 0.03719808 | 0.033998368 |
| pseudoscaff_463 | 192500 | 0.000815683 | 0.023165 | 0.028506176 | 0.027009173 |
| pseudoscaff_463 | 202500 | 0.001234531 | 0.03784 | 0.034376938 | 0.035494592 |
| pseudoscaff_463 | 217500 | 0.000135733 | 0.029316 | 0.024922425 | 0.025327726 |
| pseudoscaff_463 | 352500 | 0.001626534 | 0.011565 | 0.017997849 | 0.019783564 |
| pseudoscaff_463 | 367500 | 0.001139315 | 0.014699 | 0.018396562 | 0.01765551 |
| pseudoscaff_1014 | 12500 | 0.000693712 | 0.01298 | 0.009459699 | 0.010088014 |
| pseudoscaff_1014 | 22500 | 0.000472646 | 0.003165 | 0.002701187 | 0.003252418 |
| pseudoscaff_1014 | 27500 | 0.000636128 | 0.00572 | 0.008367373 | 0.007563729 |
| pseudoscaff_1014 | 32500 | 0.001652506 | 0.006632 | 0.00755543 | 0.007359207 |
| pseudoscaff_1014 | 37500 | 0.000341802 | 0.005014 | 0.007606794 | 0.007420896 |
| pseudoscaff_1014 | 42500 | 0 | 0.009369 | 0.011517497 | 0.009853651 |
| pseudoscaff_1014 | 47500 | 0.001178154 | 0.010818 | 0.011255602 | 0.012725575 |
| pseudoscaff_1014 | 57500 | 0.000580926 | 0.009664 | 0.008833129 | 0.007502184 |
| pseudoscaff_1014 | 67500 | 0.001504324 | 0.007815 | 0.007470252 | 0.007960715 |
| pseudoscaff_1014 | 72500 | 0.001294409 | 0.006901 | 0.006165013 | 0.005790673 |
| pseudoscaff_1014 | 87500 | 0.001338756 | 0.012397 | 0.015596465 | 0.013515059 |
| pseudoscaff_3660 | 2500 | 0.001722301 | 0.004303 | 0.005534022 | 0.005527967 |
| pseudoscaff_3660 | 7500 | 0.000850892 | 0.005713 | 0.00467094 | 0.005573155 |
| pseudoscaff_3660 | 12500 | 0 | 0.001975 | 0.003907467 | 0.004004097 |
| pseudoscaff_3660 | 17500 | 0.000516353 | 0.013661 | 0.012858974 | 0.01323791 |
| pseudoscaff_3660 | 22500 | 0.000144295 | 0.005822 | 0.006755373 | 0.006890977 |
| pseudoscaff_3660 | 32500 | 0.001662532 | 0.011396 | 0.012800823 | 0.011511466 |
| pseudoscaff_3660 | 37500 | 0.000366299 | 0.008294 | 0.005531575 | 0.0077663 |
| pseudoscaff_3660 | 42500 | 0.001552177 | 0.009764 | 0.019752072 | 0.017574815 |
| pseudoscaff_3660 | 47500 | 0 | 0.003991 | 0.010635412 | 0.009538043 |
| pseudoscaff_3660 | 52500 | 0.001361459 | 0.00927 | 0.012214192 | 0.011761364 |
| pseudoscaff_3660 | 57500 | 0.00053442 | 0.02038 | 0.008774043 | 0.008135824 |
| pseudoscaff_3660 | 62500 | 0.0011542 | 0.014157 | 0.013678623 | 0.014141393 |
| pseudoscaff_2422 | 17500 | 0.000932324 | 0.030833 | 0.032352624 | 0.033454563 |
| pseudoscaff_2422 | 22500 | 0.001725252 | 0.018409 | 0.021232469 | 0.022224576 |
| pseudoscaff_2915 | 7500 | 0.000132346 | 0.022372 | 0.017288038 | 0.014056025 |
| pseudoscaff_2915 | 27500 | 0.000345904 | 0.003479 | 0.003863862 | 0.003076622 |
| pseudoscaff_2915 | 32500 | 0.000139344 | 0.001322 | 0.001760988 | 0.00123542 |
| pseudoscaff_1775 | 217500 | 0.001951717 | 0.003339 | 0.003420357 | 0.004730152 |
| pseudoscaff_2976 | 12500 | 0.001426321 | 0.038119 | 0.039699461 | 0.041440016 |
| pseudoscaff_3248 | 212500 | 0.001324075 | 0.013277 | 0.017361589 | 0.017257658 |
| pseudoscaff_3248 | 217500 | 0.001900182 | 0.009308 | 0.008408081 | 0.008130575 |
| pseudoscaff_3248 | 227500 | 0.001837512 | 0.006174 | 0.008218844 | 0.009077918 |
| pseudoscaff_1427 | 12500 | 0.001956855 | 0.01105 | 0.024995026 | 0.015006515 |
| pseudoscaff_1427 | 22500 | 0.000136189 | 0.003476 | 0.003882288 | 0.004146264 |
| pseudoscaff_1427 | 52500 | 0.000244511 | 0.033203 | 0.022509534 | 0.028508848 |
| pseudoscaff_1427 | 72500 | 0.001703467 | 0.014649 | 0.017660879 | 0.016629645 |
| pseudoscaff_1427 | 77500 | 0.000064633 | 0.002618 | 0.002652025 | 0.002926345 |
| pseudoscaff_1427 | 82500 | 0 | 0.002902 | 0.00554767 | 0.00505538 |

| | | | | | |
|---|---|---|---|---|---|
| pseudoscaff_1427 | 87500 | 0.001000645 | 0.02091 | 0.015675985 | 0.013839753 |
| pseudoscaff_1427 | 107500 | 0.000478002 | 0.002298 | 0.003654595 | 0.003500218 |
| pseudoscaff_1427 | 112500 | 0.000283372 | 0.002567 | 0.005314179 | 0.006302544 |
| pseudoscaff_1427 | 122500 | 0.000363026 | 0.012129 | 0.010320861 | 0.008970104 |
| pseudoscaff_1427 | 127500 | 0.001981427 | 0.012574 | 0.00655211 | 0.006062228 |
| pseudoscaff_1427 | 132500 | 0.000165973 | 0.009406 | 0.005066758 | 0.003849735 |
| pseudoscaff_1427 | 137500 | 0.001453414 | 0.023963 | 0.009828066 | 0.00833707 |
| pseudoscaff_1427 | 147500 | 0.000243434 | 0.024562 | 0.025383856 | 0.019000577 |
| pseudoscaff_1427 | 152500 | 0 | 0.032766 | 0.023264678 | 0.022191172 |
| pseudoscaff_1427 | 187500 | 0.001907598 | 0.02424 | 0.021056443 | 0.019039759 |
| pseudoscaff_1427 | 192500 | 0.001547356 | 0.001774 | 0.010221398 | 0.011436335 |
| pseudoscaff_1427 | 197500 | 0.000148195 | 0.004865 | 0.028775242 | 0.026731483 |
| pseudoscaff_1427 | 202500 | 0.000487047 | 0.004475 | 0.010626144 | 0.009705213 |
| pseudoscaff_1427 | 207500 | 0.00112224 | 0.007414 | 0.017851296 | 0.01936376 |
| pseudoscaff_1427 | 212500 | 0.001801093 | 0.009192 | 0.022264487 | 0.022012323 |
| pseudoscaff_814 | 7500 | 0.000327962 | 0.004866 | 0.018923702 | 0.026331744 |
| pseudoscaff_814 | 12500 | 0.000132031 | 0.002378 | 0.018660369 | 0.023502598 |
| pseudoscaff_814 | 22500 | 0 | 0.000658 | 0.010633504 | 0.011154324 |
| pseudoscaff_814 | 27500 | 0.00116003 | 0.00185 | 0.006096976 | 0.007908855 |
| pseudoscaff_1790 | 12500 | 0.00162418 | 0.028069 | 0.030700243 | 0.032546509 |
| pseudoscaff_1790 | 17500 | 0.000878502 | 0.028375 | 0.028810721 | 0.030668063 |
| pseudoscaff_1790 | 32500 | 0.001446217 | 0.031459 | 0.028604874 | 0.028555055 |
| pseudoscaff_1790 | 37500 | 0.000427225 | 0.035506 | 0.033737333 | 0.036291749 |
| pseudoscaff_1790 | 62500 | 0.001539788 | 0.027914 | 0.027843936 | 0.028779139 |
| pseudoscaff_1790 | 77500 | 0.001825268 | 0.026988 | 0.02431507 | 0.025011999 |
| pseudoscaff_1790 | 172500 | 0.001966813 | 0.018433 | 0.019621434 | 0.021049021 |
| pseudoscaff_1790 | 177500 | 0.000651363 | 0.014329 | 0.019811079 | 0.019326875 |
| pseudoscaff_1790 | 187500 | 0.000589585 | 0.016149 | 0.019207581 | 0.021426806 |
| pseudoscaff_1790 | 202500 | 0.000113824 | 0.016927 | 0.014905808 | 0.016588897 |
| pseudoscaff_1790 | 207500 | 0.000382531 | 0.017039 | 0.020575261 | 0.020007858 |
| pseudoscaff_1790 | 237500 | 0.001129606 | 0.021654 | 0.021261519 | 0.022507596 |
| pseudoscaff_1790 | 247500 | 0.000055319 | 0.001804 | 0.002485858 | 0.002108779 |
| pseudoscaff_1790 | 252500 | 0 | 0.008768 | 0.004773894 | 0.004684111 |
| pseudoscaff_1790 | 257500 | 0.000756627 | 0.019706 | 0.020185748 | 0.018201998 |
| pseudoscaff_1790 | 287500 | 0.000354416 | 0.010982 | 0.011025428 | 0.013004 |
| pseudoscaff_1790 | 297500 | 0.000684827 | 0.017977 | 0.024113763 | 0.022331212 |
| pseudoscaff_1790 | 317500 | 0.000189397 | 0.030428 | 0.036829869 | 0.039685123 |
| pseudoscaff_1790 | 337500 | 0.000166768 | 0.014784 | 0.025464406 | 0.025286237 |
| pseudoscaff_1790 | 737500 | 0.001609584 | 0.024472 | 0.025883067 | 0.020931426 |
| pseudoscaff_1790 | 777500 | 0.001019875 | 0.033442 | 0.034516304 | 0.040660121 |
| pseudoscaff_1132 | 12500 | 0.00025536 | 0.005974 | 0.008646194 | 0.009726059 |
| pseudoscaff_1132 | 37500 | 0.001246603 | 0.008197 | 0.013130662 | 0.012945035 |
| pseudoscaff_1161 | 7500 | 0 | 0.022306 | 0.023503392 | 0.02453957 |
| pseudoscaff_1161 | 17500 | 0.001441032 | 0.018546 | 0.019449489 | 0.018628405 |
| pseudoscaff_1161 | 22500 | 0.00010435 | 0.016074 | 0.021573842 | 0.019568005 |
| pseudoscaff_1161 | 27500 | 0.000407118 | 0.023198 | 0.022979732 | 0.024802186 |
| pseudoscaff_2311 | 517500 | 0.001356418 | 0.023134 | 0.021702228 | 0.023973 |
| pseudoscaff_2311 | 817500 | 0.000482733 | 0.041704 | 0.042426975 | 0.039520977 |
| pseudoscaff_2311 | 822500 | 0 | 0.02535 | 0.033143745 | 0.033166451 |
| pseudoscaff_2311 | 827500 | 0 | 0.011526 | 0.013332973 | 0.013542622 |

| | | | | | |
|---|---|---|---|---|---|
| pseudoscaff_2311 | 832500 | 0 | 0.014868 | 0.014567039 | 0.014573255 |
| pseudoscaff_2311 | 837500 | 0.000077028 | 0.011176 | 0.01153054 | 0.011244194 |
| pseudoscaff_2311 | 842500 | 0 | 0.007483 | 0.008760998 | 0.008082464 |
| pseudoscaff_2311 | 852500 | 0.000147614 | 0.006414 | 0.008425296 | 0.007302491 |
| pseudoscaff_690 | 17500 | 0 | 0.025125 | 0.036295837 | 0.03244974 |
| pseudoscaff_690 | 27500 | 0.001714039 | 0.035234 | 0.033895654 | 0.033666196 |
| pseudoscaff_3543 | 172500 | 0.001610409 | 0.006296 | 0.012862178 | 0.01124891 |
| pseudoscaff_3915 | 7500 | 0.000426698 | 0.040497 | 0.03711997 | 0.037654316 |
| pseudoscaff_182 | 47500 | 0 | 0.022569 | 0.019529961 | 0.01986884 |
| pseudoscaff_182 | 52500 | 0.000116024 | 0.015348 | 0.015110287 | 0.015258766 |
| pseudoscaff_182 | 62500 | 0.000448694 | 0.028706 | 0.036546305 | 0.037612926 |
| pseudoscaff_2855 | 22500 | 0.001757566 | 0.004802 | 0.003198968 | 0.005714521 |
| pseudoscaff_2654 | 12500 | 0.00012441 | 0.003188 | 0.004487796 | 0.005589245 |
| pseudoscaff_2654 | 17500 | 0.001526193 | 0.006572 | 0.005554252 | 0.005332497 |
| pseudoscaff_2618 | 2500 | 0.000819327 | 0.007275 | 0.00729056 | 0.00729017 |
| pseudoscaff_2618 | 27500 | 0.001833837 | 0.024394 | 0.024066472 | 0.02149453 |
| pseudoscaff_2618 | 47500 | 0 | 0.015733 | 0.024240621 | 0.022665323 |
| pseudoscaff_2618 | 57500 | 0.000292484 | 0.023686 | 0.028945937 | 0.026878224 |
| pseudoscaff_2618 | 67500 | 0.000134206 | 0.029565 | 0.026462508 | 0.025592822 |
| pseudoscaff_2618 | 72500 | 0.000578112 | 0.03488 | 0.038441273 | 0.037688015 |
| pseudoscaff_2618 | 82500 | 0.000991154 | 0.024919 | 0.02795183 | 0.027544034 |
| pseudoscaff_2618 | 97500 | 0.000549266 | 0.020596 | 0.024757731 | 0.02606533 |
| pseudoscaff_2618 | 117500 | 0.001256724 | 0.041626 | 0.042859298 | 0.042617063 |
| pseudoscaff_2618 | 122500 | 0.000946179 | 0.03433 | 0.036257952 | 0.036948683 |
| pseudoscaff_2618 | 127500 | 0.000063661 | 0.023153 | 0.02755505 | 0.026071811 |
| pseudoscaff_2618 | 132500 | 0.000598389 | 0.021583 | 0.026495479 | 0.025039179 |
| pseudoscaff_2618 | 137500 | 0.001083367 | 0.027377 | 0.025818203 | 0.026578061 |
| pseudoscaff_2618 | 142500 | 0.000195909 | 0.005964 | 0.010876081 | 0.010225537 |
| pseudoscaff_2618 | 147500 | 0.000391362 | 0.025079 | 0.032117844 | 0.030402283 |
| pseudoscaff_2618 | 157500 | 0.000250109 | 0.012543 | 0.013014002 | 0.012536334 |
| pseudoscaff_2618 | 167500 | 0.000049227 | 0.012491 | 0.013984565 | 0.01160153 |
| pseudoscaff_2618 | 177500 | 0.000077375 | 0.010596 | 0.012066959 | 0.011808267 |
| pseudoscaff_2618 | 187500 | 0.000051515 | 0.017248 | 0.019671928 | 0.02024458 |
| pseudoscaff_2618 | 192500 | 0 | 0.0095 | 0.011510372 | 0.011786234 |
| pseudoscaff_2618 | 202500 | 0.000701977 | 0.01823 | 0.023253764 | 0.023219636 |
| pseudoscaff_3172 | 2500 | 0.000099927 | 0.011322 | 0.009113674 | 0.00908416 |
| pseudoscaff_3172 | 12500 | 0.000350968 | 0.016837 | 0.020534622 | 0.024180992 |
| pseudoscaff_3172 | 17500 | 0 | 0.015076 | 0.02178093 | 0.022224932 |
| pseudoscaff_3172 | 27500 | 0 | 0.010434 | 0.019783832 | 0.022906531 |
| pseudoscaff_3172 | 32500 | 0.000648139 | 0.027792 | 0.006866628 | 0.015495073 |
| pseudoscaff_524 | 17500 | 0.000273543 | 0.025824 | 0.036819135 | 0.034999886 |
| pseudoscaff_524 | 22500 | 0 | 0.019474 | 0.027144111 | 0.029147665 |
| pseudoscaff_524 | 32500 | 0.000401369 | 0.014798 | 0.027319424 | 0.028313191 |
| pseudoscaff_524 | 42500 | 0.00049504 | 0.003813 | 0.005731353 | 0.004748515 |
| pseudoscaff_524 | 47500 | 0 | 0.016917 | 0.012753807 | 0.012840599 |
| pseudoscaff_524 | 92500 | 0.001601522 | 0.025441 | 0.029436842 | 0.027244121 |
| pseudoscaff_524 | 122500 | 0.000237586 | 0.033795 | 0.02517891 | 0.025832097 |
| pseudoscaff_524 | 127500 | 0.000105888 | 0.022718 | 0.022540495 | 0.023682144 |
| pseudoscaff_524 | 132500 | 0 | 0.016675 | 0.022147362 | 0.022398169 |
| pseudoscaff_524 | 152500 | 0 | 0.013874 | 0.016211371 | 0.014830831 |

| | | | | | |
|---|---|---|---|---|---|
| pseudoscaff_524 | 157500 | 0.000228514 | 0.024158 | 0.022270462 | 0.022426831 |
| pseudoscaff_524 | 162500 | 0 | 0.014411 | 0.017626712 | 0.018448178 |
| pseudoscaff_524 | 167500 | 0 | 0.002738 | 0.00281329 | 0.003263834 |
| pseudoscaff_524 | 172500 | 0 | 0.005702 | 0.002937968 | 0.003066362 |
| pseudoscaff_524 | 177500 | 0.000059028 | 0.016313 | 0.020553379 | 0.019058626 |
| pseudoscaff_524 | 187500 | 0 | 0.020728 | 0.029617532 | 0.027155121 |
| pseudoscaff_524 | 192500 | 0 | 0.023491 | 0.023101911 | 0.023881608 |
| pseudoscaff_524 | 197500 | 0.000055262 | 0.019882 | 0.02133089 | 0.023211703 |
| pseudoscaff_524 | 202500 | 0.000102459 | 0.034069 | 0.035409019 | 0.034588323 |
| pseudoscaff_524 | 267500 | 0.000666051 | 0.023671 | 0.029623857 | 0.036284648 |
| pseudoscaff_524 | 282500 | 0 | 0.043925 | 0.044246312 | 0.043399568 |
| pseudoscaff_524 | 292500 | 0.001252554 | 0.020877 | 0.030383279 | 0.030509724 |
| pseudoscaff_524 | 312500 | 0.000147576 | 0.011883 | 0.012220531 | 0.013268003 |
| pseudoscaff_252 | 57500 | 0.000321486 | 0.001099 | 0.001191907 | 0.00116387 |
| pseudoscaff_710 | 12500 | 0.001334236 | 0.03948 | 0.033292751 | 0.035355196 |
| pseudoscaff_4190 | 7500 | 0.001220308 | 0.023535 | 0.028961761 | 0.027353202 |
| pseudoscaff_4190 | 12500 | 0.000766666 | 0.01319 | 0.022656791 | 0.02013245 |
| pseudoscaff_4190 | 22500 | 0.000322328 | 0.020944 | 0.025259353 | 0.024357312 |
| pseudoscaff_4190 | 27500 | 0.000243979 | 0.019022 | 0.020118459 | 0.020735882 |
| pseudoscaff_4190 | 32500 | 0.001358553 | 0.022134 | 0.023621284 | 0.023011889 |
| pseudoscaff_4190 | 37500 | 0.00031286 | 0.017892 | 0.023808459 | 0.022929711 |
| pseudoscaff_4190 | 42500 | 0.00032861 | 0.016635 | 0.022567238 | 0.026836501 |
| pseudoscaff_4190 | 47500 | 0.000340138 | 0.01737 | 0.023086668 | 0.019639565 |
| pseudoscaff_4190 | 57500 | 0.000451457 | 0.011754 | 0.023779722 | 0.024735782 |
| pseudoscaff_4190 | 62500 | 0.001765884 | 0.023974 | 0.0288828 | 0.026453575 |
| pseudoscaff_1506 | 2500 | 0 | 0.031153 | 0.036001812 | 0.0338078 |
| pseudoscaff_1506 | 12500 | 0.000413516 | 0.018046 | 0.028622805 | 0.032000608 |
| pseudoscaff_3693 | 82500 | 0.000245201 | 0.02162 | 0.019342063 | 0.02201938 |
| pseudoscaff_3693 | 127500 | 0.001154624 | 0.020602 | 0.026776102 | 0.02522825 |
| pseudoscaff_3693 | 142500 | 0.001143103 | 0.020337 | 0.023976661 | 0.021715904 |
| pseudoscaff_2667 | 27500 | 0.000600604 | 0.001114 | 0.001814422 | 0.001677901 |
| pseudoscaff_4169 | 7500 | 0.001638947 | 0.004544 | 0.007857085 | 0.007120027 |
| pseudoscaff_4169 | 17500 | 0.001692477 | 0.006121 | 0.007599666 | 0.006093508 |
| pseudoscaff_1434 | 2500 | 0.000322544 | 0.025757 | 0.032112758 | 0.03380175 |
| pseudoscaff_3851 | 12500 | 0 | 0.023251 | 0.028876939 | 0.028626737 |
| pseudoscaff_3851 | 17500 | 0.000056128 | 0.016739 | 0.020365498 | 0.022448849 |
| pseudoscaff_3851 | 22500 | 0 | 0.013849 | 0.018824209 | 0.019410085 |
| pseudoscaff_3851 | 27500 | 0 | 0.016249 | 0.017391935 | 0.018134362 |
| pseudoscaff_3851 | 32500 | 0.000823083 | 0.043766 | 0.041802265 | 0.043583762 |
| pseudoscaff_3851 | 42500 | 0.001732905 | 0.030592 | 0.030498049 | 0.030207773 |
| pseudoscaff_2241 | 2500 | 0 | 0.018938 | 0.012026053 | 0.010419035 |
| pseudoscaff_2241 | 7500 | 0 | 0.019665 | 0.016095998 | 0.015999262 |
| pseudoscaff_2241 | 12500 | 0.000084722 | 0.014587 | 0.012188733 | 0.011584075 |
| pseudoscaff_2241 | 17500 | 0 | 0.009231 | 0.00914456 | 0.007797113 |
| pseudoscaff_2241 | 22500 | 0.001771333 | 0.016861 | 0.013393224 | 0.013625475 |
| pseudoscaff_2241 | 27500 | 0.000651252 | 0.013266 | 0.010440532 | 0.011455098 |
| pseudoscaff_2241 | 32500 | 0.000083072 | 0.017331 | 0.015971239 | 0.01577983 |
| pseudoscaff_2241 | 37500 | 0.000877478 | 0.020481 | 0.019679866 | 0.019950029 |
| pseudoscaff_2241 | 42500 | 0.001432443 | 0.018521 | 0.01539055 | 0.015553273 |
| pseudoscaff_2241 | 62500 | 0.000395982 | 0.007695 | 0.004675992 | 0.005259463 |

| | | | | | |
|---|---|---|---|---|---|
| pseudoscaff_2241 | 67500 | 0.001245788 | 0.014934 | 0.014087655 | 0.013402446 |
| pseudoscaff_2241 | 77500 | 0.001682313 | 0.012243 | 0.00974496 | 0.009682906 |
| pseudoscaff_2241 | 82500 | 0.001013281 | 0.007082 | 0.00823021 | 0.008185916 |
| pseudoscaff_2241 | 97500 | 0.001268674 | 0.008077 | 0.006572594 | 0.006466429 |
| pseudoscaff_2241 | 102500 | 0.000230368 | 0.008491 | 0.009614795 | 0.010569606 |
| pseudoscaff_2517 | 7500 | 0 | 0.029425 | 0.03660722 | 0.032138101 |
| pseudoscaff_2517 | 122500 | 0.00092716 | 0.00578 | 0.007109368 | 0.007193016 |
| pseudoscaff_3098 | 7500 | 0.000236073 | 0.020792 | 0.035071475 | 0.035524611 |
| pseudoscaff_3098 | 17500 | 0 | 0.021335 | 0.027349839 | 0.028213238 |
| pseudoscaff_3098 | 22500 | 0.000427346 | 0.022339 | 0.025981577 | 0.026759846 |
| pseudoscaff_3098 | 37500 | 0.00009643 | 0.020017 | 0.023875867 | 0.02623109 |
| pseudoscaff_3098 | 42500 | 0 | 0.015821 | 0.017982047 | 0.018568346 |
| pseudoscaff_3098 | 47500 | 0 | 0.035363 | 0.03475797 | 0.03570905 |
| pseudoscaff_3098 | 82500 | 0.001636703 | 0.03316 | 0.033538977 | 0.033955243 |
| pseudoscaff_3098 | 112500 | 0.001676482 | 0.026796 | 0.038987595 | 0.038939684 |
| pseudoscaff_3098 | 132500 | 0.001529445 | 0.016037 | 0.020291857 | 0.018306953 |
| pseudoscaff_2297 | 7500 | 0.001255628 | 0 | 0.010262751 | 0.00997072 |
| pseudoscaff_2297 | 17500 | 0.000952467 | 0.000165 | 0.003046003 | 0.00323515 |
| pseudoscaff_1000 | 12500 | 0.000155584 | 0.016485 | 0.026385815 | 0.025813673 |
| pseudoscaff_2763 | 7500 | 0 | 0.034468 | 0.016742737 | 0.012030431 |
| pseudoscaff_3667 | 2500 | 0.000084732 | 0.029263 | 0.03328531 | 0.031568581 |
| pseudoscaff_1972 | 22500 | 0.000686952 | 0.028072 | 0.028067641 | 0.02847916 |
| pseudoscaff_349 | 2500 | 0 | 0.015384 | 0.020638818 | 0.022201314 |
| pseudoscaff_349 | 7500 | 0.000084175 | 0.005116 | 0.004721088 | 0.004867064 |
| pseudoscaff_349 | 12500 | 0 | 0.013858 | 0.017999722 | 0.017302177 |
| pseudoscaff_349 | 17500 | 0.000101386 | 0.0284 | 0.030487397 | 0.031009613 |
| pseudoscaff_349 | 22500 | 0.000137158 | 0.017765 | 0.021005329 | 0.023218188 |
| pseudoscaff_349 | 27500 | 0.001743375 | 0.025384 | 0.030975131 | 0.033060876 |
| pseudoscaff_349 | 47500 | 0.000198294 | 0.027102 | 0.037390157 | 0.038526814 |
| pseudoscaff_349 | 52500 | 0 | 0.032414 | 0.031905675 | 0.034021676 |
| pseudoscaff_349 | 62500 | 0.000150961 | 0.017285 | 0.024241429 | 0.027741928 |
| pseudoscaff_349 | 67500 | 0.000073095 | 0.024915 | 0.023408222 | 0.029258538 |
| pseudoscaff_349 | 72500 | 0.000701331 | 0.019514 | 0.015426827 | 0.017362282 |
| pseudoscaff_349 | 82500 | 0.001086532 | 0.030025 | 0.026139053 | 0.028030421 |
| pseudoscaff_349 | 87500 | 0 | 0.005681 | 0.013136008 | 0.012917495 |
| pseudoscaff_349 | 92500 | 0 | 0.017008 | 0.016477993 | 0.016495261 |
| pseudoscaff_349 | 97500 | 0.001265363 | 0.016116 | 0.011755537 | 0.014561429 |
| pseudoscaff_349 | 102500 | 0 | 0.010792 | 0.013097966 | 0.0145281 |
| pseudoscaff_349 | 107500 | 0.000079316 | 0.014016 | 0.022193331 | 0.022405637 |
| pseudoscaff_349 | 112500 | 0 | 0.014428 | 0.018028024 | 0.01828133 |
| pseudoscaff_349 | 117500 | 0.00123104 | 0.042424 | 0.033379668 | 0.034591789 |
| pseudoscaff_349 | 122500 | 0.000682761 | 0.021511 | 0.029993492 | 0.029024244 |
| pseudoscaff_349 | 127500 | 0.000214309 | 0.022812 | 0.016175764 | 0.017866561 |
| pseudoscaff_349 | 132500 | 0 | 0.02009 | 0.02374677 | 0.026637806 |
| pseudoscaff_349 | 137500 | 0.000508027 | 0.020398 | 0.018276194 | 0.020787394 |
| pseudoscaff_349 | 142500 | 0.000575934 | 0.029913 | 0.030231904 | 0.029966034 |
| pseudoscaff_349 | 147500 | 0 | 0.019543 | 0.022711866 | 0.024976037 |
| pseudoscaff_349 | 162500 | 0 | 0.014317 | 0.018666357 | 0.017425974 |
| pseudoscaff_349 | 167500 | 0.000967434 | 0.025897 | 0.033355387 | 0.03470153 |
| pseudoscaff_349 | 172500 | 0 | 0.021673 | 0.0290606 | 0.028563237 |

| | | | | | |
|---|---|---|---|---|---|
| pseudoscaff_349 | 177500 | 0 | 0.013505 | 0.010107397 | 0.011559751 |
| pseudoscaff_349 | 182500 | 0 | 0.013468 | 0.0131624 | 0.013554014 |
| pseudoscaff_349 | 187500 | 0.000933889 | 0.015568 | 0.012891802 | 0.012602024 |
| pseudoscaff_349 | 197500 | 0.000948341 | 0.026584 | 0.028685095 | 0.026731297 |
| pseudoscaff_349 | 207500 | 0 | 0.019468 | 0.019508629 | 0.021640239 |
| pseudoscaff_349 | 212500 | 0 | 0.025769 | 0.0232733 | 0.025069376 |
| pseudoscaff_349 | 217500 | 0.00120006 | 0.032382 | 0.031725211 | 0.027229 |
| pseudoscaff_349 | 222500 | 0 | 0.027369 | 0.024340836 | 0.024550111 |
| pseudoscaff_349 | 227500 | 0.000205053 | 0.008137 | 0.016897958 | 0.017161682 |
| pseudoscaff_349 | 232500 | 0.000064607 | 0.01058 | 0.010342531 | 0.010010799 |
| pseudoscaff_349 | 237500 | 0 | 0.014325 | 0.01613882 | 0.018789517 |
| pseudoscaff_349 | 242500 | 0 | 0.011018 | 0.013388008 | 0.013204037 |
| pseudoscaff_349 | 247500 | 0 | 0.012675 | 0.012832551 | 0.013825444 |
| pseudoscaff_349 | 252500 | 0 | 0.027073 | 0.022339818 | 0.022253266 |
| pseudoscaff_349 | 257500 | 0 | 0.006576 | 0.004998003 | 0.005363151 |
| pseudoscaff_349 | 262500 | 0 | 0.022587 | 0.027625158 | 0.027891675 |
| pseudoscaff_349 | 267500 | 0.000091454 | 0.027127 | 0.035560307 | 0.034078944 |
| pseudoscaff_349 | 272500 | 0.001819459 | 0.037393 | 0.034499029 | 0.034137334 |
| pseudoscaff_349 | 277500 | 0.000170997 | 0.023202 | 0.024953712 | 0.028761599 |
| pseudoscaff_1242 | 42500 | 0.000681866 | 0.003856 | 0.011609953 | 0.009434403 |
| pseudoscaff_2074 | 27500 | 0.000601471 | 0.028741 | 0.034759455 | 0.035810639 |
| pseudoscaff_1708 | 127500 | 0.000671599 | 0.024652 | 0.032890979 | 0.029442373 |
| pseudoscaff_1708 | 132500 | 0 | 0.022992 | 0.033894193 | 0.034159272 |
| pseudoscaff_1708 | 137500 | 0.00053607 | 0.017958 | 0.024655765 | 0.024965234 |
| pseudoscaff_1708 | 142500 | 0.000118174 | 0.021632 | 0.026890547 | 0.028189325 |
| pseudoscaff_1708 | 147500 | 0.001382963 | 0.007535 | 0.012384627 | 0.011280899 |
| pseudoscaff_1708 | 152500 | 0.000116058 | 0.018169 | 0.01745343 | 0.01759368 |
| pseudoscaff_1708 | 162500 | 0 | 0.003539 | 0.006328847 | 0.006843121 |
| pseudoscaff_1708 | 167500 | 0 | 0.012674 | 0.011395358 | 0.011472735 |
| pseudoscaff_1708 | 172500 | 0 | 0.013241 | 0.014571736 | 0.016663479 |
| pseudoscaff_1708 | 177500 | 0.000071332 | 0.024806 | 0.018463905 | 0.019428988 |
| pseudoscaff_1206 | 2500 | 0 | 0.00464 | 0.002596347 | 0.003468502 |
| pseudoscaff_1206 | 12500 | 0.000352299 | 0.0096 | 0.007696 | 0.007701916 |
| pseudoscaff_1206 | 22500 | 0 | 0.008764 | 0.00766905 | 0.007310393 |
| pseudoscaff_1206 | 37500 | 0 | 0.008522 | 0.005271559 | 0.006832986 |
| pseudoscaff_1206 | 42500 | 0.000689161 | 0.023392 | 0.01487915 | 0.018045259 |
| pseudoscaff_1825 | 2500 | 0.00140992 | 0.022679 | 0.020980178 | 0.020556985 |
| pseudoscaff_3790 | 147500 | 0.001783843 | 0.004691 | 0.008656375 | 0.005018877 |
| pseudoscaff_3790 | 177500 | 0.00120006 | 0.002127 | 0.003344219 | 0.002140355 |
| pseudoscaff_995 | 17500 | 0.000274842 | 0.01662 | 0.014594877 | 0.015095237 |
| pseudoscaff_995 | 27500 | 0.001953923 | 0.018297 | 0.022368338 | 0.022638337 |
| pseudoscaff_995 | 37500 | 0.001788162 | 0.028218 | 0.034370653 | 0.031041117 |
| pseudoscaff_995 | 57500 | 0 | 0.021128 | 0.023160433 | 0.026752936 |
| pseudoscaff_995 | 72500 | 0 | 0.03207 | 0.029288957 | 0.029422522 |
| pseudoscaff_995 | 82500 | 0.001404662 | 0.03243 | 0.031979563 | 0.032250733 |
| pseudoscaff_3756 | 62500 | 0.001204865 | 0.008246 | 0.011851682 | 0.011793076 |
| pseudoscaff_2866 | 2500 | 0 | 0.027031 | 0.034873502 | 0.037356837 |
| pseudoscaff_2866 | 7500 | 0.00006373 | 0.014696 | 0.019218113 | 0.019161249 |
| pseudoscaff_1544 | 22500 | 0.001337674 | 0.008145 | 0.009685504 | 0.010189219 |
| pseudoscaff_1544 | 27500 | 0.000740538 | 0.005265 | 0.004140318 | 0.004641748 |

| | | | | | |
|---|---|---|---|---|---|
| pseudoscaff_1112 | 2500 | 0 | 0.010892 | 0.012935592 | 0.013247694 |
| pseudoscaff_1112 | 7500 | 0.00176677 | 0.014761 | 0.020110652 | 0.019868872 |
| pseudoscaff_1112 | 12500 | 0.000338937 | 0.01064 | 0.015631005 | 0.01715388 |
| pseudoscaff_1112 | 22500 | 0.000666224 | 0.006457 | 0.008920832 | 0.007329307 |
| pseudoscaff_1112 | 27500 | 0.000136159 | 0.007086 | 0.009966823 | 0.008130985 |
| pseudoscaff_1112 | 32500 | 0.000791022 | 0.006481 | 0.005793352 | 0.005467146 |
| pseudoscaff_1112 | 37500 | 0 | 0.004351 | 0.009810531 | 0.008736844 |
| pseudoscaff_1112 | 42500 | 0.00035727 | 0.001781 | 0.00512761 | 0.00448839 |
| pseudoscaff_1112 | 47500 | 0 | 0.004409 | 0.02161549 | 0.020192074 |
| pseudoscaff_1112 | 57500 | 0.000089526 | 0.012185 | 0.010667791 | 0.012178954 |
| pseudoscaff_1112 | 62500 | 0.000197176 | 0.017862 | 0.019669962 | 0.020637119 |
| pseudoscaff_1112 | 67500 | 0 | 0.017634 | 0.013507987 | 0.013585414 |
| pseudoscaff_1112 | 97500 | 0.000238196 | 0.007003 | 0.031197187 | 0.028962275 |
| pseudoscaff_1112 | 102500 | 0.000541932 | 0.015611 | 0.013861821 | 0.015755116 |
| pseudoscaff_1112 | 112500 | 0.000651165 | 0.007338 | 0.009701124 | 0.00953275 |
| pseudoscaff_1112 | 117500 | 0.001692082 | 0.013544 | 0.015071037 | 0.014154863 |
| pseudoscaff_1112 | 122500 | 0.000430683 | 0.019098 | 0.01439816 | 0.01326834 |
| pseudoscaff_1112 | 132500 | 0.000164386 | 0.014198 | 0.019232563 | 0.018675581 |
| pseudoscaff_1112 | 142500 | 0 | 0.016188 | 0.013840467 | 0.012628237 |
| pseudoscaff_1112 | 147500 | 0 | 0.019605 | 0.020684462 | 0.019352932 |
| pseudoscaff_1112 | 152500 | 0 | 0.008307 | 0.013741236 | 0.013995612 |
| pseudoscaff_1112 | 157500 | 0 | 0.015807 | 0.023508761 | 0.023339355 |
| pseudoscaff_1112 | 167500 | 0.000264021 | 0.006413 | 0.015081336 | 0.013818328 |
| pseudoscaff_1112 | 172500 | 0 | 0.014573 | 0.011737053 | 0.011739761 |
| pseudoscaff_1112 | 177500 | 0.000084243 | 0.009323 | 0.009408802 | 0.008758216 |
| pseudoscaff_1112 | 182500 | 0.000091471 | 0.011198 | 0.00897889 | 0.009858662 |
| pseudoscaff_1112 | 187500 | 0 | 0.002065 | 0.006761409 | 0.008355009 |
| pseudoscaff_1112 | 192500 | 0.000477029 | 0.007579 | 0.011035189 | 0.014348915 |
| pseudoscaff_1112 | 197500 | 0 | 0.002351 | 0.011862455 | 0.013225805 |
| pseudoscaff_1112 | 202500 | 0 | 0.015875 | 0.01224679 | 0.012543778 |
| pseudoscaff_1112 | 207500 | 0 | 0.009352 | 0.014779086 | 0.014034632 |
| pseudoscaff_1112 | 212500 | 0 | 0.012204 | 0.01138927 | 0.01055958 |
| pseudoscaff_1112 | 217500 | 0 | 0.013231 | 0.011880266 | 0.00919468 |
| pseudoscaff_1112 | 222500 | 0 | 0.004849 | 0.006977641 | 0.004092727 |
| pseudoscaff_1112 | 227500 | 0 | 0.001988 | 0.004620061 | 0.004229758 |
| pseudoscaff_1112 | 232500 | 0 | 0.024607 | 0.011540366 | 0.013800421 |
| pseudoscaff_1112 | 237500 | 0 | 0.007288 | 0.005545154 | 0.005151551 |
| pseudoscaff_1112 | 242500 | 0.000463692 | 0.015087 | 0.037539057 | 0.031185495 |
| pseudoscaff_1112 | 252500 | 0.000659408 | 0.015544 | 0.013790327 | 0.014918366 |
| pseudoscaff_1112 | 262500 | 0 | 0.004345 | 0.004990443 | 0.004514556 |
| pseudoscaff_1112 | 267500 | 0 | 0.004567 | 0.005049016 | 0.004205411 |
| pseudoscaff_1112 | 277500 | 0.000606879 | 0.014797 | 0.027341377 | 0.02354518 |
| pseudoscaff_1112 | 307500 | 0 | 0.009936 | 0.012693326 | 0.012407067 |
| pseudoscaff_572 | 197500 | 0.000718792 | 0.022374 | 0.022898265 | 0.023350397 |
| pseudoscaff_572 | 202500 | 0.000239849 | 0.029327 | 0.033015442 | 0.034634861 |
| pseudoscaff_572 | 297500 | 0.000653592 | 0.031206 | 0.033121147 | 0.030819961 |
| pseudoscaff_572 | 302500 | 0.000069521 | 0.011889 | 0.015327008 | 0.015022327 |
| pseudoscaff_381 | 2500 | 0.000217601 | 0.006457 | 0.008004697 | 0.007829681 |
| pseudoscaff_381 | 7500 | 0.000320044 | 0.007892 | 0.008615902 | 0.008684141 |
| pseudoscaff_381 | 12500 | 0 | 0.002841 | 0.006518102 | 0.00610891 |

| | | | | | |
|---|---|---|---|---|---|
| pseudoscaff_381 | 17500 | 0 | 0.005274 | 0.007185017 | 0.006786288 |
| pseudoscaff_381 | 22500 | 0 | 0.008215 | 0.011439376 | 0.012932137 |
| pseudoscaff_381 | 27500 | 0 | 0.010755 | 0.007316809 | 0.006719367 |
| pseudoscaff_381 | 32500 | 0.001732389 | 0.040733 | 0.012047794 | 0.012701009 |
| pseudoscaff_970 | 7500 | 0.000309948 | 0.038152 | 0.037768739 | 0.037799042 |
| pseudoscaff_2951 | 7500 | 0.000495854 | 0.005862 | 0.005086675 | 0.006268723 |
| pseudoscaff_2951 | 12500 | 0.001082383 | 0.003285 | 0.007128847 | 0.009284105 |
| pseudoscaff_2951 | 17500 | 0.001127522 | 0.00933 | 0.007719988 | 0.00833255 |
| pseudoscaff_2951 | 22500 | 0 | 0.015034 | 0.015222125 | 0.016204659 |
| pseudoscaff_2951 | 37500 | 0 | 0.004733 | 0.007923085 | 0.007355401 |
| pseudoscaff_2951 | 42500 | 0.001857465 | 0.007874 | 0.008914071 | 0.008801291 |
| pseudoscaff_2951 | 47500 | 0 | 0.002704 | 0.002742859 | 0.002993326 |
| pseudoscaff_2951 | 52500 | 0 | 0.001371 | 0.002018135 | 0.001906169 |
| pseudoscaff_2951 | 57500 | 0 | 0.000339 | 0.00253247 | 0.00207052 |
| pseudoscaff_2790 | 2500 | 0.001754089 | 0.010327 | 0.011177789 | 0.010937914 |
| pseudoscaff_4062 | 352500 | 0.00112221 | 0.028525 | 0.041530615 | 0.039420798 |
| pseudoscaff_4062 | 452500 | 0.001499416 | 0.028781 | 0.025491437 | 0.024079559 |
| pseudoscaff_1369 | 2500 | 0.000096356 | 0.003873 | 0.003401554 | 0.004884753 |
| pseudoscaff_1369 | 7500 | 0.000908317 | 0.010023 | 0.017685032 | 0.016160624 |
| pseudoscaff_1369 | 12500 | 0.001625797 | 0.007823 | 0.012630356 | 0.010139173 |
| pseudoscaff_1369 | 17500 | 0.001094568 | 0.007657 | 0.00476736 | 0.006136209 |
| pseudoscaff_1369 | 22500 | 0.000893036 | 0.011406 | 0.010695506 | 0.011806989 |
| pseudoscaff_740 | 12500 | 0 | 0.000738 | 0.000502909 | 0.000644101 |
| pseudoscaff_1965 | 92500 | 0.00075004 | 0.010065 | 0.01988577 | 0.021408056 |
| pseudoscaff_1554 | 2500 | 0 | 0.003116 | 0.011204555 | 0.009675027 |
| pseudoscaff_1554 | 7500 | 0.000103669 | 0.008428 | 0.01817264 | 0.014857402 |
| pseudoscaff_1554 | 12500 | 0.00078564 | 0.003013 | 0.00410596 | 0.004391034 |
| pseudoscaff_1554 | 22500 | 0.000796816 | 0.005552 | 0.007378103 | 0.005459625 |
| pseudoscaff_1554 | 27500 | 0 | 0.010162 | 0.007367044 | 0.008377894 |
| pseudoscaff_1554 | 42500 | 0.001008641 | 0.001654 | 0.00243707 | 0.002402176 |
| pseudoscaff_1554 | 47500 | 0.000329542 | 0.001253 | 0.000822789 | 0.000999963 |
| pseudoscaff_2349 | 7500 | 0.001371198 | 0.008525 | 0.01321115 | 0.014427875 |
| pseudoscaff_2300 | 607500 | 0.000621333 | 0.000818 | 0.000747496 | 0.000879562 |
| pseudoscaff_3766 | 7500 | 0.001539201 | 0.022008 | 0.025776161 | 0.025974788 |
| pseudoscaff_750 | 7500 | 0 | 0.003382 | 0.007293413 | 0.007367356 |
| pseudoscaff_2351 | 152500 | 0.001454182 | 0.006583 | 0.010933868 | 0.010322955 |
| pseudoscaff_2351 | 162500 | 0.001697026 | 0.015064 | 0.01495126 | 0.014203126 |
| pseudoscaff_3314 | 12500 | 0.001839684 | 0.040521 | 0.042604463 | 0.044463243 |
| pseudoscaff_3161 | 7500 | 0 | 0.024466 | 0.030145059 | 0.030994688 |
| pseudoscaff_3161 | 12500 | 0.000900699 | 0.026811 | 0.024595084 | 0.027469899 |
| pseudoscaff_3161 | 17500 | 0 | 0.021195 | 0.019219691 | 0.01981733 |
| pseudoscaff_3161 | 22500 | 0.000091482 | 0.021767 | 0.022213689 | 0.020251145 |
| pseudoscaff_3161 | 27500 | 0 | 0.022189 | 0.025454829 | 0.026655131 |
| pseudoscaff_3161 | 32500 | 0 | 0.021583 | 0.033464322 | 0.029042331 |
| pseudoscaff_1512 | 12500 | 0.001870362 | 0.0348 | 0.024838048 | 0.026843563 |
| pseudoscaff_1512 | 17500 | 0.000226441 | 0.010182 | 0.006867046 | 0.008105922 |
| pseudoscaff_1512 | 22500 | 0.000152251 | 0.008495 | 0.011126529 | 0.009969618 |
| pseudoscaff_2284 | 2500 | 0 | 0.005709 | 0.009805965 | 0.009798232 |
| pseudoscaff_2284 | 7500 | 0.000055992 | 0.005971 | 0.005427157 | 0.006892501 |
| pseudoscaff_2284 | 17500 | 0.000999586 | 0.012068 | 0.014106584 | 0.014708461 |

| | | | | | |
|---|---|---|---|---|---|
| pseudoscaff_2284 | 22500 | 0.00024638 | 0.013896 | 0.009848726 | 0.012940469 |
| pseudoscaff_2284 | 27500 | 0 | 0.024982 | 0.018580736 | 0.020016738 |
| pseudoscaff_2284 | 32500 | 0.001139691 | 0.009714 | 0.014576509 | 0.015184933 |
| pseudoscaff_2284 | 37500 | 0.000542045 | 0.010416 | 0.017294959 | 0.017062248 |
| pseudoscaff_2284 | 42500 | 0.000803084 | 0.014071 | 0.011503235 | 0.010989794 |
| pseudoscaff_2284 | 47500 | 0.000149386 | 0.009926 | 0.017530628 | 0.015123702 |
| pseudoscaff_2284 | 57500 | 0.00122236 | 0.021399 | 0.02150011 | 0.018659461 |
| pseudoscaff_2284 | 67500 | 0.001868647 | 0.009575 | 0.017483041 | 0.016638561 |
| pseudoscaff_1815 | 2500 | 0.00168901 | 0.007693 | 0.009954111 | 0.008235168 |
| pseudoscaff_1815 | 12500 | 0.000383411 | 0.012185 | 0.010522428 | 0.012397666 |
| pseudoscaff_1815 | 17500 | 0.000122767 | 0.006619 | 0.006194983 | 0.005312895 |
| pseudoscaff_1815 | 22500 | 0.000139626 | 0.010326 | 0.013955868 | 0.013176932 |
| pseudoscaff_1815 | 37500 | 0 | 0.002901 | 0.004676257 | 0.005074306 |
| pseudoscaff_1079 | 27500 | 0.001575243 | 0.004526 | 0.003171903 | 0.002676572 |
| pseudoscaff_1338 | 12500 | 0.001820757 | 0.006473 | 0.008162885 | 0.007716721 |
| pseudoscaff_1086 | 12500 | 0.001418013 | 0.016727 | 0.018243862 | 0.018380701 |
| pseudoscaff_4033 | 7500 | 0.000526762 | 0.007247 | 0.007369523 | 0.006809947 |
| pseudoscaff_4033 | 12500 | 0.001975081 | 0.012947 | 0.010110632 | 0.012561939 |
| pseudoscaff_3832 | 2500 | 0 | 0.004244 | 0.003837815 | 0.003052888 |
| pseudoscaff_3832 | 7500 | 0 | 0.002241 | 0.004130114 | 0.004490038 |
| pseudoscaff_3832 | 12500 | 0 | 0.003962 | 0.002199621 | 0.002757043 |
| pseudoscaff_3832 | 17500 | 0 | 0.00351 | 0.002924703 | 0.003418173 |
| pseudoscaff_3832 | 22500 | 0 | 0.002765 | 0.007053243 | 0.007214834 |
| pseudoscaff_1088 | 17500 | 0.000849806 | 0.028607 | 0.043881337 | 0.044257138 |
| pseudoscaff_1885 | 17500 | 0 | 0.021948 | 0.026108786 | 0.027111958 |
| pseudoscaff_1885 | 22500 | 0.00183912 | 0.029213 | 0.03244337 | 0.024896603 |
| pseudoscaff_1885 | 27500 | 0.001269516 | 0.027762 | 0.034809817 | 0.036518273 |
| pseudoscaff_1885 | 37500 | 0.000237479 | 0.023183 | 0.021455152 | 0.020973496 |
| pseudoscaff_1885 | 102500 | 0.000161908 | 0.016083 | 0.022368618 | 0.022956871 |
| pseudoscaff_1885 | 172500 | 0 | 0.024121 | 0.020889037 | 0.022443329 |
| pseudoscaff_1885 | 202500 | 0 | 0.020699 | 0.03037076 | 0.027587425 |
| pseudoscaff_1885 | 207500 | 0.000457014 | 0.02708 | 0.031631593 | 0.031329288 |
| pseudoscaff_1182 | 217500 | 0.000837574 | 0.018386 | 0.018747011 | 0.022365514 |
| pseudoscaff_1182 | 222500 | 0 | 0.013674 | 0.016585932 | 0.016714339 |
| pseudoscaff_1182 | 227500 | 0.000064896 | 0.018411 | 0.017611345 | 0.017382413 |
| pseudoscaff_1182 | 232500 | 0.000088462 | 0.023065 | 0.020595579 | 0.019809477 |
| pseudoscaff_688 | 2500 | 0.000466452 | 0.003808 | 0.016580882 | 0.013187884 |
| pseudoscaff_688 | 12500 | 0.000986802 | 0.00394 | 0.019317947 | 0.015595411 |
| pseudoscaff_117 | 2500 | 0.000339942 | 0.030066 | 0.012552842 | 0.014660461 |
| pseudoscaff_117 | 7500 | 0.000398866 | 0.033702 | 0.022333482 | 0.019814566 |
| pseudoscaff_706 | 2500 | 0.00018751 | 0.005489 | 0.006015707 | 0.007861438 |
| pseudoscaff_706 | 7500 | 0.001552393 | 0.001118 | 0.002143571 | 0.003382642 |
| pseudoscaff_706 | 12500 | 0.000371377 | 0.000217 | 0.000336281 | 0.0011007 |
| pseudoscaff_706 | 17500 | 0 | 0.000854 | 0.000879747 | 0.001281881 |
| pseudoscaff_706 | 22500 | 0 | 0.00047 | 0.001111099 | 0.001796867 |
| pseudoscaff_706 | 27500 | 0 | 0.006129 | 0.008757777 | 0.006284428 |
| pseudoscaff_561 | 2500 | 0.001130829 | 0.00347 | 0.011532786 | 0.011602031 |
| pseudoscaff_561 | 12500 | 0.001055887 | 0.003826 | 0.005701374 | 0.006360826 |
| pseudoscaff_561 | 22500 | 0.001584109 | 0.008686 | 0.011799136 | 0.0122162 |
| pseudoscaff_561 | 42500 | 0.000695083 | 0.013503 | 0.015586767 | 0.013766404 |

| | | | | | |
|---|---|---|---|---|---|
| pseudoscaff_3948 | 47500 | 0 | 0.026828 | 0.024291631 | 0.027462293 |
| pseudoscaff_3948 | 52500 | 0 | 0.029747 | 0.030617065 | 0.036407563 |
| pseudoscaff_3948 | 167500 | 0.000654131 | 0.013141 | 0.021567875 | 0.024377628 |
| pseudoscaff_3948 | 337500 | 0.001472896 | 0.034146 | 0.032718031 | 0.034510934 |
| pseudoscaff_3147 | 2500 | 0 | 0.024568 | 0.009322557 | 0.011606818 |
| pseudoscaff_3147 | 12500 | 0 | 0.00426 | 0.012234348 | 0.011831916 |
| pseudoscaff_3147 | 17500 | 0 | 0.002969 | 0.002976257 | 0.002723933 |
| pseudoscaff_3147 | 27500 | 0 | 0.003385 | 0.001602864 | 0.001712147 |
| pseudoscaff_3147 | 32500 | 0.000356433 | 0.000593 | 0.000927512 | 0.000859722 |
| pseudoscaff_3147 | 37500 | 0.000346936 | 0.000829 | 0.004149974 | 0.003893025 |
| pseudoscaff_2079 | 2500 | 0.000259649 | 0.00015 | 0.000582895 | 0.001015996 |
| pseudoscaff_2079 | 7500 | 0 | 0 | 0.000062098 | 0.000021272 |
| pseudoscaff_2079 | 12500 | 0.000705048 | 0.000473 | 0.000443205 | 0.000716262 |
| pseudoscaff_2079 | 17500 | 0 | 0 | 0.000089414 | 0.0000839 |
| pseudoscaff_2079 | 22500 | 0.000560283 | 0 | 0.000040629 | 0 |
| pseudoscaff_1730 | 12500 | 0.001255479 | 0.033348 | 0.037646808 | 0.036294189 |
| pseudoscaff_1730 | 32500 | 0.001702518 | 0.036371 | 0.041030356 | 0.041914273 |
| pseudoscaff_1730 | 37500 | 0 | 0.034909 | 0.036753184 | 0.036956221 |
| pseudoscaff_1730 | 42500 | 0.00099542 | 0.041115 | 0.042218818 | 0.043792401 |
| pseudoscaff_1730 | 47500 | 0.000475045 | 0.025427 | 0.033716349 | 0.031646153 |
| pseudoscaff_1730 | 52500 | 0.000293103 | 0.02886 | 0.028347731 | 0.028950914 |
| pseudoscaff_1730 | 57500 | 0 | 0.02869 | 0.028543852 | 0.030113342 |
| pseudoscaff_1730 | 62500 | 0.000406112 | 0.040635 | 0.033339228 | 0.034929558 |
| pseudoscaff_1730 | 67500 | 0.000121465 | 0.024579 | 0.01855828 | 0.019284669 |
| pseudoscaff_1730 | 72500 | 0.000120354 | 0.022883 | 0.023291752 | 0.026808155 |
| pseudoscaff_1730 | 107500 | 0 | 0.040441 | 0.038008597 | 0.040598224 |
| pseudoscaff_1730 | 112500 | 0 | 0.021348 | 0.031039002 | 0.030356699 |
| pseudoscaff_1730 | 122500 | 0.001792228 | 0.031393 | 0.032541566 | 0.030252062 |
| pseudoscaff_1730 | 127500 | 0.00116038 | 0.028153 | 0.027124141 | 0.028272171 |
| pseudoscaff_2462 | 17500 | 0.000529658 | 0.014217 | 0.022096497 | 0.021809656 |
| pseudoscaff_3640 | 17500 | 0.001910205 | 0.010256 | 0.009544176 | 0.009870622 |
| pseudoscaff_2740 | 2500 | 0.000066625 | 0.018116 | 0.007326724 | 0.010329151 |
| pseudoscaff_2740 | 7500 | 0.00011137 | 0.008677 | 0.011071404 | 0.00945566 |
| pseudoscaff_2740 | 17500 | 0.00125152 | 0.004865 | 0.00855177 | 0.007990479 |
| pseudoscaff_2740 | 22500 | 0 | 0.003718 | 0.005085411 | 0.005514372 |
| pseudoscaff_2740 | 27500 | 0 | 0.002015 | 0.002659619 | 0.002722679 |
| pseudoscaff_2740 | 32500 | 0.000175475 | 0.003754 | 0.00437112 | 0.00491903 |
| pseudoscaff_2740 | 37500 | 0.000826196 | 0.005345 | 0.003516857 | 0.003532881 |
| pseudoscaff_2740 | 42500 | 0.001553815 | 0.007342 | 0.010378225 | 0.014189151 |
| pseudoscaff_2740 | 47500 | 0.000190902 | 0.002509 | 0.002541532 | 0.00243849 |
| pseudoscaff_2740 | 52500 | 0.001011494 | 0.001149 | 0.00220755 | 0.003162719 |
| pseudoscaff_2740 | 57500 | 0 | 0.001178 | 0.002826221 | 0.00367395 |
| pseudoscaff_2740 | 62500 | 0.000423709 | 0.00715 | 0.007853159 | 0.00768005 |
| pseudoscaff_2740 | 82500 | 0.001432822 | 0.001441 | 0.006998536 | 0.008390545 |
| pseudoscaff_2740 | 87500 | 0 | 0.002766 | 0.003100084 | 0.003040152 |
| pseudoscaff_1376 | 7500 | 0.000556592 | 0.002317 | 0.008527151 | 0.007464692 |
| pseudoscaff_1376 | 17500 | 0 | 0.000746 | 0.000778376 | 0.000890914 |
| pseudoscaff_1376 | 22500 | 0.000639146 | 0.002749 | 0.012689461 | 0.010228228 |
| pseudoscaff_2110 | 7500 | 0.001639879 | 0.006304 | 0.015035668 | 0.010344798 |
| pseudoscaff_2110 | 12500 | 0.000583122 | 0.00476 | 0.007577284 | 0.006915065 |

| | | | | | |
|---|---|---|---|---|---|
| pseudoscaff_2110 | 17500 | 0.000300734 | 0.008199 | 0.00398944 | 0.006451269 |
| pseudoscaff_2110 | 22500 | 0.00162398 | 0.007872 | 0.007934581 | 0.008254015 |
| pseudoscaff_2110 | 27500 | 0.000402607 | 0.014513 | 0.007772617 | 0.008962598 |
| pseudoscaff_3659 | 17500 | 0.000196651 | 0.028808 | 0.037070501 | 0.033976707 |
| pseudoscaff_3659 | 22500 | 0 | 0.027553 | 0.034259263 | 0.032373527 |
| pseudoscaff_3659 | 27500 | 0.000061712 | 0.022881 | 0.02634334 | 0.026520832 |
| pseudoscaff_3659 | 32500 | 0 | 0.022949 | 0.026539594 | 0.026983374 |
| pseudoscaff_3659 | 37500 | 0 | 0.019316 | 0.024000295 | 0.024283694 |
| pseudoscaff_3659 | 42500 | 0 | 0.02367 | 0.025568629 | 0.026381148 |
| pseudoscaff_3206 | 2500 | 0 | 0.01147 | 0.014674967 | 0.015480539 |
| pseudoscaff_3206 | 7500 | 0 | 0.011155 | 0.010398536 | 0.010337541 |
| pseudoscaff_3206 | 12500 | 0 | 0.024731 | 0.021893169 | 0.022967873 |
| pseudoscaff_3206 | 17500 | 0 | 0.020548 | 0.022973405 | 0.022112979 |
| pseudoscaff_3206 | 22500 | 0 | 0.018079 | 0.027690051 | 0.025101926 |
| pseudoscaff_3206 | 27500 | 0.000827546 | 0.019573 | 0.01569899 | 0.014367766 |
| pseudoscaff_3206 | 32500 | 0.000100228 | 0.015531 | 0.016881249 | 0.017017738 |
| pseudoscaff_3206 | 37500 | 0.000574671 | 0.021763 | 0.02046564 | 0.020082671 |
| pseudoscaff_3206 | 42500 | 0 | 0.024807 | 0.026048054 | 0.025864943 |
| pseudoscaff_3206 | 52500 | 0 | 0.016504 | 0.018332674 | 0.017343055 |
| pseudoscaff_3206 | 57500 | 0 | 0.01354 | 0.016942649 | 0.019951967 |
| pseudoscaff_3206 | 62500 | 0 | 0.019685 | 0.02042448 | 0.017020357 |
| pseudoscaff_3206 | 67500 | 0 | 0.015938 | 0.013381814 | 0.011200907 |
| pseudoscaff_3206 | 72500 | 0.000257537 | 0.012706 | 0.02107227 | 0.014851751 |
| pseudoscaff_3206 | 77500 | 0 | 0.028616 | 0.027273999 | 0.025763141 |
| pseudoscaff_3206 | 82500 | 0 | 0.025263 | 0.024599097 | 0.024218469 |
| pseudoscaff_3206 | 92500 | 0 | 0.015352 | 0.016655205 | 0.017435226 |
| pseudoscaff_3206 | 102500 | 0.000707107 | 0.037218 | 0.029965368 | 0.03210376 |
| pseudoscaff_3206 | 117500 | 0.000192032 | 0.030232 | 0.020873801 | 0.02316701 |
| pseudoscaff_704 | 2500 | 0 | 0.005789 | 0.006800327 | 0.00663619 |
| pseudoscaff_704 | 7500 | 0.001448402 | 0.006584 | 0.009643205 | 0.008709824 |
| pseudoscaff_704 | 17500 | 0.000725799 | 0.004739 | 0.009744361 | 0.009042706 |
| pseudoscaff_704 | 32500 | 0.000710945 | 0.008393 | 0.008684426 | 0.009011769 |
| pseudoscaff_704 | 37500 | 0.001211874 | 0.010297 | 0.00553855 | 0.008459902 |
| pseudoscaff_704 | 47500 | 0.001698779 | 0.007423 | 0.012557738 | 0.011957052 |
| pseudoscaff_704 | 52500 | 0.000821946 | 0.006114 | 0.007071905 | 0.006902261 |
| pseudoscaff_704 | 62500 | 0.00195888 | 0.00697 | 0.009205849 | 0.011282795 |
| pseudoscaff_704 | 72500 | 0.001662415 | 0.016725 | 0.011285857 | 0.013607846 |
| pseudoscaff_704 | 82500 | 0 | 0.016508 | 0.011412971 | 0.013952422 |
| pseudoscaff_704 | 97500 | 0.001978784 | 0.006545 | 0.006885464 | 0.006550619 |
| pseudoscaff_704 | 107500 | 0.001972002 | 0.003623 | 0.004203858 | 0.00501389 |
| pseudoscaff_704 | 117500 | 0.000947129 | 0.002687 | 0.004569457 | 0.003896807 |
| pseudoscaff_704 | 147500 | 0.001785448 | 0.012288 | 0.005775512 | 0.005683123 |
| pseudoscaff_559 | 12500 | 0 | 0.024001 | 0.030313666 | 0.031368456 |
| pseudoscaff_559 | 17500 | 0.000077954 | 0.027091 | 0.03178816 | 0.033429379 |
| pseudoscaff_559 | 22500 | 0.001620408 | 0.026467 | 0.022229325 | 0.024398438 |
| pseudoscaff_1591 | 2500 | 0.000924526 | 0.001713 | 0.001653296 | 0.001783004 |
| pseudoscaff_1591 | 12500 | 0.000080148 | 0.002528 | 0.002442061 | 0.002233897 |
| pseudoscaff_1591 | 17500 | 0.001719299 | 0.001946 | 0.004650579 | 0.00313519 |
| pseudoscaff_1591 | 22500 | 0 | 0.004573 | 0.002488912 | 0.002737157 |
| pseudoscaff_1591 | 27500 | 0 | 0.00543 | 0.004082962 | 0.003789437 |

| | | | | | |
|---|---|---|---|---|---|
| pseudoscaff_1591 | 37500 | 0.001888672 | 0.037736 | 0.024292665 | 0.025763081 |
| pseudoscaff_1394 | 47500 | 0.001759928 | 0.0035 | 0.00325096 | 0.002663386 |
| pseudoscaff_1260 | 12500 | 0.00113884 | 0.002715 | 0.006284981 | 0.006496983 |
| pseudoscaff_1260 | 17500 | 0.001337779 | 0.003685 | 0.007934002 | 0.007372445 |
| pseudoscaff_1260 | 22500 | 0.000888858 | 0.004518 | 0.008518839 | 0.009476939 |
| pseudoscaff_2114 | 12500 | 0.000064888 | 0.020536 | 0.021270145 | 0.02001099 |
| pseudoscaff_2114 | 17500 | 0.001349241 | 0.025866 | 0.030316154 | 0.027283371 |
| pseudoscaff_2114 | 22500 | 0.000267007 | 0.023681 | 0.025723336 | 0.02678003 |
| pseudoscaff_2569 | 2500 | 0.000263416 | 0.0162 | 0.027610704 | 0.029029547 |
| pseudoscaff_2569 | 7500 | 0 | 0.007624 | 0.010572389 | 0.010769184 |
| pseudoscaff_2569 | 12500 | 0.000701525 | 0.029322 | 0.027474112 | 0.025563252 |
| pseudoscaff_2569 | 17500 | 0.000124715 | 0.023276 | 0.022739196 | 0.024178282 |
| pseudoscaff_2569 | 27500 | 0 | 0.011149 | 0.010496276 | 0.011422732 |
| pseudoscaff_1528 | 222500 | 0.000167502 | 0.035248 | 0.035290017 | 0.035533583 |
| pseudoscaff_1528 | 227500 | 0.000044422 | 0.025402 | 0.036423231 | 0.033362045 |
| pseudoscaff_1528 | 242500 | 0.000110021 | 0.019896 | 0.024670202 | 0.023967116 |
| pseudoscaff_1528 | 272500 | 0.000194177 | 0.028539 | 0.028448018 | 0.032128113 |
| pseudoscaff_1528 | 342500 | 0.000854802 | 0.033573 | 0.042770628 | 0.041338576 |
| pseudoscaff_1528 | 362500 | 0.001469681 | 0.03127 | 0.052323918 | 0.047553333 |
| pseudoscaff_1528 | 627500 | 0.000868723 | 0.0281 | 0.042895811 | 0.044470002 |
| pseudoscaff_428 | 7500 | 0 | 0.023098 | 0.031747074 | 0.029448489 |
| pseudoscaff_428 | 37500 | 0.000199658 | 0.048813 | 0.017756366 | 0.02160119 |
| pseudoscaff_1415 | 27500 | 0.000405461 | 0.043153 | 0.042299486 | 0.046591309 |
| pseudoscaff_538 | 92500 | 0.001887133 | 0.002655 | 0.003169797 | 0.0031976 |
| pseudoscaff_146 | 2500 | 0.001011463 | 0.02012 | 0.021159306 | 0.021220498 |
| pseudoscaff_343 | 12500 | 0 | 0.028041 | 0.03403642 | 0.035914179 |
| pseudoscaff_1782 | 7500 | 0.001529973 | 0.002162 | 0.004704635 | 0.005351275 |
| pseudoscaff_3111 | 12500 | 0.000162737 | 0.026983 | 0.032498672 | 0.033134027 |
| pseudoscaff_3111 | 17500 | 0 | 0.043541 | 0.045969769 | 0.043929501 |
| pseudoscaff_3111 | 22500 | 0.000937632 | 0.044779 | 0.039261408 | 0.037397347 |
| pseudoscaff_1267 | 2500 | 0 | 0.01517 | 0.020368811 | 0.019665052 |
| pseudoscaff_1267 | 7500 | 0 | 0.018748 | 0.017844003 | 0.01832084 |
| pseudoscaff_1267 | 12500 | 0.000272222 | 0.01787 | 0.011725924 | 0.012027142 |
| pseudoscaff_1267 | 17500 | 0.000160869 | 0.011312 | 0.006323066 | 0.005511626 |
| pseudoscaff_1267 | 22500 | 0 | 0.008765 | 0.009026986 | 0.010896993 |
| pseudoscaff_1267 | 27500 | 0 | 0.007371 | 0.010068231 | 0.010066476 |
| pseudoscaff_1267 | 32500 | 0 | 0.004051 | 0.004615136 | 0.004445758 |
| pseudoscaff_1267 | 37500 | 0 | 0.004016 | 0.007716194 | 0.007537854 |
| pseudoscaff_1267 | 42500 | 0.000365096 | 0.001789 | 0.003254383 | 0.003108578 |
| pseudoscaff_1267 | 47500 | 0.000116208 | 0.00105 | 0.004075073 | 0.003639757 |
| pseudoscaff_1267 | 52500 | 0 | 0.003491 | 0.003594529 | 0.002781224 |
| pseudoscaff_1267 | 57500 | 0 | 0.026341 | 0.011597376 | 0.010870637 |
| pseudoscaff_1267 | 62500 | 0 | 0.009348 | 0.009314842 | 0.009856532 |
| pseudoscaff_1267 | 67500 | 0 | 0.00749 | 0.007756305 | 0.010657293 |
| pseudoscaff_1267 | 72500 | 0 | 0.01182 | 0.005148915 | 0.006760567 |
| pseudoscaff_1267 | 82500 | 0.000205314 | 0.005211 | 0.008546023 | 0.006230027 |
| pseudoscaff_1267 | 87500 | 0.001040561 | 0.006736 | 0.019833635 | 0.014687654 |
| pseudoscaff_1267 | 92500 | 0.001864078 | 0.011201 | 0.011014852 | 0.011057919 |
| pseudoscaff_2988 | 7500 | 0 | 0.004648 | 0.007196989 | 0.006697938 |
| pseudoscaff_2988 | 12500 | 0 | 0.004985 | 0.003099318 | 0.002408443 |

| | | | | | |
|---|---|---|---|---|---|
| pseudoscaff_2988 | 17500 | 0 | 0.003783 | 0.002892744 | 0.004217157 |
| pseudoscaff_2988 | 22500 | 0.000225188 | 0.006937 | 0.007520851 | 0.006740018 |
| pseudoscaff_2988 | 27500 | 0 | 0.002227 | 0.003637004 | 0.003551723 |
| pseudoscaff_2988 | 32500 | 0 | 0.001261 | 0.003864875 | 0.003559848 |
| pseudoscaff_2988 | 37500 | 0.000080046 | 0.003398 | 0.005389095 | 0.004928662 |
| pseudoscaff_2988 | 47500 | 0.001904768 | 0.007073 | 0.004597171 | 0.003452482 |
| pseudoscaff_2988 | 57500 | 0.001710388 | 0.004392 | 0.008641803 | 0.008044778 |
| pseudoscaff_1229 | 2500 | 0.000265583 | 0.014284 | 0.004824504 | 0.006214136 |
| pseudoscaff_1229 | 7500 | 0.000345152 | 0.014566 | 0.007609693 | 0.00918282 |
| pseudoscaff_1229 | 17500 | 0 | 0.011446 | 0.006986613 | 0.008993758 |
| pseudoscaff_1229 | 22500 | 0 | 0.011275 | 0.001382348 | 0.00242066 |
| pseudoscaff_1229 | 32500 | 0.000564777 | 0.008092 | 0.002127256 | 0.002206398 |
| pseudoscaff_1229 | 37500 | 0 | 0.006307 | 0.002499737 | 0.003036305 |
| pseudoscaff_1229 | 42500 | 0 | 0.00461 | 0.001780917 | 0.002263388 |
| pseudoscaff_1229 | 47500 | 0 | 0.003088 | 0.001185512 | 0.002654762 |
| pseudoscaff_1229 | 52500 | 0.000156344 | 0.012418 | 0.001910065 | 0.003270431 |
| pseudoscaff_1229 | 57500 | 0 | 0.007526 | 0.001286905 | 0.002832621 |
| pseudoscaff_1229 | 67500 | 0 | 0.005359 | 0.001670553 | 0.00341099 |
| pseudoscaff_1229 | 72500 | 0 | 0.005139 | 0.001365428 | 0.002830215 |
| pseudoscaff_1229 | 77500 | 0 | 0.00648 | 0.001941724 | 0.002543115 |
| pseudoscaff_1229 | 82500 | 0 | 0.007473 | 0.001651752 | 0.002509804 |
| pseudoscaff_3316 | 147500 | 0 | 0.028309 | 0.030217719 | 0.029631594 |
| pseudoscaff_3399 | 2500 | 0.00017575 | 0.001801 | 0.003467785 | 0.002663162 |
| pseudoscaff_2514 | 2500 | 0.001693148 | 0.005232 | 0.004796017 | 0.005098424 |
| pseudoscaff_2514 | 12500 | 0.001451615 | 0.002889 | 0.001472824 | 0.001651372 |
| pseudoscaff_2514 | 22500 | 0.001736693 | 0.001004 | 0.004859847 | 0.008013116 |
| pseudoscaff_2514 | 27500 | 0.001872687 | 0.008483 | 0.009641816 | 0.010066324 |
| pseudoscaff_2514 | 37500 | 0.000690703 | 0.007618 | 0.008277048 | 0.011412207 |
| pseudoscaff_2514 | 42500 | 0.000448565 | 0.005717 | 0.004660188 | 0.011103178 |
| pseudoscaff_52 | 2500 | 0.00020836 | 0.003889 | 0.0087599 | 0.010569629 |
| pseudoscaff_1588 | 2500 | 0 | 0.014256 | 0.025279259 | 0.021167263 |
| pseudoscaff_1588 | 7500 | 0.000631719 | 0.015135 | 0.017219088 | 0.019160149 |
| pseudoscaff_1588 | 27500 | 0 | 0.014544 | 0.020944268 | 0.02110352 |
| pseudoscaff_1588 | 47500 | 0 | 0.00665 | 0.006952438 | 0.007317673 |
| pseudoscaff_1588 | 52500 | 0 | 0.022364 | 0.020701203 | 0.022733913 |
| pseudoscaff_1588 | 57500 | 0 | 0.009212 | 0.01141695 | 0.012060799 |
| pseudoscaff_5 | 267500 | 0.000179881 | 0.004405 | 0.000934401 | 0.000971984 |
| pseudoscaff_5 | 272500 | 0 | 0.012398 | 0.000318032 | 0.000375669 |
| pseudoscaff_5 | 277500 | 0.000101144 | 0.016596 | 0.00015696 | 0.000114345 |
| pseudoscaff_5 | 292500 | 0 | 0.003182 | 0.003645847 | 0.002428756 |
| pseudoscaff_5 | 302500 | 0.001927963 | 0.016487 | 0.005793904 | 0.006129458 |
| pseudoscaff_5 | 307500 | 0.001686073 | 0.016316 | 0.023136268 | 0.022145754 |
| pseudoscaff_5 | 317500 | 0.001895124 | 0.001084 | 0.013327653 | 0.012665148 |
| pseudoscaff_5 | 332500 | 0 | 0 | 0.000474502 | 0.000508075 |
| pseudoscaff_5 | 337500 | 0.000584717 | 0.012133 | 0.010380233 | 0.010298373 |
| pseudoscaff_5 | 342500 | 0 | 0.000802 | 0.001053108 | 0.00119052 |
| pseudoscaff_2625 | 7500 | 0.001112276 | 0.024543 | 0.023195872 | 0.022446132 |
| pseudoscaff_2625 | 12500 | 0.000111606 | 0.029934 | 0.029277752 | 0.029961009 |
| pseudoscaff_2625 | 17500 | 0 | 0.031871 | 0.025518593 | 0.026922196 |
| pseudoscaff_107 | 17500 | 0.000225593 | 0.037655 | 0.042617644 | 0.050210908 |

| | | | | | |
|---|---|---|---|---|---|
| pseudoscaff_3540 | 12500 | 0.001869792 | 0.034729 | 0.035319484 | 0.035153869 |
| pseudoscaff_3540 | 17500 | 0.001719798 | 0.030798 | 0.038407059 | 0.040699761 |
| pseudoscaff_3540 | 42500 | 0.000632035 | 0.015491 | 0.020420409 | 0.019742979 |
| pseudoscaff_3540 | 77500 | 0.000467744 | 0.028355 | 0.037262752 | 0.038986416 |
| pseudoscaff_4206 | 252500 | 0.000610031 | 0.038078 | 0.041415078 | 0.044491176 |
| pseudoscaff_4206 | 272500 | 0.000351378 | 0.005498 | 0.007115055 | 0.00739888 |
| pseudoscaff_4206 | 277500 | 0.000052035 | 0.005769 | 0.00591044 | 0.00663254 |
| pseudoscaff_4206 | 282500 | 0 | 0.011049 | 0.016605034 | 0.016036533 |
| pseudoscaff_4206 | 287500 | 0.000093519 | 0.006358 | 0.006487716 | 0.006479098 |
| pseudoscaff_4206 | 292500 | 0 | 0.004905 | 0.005791162 | 0.006004811 |
| pseudoscaff_4206 | 297500 | 0 | 0.018562 | 0.017178988 | 0.017004045 |
| pseudoscaff_2603 | 2500 | 0.001631367 | 0.005455 | 0.009023961 | 0.008703368 |
| pseudoscaff_2603 | 7500 | 0.001648635 | 0.006218 | 0.01152389 | 0.008612627 |
| pseudoscaff_1552 | 7500 | 0 | 0.022003 | 0.017666314 | 0.017195857 |
| pseudoscaff_2766 | 2500 | 0 | 0.015498 | 0.004856445 | 0.005601374 |
| pseudoscaff_3513 | 22500 | 0.000100247 | 0.008774 | 0.006477139 | 0.00606043 |
| pseudoscaff_3513 | 27500 | 0 | 0.002037 | 0.003369785 | 0.002453396 |
| pseudoscaff_3513 | 32500 | 0 | 0.005496 | 0.004461985 | 0.00382345 |
| pseudoscaff_3513 | 52500 | 0.000155586 | 0.009714 | 0.011159116 | 0.012152813 |
| pseudoscaff_3513 | 57500 | 0.000312231 | 0.017853 | 0.018310354 | 0.020271555 |
| pseudoscaff_3513 | 62500 | 0 | 0.01299 | 0.015620835 | 0.016065303 |
| pseudoscaff_3513 | 82500 | 0 | 0.028226 | 0.015952263 | 0.015027202 |
| pseudoscaff_3513 | 87500 | 0.000056725 | 0.028688 | 0.029017035 | 0.029491512 |
| pseudoscaff_3513 | 92500 | 0.000639865 | 0.032068 | 0.024676616 | 0.025390922 |
| pseudoscaff_3513 | 112500 | 0 | 0.014011 | 0.011139947 | 0.014226878 |
| pseudoscaff_3513 | 127500 | 0.000154553 | 0.023954 | 0.020097414 | 0.020329906 |
| pseudoscaff_3513 | 137500 | 0.001391056 | 0.027303 | 0.03010965 | 0.030080147 |
| pseudoscaff_3499 | 12500 | 0.000550985 | 0.001781 | 0.002536582 | 0.001755742 |
| pseudoscaff_1717 | 17500 | 0.001161457 | 0.028271 | 0.028452372 | 0.026126619 |
| pseudoscaff_4034 | 17500 | 0.000646725 | 0.00255 | 0.020759873 | 0.015724531 |
| pseudoscaff_3765 | 7500 | 0 | 0.037154 | 0.040588587 | 0.038298614 |
| pseudoscaff_2920 | 57500 | 0.000381178 | 0.03969 | 0.033264213 | 0.037277906 |
| pseudoscaff_3418 | 32500 | 0.001831521 | 0.000953 | 0.00305368 | 0.003972507 |
| pseudoscaff_3418 | 57500 | 0.001010909 | 0.001171 | 0.001188422 | 0.001174871 |
| pseudoscaff_2368 | 147500 | 0.000085434 | 0.014305 | 0.015017073 | 0.015613454 |
| pseudoscaff_2368 | 157500 | 0.000143223 | 0.025022 | 0.024561753 | 0.025449983 |
| pseudoscaff_2368 | 172500 | 0.00027559 | 0.017377 | 0.021295629 | 0.021752283 |
| pseudoscaff_2368 | 177500 | 0.000649903 | 0.012176 | 0.021614751 | 0.021510444 |
| pseudoscaff_2368 | 187500 | 0.000679224 | 0.012602 | 0.013306759 | 0.0135245 |
| pseudoscaff_2368 | 202500 | 0 | 0.016329 | 0.022539538 | 0.020807733 |
| pseudoscaff_2368 | 217500 | 0.000055606 | 0.019646 | 0.022971149 | 0.02113637 |
| pseudoscaff_2368 | 232500 | 0.000349114 | 0.019645 | 0.015421628 | 0.016957406 |
| pseudoscaff_2368 | 347500 | 0.000403855 | 0.024266 | 0.028353806 | 0.028847926 |
| pseudoscaff_1811 | 12500 | 0.001468453 | 0.016794 | 0.019309487 | 0.021739619 |
| pseudoscaff_1876 | 47500 | 0.001724835 | 0.007739 | 0.010720379 | 0.010022981 |
| pseudoscaff_905 | 27500 | 0.001399328 | 0.027675 | 0.03640679 | 0.035229175 |
| pseudoscaff_3546 | 2500 | 0.000525417 | 0.016122 | 0.025560451 | 0.025412015 |
| pseudoscaff_3546 | 7500 | 0.001000077 | 0.001603 | 0.014292518 | 0.012959918 |
| pseudoscaff_3546 | 12500 | 0.000605325 | 0.000336 | 0.008677662 | 0.008664978 |
| pseudoscaff_3546 | 17500 | 0.001175812 | 0.000205 | 0.007350117 | 0.006664952 |

| | | | | | |
|---|---|---|---|---|---|
| pseudoscaff_3546 | 22500 | 0 | 0.000535 | 0.006080483 | 0.006170906 |
| pseudoscaff_3546 | 27500 | 0 | 0.003522 | 0.025605282 | 0.02443702 |
| pseudoscaff_3546 | 42500 | 0.001524934 | 0.001628 | 0.007899981 | 0.00646696 |
| pseudoscaff_3546 | 47500 | 0 | 0.000208 | 0.008594397 | 0.007636268 |
| pseudoscaff_3546 | 52500 | 0 | 0.000861 | 0.004596734 | 0.004355556 |
| pseudoscaff_3546 | 57500 | 0.000078289 | 0.001193 | 0.009122502 | 0.010807429 |
| pseudoscaff_3546 | 67500 | 0 | 0.000592 | 0.00462535 | 0.005144364 |
| scaffold_4233 | 2500 | 0 | 0.020965 | 0.019478761 | 0.018108064 |
| scaffold_4233 | 7500 | 0 | 0.00744 | 0.009946915 | 0.007602817 |
| scaffold_13812 | 2500 | 0.000267399 | 0 | 0.000163229 | 0.000250069 |
| scaffold_706 | 27500 | 0.001665465 | 0.001783 | 0.012905279 | 0.010431357 |
| scaffold_706 | 32500 | 0.000187225 | 0.002426 | 0.002893893 | 0.004009174 |
| scaffold_706 | 42500 | 0.000096027 | 0.00097 | 0.004753665 | 0.003818873 |
| scaffold_706 | 62500 | 0 | 0.002376 | 0.002296094 | 0.001679709 |
| scaffold_706 | 67500 | 0 | 0.00045 | 0.001731458 | 0.001740386 |
| scaffold_706 | 72500 | 0.001494606 | 0.005435 | 0.003842718 | 0.00395863 |
| scaffold_706 | 77500 | 0.000985847 | 0.020817 | 0.009651636 | 0.009234381 |
| scaffold_706 | 87500 | 0.000533407 | 0.000794 | 0.001240633 | 0.001745811 |
| scaffold_706 | 92500 | 0.000156448 | 0.004644 | 0.001575926 | 0.001772133 |
| scaffold_706 | 97500 | 0.001683833 | 0.003182 | 0.002619577 | 0.003148055 |
| scaffold_8073 | 7500 | 0.00178447 | 0.002315 | 0.001868934 | 0.001748243 |
| superscaffold_31 | 7500 | 0.001961529 | 0.012893 | 0.012140279 | 0.012277152 |
| superscaffold_31 | 12500 | 0.001393088 | 0.01302 | 0.009359503 | 0.009162466 |
| scaffold_1825 | 12500 | 0.001018956 | 0.003202 | 0.002027555 | 0.003363983 |
| scaffold_281 | 27500 | 0.000839935 | 0.007953 | 0.005921505 | 0.005950248 |
| scaffold_281 | 32500 | 0.001681093 | 0.007455 | 0.006872406 | 0.007235603 |
| scaffold_281 | 37500 | 0 | 0 | 0.001917674 | 0.001865513 |
| scaffold_281 | 62500 | 0.001141025 | 0.000503 | 0.005770509 | 0.005503795 |
| scaffold_281 | 157500 | 0.001930249 | 0.008151 | 0.00463987 | 0.00458145 |
| scaffold_281 | 172500 | 0.000381171 | 0.002448 | 0.001565561 | 0.001499294 |
| scaffold_18109 | 2500 | 0.00081998 | 0.005733 | 0.017640268 | 0.016974844 |
| scaffold_1006 | 57500 | 0 | 0.011577 | 0.01331074 | 0.013917449 |
| scaffold_1006 | 62500 | 0.000412812 | 0.025833 | 0.017386261 | 0.018963018 |
| scaffold_16942 | 2500 | 0 | 0.018153 | 0.009799672 | 0.014045426 |
| scaffold_22130 | 2500 | 0.000833517 | 0.00127 | 0.003800057 | 0.004639689 |
| superscaffold_1045 | 47500 | 0.001498965 | 0.017835 | 0.017563737 | 0.019374154 |
| scaffold_848 | 57500 | 0 | 0.003053 | 0.004111401 | 0.004150192 |
| scaffold_848 | 72500 | 0.001330884 | 0.018146 | 0.026384892 | 0.026041062 |
| scaffold_1124 | 72500 | 0.000070622 | 0.017016 | 0.014059671 | 0.014388901 |
| scaffold_207 | 92500 | 0 | 0.035067 | 0.032147769 | 0.032972877 |
| superscaffold_1025 | 47500 | 0 | 0.025559 | 0.030853436 | 0.033151845 |
| superscaffold_1025 | 62500 | 0.000529554 | 0.057479 | 0.049521178 | 0.048915686 |
| scaffold_21407 | 2500 | 0.0017757 | 0.005066 | 0.005148324 | 0.004405857 |
| scaffold_9080 | 2500 | 0 | 0.003987 | 0.005784959 | 0.008460965 |
| scaffold_9080 | 7500 | 0 | 0.002607 | 0.00371189 | 0.003851501 |
| scaffold_5750 | 7500 | 0.000291866 | 0.007862 | 0.009955038 | 0.010560741 |
| scaffold_17216 | 2500 | 0 | 0.029469 | 0.007778886 | 0.007140463 |
| superscaffold_630 | 87500 | 0.000127616 | 0.014729 | 0.024179289 | 0.022182346 |
| superscaffold_630 | 92500 | 0.001483181 | 0.021872 | 0.028531958 | 0.026792544 |
| scaffold_4615 | 7500 | 0.001677457 | 0.002108 | 0.002975549 | 0.004202804 |

| | | | | | |
|---|---|---|---|---|---|
| superscaffold_147 | 2500 | 0.000195437 | 0.011119 | 0.016425382 | 0.017298856 |
| superscaffold_219 | 152500 | 0.000931047 | 0.015139 | 0.020040726 | 0.020679844 |
| superscaffold_219 | 162500 | 0.000388377 | 0.027705 | 0.024145499 | 0.025790106 |
| superscaffold_219 | 167500 | 0.000102268 | 0.020473 | 0.019839016 | 0.019355388 |
| superscaffold_219 | 172500 | 0.00009097 | 0.028636 | 0.032405987 | 0.031322579 |
| superscaffold_219 | 177500 | 0 | 0.02535 | 0.027244434 | 0.027703685 |
| superscaffold_219 | 182500 | 0.000125914 | 0.020447 | 0.02298508 | 0.02297619 |
| superscaffold_219 | 187500 | 0.000156283 | 0.02079 | 0.030233722 | 0.028157965 |
| superscaffold_219 | 192500 | 0.000157 | 0.017076 | 0.01709757 | 0.01937273 |
| superscaffold_219 | 202500 | 0 | 0.021927 | 0.018303436 | 0.018252229 |
| superscaffold_219 | 207500 | 0.000052809 | 0.01717 | 0.018664766 | 0.018791166 |
| superscaffold_219 | 212500 | 0.000748626 | 0.026194 | 0.020741574 | 0.021944546 |
| superscaffold_219 | 217500 | 0 | 0.005889 | 0.009925194 | 0.008728464 |
| superscaffold_219 | 252500 | 0.001616077 | 0.029744 | 0.029011984 | 0.030182772 |
| superscaffold_219 | 277500 | 0.001788405 | 0.014845 | 0.019087371 | 0.016182111 |
| superscaffold_219 | 282500 | 0.000518383 | 0.004251 | 0.008922982 | 0.007840105 |
| superscaffold_219 | 287500 | 0.000065042 | 0.007039 | 0.005937727 | 0.006321507 |
| superscaffold_219 | 292500 | 0.000337141 | 0.019654 | 0.022949398 | 0.023404836 |
| superscaffold_219 | 297500 | 0.001019328 | 0.018137 | 0.018884635 | 0.016970587 |
| superscaffold_219 | 312500 | 0.001199003 | 0.037762 | 0.039398839 | 0.038249613 |
| scaffold_8176 | 2500 | 0.000945453 | 0.006279 | 0.007472912 | 0.007378519 |
| scaffold_16962 | 2500 | 0.001408881 | 0.044906 | 0.039517276 | 0.0408369 |
| scaffold_18140 | 2500 | 0.000200148 | 0.015787 | 0.015570176 | 0.013575946 |
| scaffold_15006 | 2500 | 0.00107101 | 0.021221 | 0.005223336 | 0.005835797 |
| scaffold_8040 | 7500 | 0.001098436 | 0.036762 | 0.045739891 | 0.045524759 |
| superscaffold_58 | 127500 | 0.000437947 | 0.02354 | 0.021059019 | 0.021791485 |
| superscaffold_58 | 262500 | 0.00182561 | 0.030551 | 0.032997593 | 0.032752281 |
| superscaffold_58 | 277500 | 0.001607371 | 0.032542 | 0.030214704 | 0.033287303 |
| superscaffold_58 | 282500 | 0.00022907 | 0.015465 | 0.024973718 | 0.02611131 |
| scaffold_4288 | 2500 | 0.000891325 | 0.000789 | 0.00138177 | 0.001890899 |
| scaffold_4509 | 2500 | 0 | 0.015037 | 0.01164378 | 0.011203947 |
| scaffold_4509 | 7500 | 0.000822968 | 0.00996 | 0.017236044 | 0.015625236 |
| scaffold_4509 | 12500 | 0.000200221 | 0.021501 | 0.024531838 | 0.023007177 |
| scaffold_4509 | 17500 | 0.000110228 | 0.024531 | 0.027646894 | 0.025784115 |
| scaffold_17641 | 2500 | 0 | 0.007967 | 0.023388794 | 0.026906458 |
| superscaffold_528 | 47500 | 0.00112565 | 0.001756 | 0.002072235 | 0.002438649 |
| superscaffold_528 | 52500 | 0.00081869 | 0.00904 | 0.013873902 | 0.011091381 |
| superscaffold_528 | 57500 | 0.001553859 | 0.014382 | 0.019377478 | 0.015992502 |
| superscaffold_500 | 7500 | 0.000567328 | 0.015028 | 0.022889059 | 0.020848845 |
| superscaffold_500 | 12500 | 0.001416609 | 0.017255 | 0.018535237 | 0.019157376 |
| superscaffold_500 | 17500 | 0 | 0.022004 | 0.026658049 | 0.026329812 |
| superscaffold_500 | 27500 | 0.00068121 | 0.016257 | 0.01641517 | 0.017071361 |
| superscaffold_500 | 42500 | 0.001765171 | 0.032046 | 0.043634813 | 0.042210734 |
| superscaffold_500 | 52500 | 0.001095186 | 0.030201 | 0.032625953 | 0.031353083 |
| superscaffold_500 | 57500 | 0 | 0.010791 | 0.014079981 | 0.014960569 |
| superscaffold_500 | 62500 | 0.000775613 | 0.01563 | 0.017521028 | 0.018487015 |
| superscaffold_500 | 77500 | 0 | 0.013177 | 0.017464004 | 0.017261295 |
| superscaffold_500 | 82500 | 0.001831863 | 0.016806 | 0.019948077 | 0.019126502 |
| superscaffold_500 | 87500 | 0.000335764 | 0.020717 | 0.020600079 | 0.019599298 |
| superscaffold_500 | 92500 | 0.00046768 | 0.013499 | 0.014068878 | 0.014358246 |

| | | | | | |
|---|---|---|---|---|---|
| superscaffold_500 | 97500 | 0.001348203 | 0.023183 | 0.0239782 | 0.023878967 |
| scaffold_1 | 212500 | 0.001696977 | 0.003201 | 0.010120698 | 0.012072869 |
| scaffold_7819 | 7500 | 0 | 0.027886 | 0.037146129 | 0.036105707 |
| scaffold_14552 | 2500 | 0 | 0.03156 | 0.033730653 | 0.035054637 |
| scaffold_16823 | 2500 | 0.001935397 | 0.02071 | 0.016425711 | 0.015859499 |
| superscaffold_725 | 12500 | 0.001832391 | 0.01168 | 0.011940526 | 0.00983968 |
| superscaffold_725 | 32500 | 0.001628421 | 0.003116 | 0.002287274 | 0.001818454 |
| scaffold_3 | 502500 | 0.001038656 | 0.002421 | 0.013177638 | 0.011327963 |
| scaffold_21179 | 2500 | 0.001782586 | 0.010122 | 0.009582388 | 0.01028674 |
| scaffold_9015 | 7500 | 0.000655667 | 0.020438 | 0.023432988 | 0.027594973 |
| scaffold_13632 | 2500 | 0.000745982 | 0.014269 | 0.024124855 | 0.022442474 |
| superscaffold_990 | 37500 | 0.000331634 | 0.006076 | 0.011782632 | 0.011622575 |
| scaffold_622 | 7500 | 0.000158434 | 0.000472 | 0.000645932 | 0.000332377 |
| scaffold_622 | 22500 | 0.000113555 | 0.005472 | 0.007856485 | 0.008295685 |
| scaffold_622 | 27500 | 0 | 0.002207 | 0.000660876 | 0.000821987 |
| scaffold_622 | 32500 | 0 | 0.004436 | 0.003563824 | 0.002645304 |
| scaffold_622 | 77500 | 0.000780787 | 0.001595 | 0.002935251 | 0.002825456 |
| scaffold_622 | 87500 | 0.000899644 | 0.004612 | 0.00750736 | 0.005195276 |
| scaffold_298 | 57500 | 0 | 0.024876 | 0.024966533 | 0.026302837 |
| scaffold_10708 | 2500 | 0.000324357 | 0.000115 | 0.004049986 | 0.00404339 |
| scaffold_16997 | 2500 | 0.000071293 | 0.04139 | 0.03871741 | 0.038642913 |
| scaffold_5829 | 2500 | 0.00058268 | 0.006645 | 0.021920505 | 0.019099773 |
| scaffold_168 | 142500 | 0.000420788 | 0.027327 | 0.025918284 | 0.026804314 |
| scaffold_168 | 152500 | 0.000064535 | 0.021695 | 0.02124113 | 0.020816115 |
| scaffold_168 | 157500 | 0.001258559 | 0.01703 | 0.022671254 | 0.022818327 |
| scaffold_2965 | 17500 | 0.001631887 | 0.004762 | 0.005550393 | 0.005304204 |
| superscaffold_1014 | 12500 | 0.000120297 | 0.023681 | 0.026791737 | 0.028187733 |
| superscaffold_1014 | 82500 | 0.001518387 | 0.010537 | 0.010632734 | 0.011519573 |
| superscaffold_1014 | 142500 | 0.00020357 | 0.004616 | 0.006406303 | 0.006036121 |
| superscaffold_1014 | 147500 | 0.001739473 | 0.008198 | 0.005725393 | 0.006308584 |
| scaffold_3363 | 2500 | 0 | 0.012297 | 0.012613452 | 0.012587374 |
| scaffold_3363 | 7500 | 0.000197517 | 0.01997 | 0.0147599 | 0.014024709 |
| scaffold_3363 | 12500 | 0 | 0.018145 | 0.020278508 | 0.020407727 |
| scaffold_3363 | 17500 | 0.000100825 | 0.029511 | 0.031679248 | 0.030858117 |
| scaffold_3363 | 22500 | 0.000055798 | 0.028187 | 0.033312352 | 0.029710652 |
| scaffold_3363 | 27500 | 0 | 0.024098 | 0.020486391 | 0.018599746 |
| scaffold_6263 | 2500 | 0 | 0.047939 | 0.042755889 | 0.043582394 |
| scaffold_6263 | 7500 | 0 | 0.035553 | 0.037366774 | 0.040878573 |
| scaffold_6263 | 12500 | 0 | 0.00194 | 0.003993603 | 0.00382151 |
| scaffold_4776 | 7500 | 0.000136072 | 0.000939 | 0.001768642 | 0.000532529 |
| scaffold_4534 | 17500 | 0.000231071 | 0.002876 | 0.00319708 | 0.003338179 |
