## Supplementary figures and images for "There is more than chitin synthase in insect resistance to benzoylureas: Molecular markers associated with teflubenzuron resistance in *Spodoptera frugiperda*"

### Figure S4

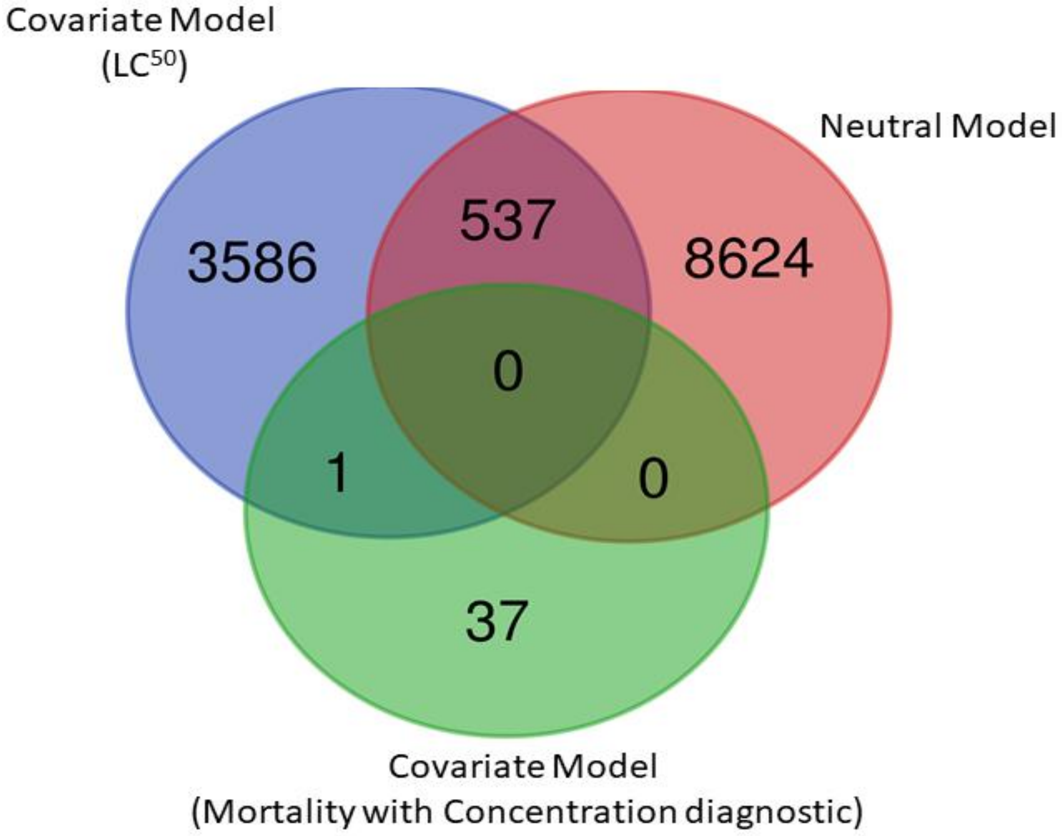


Figure S4. Venn diagram representing significant SNP from GWAS by Baypass.

### Figure S5

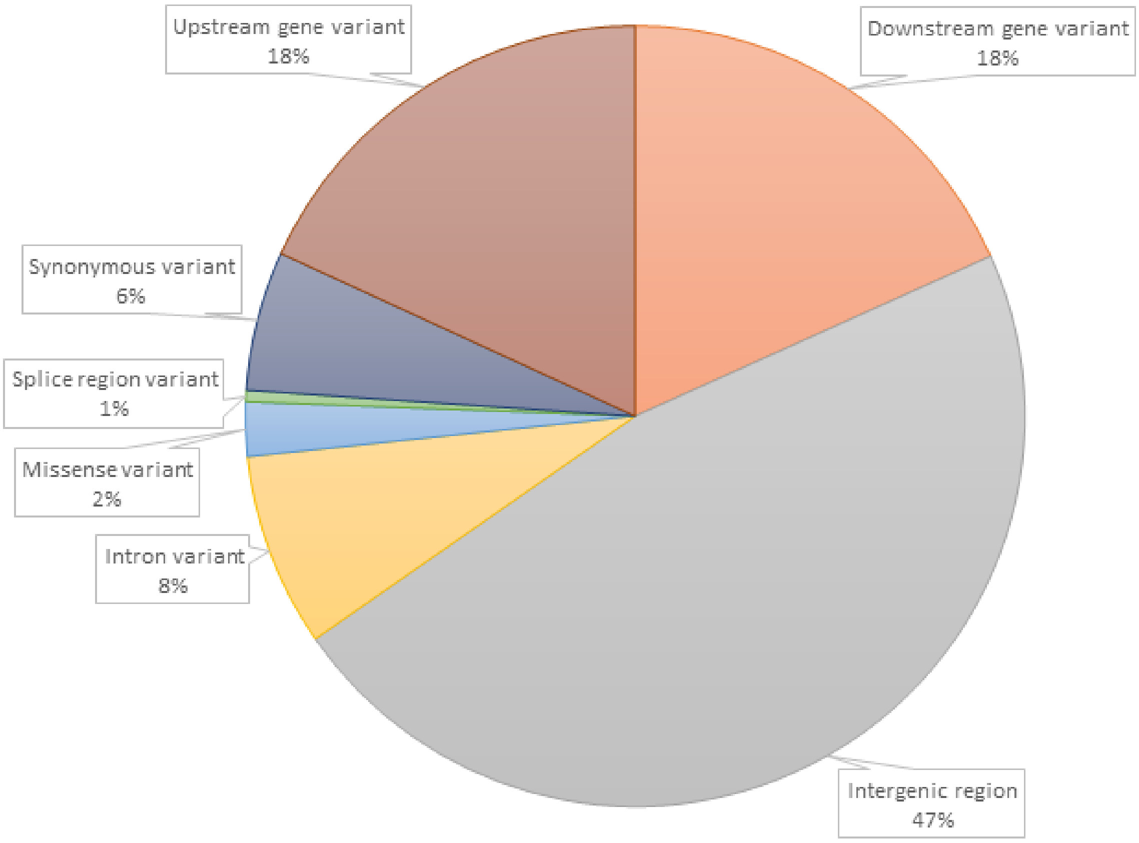


Figure S5. Distribution of SNPs under selection based on regonig regions.
