## Supplementary material for "There is more than chitin synthase in insect resistance to benzoylureas: Molecular markers associated with teflubenzuron resistance in *Spodoptera frugiperda*": Figure S2

Figure S2 - Pairwise FST estimates (ANOVA).

| | **Tef-rr** | **BC-Control** | **Sf-ss** | **BC-Treated** |
|---|---|---|---|---|
| **Tef-rr** | 0.000000 | | | |
| **BC-Control** | 0.072168 | 0.000000 | | |
| **Sf-ss** | 0.375816 | 0.20461365 | 0.00000 | |
| **BC-Treated** | 0.070699 | -0.01454088 | 0.211865 | 0.000000 |

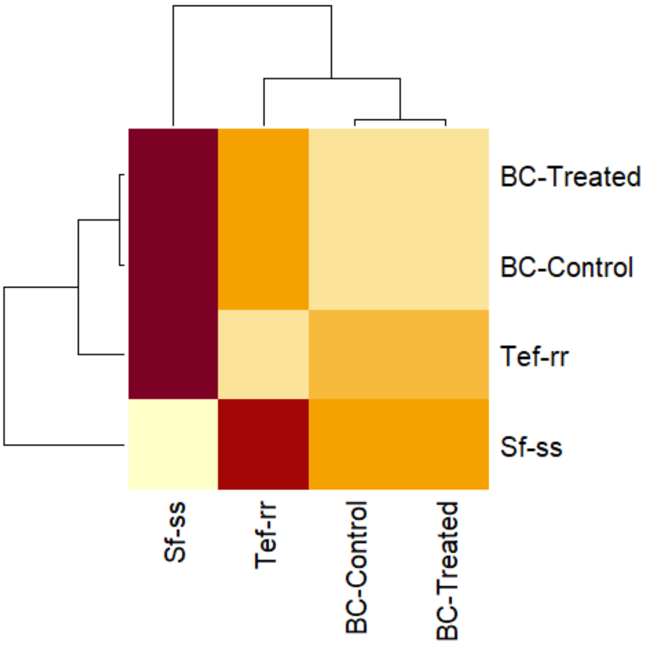


Fig. 1 Heatmap FST pairware
